# Genome-enabled analysis of population dynamics and virulence associated loci in the oat crown rust fungus *Puccinia coronata* f. sp. *avenae*

**DOI:** 10.1101/2023.09.18.557855

**Authors:** Tim C. Hewitt, Eva C. Henningsen, Danilo Pereira, Kerensa McElroy, Eric S. Nazareno, Sheshanka Dugyala, Hoa Nguyen-Phuc, Feng Li, Marisa E. Miller, Botma Visser, Zacharias A. Pretorius, Willem H.P. Boshoff, Jana Sperschneider, Eva H. Stukenbrock, Shahryar F. Kianian, Peter N. Dodds, Melania Figueroa

**Affiliations:** Commonwealth Scientific and Industrial Research Organisation, Agriculture and Food, Canberra, ACT 2600, Australia; Christian Albrechts University of Kiel and Max Planck Institute of Evolutionary Biology, Kiel and Ploen, Germany; Department of Plant Pathology, University of Minnesota, St. Paul, MN 55108, USA; Department of Plant Sciences, University of the Free State, Bloemfontein 9300, South Africa; USDA-ARS Cereal Disease Laboratory, St. Paul, MN 55108, U.S.A

## Abstract

*Puccinia coronata* f. sp. *avenae* (*Pca*) is an important fungal pathogen causing crown rust that impacts oat production worldwide. Genetic resistance for crop protection against *Pca* is often overcome by the rapid virulence evolution of the pathogen. This study investigated the factors shaping adaptive evolution of *Pca* using pathogen populations from distinct geographic regions within the USA and South Africa (SA). Phenotypic and genome-wide sequencing data of these diverse *Pca* collections, including 217 isolates, uncovered phylogenetic relationships and established distinct genetic composition between populations from northern and southern regions from the USA and SA. The population dynamics of *Pca* involve a bidirectional movement of inoculum between northern and southern regions of the USA and contributions from clonality and sexuality. The population from SA is solely clonal. A genome-wide association study (GWAS) employing a haplotype-resolved *Pca* reference genome was used to define eleven virulence-associated loci corresponding to twenty-five oat differential lines. These regions were screened to determine candidate *Avr* effector genes. Overall, the GWAS results allowed us to identify the underlying genetic traits controlling pathogen recognition in an oat differential set used in the USA to assign pathogen races (pathotypes). Key GWAS findings support complex genetic interactions in several oat lines suggesting allelism among resistance genes or redundancy of genes included in the differential set, multiple resistance genes recognising genetically linked *Avr* effector genes, or potentially epistatic relationships. A careful evaluation of the composition of the oat differential set accompanied by the development or implementation of molecular markers is recommended.

## INTRODUCTION

*Puccinia coronata* f. sp. *avenae* (*Pca*) is an important foliar pathogen that causes oat crown rust, which impacts oat production around the world. (Nazareno et al., 2018). An effective method to control rust diseases is the use of genetic resistance in crops through the incorporation of resistance (*R*) genes (Ellis et al., 2014). Most *R* genes encode immune receptors that can recognise specific secreted proteins from the pathogen, known as avirulence (Avr) effectors (Dodds and Rathjen, 2010) Such a recognition event is essential to mount an immune response that prevents the pathogen’s growth in the plant. Despite the potential benefits of genetic resistance in agriculture, the use of *R* genes to manage *Pca* infections in the field has not been as effective as needed given that most released oat *R* genes have shown limited durability against crown rust (Nazareno et al., 2018; Figueroa et al., 2020). Sexual recombination, random (sequential) mutation and somatic hybridisation are some of mechanisms that allow rust pathogens such as *Pca* to alter their genetic make-up and gain virulence in otherwise resistant cultivars (Möller and Stukenbrock, 2017; Figueroa et al., 2020; Duplessis et al., 2021).

*Pca* shares a similar life cycle with other *Puccinia* species that infect cereals. This involves alternation between an asexual cereal infection phase mediated by dikaryotic urediniospores (containing two different haploid nuclei), and a sexual phase that occurs on an alternate host. Thus, populations of cereal rust fungi can be highly sexual when the alternate host is present, but in its absence, the asexual phase can persist indefinitely giving rise to long-lived clonal populations (Figueroa et al., 2020). For instance, populations of wheat stem and leaf rust fungi (*Puccinia graminis* f. sp. *tritici* (*Pgt*) and *Puccinia triticina* (*Pt*), respectively) are predominately clonal due to the absence of the alternate sexual hosts (*Berberis* spp. and *Thalictrum spp.*, respectively) in most parts of the world (Bolton et al., 2008; Saunders et al., 2019; Patpour et al., 2022). In the northern USA, the oat crown rust fungus is capable of both sexual and asexual (clonal) reproduction due to the wide prevalence of the alternate host common buckthorn (*Rhamnus cathartica*), which permits the sexual life cycle to occur on a seasonal basis (Nazareno et al., 2018). However, buckthorn is absent from southern USA oat growing regions where asexual reproduction is predicted to play a major role (Rawlins et al., 2018). In some geographic regions the epidemiology of *Pca* is also impacted by the presence of wild oats near cropping fields, which can also facilitate asexual reproduction. Ongoing pathology surveys in the USA have found an extreme diversity of virulence phenotypes in the *Pca* population (Carson, 2011). The Rust Surveillance Annual Surveys assign race pathotypes to *Pca* isolates based on infection phenotypes on the North American oat differential set using 4-letter and 10-letter codes corresponding to virulence scores on 16 and 40 oat differential lines respectively (Chong et al., 2000). For example, between 2006 and 2009, 201 races were found among the 357 isolates from the spring oat region of the north-central USA, and 140 races were found among 214 isolates from the southern winter oat region (Carson, 2011).

Over a hundred *R* genes against oat crown rust (commonly designated as *Pc* genes) have been postulated (Nazareno et al., 2018). However, no *Pc* gene has yet been cloned and the limited molecular and genetic data for most resistance sources makes it difficult to assess whether they contain unique *Pc* genes or combinations of several *Pc* genes. The complex rearrangements observed between the sequenced oat genomes (PepsiCo, 2021; Kamal et al., 2022; Peng et al., 2022) mean that orthologous *Pc* genes may not always occur in the same location, compounding the difficulties in assigning unique *Pc* gene designations. Consequently, the limited durability of introduced *Pc* genes in the field may be partially caused by reintroduction of pre-existing genes or their allelic variants in breeding programs. Members of differential sets to complete race assignments are often selected to represent unique *R* genes; however, in the absence of genetic characterisation and molecular data of *Pc* genes it has been noted various members of oat differential sets could include the same or alleles of the same *Pc* gene (Miller et al., 2020). No *Avr* genes have yet been isolated from *Pca*. Until recently, the complex dikaryotic genomes of rust fungi have hampered genetic analyses of virulence genes, with the only Avr effectors isolated from cereal rusts so far being from the wheat stem rust pathogen (*Pgt*) using mutational and high throughput screening approaches (Chen et al., 2017; Salcedo et al., 2017; Upadhyaya et al., 2021; Arndell et al., 2023). Genome Wide Association Studies (GWAS) are emerging as a powerful approach to identify *Avr* loci in plant pathogens (Gao et al., 2016; Zhong et al., 2017; Martin et al., 2020; Kariyawasam et al., 2022; Kloppe et al., 2023), but require highly complete genome references and population data from sexually reproducing populations. Advances in sequencing technologies and computational pipelines for haplotype phasing (Li et al., 2019; Duan et al., 2022; Sperschneider et al., 2022) have created new opportunities for applying such genome wide approaches . Miller et al. (2018) generated the first partially haplotype-separated genome references for two *Pca* isolates, 12SD80 and 12NC29. These resources facilitated a population genomics study of a limited set of USA *Pca* isolates from 1990 and 2015, finding a significant shift in population genetics and virulence over this time (Miller et al., 2020). A GWAS of this population also identified seven genomic regions associated with avirulence phenotypes on fifteen *Pc* genes, and indicated for the first time that some *Pc* genes may detect *Avr* genes at similar genomic locations. However, the 12SD80 and 12NC29 genome assemblies are quite fragmented and many of the *Pc* genes elicited association peaks across multiple assembled contigs whose relative positions in the *Pca* genome were not clear. We recently developed a complete chromosome-level and nuclear-phased reference genome of *Pca* (Pca203) (Henningsen et al., 2022), and here we use this resource to expand on the initial findings reported by Miller et al. (2020) through analysis of a much larger population of USA *Pca* isolates. This reveals a primary role for local sexual reproduction in the northern USA population and extensive migration between southern and northern regions. Inclusion of the SA isolates allowed us to investigate pathotype diversity in the absence of the sexual phase, demonstrating likely single mutation events leading to multiple virulence gains. Our GWAS, using a larger *Pca* set with the Pca203 genome reference, identified a total of 11 <u>v</u>irulence-associated <u>g</u>enomic intervals (VGIs) associated with virulence phenotypes on 25 oat differential lines postulated to carry different *Pc* genes. Multiple oat lines have several corresponding VGIs, suggesting the presence of multiple *Pc* genes, many of which overlap between different oat differential lines. These results allowed us to identify *Avr* effector candidates from these regions for future testing and validation and may also expedite the isolation of *Pc* genes in the host.

## RESULTS

### *Pca* populations exhibit increased virulence in geographic regions where the sexual host is present

To assess pathotype (race) variation in the USA *Pca* population, we compared virulence scores for 185 isolates on the set of 40 oat resistance differential lines routinely used in the USA (Nazareno et al., 2018; Chong et al., 2000) (**Supp. Data S1**). These include 30 isolates collected in 1990 and 30 from 2015 that were previously characterised (Miller et al., 2020), the three reference isolates Pca203, 12SD80 and 12NC29 (Henningsen et al., 2022; Miller et al., 2018), and an additional 122 isolates collected from 2015 to 2018. Of this total set, 93 isolates were sampled from the Minnesota Matt Moore Buckthorn Nursery either as aeciospores directly from buckthorn plants (46 isolates) or as urediniospores from oats growing in the nursery and infected from the adjacent buckthorn sources (47 isolates). A total of 147 unique races were detected out of 185 isolates.

To assess recent regional diversity of *Pca*, we focussed on isolates collected from 2015-2018, since a substantial virulence shift was documented post 1990 (Miller et al., 2020). Consistent with their recent sexual recombination history, all 93 of the buckthorn nursery isolates had a unique race. Field derived isolates were also highly diverse, with 65 races detected amongst 69 isolates. However, the buckthorn nursery isolates showed a significantly higher frequency of virulence phenotypes compared to field isolates (Wilcoxon rank sum test; *p*=6.795×10^-10^) (**Supp. Fig. S1**). Amongst the field-derived samples, pathotype diversity was similarly high in northern states with prevalent buckthorn (30 races/31 isolates) and southern states where buckthorn is rare or absent (25 races/26 isolates). However, there was also a higher frequency of virulence phenotypes in the northern versus southern states (*p*=1.673×10^-6^, Wilcoxon rank sum test) (**Supp. Fig. S2**). We also observed differences in virulence distributions on individual oat differential lines between these populations. Nursery-derived isolates showed significantly higher virulence scores (*p*<0.01, Wilcoxon rank sum test) compared to field isolates on the differential lines Pc91, HiFi, Pc48, Pc52, Pc68, Pc53, Pc50, Stainless, Belle, TAM-O-405, Pc96, Pc94, and Leggett (**Fig. 1A**). The nursery-derived isolates displayed lower virulence on the Pc67 and Pc63 lines than field-derived isolates. Similarly, the northern isolates showed higher virulence (*p*<0.01, Wilcoxon rank sum test) compared to southern isolates on Pc91, HiFi, Pc48, Pc52, Pc68, Pc50, Pc63, Pc45 and IAB605Xsel lines (**Fig. 1B**).

**Figure 1.**
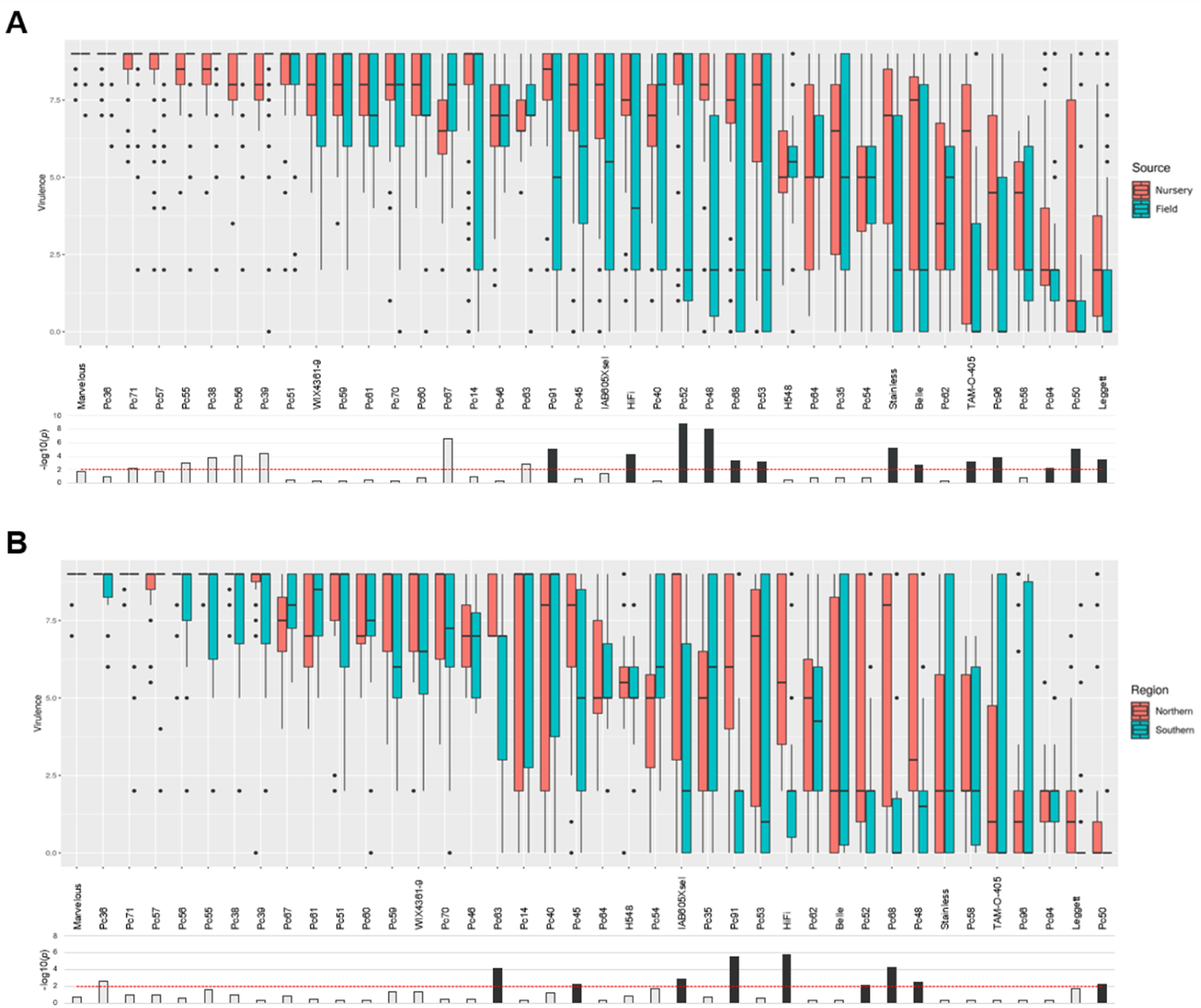
Boxplots of virulence scores on each differential comparing isolate groups. **(A)** Buckthorn nursery (red bars) versus field (blue bars) *Pca* isolates. **(B)** Northern (red bars) versus southern (blue bars) *Pca* isolates. Black dots represent outliers. Plots ordered from highest to lowest mean virulence. Bottom panels display bar graphs of –log10 of *p*-values from two-tailed Wilcoxon rank sum test of difference between groups. The red, dashed lines indicate a significance threshold of *p*=0.01. Bars with black fill mark differentials on which the virulence distribution is significantly higher in the nursery or northern group.

In contrast to the USA *Pca* population, we only recorded six different races in a set of 32 *Pca* isolates from SA (**Supp. Data S1**; **Supp. Fig. S3** and Boshoff et al., 2020). No virulence was recorded on differential lines Pc36, Pc50, Pc56, Pc60, Pc62, Pc64, Pc68, Pc91, Pc94, Pc96, Pc-H548 and Pc-WIX 1,2 for which virulence commonly occurs in the USA. This lower virulence prevalence and variation of South African isolates is consistent with the absence of buckthorn in this country limiting *Pca* to asexual reproduction.

### Structure and diversity of *Pca* populations varies between geographic regions

We generated Illumina whole-genome DNA sequencing data for all 154 new *Pca* isolates and combined this with existing sequence data for 63 isolates (Miller et al., 2018, 2020; Henningsen et al., 2022) to examine population genetic diversity. Reads were aligned to the 12SD80 reference assembly for variant calling. All samples showed a genome-wide average of at least 30X read coverage (**Supp. Fig. S4**) with normal distributions of allele frequencies (**Supp. Fig. S5**). A neighbour-net network generated from 922.1K biallelic SNPs (**Fig. 2**) showed that the USA population is dominated by a large, unstructured reticulated grouping consistent with a history of sexual recombination. A pairwise homoplasy index (PHI) test on this group strongly supported the contribution of recombination (*p* < 10^-12^) to the diversity of the population. There was also a single distinct outgroup of USA isolates, mainly from 1990, that was divergent from the main group. The SA population formed another more divergent outgroup that was very tightly clustered and a PHI test did not provide strong support for a contribution of recombination to genetic diversity (*p*=0.0464). Interestingly, the three wild oat derived isolates formed a separate cluster in this group.

**Figure 2.**
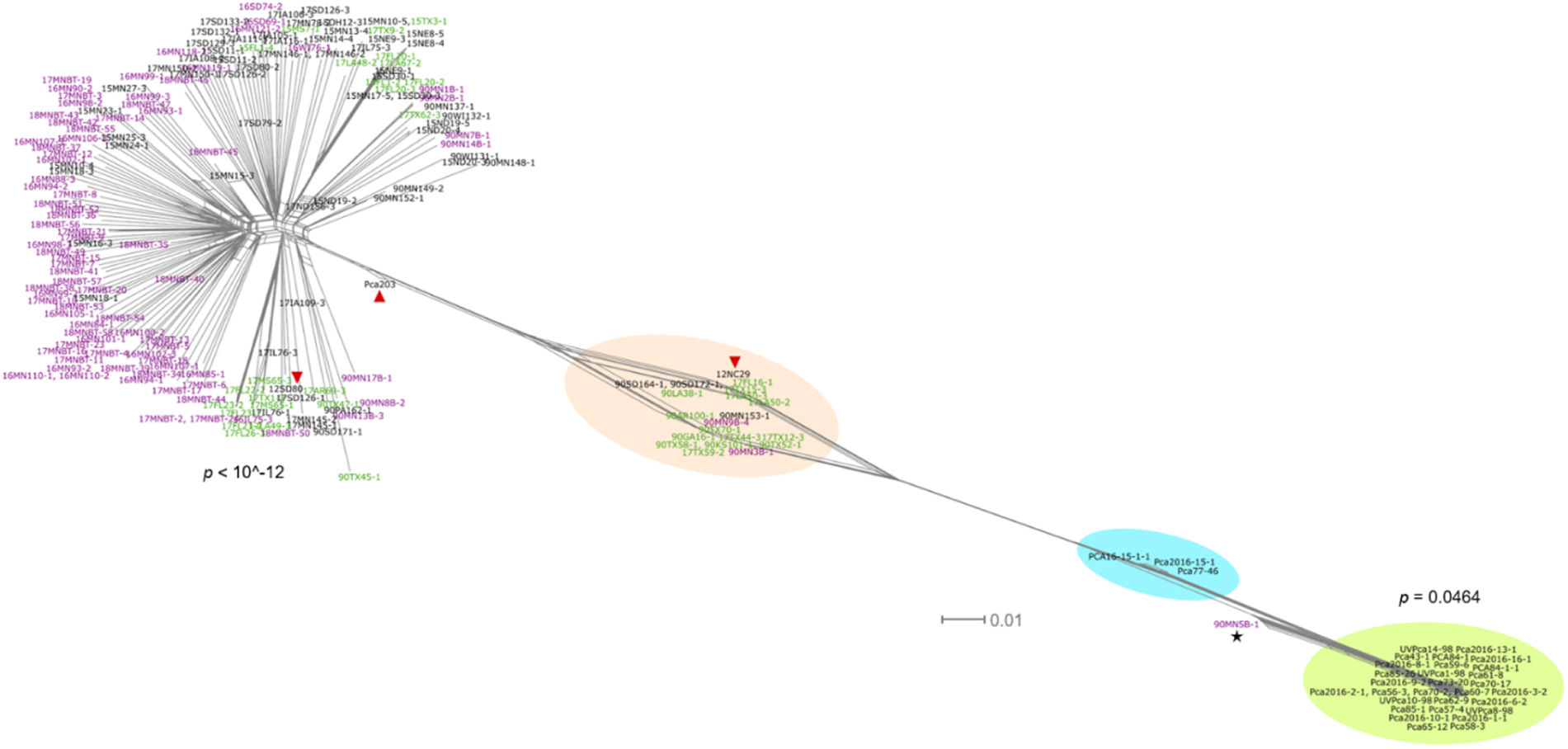
Neighbour-net graph of *Pca* isolates from the USA and South Africa. Network was generated based on 922,125 biallelic SNPs called against the 12SD80 genome. Groups of interest coloured according to legend inset. Outlying USA isolate 90MN5B-1 is indicated with a black star. Scale bar indicates nucleotide substitutions per site. *p*-values from PHI tests for recombination for the main USA population and the South African population from cultivated oats are shown.

These overall relationships were also evident in a maximum likelihood (ML) phylogenetic tree (**Fig. 3**), where the sexual recombination history in the USA population is reflected in the low bootstrap support values for most nodes, especially among northern and nursery derived isolates. However, there are several clonal groups evident (A to E) that consist of isolates separated by extremely short branch lengths connected by nodes with high bootstrap values. These clonal lineages include mostly southern USA derived isolates, consistent with absence of the sexual host in those regions (Miller et al., 2020). However, the presence of both northern and southern isolates in some of these clonal groups (B, D and E) suggests migration of rust isolates between geographic regions. Again, the SA population is divergent from the USA population and falls into just two clonal groups, one consisting of the three isolates collected from wild oats and the other containing all 29 isolates from cultivated oat. The latter group comprises isolates collected between 1998 and 2018 from a broad geographical region, which is consistent with an asexual population dominated by a single clonal lineage in South Africa. Interestingly, a single USA isolate 90MN5B-1 collected from the buckthorn nursery in 1990 appears closely related to the SA population in both the neighbour-net network and phylogenetic analyses, suggesting it may be part of a globally dispersed lineage. Genotypic variation was very high in the greater USA population (*n*=165), with variation at 821,833 out of 831,051 SNP sites, and the USA outgroup (*n*=20) with variation at 652,504 out of 756,980 SNP sites. In contrast, only 47,369 out of 548,091 SNP sites were variable within the SA cultivated-oat-derived population. Again, this is consistent with the contrasting clonal and sexual reproduction of these two populations.

**Figure 3.**
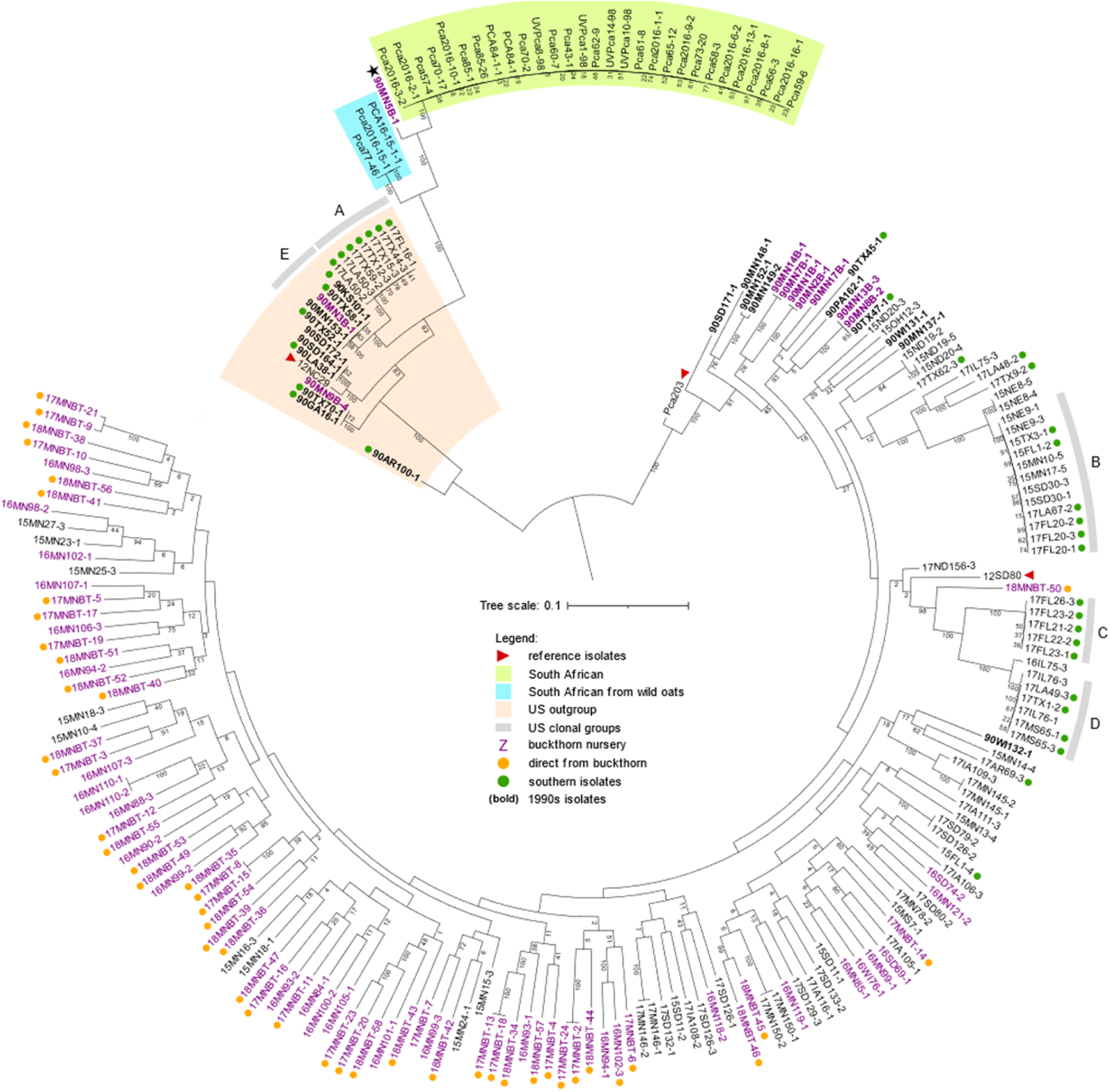
Maximum likelihood phylogenetic tree of *Pca* isolates from the USA and South Africa. The phylogeny is mid-rooted and based on 922,125 SNP variants called against the 12SD80 genome reference with support values from 500 bootstraps shown below branches. Scale bar indicates nucleotide substitutions per site. Groups of interest are coloured according to legend inset. Outlying USA isolate 90MN5B-1 is indicated with a black asterisk. Clonal groups with > 3 members are labelled A, B, C, D and E.

To determine the population structure of the pathogen population, the SNP dataset was pruned to a set of 132,571 unlinked SNPs. A principal component analysis (PCA) (**Supp. Fig. S6A)** showed a clear differentiation of the cultivated oat derived South African isolates from all other isolates, but no other clear groupings. Cluster membership analysis (**Fig. 4A, Supp. Fig. S6B**) revealed that the nursery-derived isolates displayed a high degree of admixture, which is consistent with a history of sexual recombination. Some USA field-derived isolates showed single cluster membership, consistent with the presence of asexually reproducing clonal groups, yet admixture is still evident in most isolates, especially those originating from northern regions. Most clusters are detected in both northern and southern USA *Pca* isolates, except for clusters 4 and 5, which comprise clonal groups A and C (**Fig. 3**) which each show homogeneous cluster memberships. This supports the hypothesis that divergent lineages persist asexually in the southern USA. The SA isolates from cultivated oats display no admixture and belong to a single cluster (1), while the three isolates from wild oats showed admixture between cluster 1 and 4, suggesting a relationship with the southern USA clonal group A. Overall, these population analyses all support that the USA *Pca* population is predominantly sexual with some clonal lineages persisting over time particularly in the southern regions, in contrast to the SA population, which is entirely clonal and consists of a single lineage.

**Figure 4.**
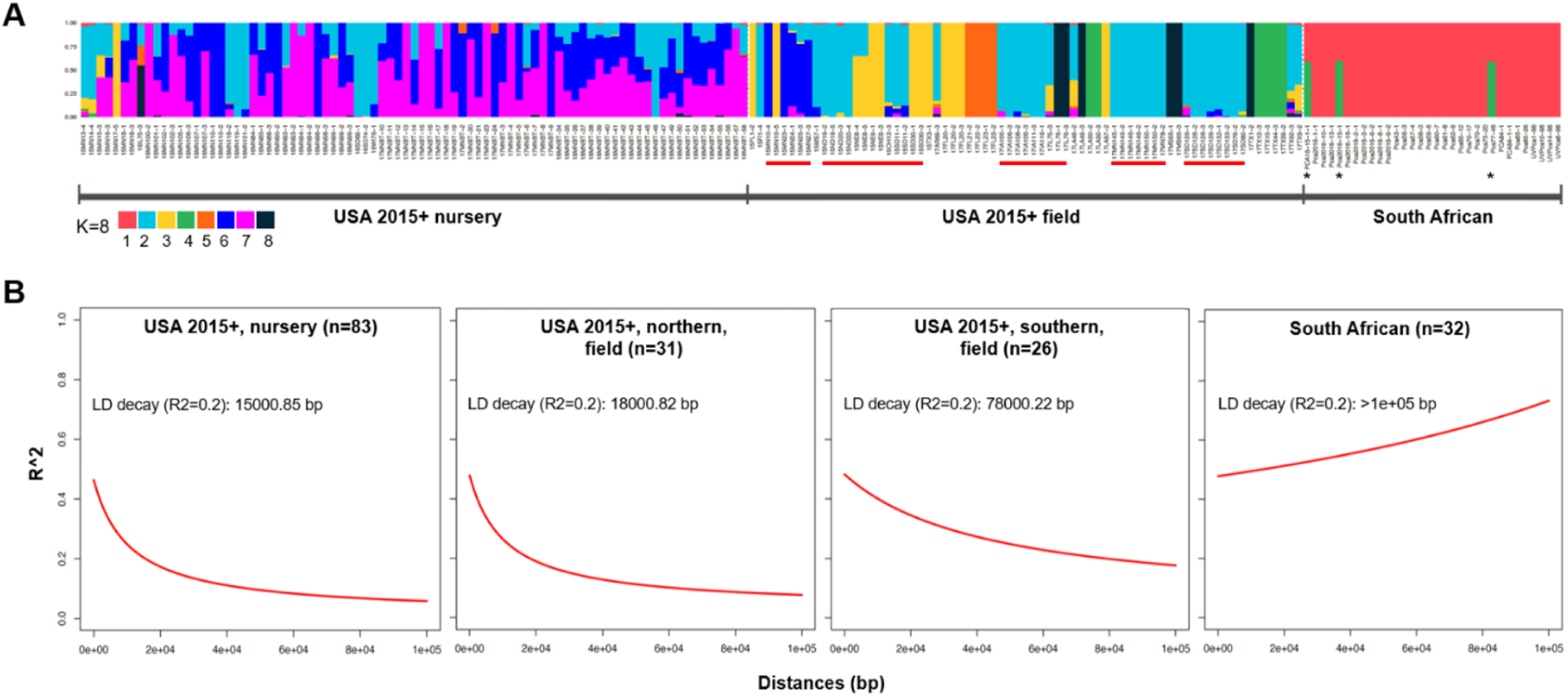
Population structure analysis and linkage disequilibrium (LD) decay analysis of *Pca* isolates from the USA and South Africa. **(A)** Population membership structure bar plot using 132,571 unlinked SNPs and based on K=8 clusters, the optimal number determined by cross-validation. Isolates underlined red were collected from field oats in northern USA states where buckthorn grows. The three South African isolates marked with black asterisks originated from wild oats. Cluster ID is shown as colours represented in the key. **(B)** LD decay plots based on 21,309 and 10,341 SNP variants identified against chromosome 1A of the Pca203 reference from USA isolates (only isolates collected in 2015 and subsequent years were included) and South African isolates, respectively. Population subsets labelled for each plot with LD decay indicated as a function of distance (bp).

Analysis of linkage disequilibrium (LD) was performed using SNPs called on chromosome 1A of the Pca203 genome reference (Henningsen et al., 2022) (**Fig. 4B**) to enable longer physical distance-based calculations. The highest rate of LD decay was shown in the USA population from the buckthorn nursery. The northern field population exhibited a similar rate of LD decay, consistent with a high frequency of sexual recombination in these populations due to the presence of buckthorn. Conversely, the southern field population showed a notably lower rate of LD decay, consistent with a lack of local recombination in the absence of buckthorn but with some LD decay maintained via migration occurring from regions with buckthorn. For the South African population, SNP sites were in complete disequilibrium and no LD decay was detected, consistent with asexual reproduction.

### Virulence phenotypes do not inform genetic relationships among diverse *Pca* isolates

Most rust surveillance programs rely heavily on pathotype (race) analysis using a limited set of host resistant differential lines, but it is not always clear whether this provides sufficient discrimination between lineages. We compared the virulence profiles and genotypes of a subset of 65 USA isolates selected to included representatives of diverse clonal groups. These data showed different groupings when clustered by virulence phenotype profile or by SNP phylogeny (**Supp. Fig. S7**). While most clonally related *Pca* isolates showed near-matching virulence profiles, the reverse was not always true as some isolates with highly similar virulence profiles belonged to genetically diverse groups. For instance, all isolates of clonal groups C and D had highly similar virulence profiles and clustered together by phenotype but were clearly separated as genetically distinct groups by phylogeny. Likewise, eight isolates collected from the buckthorn nursery in 2017 had very similar pathotypes and clustered together phenotypically, but belonged to four genetically distinct clonal pairs that were widely separated in the phylogeny. These observations argue that race assignments are poor indicators of genetic relationships in rust populations with high levels of diversity.

In contrast, comparison of phenotypes of South African *Pca* isolates (n=32) with a phylogenetic tree indicated a much clearer relationship, consistent with stepwise mutation of this single clonal lineage into groups with different pathotypes (**Supp. Fig. S8**). Several mutation events can be placed onto the phylogenetic tree of this lineage, with branches showing co-mutation to virulence on multiple *Pc* genes (**Supp. Fig S8**). Branch A shows simultaneous mutation to virulence on *Pc39*, *Pc55* and *Pc71*. Similarly, branch B shows mutation to virulence on both *Pc48* and *Pc52.* Finally, branch C includes one or more mutations to virulence on eight *Pc* genes *Pc39*, *Pc55*, *Pc71*, *Pc38*, *Pc57*, *Pc58*, *Pc59* and *Pc63*.

### Nuclear phased chromosome assembly enhances identification of virulence loci

To identify genomic regions linked to virulence on specific *Pc* genes, we conducted a GWAS analysis on isolates of the sexually recombining USA population. SNP genotypes were called separately against five reference assemblies: 12SD80 primary contigs; 12NC29 primary contigs; Pca203 full diploid genome; and the separate A and B haplotypes of Pca203. These data sets were analysed for genetic associations with the virulence scores of these isolates on individual oat differential lines (**Supp. Data S2**). In general, the use of the phased chromosome-scale Pca203 reference resulted in better discrimination of association regions than the more fragmented 12SD80 and 12NC29 references (**Fig. 5** and **Supp. Data S2**). For instance, association peaks that were split between multiple contigs in 12SD80 and 12NC29 were resolved to single peaks in the Pca203 assembly for 16 oat lines (Pc38, Pc39, Pc48, Pc51, Pc52, Pc53, Pc55, Pc57, Pc61, Pc91, Pc63, Pc68, Pc70, Pc71, Belle, HiFi). In contrast, for Pc35, Pc58, Pc62 and Pc64 single association peaks were detected on the 12SD80 and 12NC29 assemblies and were stronger than on Pca203. In all cases, contigs from 12SD80 and 12NC29 showing strong associations were syntenic with the chromosomal locations of the equivalent peaks in the Pca203 genome (**Supp. Fig. S9, Supp. Table S1**). For oat lines H548 and Pc54, an association peak was detected only in 12SD80 and 12NC29 references but no peak was detected in Pca203 (**Supp. Data S2**). Nonetheless, both 12SD80 and 12NC29 regions displayed synteny with each other and mapped to the same region on chromosome 1 of the Pca203 reference.

**Figure 5.**
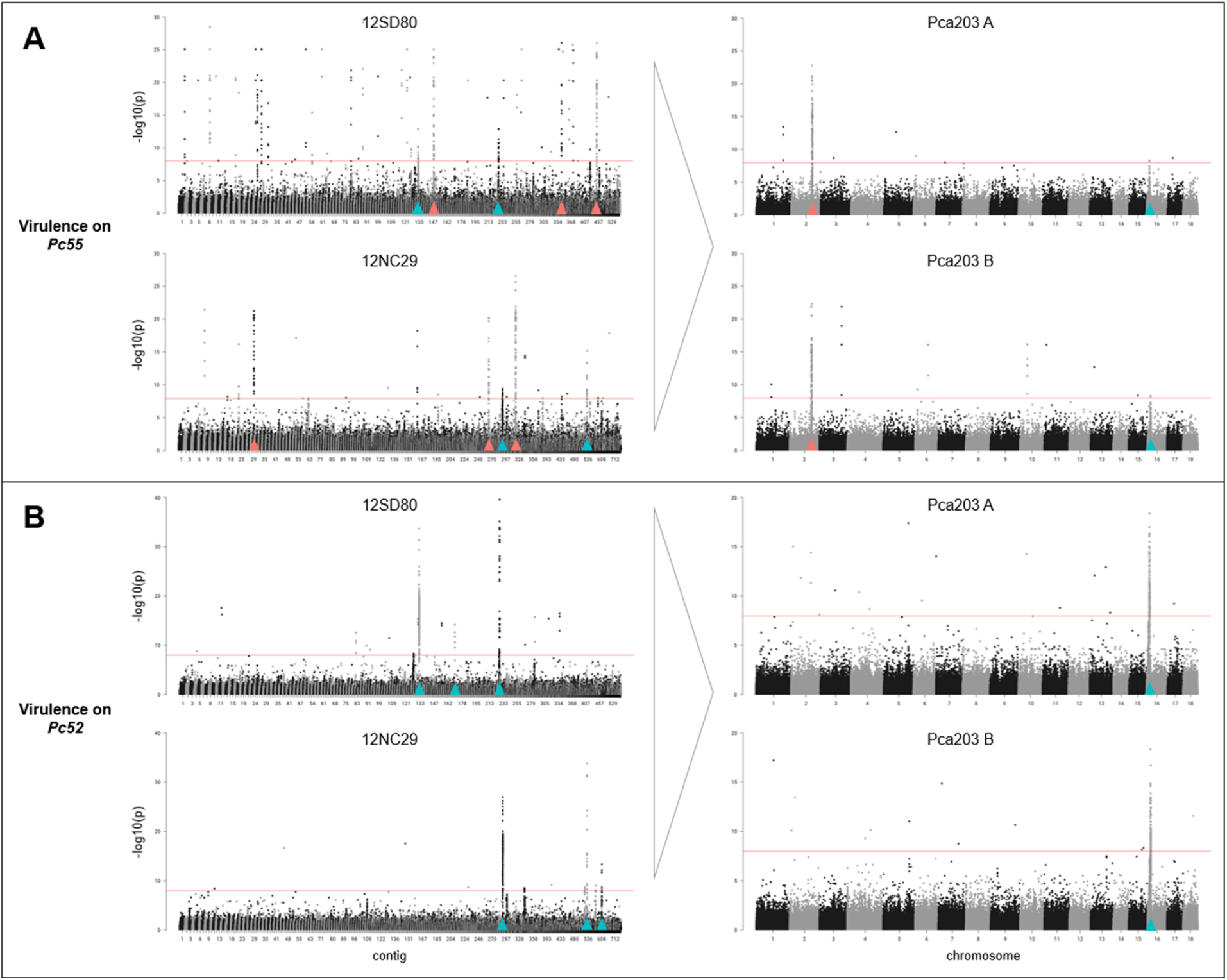
Manhattan plots derived from GWAS for virulence of USA isolates showing reduction in the number of association peaks using the fully phased Pca203 reference (separate A and B haplotypes) compared to previous reference assemblies (12SD80, 12NC29). **(A)** Association with virulence on *Pc55*. **(B)** Association with virulence on *Pc52*. Bonferroni significance thresholds (α = 0.01/total number of markers) are indicated by red horizontal lines. Peak regions marked in red or blue in 12SD80 or 12NC29 correspond to peaks marked with the same colour in Pca203 (A or B) based on sequence homology.

In total, significant peaks of virulence associated SNPs were detected for 25 of the oat differentials. Many of these locations overlapped between different oat lines, giving a total of 11 significant VGIs (**Table 1**, **Fig. 6**). These VGIs span between 10 and 320 kbp on the Pca203 genome. For 12 of the oat differential lines (TAM-O-405, Pc38, Pc39, Pc48, Pc51, Pc52, Pc55, Pc57, Pc61, Pc63, Pc70, and Pc71), the VGIs detected here correspond to those previously detected by Miller et al. (2020) using the 12SD80 and 12NC29 references and a smaller *Pca* population, although further refined here using the Pca203 reference. Associations with the other 13 differentials were not previously detected, showing the value of the larger population size and chromosome scale assembly. Only four VGIs were exclusive to a single oat line (VGI #5/Pc51, VGI #6/TAM-O-405, VGI #7/Pc53, VGI #9/Belle), while the remainder were detected by more than one oat differential line (**Fig. 6**). For example, four pairs of oat lines each detected a single strong association peak: Pc62 and Pc64 detecting VGI #1, Pc54 and H548 detecting VGI #2, Pc35 and Pc58 detecting VGI #8, and Pc48 and Pc52 detecting VGI #10. The latter, VGI #10, also appears as a minor peak for 10 other oat differential lines, while VGI #3 appears as a strong association peak for oat differential lines Pc38, Pc39, Pc70, Pc55, Pc63 and Pc71, and is also detected as a minor peak for Pc57, Pc61, Pc91 and WIX4361-9. Similarly, a common association on VGI #11 was detected for virulence on oat lines Pc36, Pc68, Pc91 and HiFi. Conversely, for 15 oat differential lines we detected a single VGI (**Fig. 6, Supp. Data S2**), consistent with these lines containing a single effective *R* gene with a single corresponding *Avr* locus. However, the other ten lines detected association peaks in two or more VGIs, suggesting that these oat differential lines may contain multiple different *R* genes.

**Figure 6.**
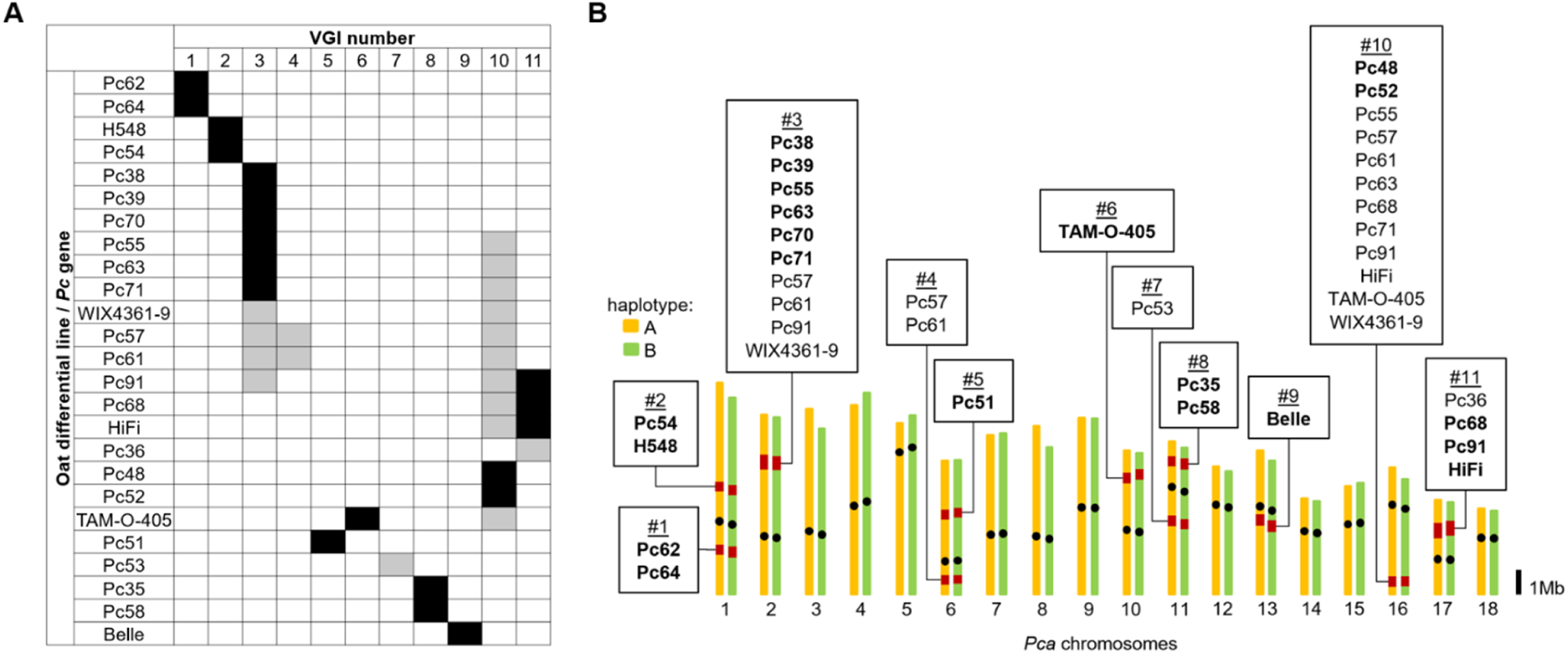
Chromosomal map of Pca203 showing locations of virulence associations. **(A)** Matrix indicating presence of GWAS peaks for virulence-associated genomic intervals (VGIs) in different oat differential lines. Black boxes denote presence of primary association peaks and grey boxes denote presence of secondary or minor association peaks. *Pc* genes/oat lines are grouped by shared VGIs. **(B)** Chromosomal map of Pca203 showing locations of VGIs in red labelled with *Pc* genes/oat lines having significant associations. Bold text indicates *Pc* genes/oat lines having a primary association peak for a given VGI whereas normal text indicates *Pc* genes/oat lines having a secondary or minor association peak for a given VGI. Black dots denote approximate locations of centromeres.

**Table 1.**
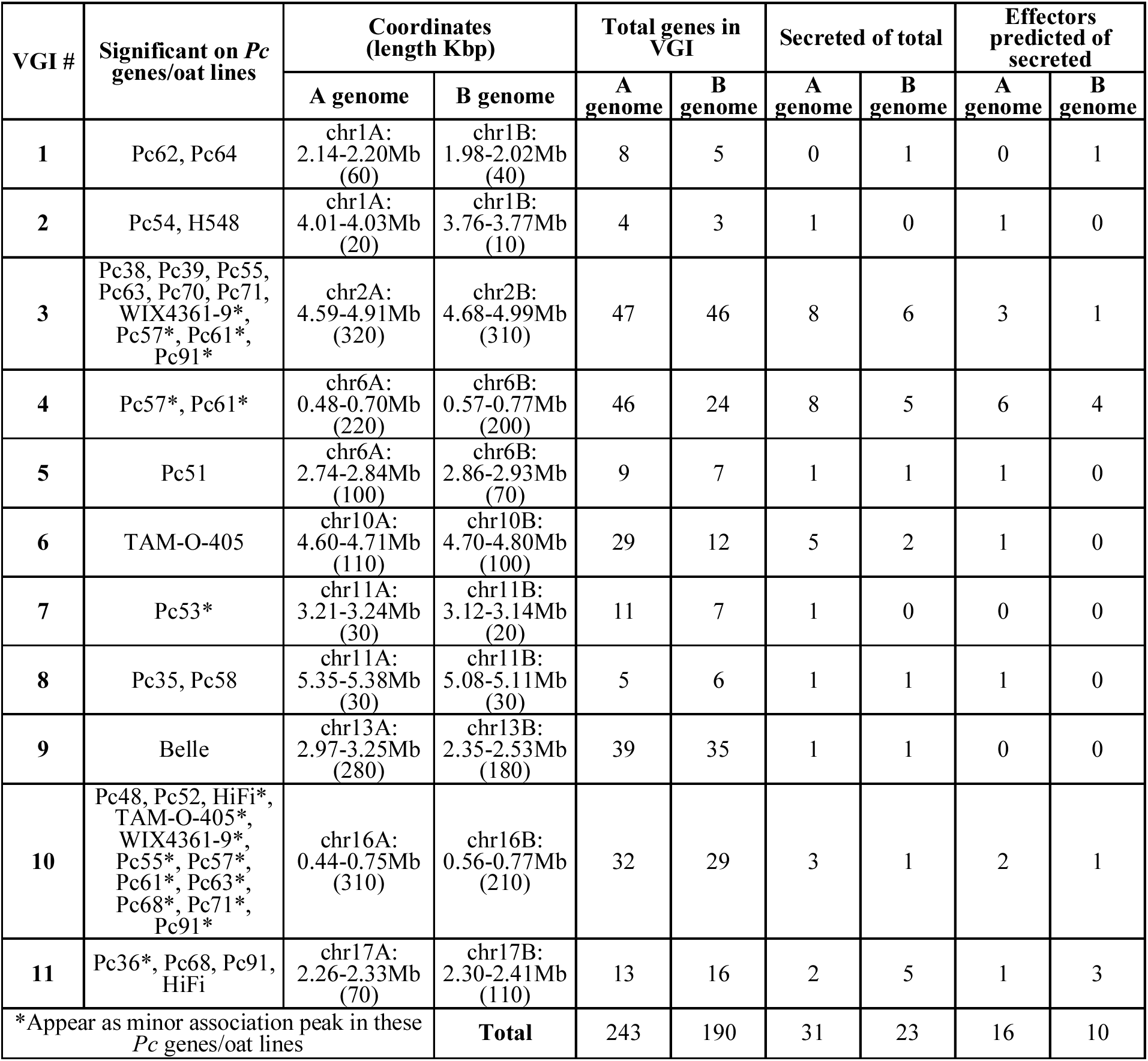
Virulence-associated genomic intervals (VGIs) determined from significant association peaks along with counts of annotated genes and predicted effectors within each VGI.

| VGI # | Significant on *Pc* genes/oat lines | Coordinates (length Kbp) |  | Total genes in VGI |  | Secreted of total |  | Effectors predicted of secreted |  |
| --- | --- | --- | --- | --- | --- | --- | --- | --- | --- |
|  |  | A genome | B genome | A genome | B genome | A genome | B genome | A genome | B genome |
| 1 | Pc62, Pc64 | chr1A:<br>2.14-2.20Mb<br>(60) | chr1B:<br>1.98-2.02Mb<br>(40) | 8 | 5 | 0 | 1 | 0 | 1 |
| 2 | Pc54, H548 | chr1A:<br>4.01-4.03Mb<br>(20) | chr1B:<br>3.76-3.77Mb<br>(10) | 4 | 3 | 1 | 0 | 1 | 0 |
| 3 | Pc38, Pc39, Pc55,<br>Pc63, Pc70, Pc71,<br>WIX4361-9*,<br>Pc57*, Pc61*,<br>Pc91* | chr2A:<br>4.59-4.91Mb<br>(320) | chr2B:<br>4.68-4.99Mb<br>(310) | 47 | 46 | 8 | 6 | 3 | 1 |
| 4 | Pc57*, Pc61* | chr6A:<br>0.48-0.70Mb<br>(220) | chr6B:<br>0.57-0.77Mb<br>(200) | 46 | 24 | 8 | 5 | 6 | 4 |
| 5 | Pc51 | chr6A:<br>2.74-2.84Mb<br>(100) | chr6B:<br>2.86-2.93Mb<br>(70) | 9 | 7 | 1 | 1 | 1 | 0 |
| 6 | TAM-O-405 | chr10A:<br>4.60-4.71Mb<br>(110) | chr10B:<br>4.70-4.80Mb<br>(100) | 29 | 12 | 5 | 2 | 1 | 0 |
| 7 | Pc53* | chr11A:<br>3.21-3.24Mb<br>(30) | chr11B:<br>3.12-3.14Mb<br>(20) | 11 | 7 | 1 | 0 | 0 | 0 |
| 8 | Pc35, Pc58 | chr11A:<br>5.35-5.38Mb<br>(30) | chr11B:<br>5.08-5.11Mb<br>(30) | 5 | 6 | 1 | 1 | 1 | 0 |
| 9 | Belle | chr13A:<br>2.97-3.25Mb<br>(280) | chr13B:<br>2.35-2.53Mb<br>(180) | 39 | 35 | 1 | 1 | 0 | 0 |
| 10 | Pc48, Pc52, HiFi*,<br>TAM-O-405*,<br>WIX4361-9*,<br>Pc55*, Pc57*,<br>Pc61*, Pc63*,<br>Pc68*, Pc71*,<br>Pc91* | chr16A:<br>0.44-0.75Mb<br>(310) | chr16B:<br>0.56-0.77Mb<br>(210) | 32 | 29 | 3 | 1 | 2 | 1 |
| 11 | Pc36*, Pc68, Pc91,<br>HiFi | chr17A:<br>2.26-2.33Mb<br>(70) | chr17B:<br>2.30-2.41Mb<br>(110) | 13 | 16 | 2 | 5 | 1 | 3 |
| *Appear as minor association peak in these *Pc* genes/oat lines |  |  | **Total** | 243 | 190 | 31 | 23 | 16 | 10 |

Comparison of the resistance profiles of differential lines that detect the same VGIs showed substantial overlap in their recognition specificity, but also some differences (**Fig. 7**). For example, Pc35 and Pc58 both detected VGI #8 and showed similar profiles (**Fig. 7A**), although the Pc58 differential gave intermediate responses to isolates that were fully virulent on Pc35. Similarly, Pc48 and Pc52 (VGI #10) showed similar profiles, but with some isolates fully virulent on Pc52 giving intermediate responses on Pc48 (**Fig. 7B**). Similar observations were made for other lines detecting common VGIs (**Fig. 7C-G**). Overall, these data suggest that *R* genes in some differential lines recognise *Avr* genes in *Pca* that occur at the same loci, either as allelic variants or closely linked genes.

**Figure 7.**
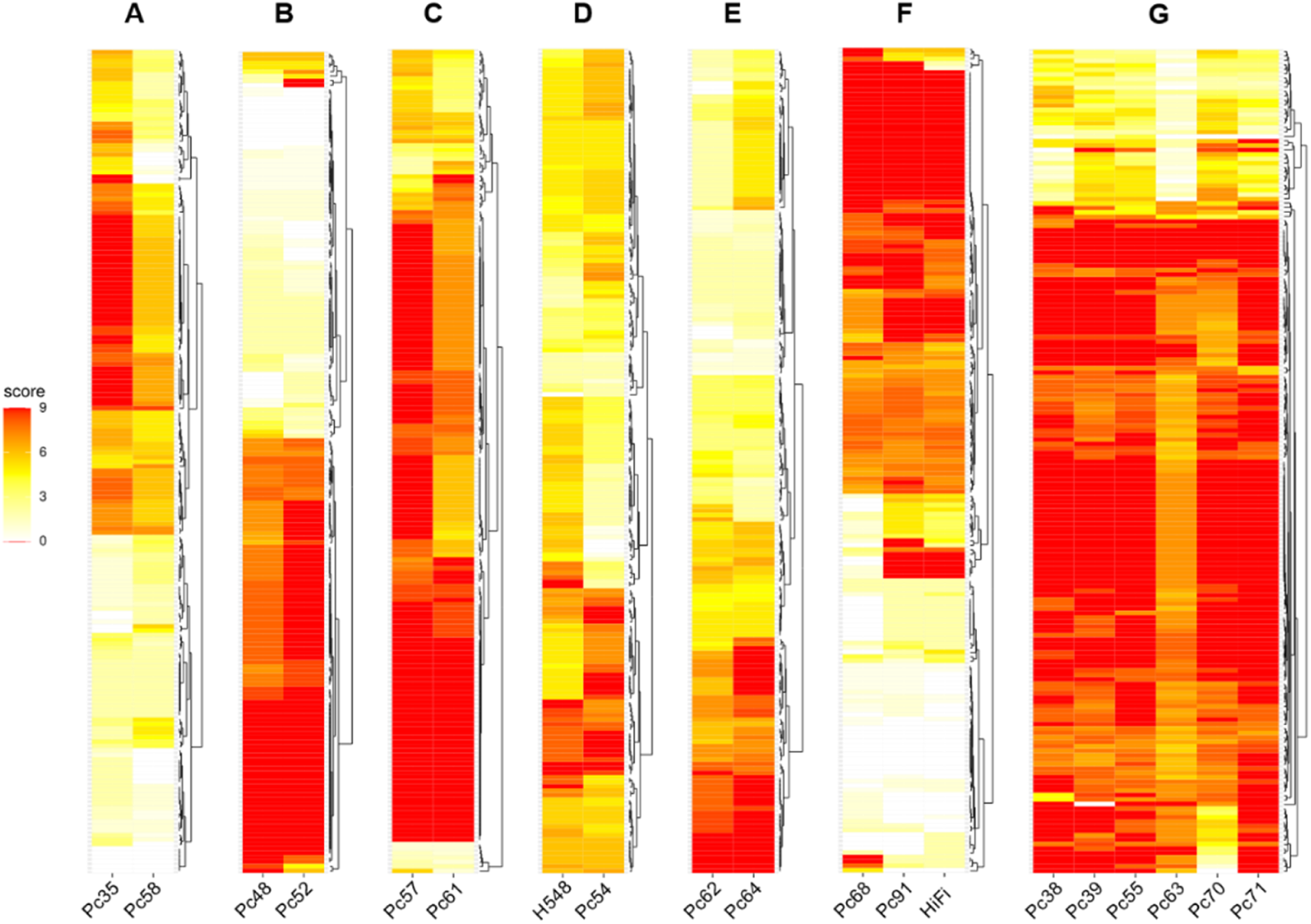
Heatmaps of virulence of 185 USA *Pca* isolates scored on oat differential lines grouped by shared genomic associations in *Pca*. High scores (red) indicating high virulence (disease susceptibility) and lower scores (yellow/white) indicate avirulence (disease resistance). Isolates of Pca (y-axis) are ordered independently according to hierarchical clustering of scores. Oat lines (x-axis) producing major virulence-associated genomic intervals (VGI) in common are grouped accordingly: **(A)** VGI #8; **(B)** VGI #10; **(C)** VGI #4; **(D)** VGI #2; **(E)** VGI #1; **(F)** VGI #11; **(G)** VGI #3.

To identify potential *Avr* gene candidates within the VGIs (**Supp. Data S1**), we examined existing gene models (Henningsen et al., 2022) within each VGI (**Table 1**). All the VGIs contained at least one and up to eight gene models encoding secreted proteins in either haplotype, and between one and six of those were predicted as cytoplasmic or dual-localized effectors by EffectorP3.0 (Sperschneider and Dodds, 2022), except in VGIs #7 (Pc53) and #9 (Belle) in which no effectors were predicted. VGI #3 corresponded to the highest number of virulence associations with oat differential lines, which included Pc38, Pc39, Pc55, Pc63, Pc70 and Pc71 and contained 8 and 6 predicted secreted protein genes in the A and B haplotypes respectively, with 3 (A haplotype) and 1 (B haplotype) predicted as cytoplasmic effectors. The presence of multiple effector candidates in this region is consistent with the possibility that different oat *R* genes could recognise closely linked *Avrs*. On the other hand, VGI #10, which was also associated with virulence on a large number of differential lines contained only one or two predicted effectors, suggesting that these differentials may recognise the same *Avr* gene, or alleles thereof.

To assess whether VGIs coincided with signals of recent host adaptation in *Pca* we conducted a selective sweep analysis of the nursery isolates (**Supp. Fig. S10**). A total of 54 sweep loci were identified, with 10 in the A haplotype and 44 in the B haplotype and ranging in size from 60kbp to 183.5kbp and spanning from 2 to 34 genes. Collectively, detected sweep loci summed to 611kbp from the A sub-genome and 3,186kbp from the B sub-genome. However, just 13 sweep loci on chr12B made up 1,247kbp of this. Of all sweep loci detected, only two coincided with identified VGIs: VGI #3 on chromosome 2B associated with virulence on Pc38/39/55/63/70/71, and VGI #9 on chromosome 13A associated with virulence on Belle (**Supp. Fig. S10**).

## DISCUSSION

Knowledge of pathogen population dynamics and the genetic architecture of host-pathogen interactions is important to understand the emergence of new virulence traits in response to the deployment of race-specific *R* genes in crops. The oat crown rust pathogen, *Pca*, has shown particularly rapid evolution of new virulence phenotypes in the USA, which is thought to be due to the involvement of the alternate host, common buckthorn, allowing sexual reproduction and reassortment of virulence alleles. While investigating this hypothesis, Miller et al. (2020) found evidence for sexual recombination from genome sequence data of a limited sample of USA *Pca* isolates but also observed the accumulation of some clonal lineages. However, Miller et al. (2020) was not able to discriminate between different geographic regions in the USA where buckthorn is present or absent to establish a link between virulence evolution and sexuality. Here we took advantage of a new chromosomal-scale and nuclear phased reference genome of *Pca* (Henningsen et al., 2022) and a substantially larger collection of recent *Pca* isolates from the USA to expand on our understanding of the effects of geography and sexuality on the genetic diversity of *Pca*. We found that the *Pca* population in the northern USA, where buckthorn is prevalent, is almost entirely sexual in nature, as evidenced by the strong support for recombination in the neighbour-net tree, the extreme genetic separation of individual isolates in the phylogenetic tree and the rapid decay of LD in this population. This suggests that the northern USA *Pca* population is mainly derived each growing season from sexually recombined aeciospores produced on buckthorn bushes during the previous winter. This is in contrast to the situation observed in the USA for wheat and barley rust diseases, for which spore dispersal follows the well-known “Puccinia Pathway” with infections manifesting from south to north as the season progresses and wind-borne inoculum from the earlier planting southern regions spreads north as the season progresses (Stakman and Harrar, 1957; Hamilton and Stakman, 1967; Fetch et al., 2011). Initiation of these rust infections in northern regions are dependent on urediniospore inoculum spread from the south, since these pathogens cannot overwinter in the north due to the absence of a green bridge. However, the presence of the alternate host buckthorn allows *Pca* to overwinter in the north and initiate locally derived infections on oat at the start of the growing season through sexually derived aeciospores produced on buckthorn. This also underpins the high levels of genetic diversity in this population. *Pca* populations in the southern regions of the USA show more prevalent clonal lineages but nevertheless are still highly diverse and with low levels of LD. This indicates that southern populations are influenced by two separate factors; local propagation and maintenance of clonal lineages; as well as regular introductions of new sexually derived lineages by migration from the north. Survival of *Pca* lineages in the south could also be facilitated through infection of wild oats acting as an additional reservoir for the pathogen between growing seasons. Some genotypes detected in 2015 in the northern USA were found in southern areas in 2017, with some clonal groups comprising both northern and southern isolates. Thus, north to south migration is also a significant factor in epidemiology of this disease, in addition to the traditional south to north “Puccinia Pathway”. Consistent with this, annual rust surveys in the USA have detected a sudden spread of virulence traits in the south in recent years which appeared to be preceded by a steady accumulation of virulence traits in the north (Moreau and Kianian et al., unpublished). Interestingly, our study also detected a long-lasting lineage still present in 2017 that has persisted in the USA since at least 1990 and includes isolates collected from Texas, Louisiana, Georgia, Minnesota, South Dakota, Kansas, and North Carolina. Miller et al. (2020) found a substantial shift in the population towards higher virulence on numerous *Pc* genes between 1990 and 2015, and our data shows evidence for this trend continuing. For example, virulence frequency for Pc68, Pc91 and HiFi oat differential lines rose from 26-30% in 2015 to 43-46% in 2017. This is consistent with reports from Canada (Menzies et al., 2019), particularly for *Pc91* whose virulence frequency increased from 0% in 2011 to 66% by 2015, and in more recent USA rust surveys reporting up to 97% since 2020 from just 41% in 2015 (Moreau and Kianian et al., unpublished).

The SA *Pca* population (Boshoff et al., 2020) shows a sharp contrast with the USA population. SA isolates collected from cultivated oat across the country between 1998 and 2018 represent a single clonal lineage with genetic markers in complete LD. Interestingly, three isolates collected from wild oats in this region represented a different clonal lineage, although genetically related. A more in-depth sampling of the SA *Pca* population, particularly from wild oats, will be required to determine the relationships between *Pca* on these different hosts. Interestingly, the USA isolate 90MN5B-1, collected in 1990, appears related to the SA *Pca* lineages. It is possible that 90MN5B-1 represents an exotic introduction with a founding effect in SA or that 90MN5B-1 and the SA population are commonly derived from a broader intercontinental lineage divergent from the USA population. Further studies on international collections of *Pca* will shed light on the distribution and diversity of different global lineages.

The use of a nuclear phased genome assembly for Pca203 and a larger population resulted in identification of virulence associations for more oat differential lines than was previously possible (Miller et al., 2020) and allowed for better resolution of VGIs in the *Pca* genome. However, the complementary use of the alternate references also aided in detecting associations that were significant in one reference but absent or weak in another; for instance, Pc54/H548 detected a strongly associated region on the 12SD80 and 12NC29 references, but not Pca203, although the latter reference helped to map this VGI to a chromosomal location. Such variation between references may result from divergence between the reference isolate and the segregating population, reducing the ability to map sequence reads and call SNPs in some regions. Overall, we detected 11 VGIs in the *Pca* genome corresponding to 25 of the oat differential lines (**Table 1**, **Fig. 6**). In four cases there were one-to-one relationships between single differential lines and VGIs: Pc51 (VGI#5), Pc57 (VGI#7), TAM-O-405 (VGI #6), and Belle (VGI #9), consistent with a simple gene-for-gene relationship between unique *R* and Avr gene pairs. In other cases, multiple oat lines detected the same VGI, such as Pc62 and Pc64 (VGI #1) Pc54 and H548 (VGI #2), Pc35 and Pc58 (VGI #8), and Pc48 and Pc52 (VGI #10). These may represent cases of the same *Pc* gene or an allelic variant recognising the same *Avr* gene being incorporated into the differential set independently. In other cases, one *Pc* differential detected multiple VGIs, suggesting the presence of multiple *R* genes in the same line. These more complex situations urge careful evaluation of the composition of the oat differential set. Although many of the differential lines have had very limited genetic analysis to determine the number or identity of *R* genes present, several reports in the oat crown rust pathosystem accompanied by virulence phenotypic comparisons can assist the interpretation of these GWAS results. For example, the genes *Pc39*, *Pc55* and *Pc71* were shown to be closely linked and postulated as either allelic or the same gene (Kiehn et al., 1976; Leonard et al., 2005). This is consistent with the observation that they all detect VGI #3 in the GWAS analysis. Furthermore, we observed a single branch (A) in the South African *Pca* clonal lineage with mutation to virulence on these three genes, consistent with these three genes recognising the same *Avr* gene and virulence resulting from a single mutational event. Similarly, Harder et al. (1980) proposed genetic linkage or allelism of *Pc38* and *Pc63*, which also detect VGI #3. However, the South African mutant branch A clearly differentiates these two groups of genes, although branch C shows simultaneous mutation to virulence on all these lines. Other individual mapping studies have assigned *Pc38, Pc71* and *Pc39* to different locations in the oat linkage map (Bush and Wise, 1998; Wight et al., 2005; Sowa and Paczos-Grzęda, 2020). Given the complex rearrangements seen in the oat genome references, this may reflect the translocation of syntenic regions to different positions in the genome through independent introgressions. These *Pc* genes may detect separate but closely linked *Avr* loci, or potentially have different recognition of the allelic variants of the same *Avr* gene.

The Pc48 and Pc52 lines both detected the same *Pca* locus (VGI#10), which is consistent with the reported positive association for *Pc48* and *Pc52* virulence by Chong and Zegeye (2004). Again, a single branch (B) in the South African clonal lineage showed mutation to virulence on both lines, supporting their recognition of the same *Avr* locus. However, although Pc48 and Pc52 (VGI #10) showed similar virulence profiles, they were differentiated by some isolates fully virulent on Pc52 that gave intermediate responses on Pc48 (**Fig. 7B**). This suggests that they may carry related *R* genes or alleles of the same *R* gene recognising the same Avr effector but with some quantitative differences in recognition of *Avr* gene variants. Similarly, Pc35 and Pc58 both detected VGI #8 and showed similar resistance profiles (**Fig. 7A**), although the Pc58 differential gave intermediate responses to isolates that were fully virulent on Pc35. Mapping studies have also suggested *Pc58* resistance is conditioned by a cluster of three genes (Hoffman et al., 2006), supporting a possible distinction from *Pc35*. Overall, these data suggest that *R* genes in some differential lines recognise *Avr* genes in *Pca* that occur at the same loci, either as allelic variants or closely linked genes.

Several oat differential lines detected virulence associations at multiple VGIs suggesting they may contain multiple *R* genes. For instance, the cultivar WIX4361-9 was reported to carry two unspecified *Pc* genes (Bonnett, 1996), and detected VGIs #3 and #10. Given that both VGI #3 and #10 are also associated to virulence on multiple oat lines, the identity of those unknown *Pc* genes in WIX4361-9 remains unclear but could include variants of the Pc38/39/55/63/71 and Pc48/52 groups. The *Pc91* oat differential detected three VGIs, (#3, #10 and #11). Menzies et al. (2019) found a positive correlation of virulence to *Pc91* with both *Pc48*, *Pc39*, which is consistent with these latter two detecting the VGI #3 and VGI #10 respectively and may indicate the presence of three different *R* loci in this line. Given the high prevalence of virulence to the Pc48/52 and Pc38/39 groups in current populations of *Pca*, many isolates would not detect such background genes, explaining the original postulation of *Pc91*. The oat cultivar HiFi showed a very similar virulence profile to Pc91 and detected VGI #10 and #11. HiFi was developed by a series of crosses including Amagalon, the differential line carrying *Pc91* (McMullen et al., 2005), so it is likely that it carries the same *Pc* gene(s). Pc68 also detected VGI #10 and #11 and showed a similar resistance profile to Pc91 and HiFi, but these lines were distinguished by several isolates with reciprocal contrasting virulence for *Pc68* and *Pc91*. Thus, these differentials may all contain multiple *R* genes with some of these shared between Pc68 and Pc91. This suggests that researchers independently identified and transfer alleles or perhaps the even same variant, called it *Pc68* or *Pc91*, from wild relatives *A. sterilis* and *A. magna*, respectively (Wong et al., 1983; Rooney et al., 1994).

The complex virulence relationships on some oat lines involving multiple loci with overlapping associations highlight limitations in our knowledge of the genetics of resistance in some of these differentials, and the molecular basis of recognition. Other than the presence of multiple genes in some differentials as described above, other explanations for this complexity include epistatic interactions between virulence loci and the presence of multiple *Avr* genes in a genetic cluster. For example, two *Avr* genes are separated by only 15kbp in the genome of wheat stem rust, yet are specific to unrelated *R* genes (*Sr35*, *Sr50*) (Li et al., 2019). Conversely, the wheat powdery mildew *R* gene *Pm1* was found to detect two separate, unlinked *Avr* loci, each encoding proteins predicted to have structural similarity (Kloppe et al., 2023). Epistasis may also present as alternative pathogen loci that moderate the virulence response, such as inhibitor loci that suppress recognition of certain *Avr* genes, as observed in flax rust (Jones, 1988) and wheat powdery mildew (Bourras et al., 2015).

Here, we established the role of geography, sexuality and clonality as factors shaping the virulence evolution of *Pca*. Future exploration on the influence of wild oat populations is also warranted. Nevertheless, the possible contribution of somatic hybridisation to the genetic diversity and evolutionary capacity of the pathogen cannot be ruled out until additional haplotype genome references are available. Ongoing work to generate a pangenome of *Pca* and capture haplotype diversity across geographic regions and clonal lineages will help to investigate this possibility, and will likely improve the capacity to identify *Avr* loci by GWAS and isolate candidate genes. Rust pathology and surveillance has relied heavily on the characterisation of virulence profiles and subsequent race (pathotypes) assignment to infer genetic relationships among rust isolates and populations. While such practice can be informative in a wholly clonal population carrying lineage specific mutations (e.g. South African subpopulations), the occurrence of incursions, sexual recombination, somatic hybridisation and presence of multiple *Avr* loci can confound associations between phylogeny and race assignment as shuffling of virulence alleles allows pathotype combinations to emerge more than once in recombining populations. The integration of phenotypic data with high resolution genotypic data from key rust lineages and haplotype combinations will transform rust surveillance programs by bringing speed, depth, and consistency to surveys around the globe. In the meantime, it would be beneficial to invest in a thoroughly characterised and curated differential set of non-redundant isogenic lines. For this, additional efforts to genetically map *Pc* genes as well as develop and implement high quality molecular markers would be instrumental, not only to assign pathotypes in a robust manner, but also to screen and identify novel sources of oat crown rust disease resistance.

## MATERIALS & METHODS

### Rust sampling, plant inoculations, and virulence phenotyping and comparisons

*P. coronata* f. sp. *avenae* (*Pca*) isolates from the annual surveys by the USDA-ARS Cereal Disease Laboratory (Saint Paul, MN, USA) were accessed as single pustule cultures either from -80°C storage or through the oat cropping season. Isolates were subjected to a second single-pustule purification and increased and tested for purity (Miller et al., 2018). Forty-six isolates were collected from aecia on buckthorn leaves at the Minnesota Matt Moore buckthorn nursery (Saint Paul, MN, USA) (**Supp. Data S1**) and inoculated onto oat cultivar ‘Marvelous’ for a two-step single-pustule purification followed by spore increase. Thirty-two *Pca* isolates were collected in SA in 1998, 2005, 2016, 2017 and 2018, including three isolates from wild oats (Boshoff et al., 2020), and were increased from single pustules on the oat cultivar ‘Makuru’. The virulence pathotypes for USA isolates were defined using a set of 40 North American oat differential lines (Nazareno et al., 2018) while SA pathotypes were defined on a smaller differential set with some common lines (Boshoff et al., 2020). Infection types were scored 10–12 days after inoculation, with a scale of “0”, “0;”, “;”, “;C”, “1;”, “1”, “2”, “3”,”3+”, and “4” (Nazareno et al., 2018) which was converted to a 0–9 numeric scale and mean values from two independent score readings were used for statistical analysis. Race assignments were made according to standard four-letter and ten-letter letter nomenclatures (Chong et al., 2000; Nazareno et al., 2018).

Wilcoxon rank sum tests were performed as two-tailed operations using the ‘wilcox.test’ function in RStudio (v4.0.2) (RStudio-Team, 2022) to compare overall virulence between isolate groups using pooled scores from the entire oat differential set as well as that of individual differential lines. Violin plots of overall virulence distribution along with boxplots of gene-specific virulence distribution between isolate groups were generated using ggplot2 (v3.3.6) (Wickham, 2016). Clustered heatmaps were made from the scoring matrices using the R package ComplexHeatmap (v1.14.0) (Gu et al., 2016).

For geographical comparisons, northern USA states with buckthorn prevalent were considered as Minnesota (MN), Wisconsin (WI), North Dakota (ND), Nebraska (NE), Iowa (IA), Illinois (IL), and Pennsylvania (PA), while southern states where buckthorn is rare or absent were Kansas (KS), Arkansas (AR), Texas (TX), Louisiana (LA), Missouri (MS), Georgia (GA), and Florida (FL). Isolates from South Dakota (SD) (n=12) were excluded from this analysis because this state is geographically northern but has a low prevalence of buckthorn (Rawlins et al. (2018).

### DNA sequencing, read mapping and variant calling

DNA was extracted from 20 mg of urediniospores of each isolate with the G-Biosciences Omniprep DNA isolation kit, and the Illumina TruSeq Nano DNA library preparation protocol was used for sequencing with Illumina NovaSeq on an S2 flow cell to generate 150 bp paired-end reads as described in Henningsen et al. (2022). FASTQ reads were trimmed with Trimmomatic (v0.38) (Bolger et al., 2014) with a sliding window of 15bp (incremented by 4bp), leading and trailing low quality (<10) bases removed, adapter clipping (seed mismatches set to 2, palindrome threshold 30, simple clip threshold 10, minimum adapter length of 2), and reads less than 100bp discarded. Trimmed reads were aligned independently to the reference genomes (12SD80 and 12NC29 primary contigs, Pca203 diploid chromosomes) using the ‘bwa mem’ algorithm of BWA (v0.7.71) (Li and Durbin, 2009). BAM files were processed with SAMtools (v1.12) (Li et al., 2009) and Picard (v2.26.9) (https://broadinstitute.github.io/picard/), including removal of duplicate reads and relabelling of reads by sample. Coverage statistics were generated using ‘samtools depth -a’ and piped to a custom AWK script then plotted using R (https://github.com/TC-Hewitt/OatCrownRust).

Variant calling was performed separately for each reference using FreeBayes (v1.3.5) (Garrison and Marth, 2012) with option ‘--use-best-n-alleles 6’ and VCF files were filtered using *vcffilter* of vcflib v1.0.1 (https://github.com/vcflib/vcflib) with the parameters ‘QUAL > 20 & QUAL/AO > 10 & SAF > 0 & SAR > 0 & RPR > 1 & RPL > 1 & AC > 0.’ VCFtools (v0.1.16) (Danecek et al., 2011) was then used to select biallelic sites with less than 10% missing data and minor allele frequency of 5% or greater. Allele balance plots were generated from SNPs called against 12NC29 using a custom R script (https://github.com/henni164/Pca203_assembly/blob/master/figure_s2/203_frequencies.R). Genome assemblies and accompanying annotations of 12SD29 and 12NC80 were sourced from the DOE-JGI Mycocosm Portal (http://genome.jgi.doe.gov/PuccoNC29_1 and http://genome.jgi.doe.gov/PuccoSD80_1).

### Phylogenetic analysis

VCF output was converted to NEXUS and PHYLIP formats with vcf2phylip (https://github.com/edgardomortiz/vcf2phylip). Phylogenetic analysis was performed in RAxML (v8.2.12) (Stamatakis, 2014) using SNPs called in 12SD80 using PHYLIP input with ML criterion, 500 bootstrap replicates and a general time reversible CAT model (GTRCAT). Trees were visualised in iTOL v5 (Letunic and Bork, 2021).

For phylogenetic analysis of only South African isolates, a subset was taken from the raw VCF of variant calls against 12SD80, with filtering and selection performed as above. Analysis was performed using RAxML as above except with 1000 bootstrap replicates. The R package ggtree (v3.4.0) (Yu et al., 2017) was used to illustrate the phylogenetic tree and virulence heatmap using the ‘gheatmap’ function. Likewise, phylogenetic analysis of 65 clonal isolates from the USA population were taken as a subset from the original VCF file and processed as described above. The phylogenetic tree was created with ggtree as above while the cluster dendrogram was generated using the R package ggdendro (v0.1.23) (https://github.com/andrie/ggdendro). Accompanying virulence heatmaps ordered by cluster or by phylogeny were generated separately using ggplot2 (v3.3.6). An unrooted phylogenetic network was created from the NEXUS file and PHI-tests (Bruen et al., 2006) for signatures of past recombination were performed using SplitsTree (v4.16.2) (Huson and Bryant, 2005).

### Population structure, linkage disequilibrium and selective sweep analysis

Population structure was investigated using a model-based approach implemented using fastSTRUCTURE (Raj et al., 2014) on SNPs called against 12SD80. SNP pairs were pruned, maintaining only markers with r^2^ < 0.6 in 500kbp windows, using ‘+prune’ in BCFtools (v1.15.1) (Danecek et al., 2021). Elbow plots were generated using the script ‘chooseK.py’ from fastSTRUCTURE and membership assignment barplots generated using the R package pophelper (v1.2.0) and the program CLUMPP (v1.1.2) (Jakobsson and Rosenberg, 2007; Francis, 2017). PCA was performed on pruned SNPs using the ‘--pca’ function in PLINK v2.0 (Chang et al., 2015) and plots generated in RStudio (v4.0.2) using ggplot2 (v3.3.6). LD decay was estimated using SNPs called against chromosome 1 A of Pca203 in windows of 100kb using the function ‘--geno-r2’ and option ‘--ld-window-bp 100000’ in VCFtools.

Selective sweeps were identified based on SNPs called against the Pca203 diploid reference using a composite likelihood ratio (CLR) method implemented in SweeD (v3.0) (Pavlidis et al., 2013). Only isolates sampled from the buckthorn nursery were included based on membership analysis using K=5 and a membership value above 0.9 for cluster 1. Isolates systematically assigned to a different genetic cluster in the fastSTRUCTURE results were removed resulting in a final dataset of 41 isolates. SweeD was run against individual chromosomes using the option ‘-folded’ in 1kb grids with significance threshold set at the 999^th^ percentile. SNP density was determined across the genome in nonoverlapping windows of 50kbp and only CLR values within 50kbp regions with at least 35 SNPs were kept. Karyoplots illustrating chromosomal positions of VGIs and selective sweep loci were created using the R package karyoploteR (Gel and Serra, 2017) with manually generated GRanges input.

### Genome-Wide Association Study (GWAS)

Virulence association testing was performed using 182 U.S. *Pca* isolates (**Supp Data S1**) based on scored phenotypes (0-9 point scale) for each oat differential line. Filtered biallelic SNPs were called for this dataset against primary contigs of 12SD80 (1,012,613 sites) and 12NC29 (856,274 sites), and chromosomes of Pca203 diploid (372,275 sites) and A and B haploid (937,120 and 956,257 sites) genome references as described above. Marker-trait associations were conducted for virulence scores on each of the forty oat differential lines in TASSEL (v5.2.63) (Bradbury et al., 2007) using a Mixed Linear Model (MLM), with population structure and kinship calculated using four principal components and centred identity by state (IBS), respectively. No compression was performed, and variance components were re-estimated after each marker. Quantile-quantile (QQ) and Manhattan plots were generated in R using the qqman package (v0.1.8) (Turner, 2018). The R package RAINBOWR (Hamazaki and Iwata, 2020) was used to compute false discovery rates (FDR) at 5%. Bonferroni correction thresholds were calculated by dividing the p-value of 0.01 (or 0.05 for Pca203 diploid genome) by the number of markers in each reference. This resulted in a threshold of 9.875×10^-9^ for 12SD80, 1.168×10^-8^ for 12NC29, 1.0671×10^-8^ for Pca203 A sub-genome, and 1.0457×10^-8^ for Pca203 B sub-genome. As for the Pca203 diploid genome, which had fewer markers, an α = 0.05 was used, resulting in a threshold of 1.3431×10^-7^.

VGIs on A and B sub-genomes of Pca203 were defined based on the start and end positions of the set of SNP markers within an association peak that exceed the Bonferroni significance threshold, expanded to the nearest kilobase. For associations with few markers above the Bonferroni threshold, the FDR threshold was used instead (VGI #1, #7, #8, #9).

### Synteny analysis and effector prediction

To check for synteny between virulence associated regions of the three genome references, alignments were visualised using D-Genies (Cabanettes and Klopp, 2018). Whole genome alignment of 12SD80 and 12NC29 haplotigs to Pca203 chromosomes was performed using minimap2 (v2.24) (Li, 2018). SAMtools (v1.12) was used for extraction of significant regions and haplotigs from reference FASTAs, and minimap2 was again used for all-vs-all alignments of extracted sequences using option ‘-X’ to avoid self-hits. Synteny plots were generated using the R package gggenomes (Hackl and Ankenbrand, 2022) in RStudio (v4.0.2). Protein sequences were compared using BLAST+ (v2.13.0) (Camacho et al., 2009) and ClustalOmega (Sievers and Higgins, 2018). Secreted proteins were predicted using SignalP (v4.1) (-t euk -u 0.34 -U 0.34) (Petersen et al., 2011) and TMHMM (v2.0) (Krogh et al., 2001). A fungal protein was considered secreted if it was predicted to have a signal peptide and had no transmembrane domains outside the N-terminal region. Effector proteins were predicted with EffectorP (v3.0) (Sperschneider and Dodds, 2022).

## DATA AVAILABILITY

Raw Illumina sequence reads of 162 USA and South African isolates used in this study are available in the NCBI BioProject PRJNA660269. Reads of 62 USA isolates used in this study generated by Miller et al., (2020) are available in the NCBI BioProject PRJNA398546. Custom scripts and programs used for analysis in this publication are available at https://github.com/TC-Hewitt/OatCrownRust. VCF files for future marker assisted diagnosis used in this study are available at the CSIRO Data Portal https://data.csiro.au/collection/csiro:60078. Pca203 assembly and RNAseq reads (Henningsen et al., 2022) used in this study are available at https://data.csiro.au/collection/csiro:53477.

## AUTHOR CONTRIBUTIONS

TCH: phylogenetic analysis, variant analysis, GWAS, interpretation and original drafting; DP: population structure, LD decay and selective sweep analysis; ECH: rust infections of USA *Pca* populations, race assignments, DNA extractions, data visualisation; KM: data curation, GWAS methodology; SD, HNP, ESN, FL: USA rust collections and infections; BV, ZAP, WHPB: South Africa rust resources, curation, race assignments; MEM: USA rust collections; JS: supervision, effector prediction, drafting; SFK: conceptualisation, USA rust collection; EHS: conceptualisation, methodology; PND, conceptualisation, original drafting, supervision; MF: conceptualisation, project administration, supervision, original drafting, USA rust collection and infections. All authors contributed to editing and review of the manuscript.

## ACKNOWLEDGEMENTS

Thanks Drs Alex Whan and Shannon Dillon at CSIRO for helpful discussions regarding GWAS, as well as Jakob Riddle and Roger Caspers at the USDA-ARS for assistance with race assignments.

## SUPPLEMENTARY MATERIALS

**Figure S1.**
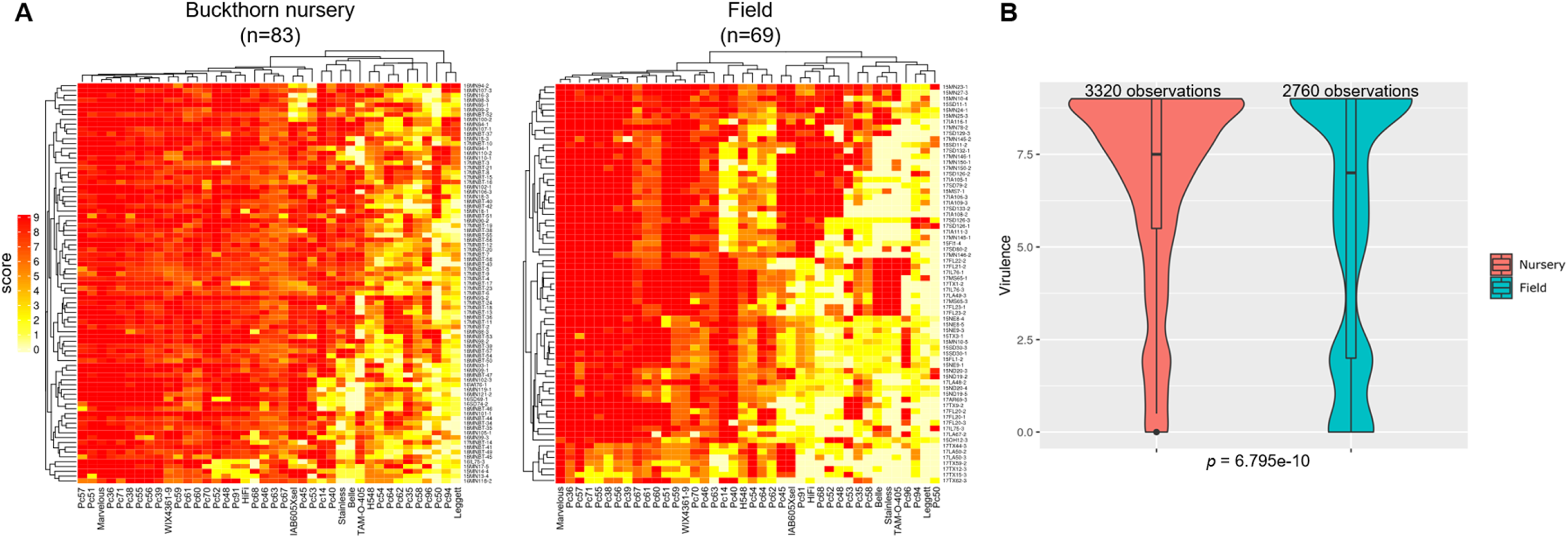
Comparison of virulence profiles of *Pca* isolates from buckthorn nursery and field in the USA. **(A)** Heatmaps to visualise virulence scores of *Pca* isolates (y-axis) on 40 oat differential lines (x-axis) comparing isolates from 2015 or later. High infection scores (red) indicating high virulence and lower scores (yellow/white) indicate avirulence. Order of *Pca* isolates is presented according to hierarchical clustering of virulence scores. **(B)** Violin plots of virulence score distribution in buckthorn nursery and field isolates. Above each plot number of observations (n=*Pca* isolate x oat differential line (*Pc* gene)) is defined. A *p*-value from a two-tailed Wilcoxon rank sum test is shown.

**Figure S2.**
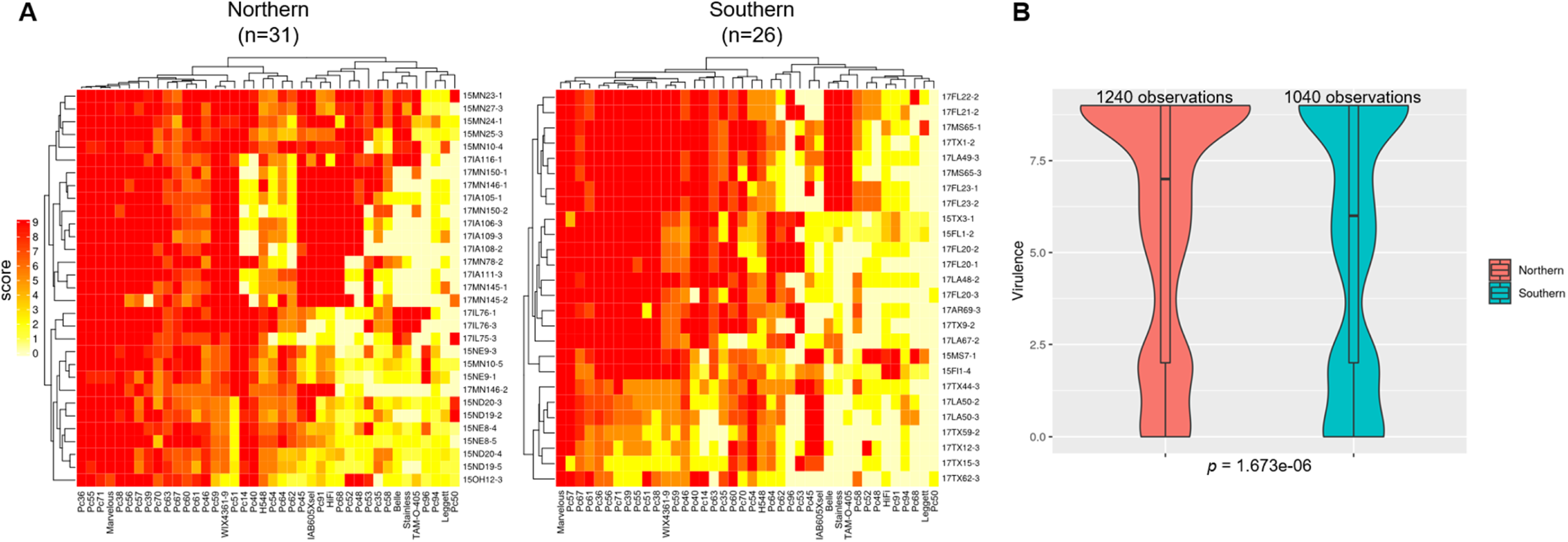
Comparison of virulence profiles of USA *Pca* isolates from northern and southern USA. **(A)** Heatmaps of virulence scores of USA *Pca* isolates on 40 oat differential lines comparing isolates collected in 2015 or later. High scores (red) indicating high disease virulence (disease susceptibility) and lower scores (yellow/white) indicate avirulence (disease resistance). Order of *Pca* isolates (y-axis) is presented according to hierarchical clustering of scores. **(B)** Data shown in heatmaps is summarised in violin plots, which depict distribution of all virulence scores for each group. Above each plot number of observations (n=*Pca* isolate x oat differential line (*Pc* gene)) is defined. A *p*-value from a Wilcoxon rank sum test is also shown.

**Figure S3.**
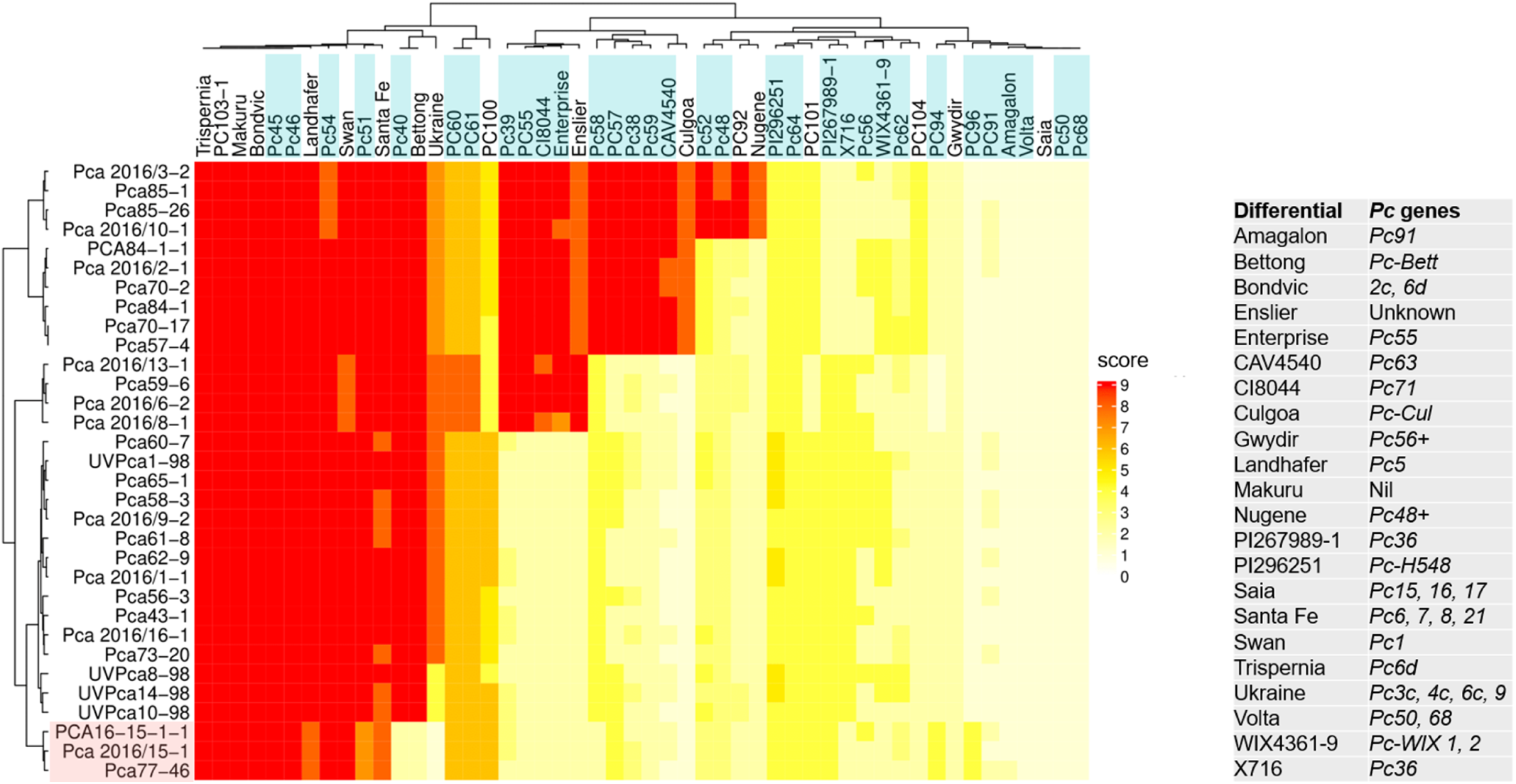
Virulence profile of *Pca* isolates from South Africa. Heatmap of virulence scores of *Pca* isolates (y-axis) on 50 oat differential lines used routinely in South Africa (x-axis). This oat differential set includes 32 postulated *Pc* genes in common with the USA differential set. High scores (red) indicating high disease virulence (disease susceptibility) and lower scores (yellow/white) indicate avirulence (disease resistance). Order of *Pca* isolates is presented according to hierarchical clustering of scores. *Pca* isolates highlighted in pink were collected from wild oats and the rest of isolates were collected from cultivated oats. Oat differential lines highlighted in blue show overlap with a *Pc* gene also included in the USA differential set. The table on the right lists postulated *Pc* genes in each cultivar used as part of the oat differential set of South Africa.

**Figure S4.**
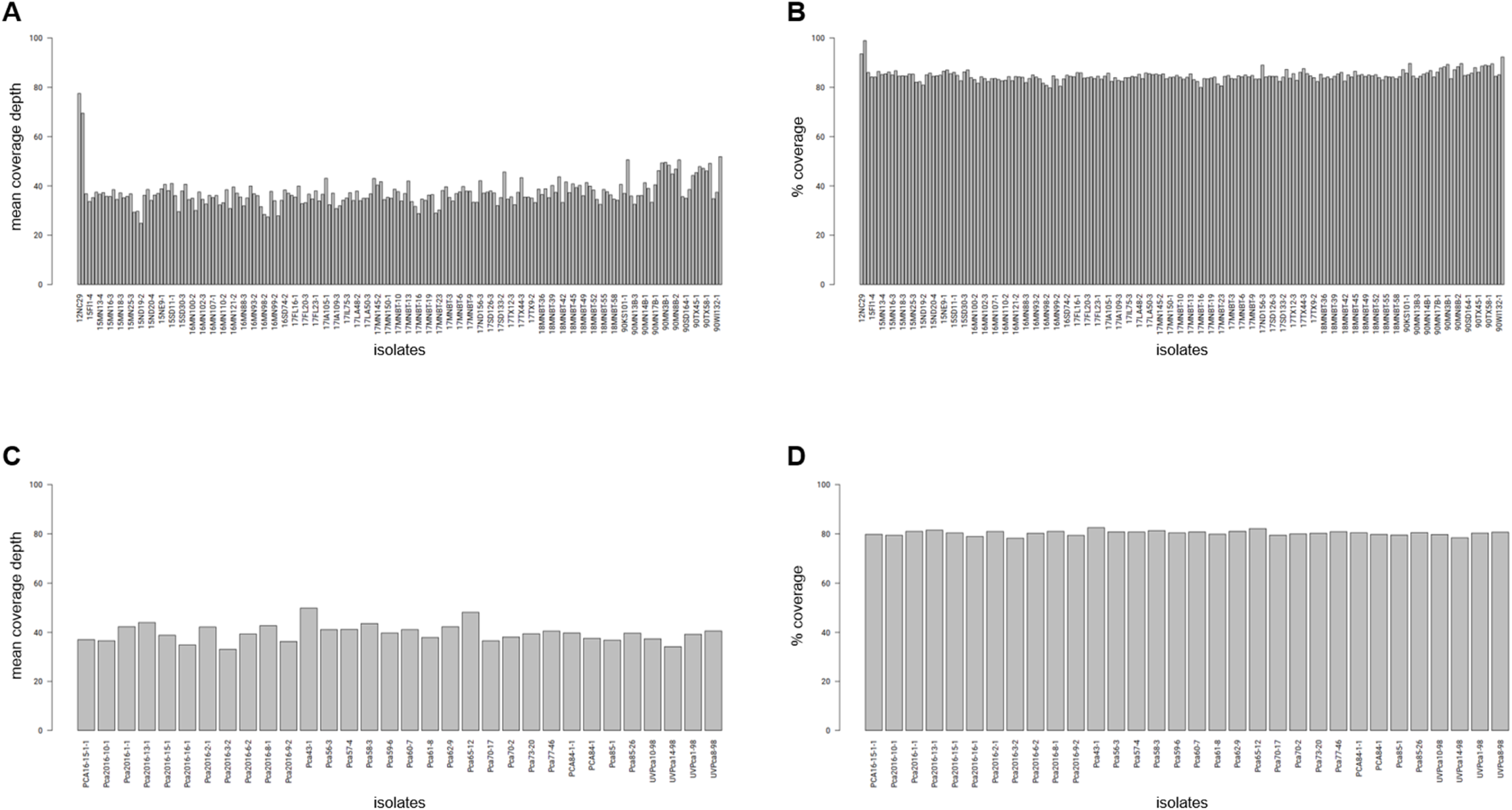
Sample coverage statistics of short-read alignments to *Pca* genome reference 12SD80 after removal of duplicate reads. **(A)** Mean overall coverage of *Pca* isolates from the USA in the y-axis. **(B)** Percent of total reference length with 10X or more coverage of isolates from the USA in the y-axis. All USA samples are shown but not all are labelled. **(C)** Mean overall coverage of South African *Pca* isolates in the y-axis. **(D)** Percent of total reference length with 10X or more coverage of South African isolates in the y-axis.

**Figure S5.**
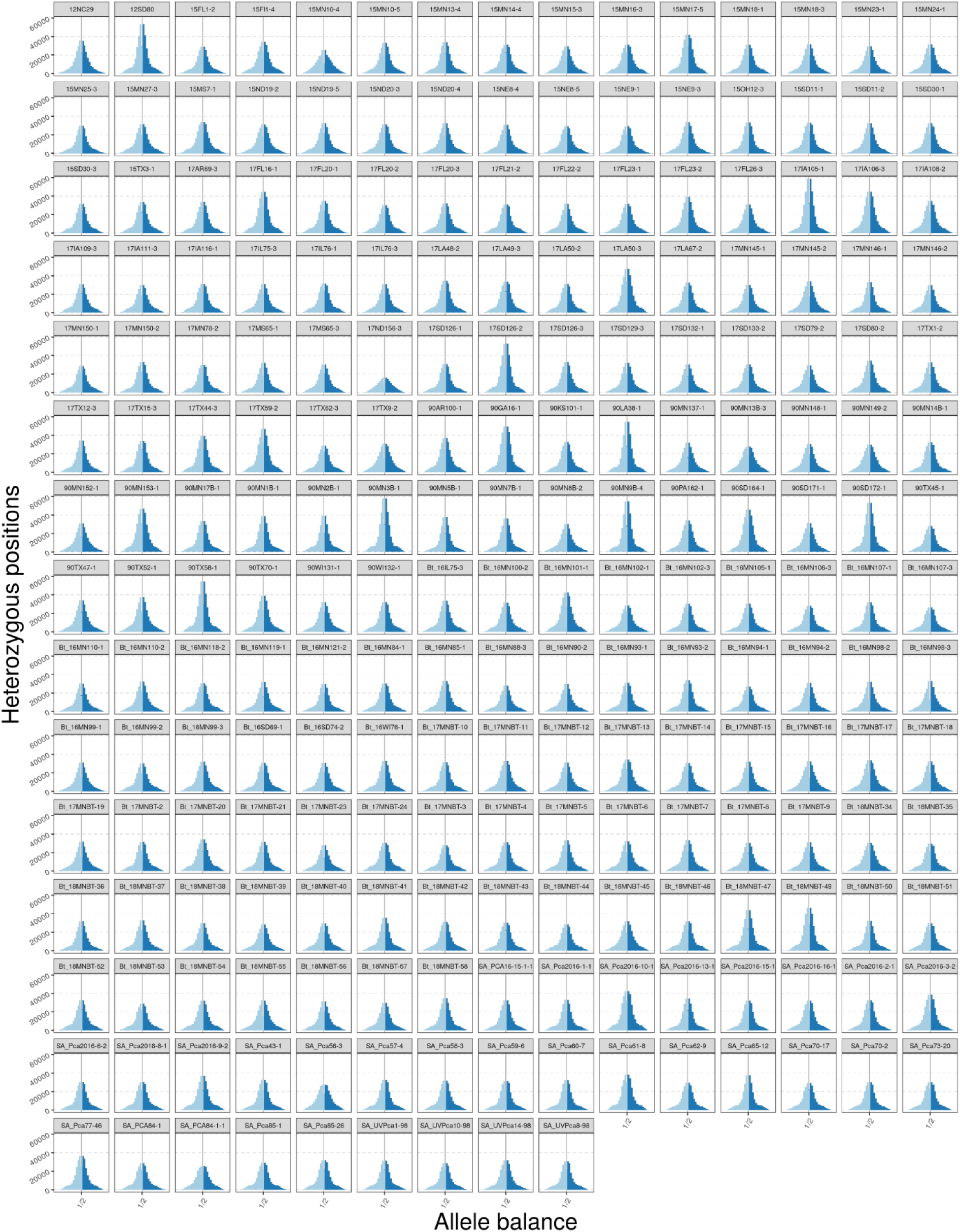
Allele balance at heterozygous positions to assess genotype contamination for USA and South Africa derived *Pca* isolates used in this study.

**Figure S6.**
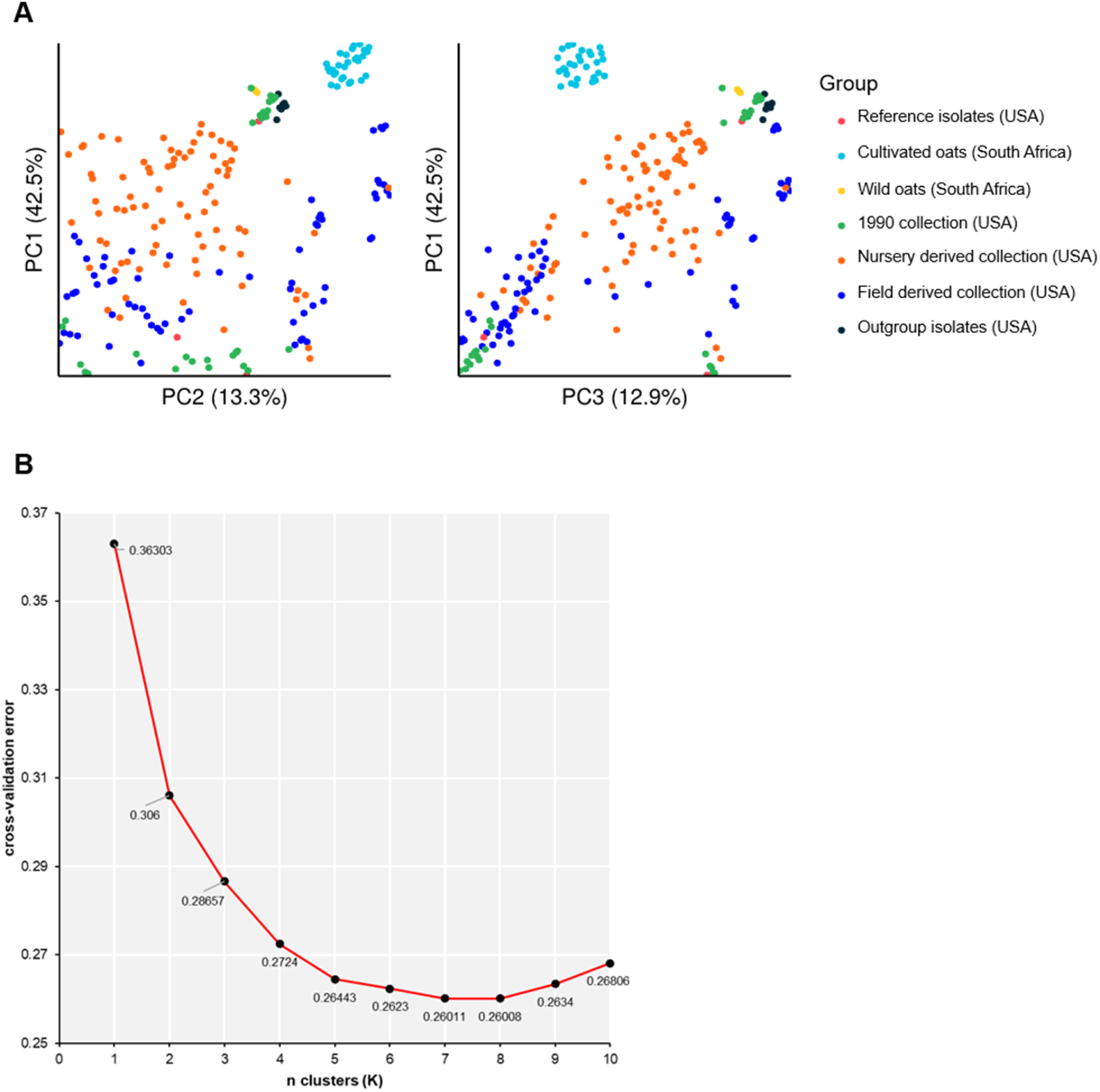
Cluster analysis of *Pca* isolates using SNPs called on 12SD80. **(A)** Principal component analysis (PCA) of *Pca* isolates coloured by different population groups. Analysis is based on linkage pruned SNPs called against 12SD80. All USA groups apart from “1990 collection” consist of isolates collected in 2015 or later. The left plot compares principal components 1 and 2. The right plot compares principal components 1 and 3. Contribution to variance indicated as percentages next to each principal component. **(B)** Elbow plot from K-means clustering showing K=8 as optimal based on the lowest cross-validation error to conduct membership analysis.

**Figure S7.**
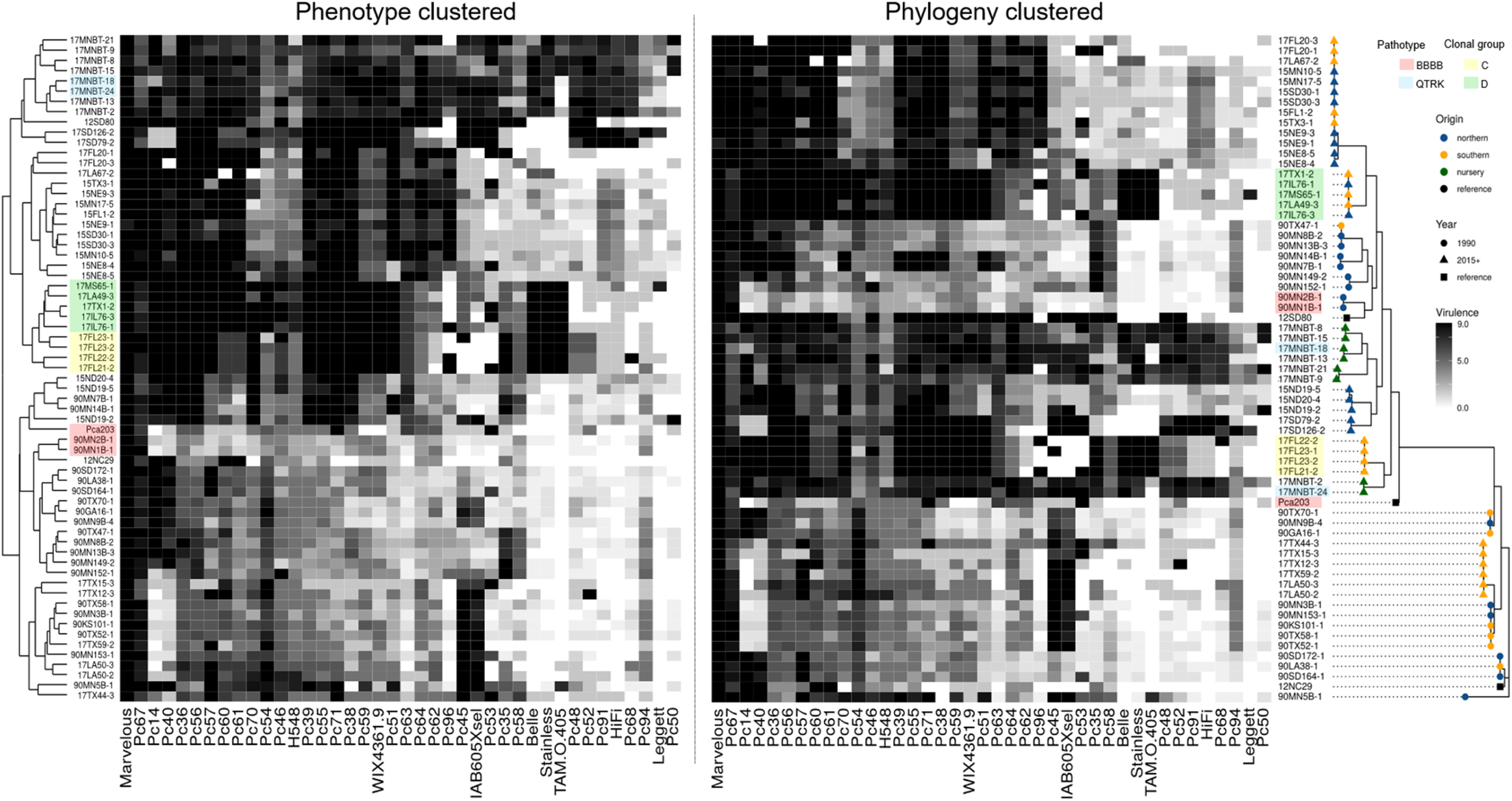
Clustering of a subset of USA *Pca* isolates according to phylogeny and virulence scores on the North American set of oat differential lines. Heatmaps of virulence scores of 65 USA *Pca* isolates derived from clonal groups comparing arrangement by virulence phenotype (left) or phylogeny (right). Isolates scored on 40 differential oat lines (x-axis). Infection scores were converted to a numeric scale (0 = resistance shown in white to 9 = susceptibility shown in black for heatmap generation. Right, ML phylogenetic tree based on 397,562 SNP variants called against 12SD80 (500 bootstraps) is shown for a subset of *Pca* isolates. Year and origin of collection is defined by shapes and colours, respectively, at the tip of the branches. *Pca* isolates with an existing genome reference are also indicated with a black square at the tip of the branch. Right, *Pca* isolates are clustering according to relationships solely based on virulence phenotypes. Blocks highlighted by pathotype demonstrate isolates of the same pathotype yet are phylogenetically distant. Similarly, blocks highlighted by clonal group demonstrate groups with highly similar virulence profiles but genetically distinct.

**Figure S8.**
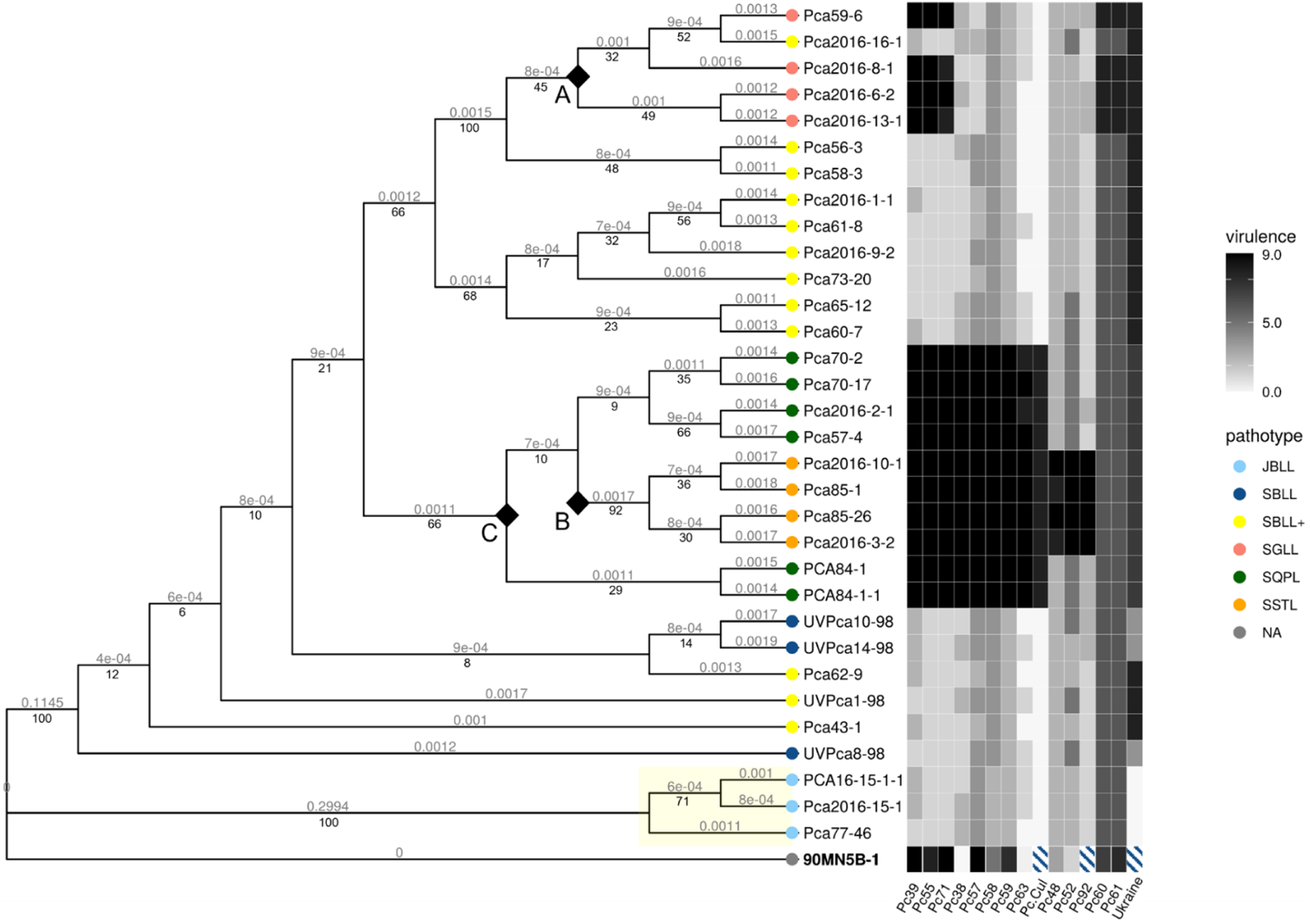
Phylogeny of South African *Pca* isolates next to virulence scores on a South African set of differential oat lines. ML phylogenetic tree based on 410,334 SNP variants called against 12SD80 (1000 bootstraps). Branch lengths in the diagram of ML were modified to easily visualise content. True branch lengths are indicated above each branch. Support values indicated below branches. The clade from wild oats is highlighted in yellow. Races are indicated by tip colour and defined on the right of the figure. Key branch points distinguishing postulated virulence mutations are indicated by black diamonds and labelled A, B and C. ML phylogenetic tree was rooted on USA isolate 90MN5B-1. Race for 90MN5B-1 (race SGML) is not shown because virulence scores of 90MN5B-1 were based on separate USA differential oat set. A heatmap is shown to illustrate virulence profiles of isolates on the right. Infection scores were converted to a numeric scale (0 = resistance shown in white to 9 = susceptibility shown in black) for heatmap generation. Striped boxes indicate unscored values for 90MN5B-1 as virulence profile of this isolate was acquired using the USA oat differential set.

**Figure S9.**
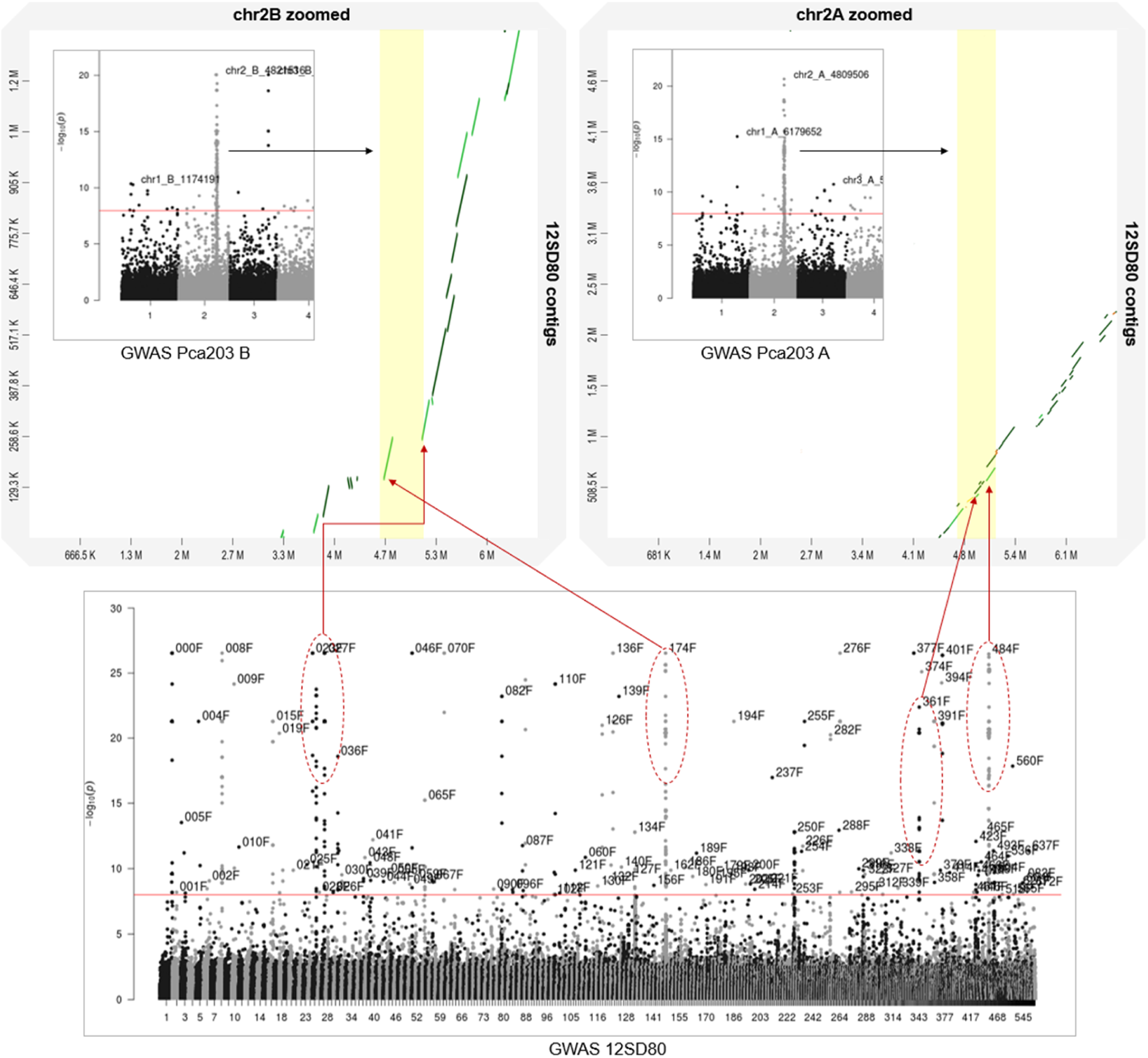
Improved resolution of GWAS associations on the oat differential line Pc71. A comparison of virulence genomic intervals (VGIs) between Pca203 and 12SD80 is shown. Association peaks in the 12SD80 genome reference (bottom panel) and their corresponding contigs map to a single VGI on either haplotype of Pca203 (top panel, A or B haplotypes) according to chromosomes are presented as a zoomed region of the Manhattan plots. VGIs on Pca203 highlighted in yellow on the alignment dot plots indicating syntenic regions. Lighter green segments show alignments with 50% or more identity while dark green segments show alignments with 75% or more identity.

**Figure S10.**
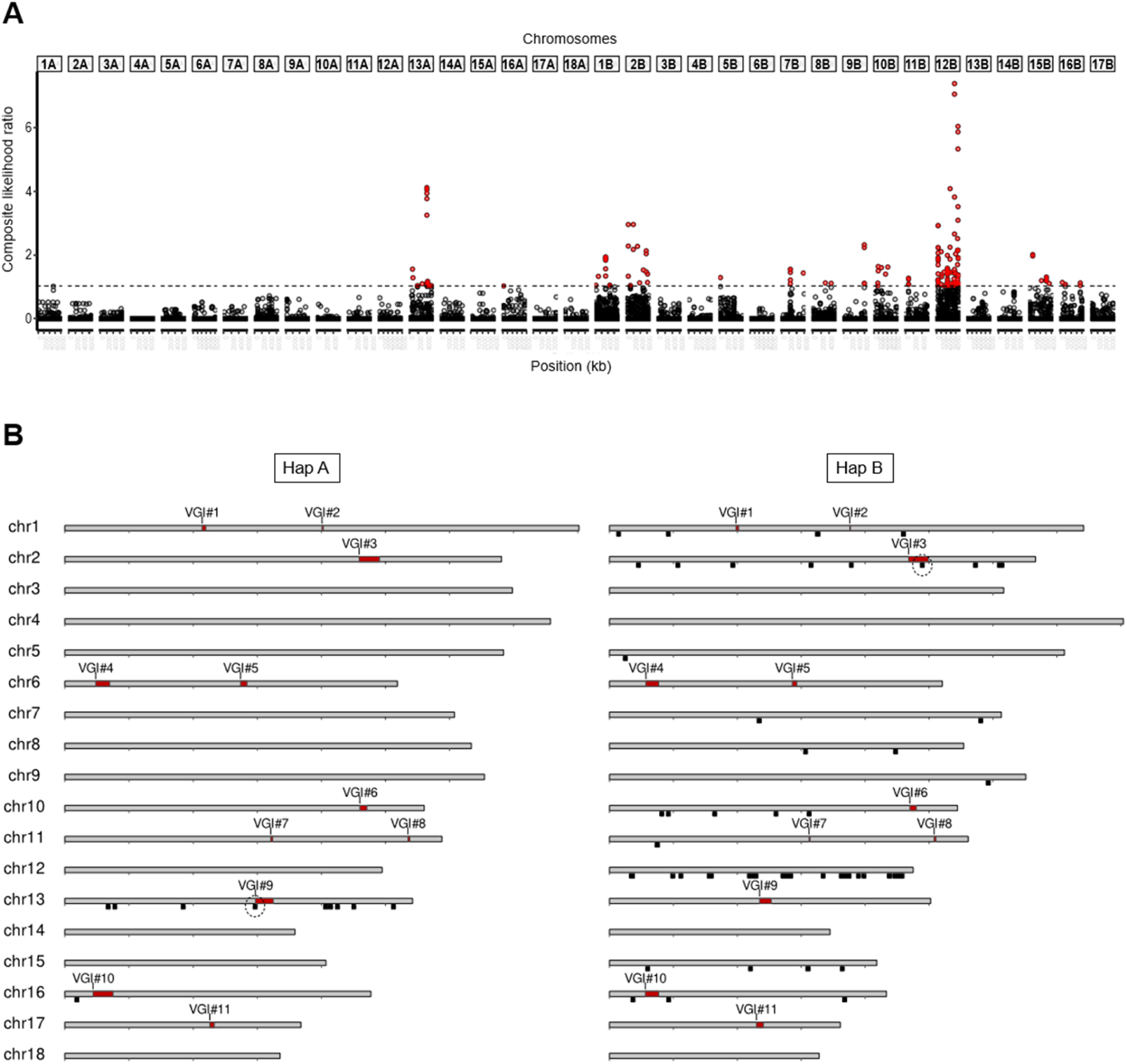
Results of selective sweep analysis in USA *Pca* isolates. **(A)** Selective sweep signals of nursery isolates versus field isolates plotted against the Pca203 genome using 372,275 SNP markers. The x-axis denotes the chromosome (labelled along top) and position, and the y-axis shows the composite likelihood ratio (CLR) evaluated by SweeD. Markers exceeding a CLR of 1.0 (dashed line) are highlighted red. Each point represents an interval of 1kb. **(B)** Pca203 chromosomes of haplotypes A (left) and B (right) displaying virulence-associated genomic intervals (VGIs) determined from GWAS indicated as red regions. Black bars indicate regions with selective sweep signal from USA buckthorn nursery derived isolates. Regions where a selective sweep overlaps a VGI are circled.

**Table S1.** Primary contigs and haplotigs from 12SD80 and 12NC29 that are syntenic with virulence-associated genomic intervals (VGIs) in Pca203.

| VGI # | Interval (length Kbp) |  | orthotigs |  |
| --- | --- | --- | --- | --- |
|  | A genome | B genome | 12SD80 | 12NC29 |
| 1 | chr1A:<br>2.14-2.20Mb<br>(60) | chr1B:<br>1.98-2.021Mb<br>(40) | 000036F_006, 000036F,<br>000036F_004 | 000176F, 000176F_002 |
| 2 | chr1A:<br>4.01-4.03Mb<br>(20) | chr1B:<br>3.76-3.773Mb<br>(10) | 000031F, 000031F_002 | 000007F, 000429F |
| 3 | chr2A:<br>4.59-4.91Mb<br>(320) | chr2B:<br>4.68-4.99Mb<br>(310) | 000651F, 000174F_005,<br>000174F, 000174F_002,<br>000484F, 000361F | 000014F, 000268F_001,<br>000359F_002,<br>000359F_001,<br>000032F_001, 000707F,<br>000268F, 000359F |
| 4 | chr6A:<br>0.48-0.70Mb<br>(220) | chr6B:<br>0.57-0.77Mb<br>(200) | 000065F_004, 000065F,<br>000065F_003 | 000020F, 000020F_001 |
| 5 | chr6A:<br>2.74-2.84Mb<br>(100) | chr6B:<br>2.86-2.93Mb<br>(70) | 000082F, 000082F_005 | 000300F_001, 000300F,<br>000344F_001 |
| 6 | chr10A:<br>4.60-4.71Mb<br>(110) | chr10B:<br>4.70-4.802Mb<br>(100) | 000647F, 000260F_002,<br>000153F, 000153F_001,<br>000153F_004,<br>000301F_003, 000301F | 000637F_001, 000024F,<br>000024F_002 |
| 7 | chr11A:<br>3.21-3.24Mb<br>(30) | chr11B:<br>3.12-3.14Mb<br>(20) | 000240F, 000240F_002 | 000082F |
| 8 | chr11A:<br>5.35-5.38Mb<br>(30) | chr11B:<br>5.08-5.11Mb<br>(30) | 000016F, 000016F_002 | 000006F, 000006F_002 |
| 9 | chr13A:<br>2.97-3.25Mb<br>(280) | chr13B:<br>2.349-2.53Mb<br>(180) | 000197F, 000197F_001,<br>000155F_004,<br>000155F_001, 000155F,<br>000155F_003,<br>000155F_002,<br>000155F_005 | 000375F, 000431F,<br>000336F, 000582F,<br>000582F_001 |
| 10 | chr16A:<br>0.435-0.75Mb<br>(310) | chr16B:<br>0.56-0.77Mb<br>(210) | 000133F, 000134F,<br>000250F_001, 000250F | 000126F, 000512F,<br>000647F_001, 000647F,<br>000566F, 000310F,<br>000310F_001, 000360F |
| 11 | chr17A:<br>2.26-2.33Mb<br>(70) | chr17B:<br>2.30-2.41Mb<br>(110) | 000316F_002, 000316F,<br>000107F_004 | 000314F_001, 000314F |

