## Supplementary Data S2 for "Genome-enabled analysis of population dynamics and virulence associated loci in the oat crown rust fungus *Puccinia coronata* f. sp. *avenae*"

**Supplementary Data S2**. Results from genome-wide association study (GWAS) for virulence of North American *Puccinia coronata* f. sp. *avenae* (*Pca*) isolates towards oat differential lines using multiple *Pca* reference assemblies (12SD80, 12NC29, Pca203 diploid chromosomes, Pca203 haplotype A chromosomes, Pca203 haplotype B chromosomes).

**Fig 1A. quantile-quantile plots of SNP association values for virulence of *Pca* to oat line Belle.** Red lines indicate expected P-value distributions if P-values follow a uniform distribution (null hypothesis).

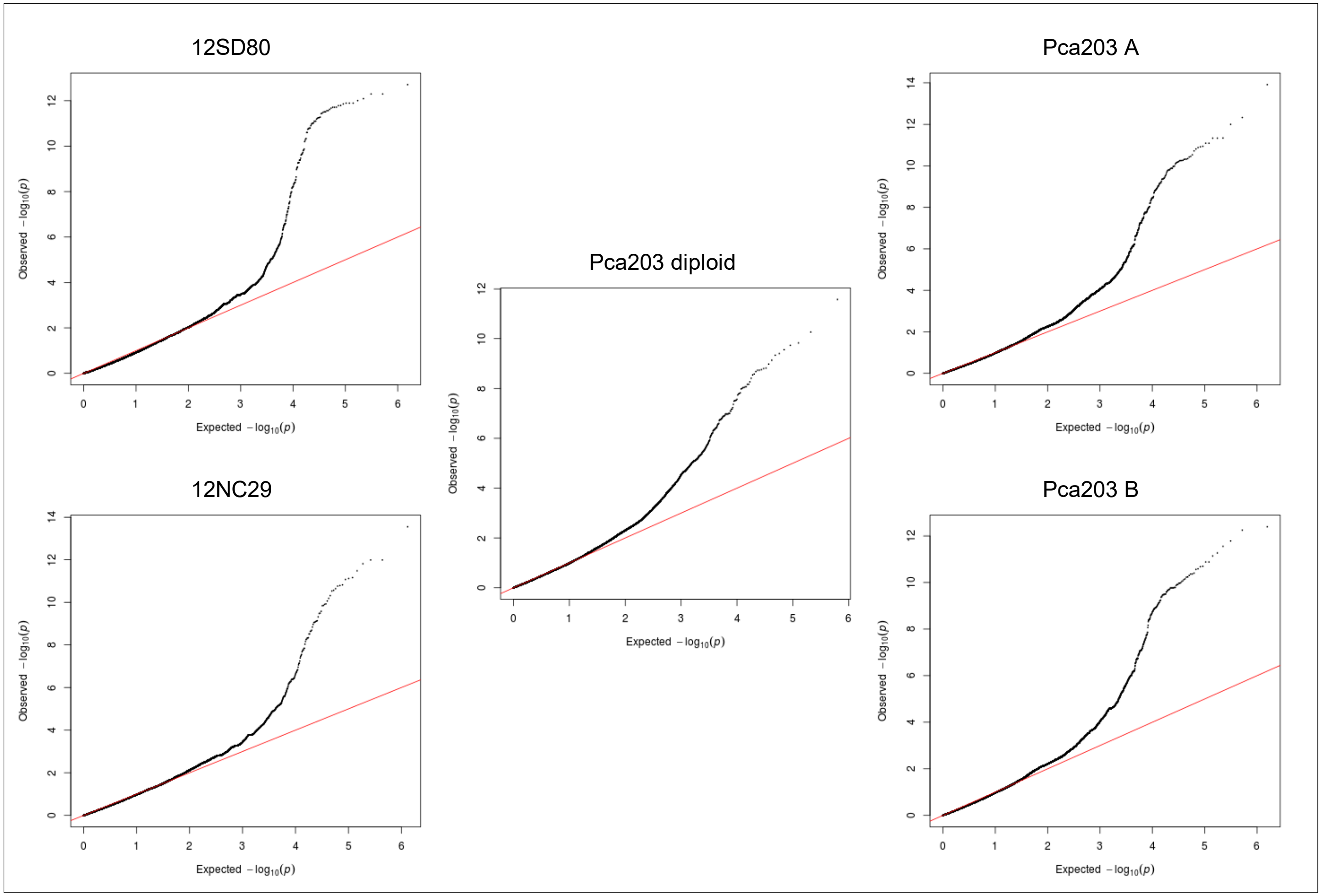

**Fig 1B. *Pca* isolate virulence score distribution and Manhattan plots showing SNP association values for virulence to oat line Belle.** Red and blue horizontal lines denote Bonferroni significance threshold (α = 0.01/total number of markers) and 5% false discovery rate threshold, respectively. Significant association peaks are labelled with contig or chromosome number and assigned VGI number.

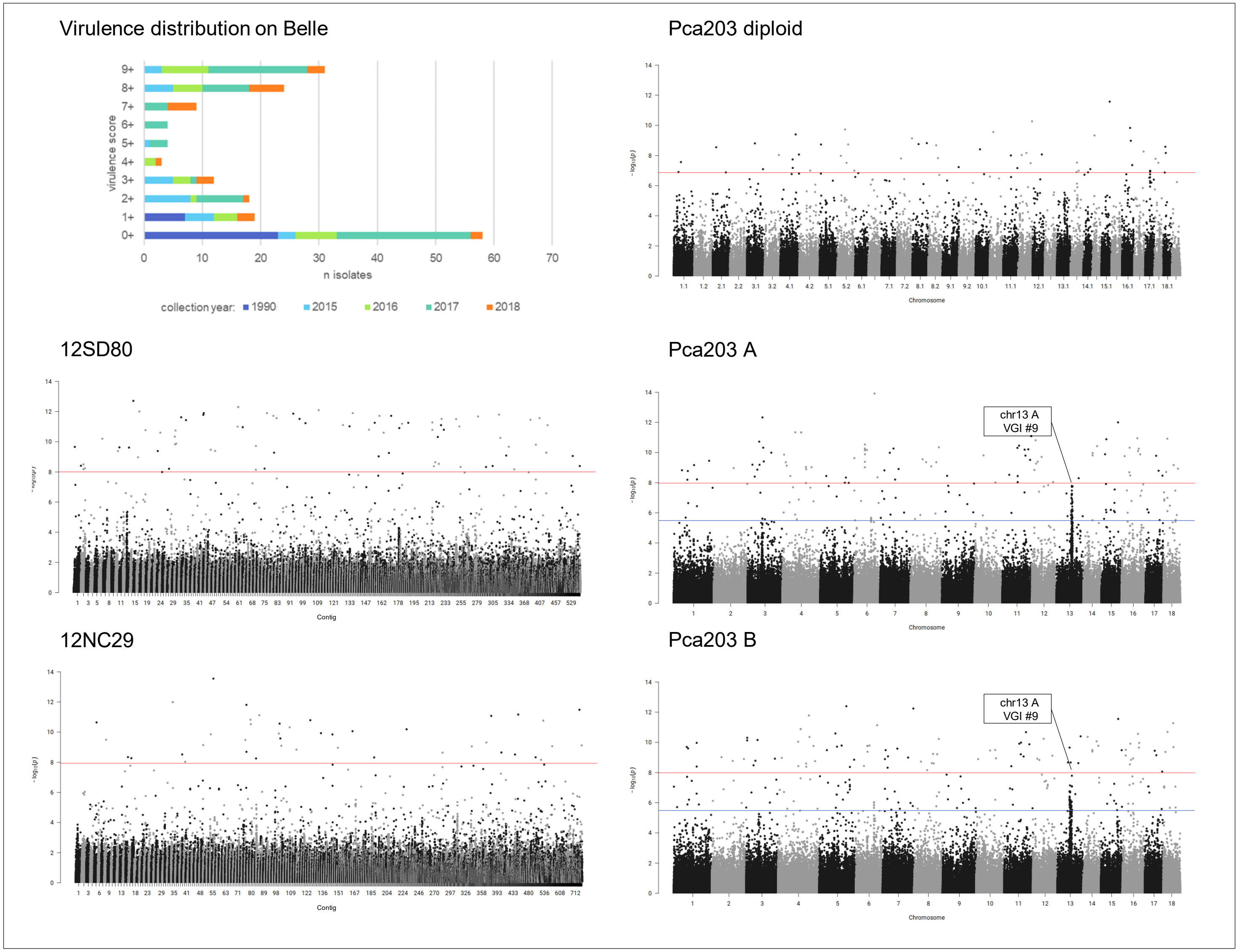

**Fig 2A. quantile-quantile plots of SNP association values for virulence of *Pca* to oat line H548.** Red lines indicate expected P-value distributions if P-values follow a uniform distribution (null hypothesis).

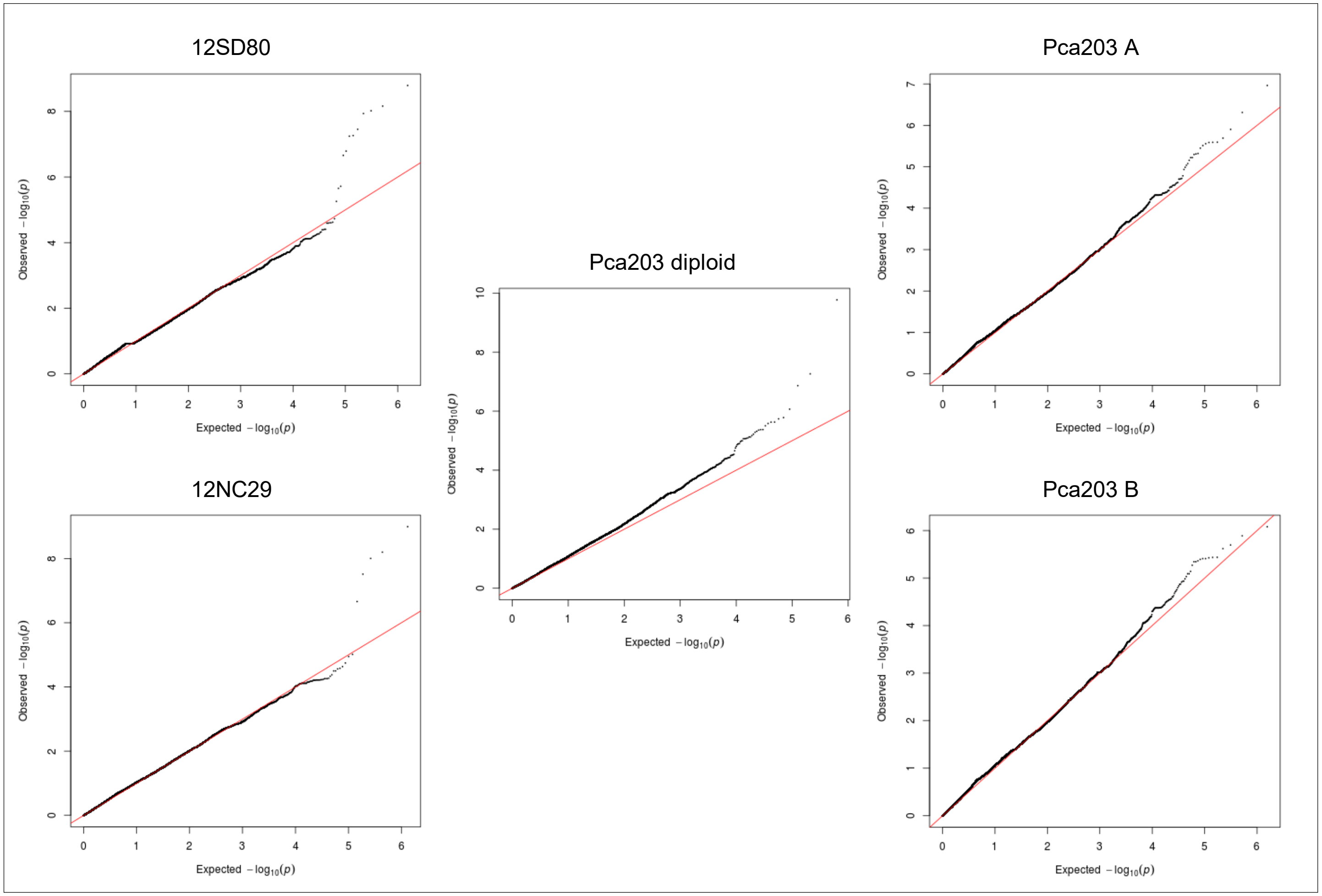

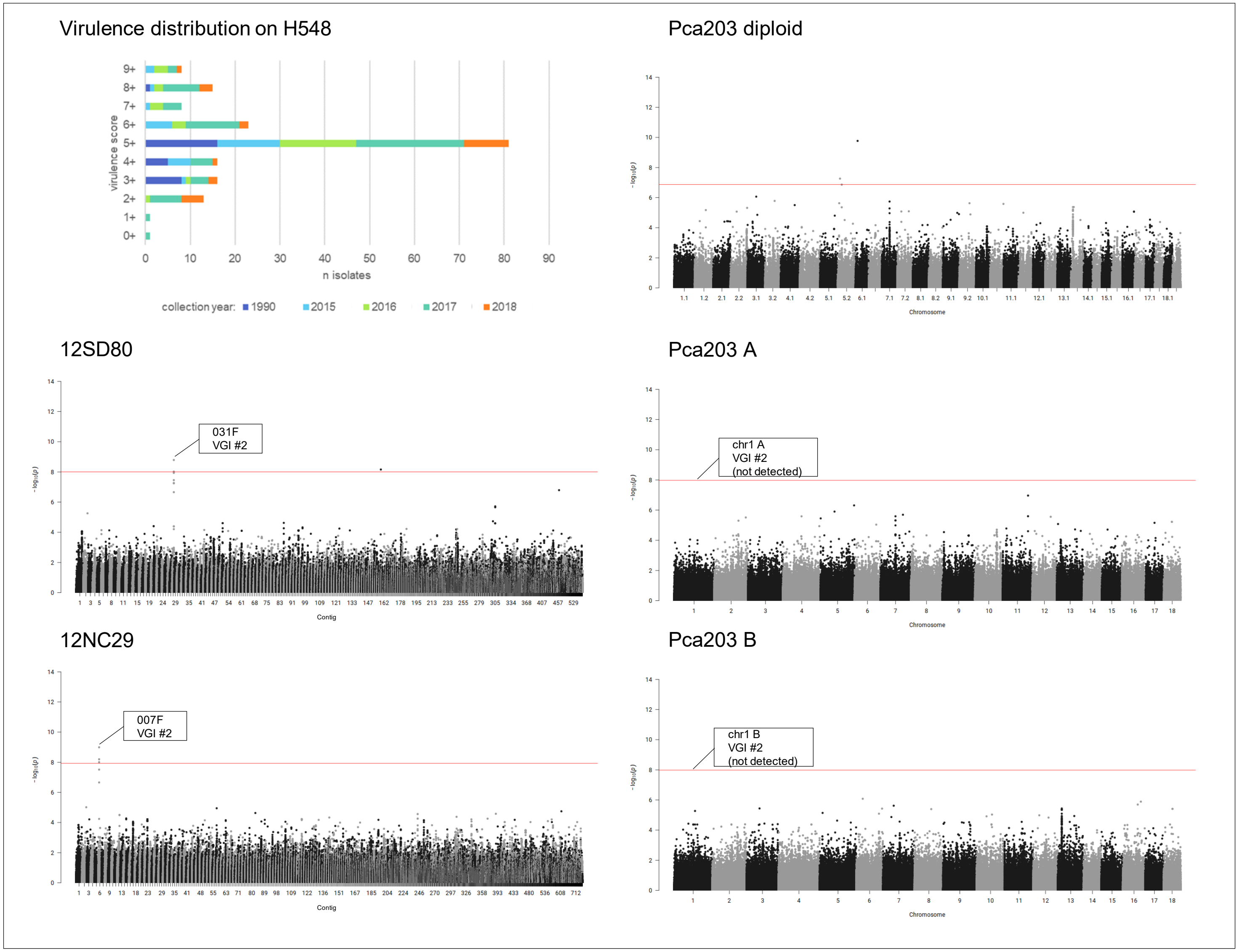


**Fig 2B. *Pca* isolate virulence score distribution and Manhattan plots showing SNP association values for virulence to oat line H548.** Red horizontal lines denote Bonferroni significance threshold (α = 0.01/total number of markers). Significant association peaks in 12SD80 and 12NC29 are labelled with contig number and assigned VGI number while labels in Pca203 A and B indicate corresponding homologous regions.

**Fig 3A. quantile-quantile plots of SNP association values for virulence of *Pca* to oat line HiFi.** Red lines indicate expected P-value distributions if P-values follow a uniform distribution (null hypothesis).

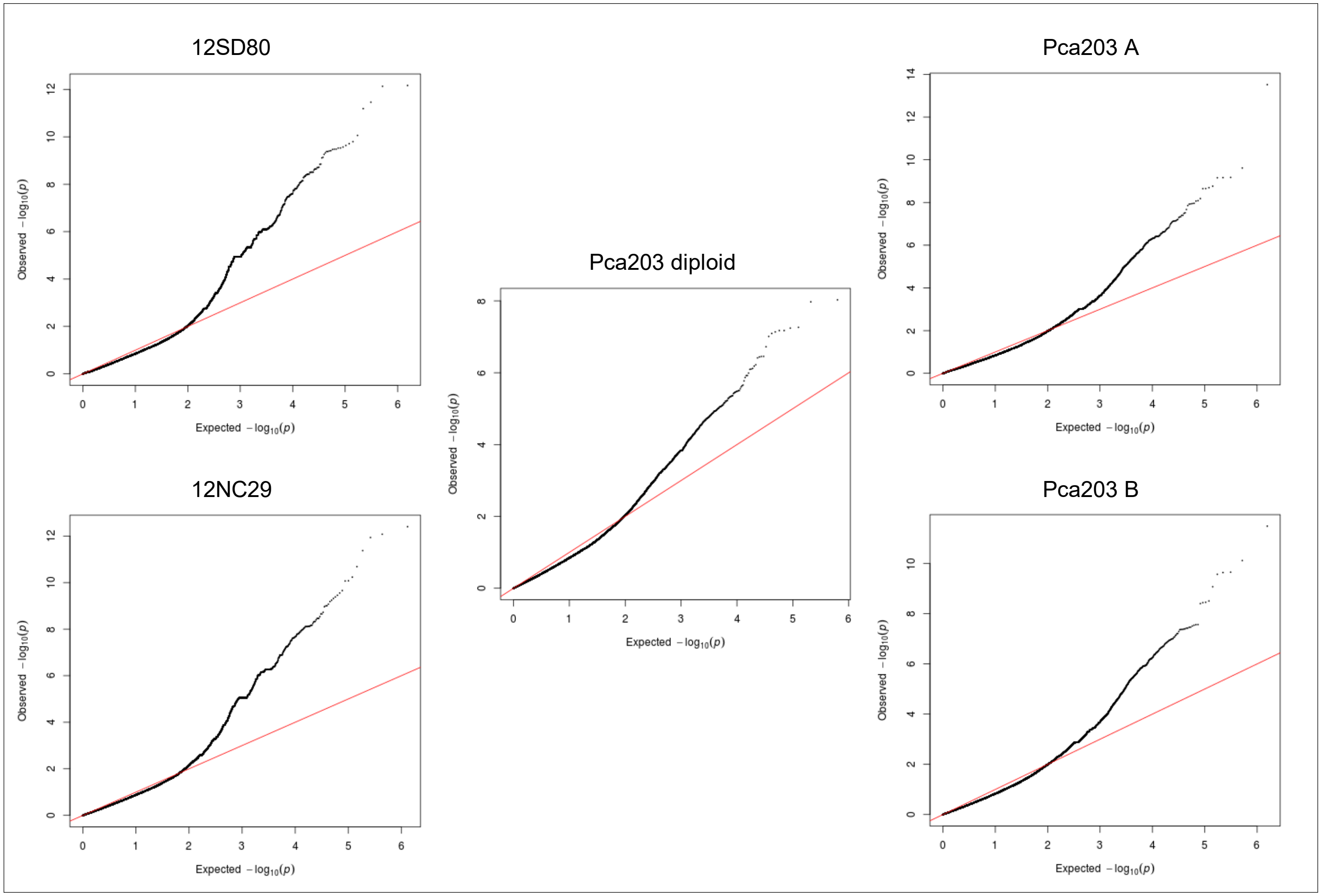

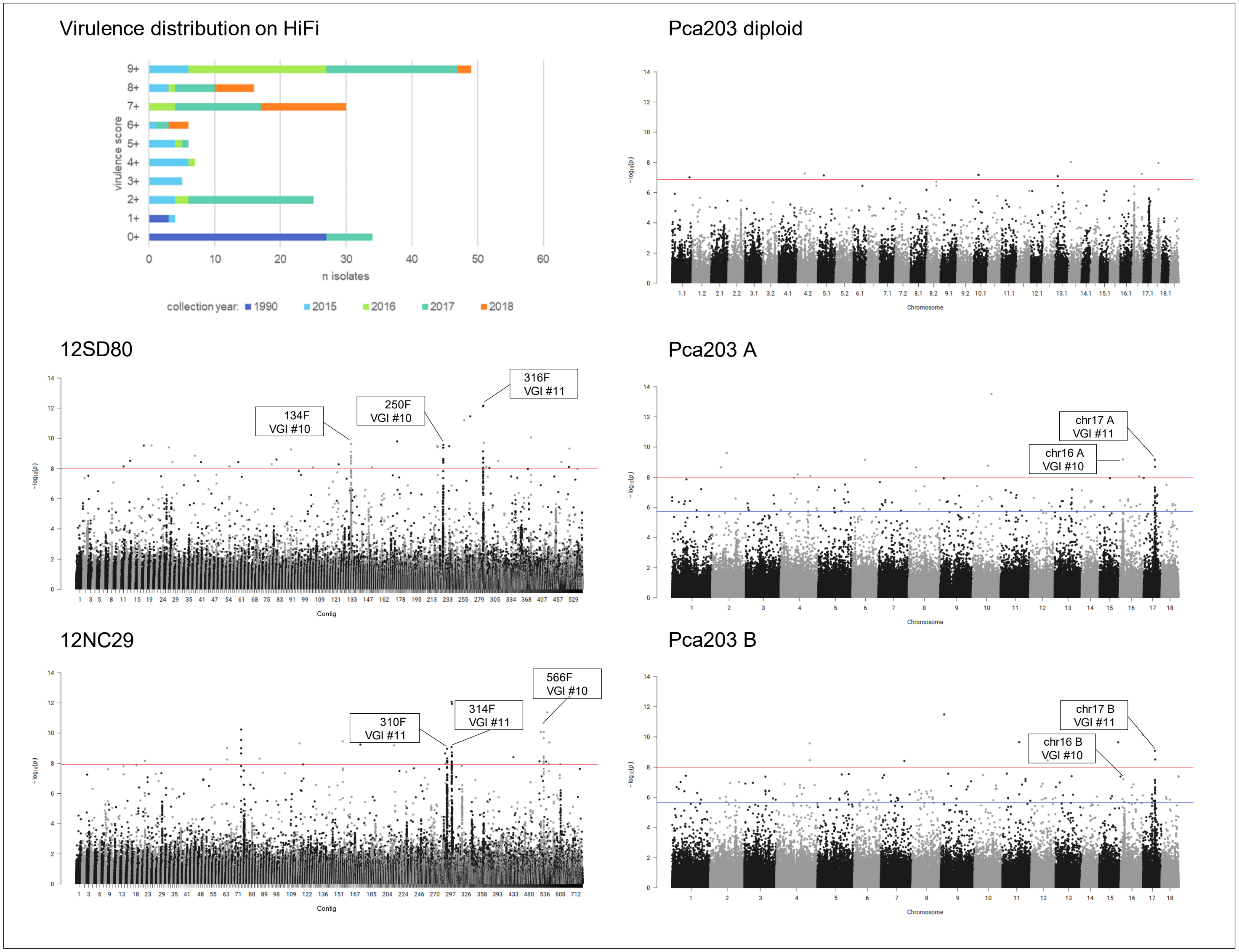


**Fig 3B. *Pca* isolate virulence score distribution and Manhattan plots showing SNP association values for virulence to oat line HiFi.** Red and blue horizontal lines denote Bonferroni significance threshold (α = 0.01/total number of markers) and 5% false discovery rate threshold, respectively. Significant association peaks are labelled with contig or chromosome number and assigned VGI number.

**Fig 4A. quantile-quantile plots of SNP association values for virulence of *Pca* to oat line IAB605Xsel.** Red lines indicate expected P-value distributions if P-values follow a uniform distribution (null hypothesis).

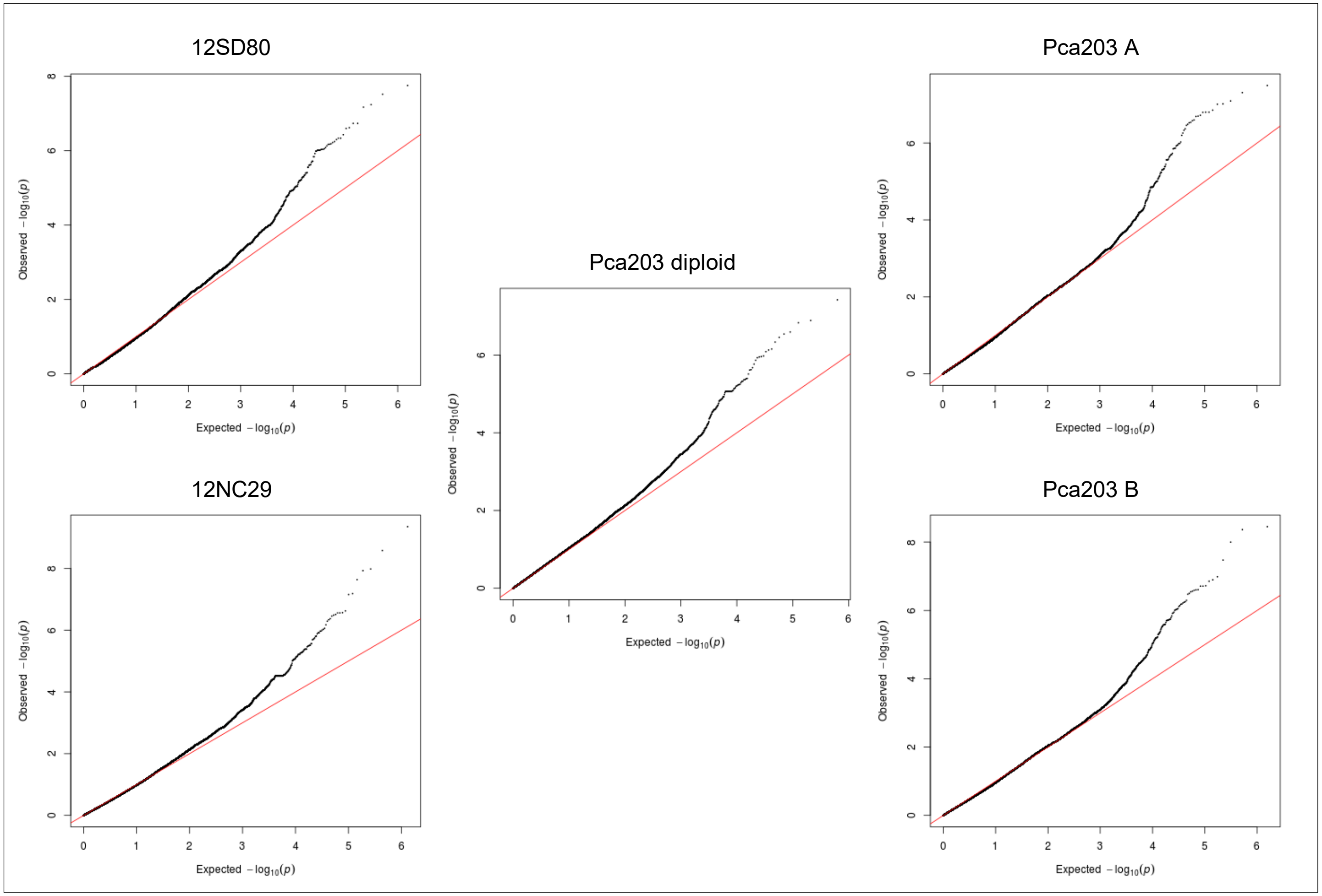

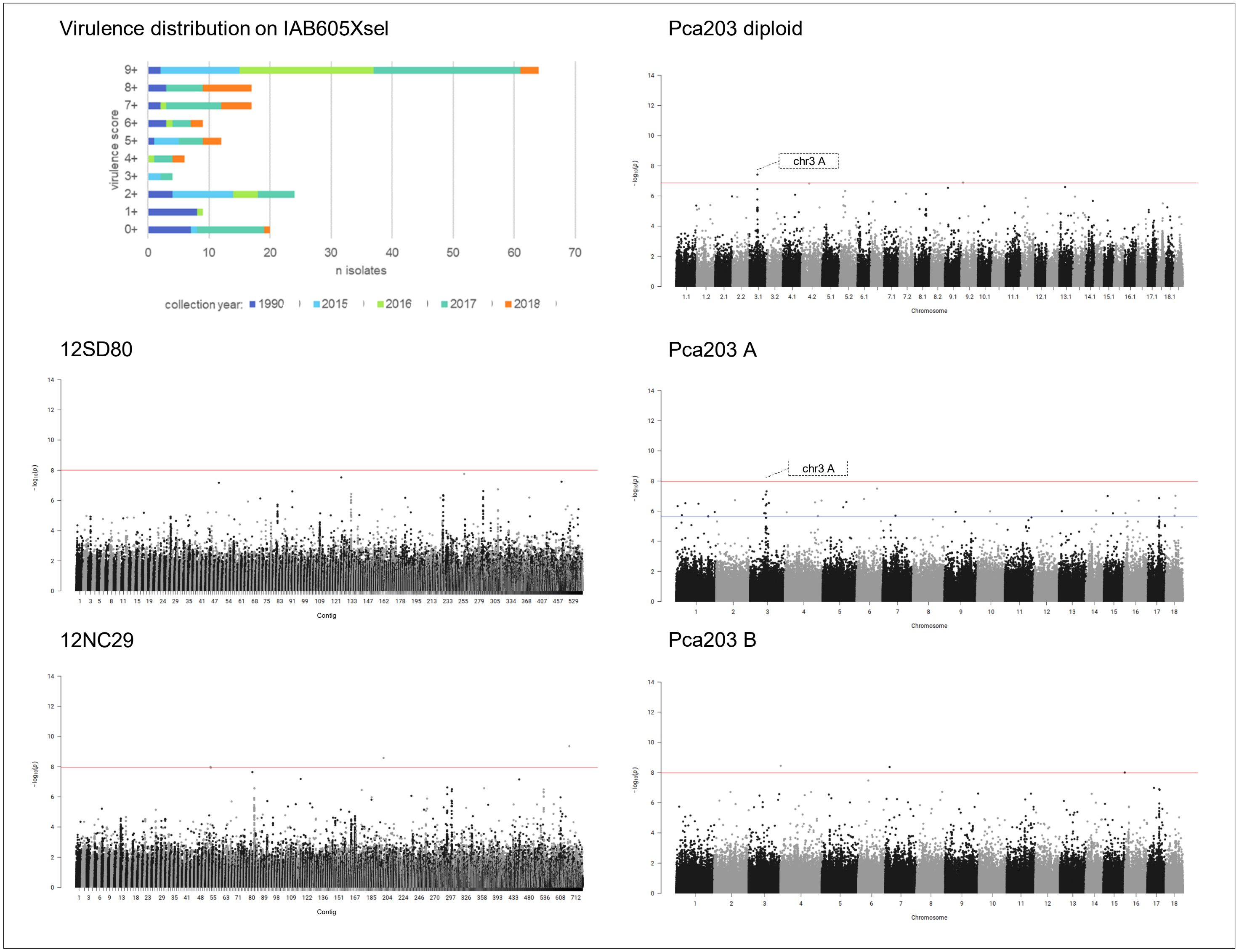


**Fig 4B. *Pca* isolate virulence score distribution and Manhattan plots showing SNP association values for virulence to oat line IAB605Xsel.** Red and blue horizontal lines denote Bonferroni significance threshold (α = 0.01/total number of markers) and 5% false discovery rate threshold, respectively. Significant association peaks are labelled with contig or chromosome number and assigned VGI number.

**Fig 5A. quantile-quantile plots of SNP association values for virulence of *Pca* to oat line Leggett.** Red lines indicate expected P-value distributions if P-values follow a uniform distribution (null hypothesis).

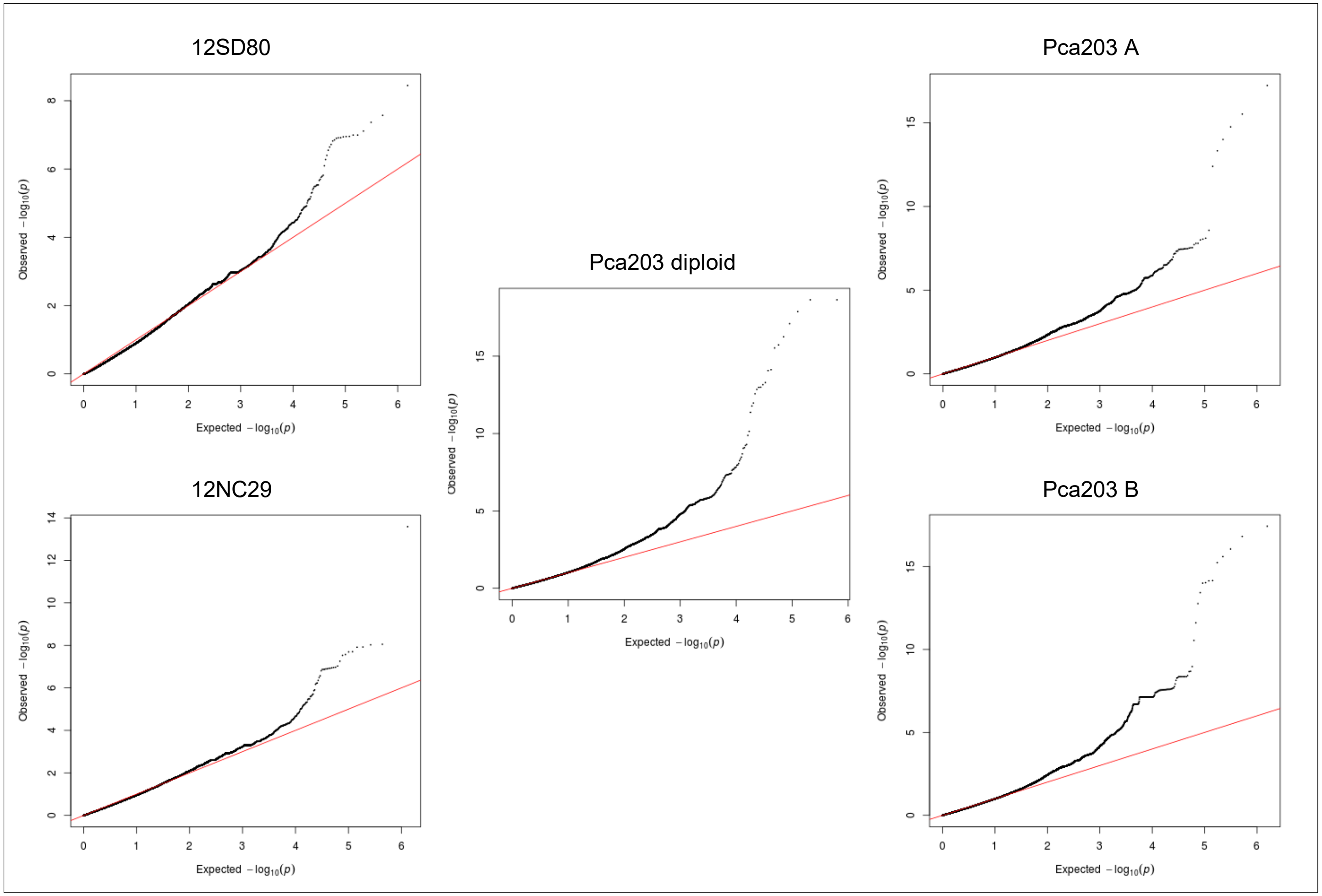

**Fig 5B. *Pca* isolate virulence score distribution and Manhattan plots showing SNP association values for virulence to oat line Leggett.** Red horizontal lines denote Bonferroni significance threshold (α = 0.01/total number of markers).

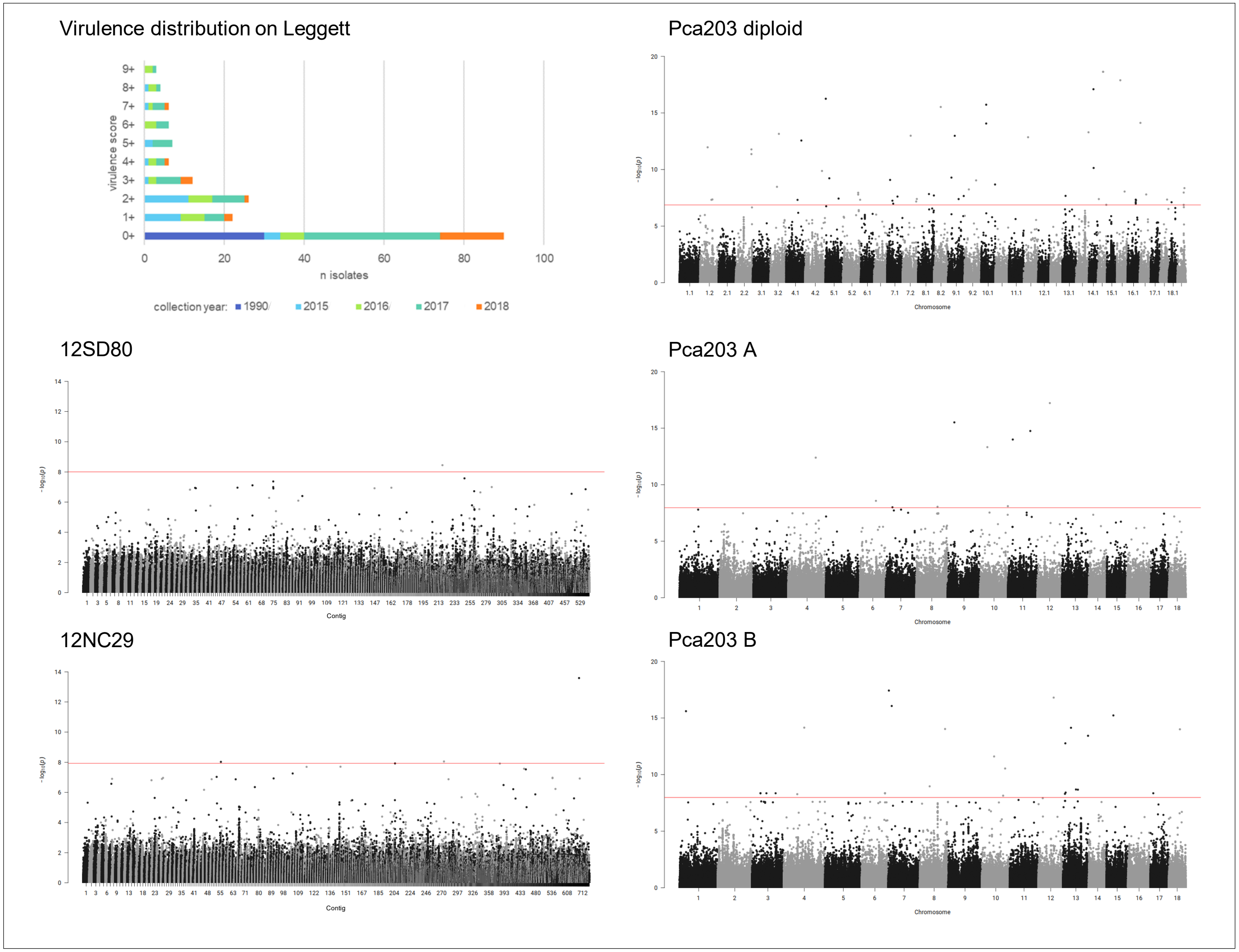

**Fig 6A. quantile-quantile plots of SNP association values for virulence of *Pca* to oat line Marvelous.** Red lines indicate expected P-value distributions if P-values follow a uniform distribution (null hypothesis).

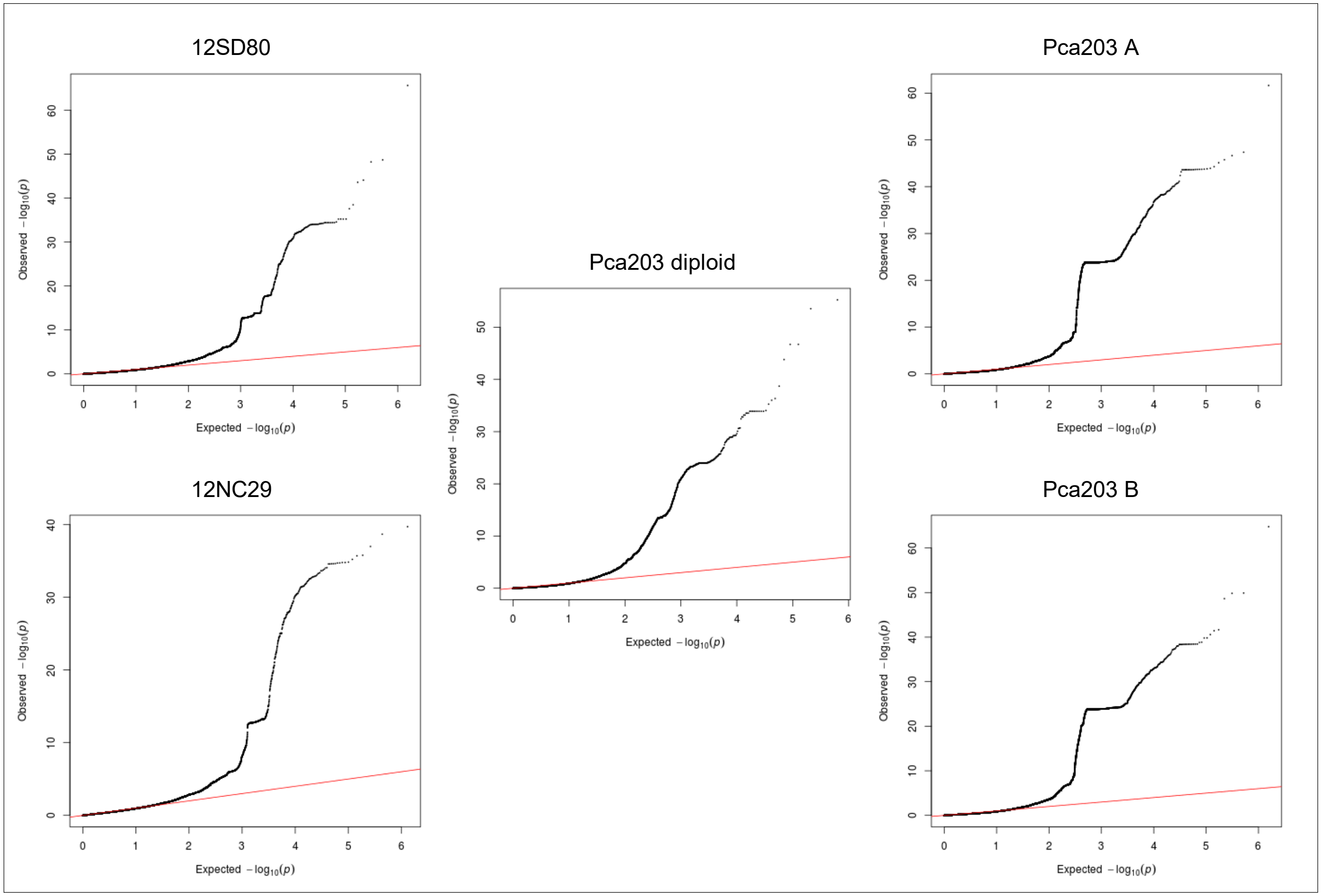

**Fig 6B. *Pca* isolate virulence score distribution and Manhattan plots showing SNP association values for virulence to oat line Marvelous.** Red horizontal lines denote Bonferroni significance threshold (α = 0.01/total number of markers).

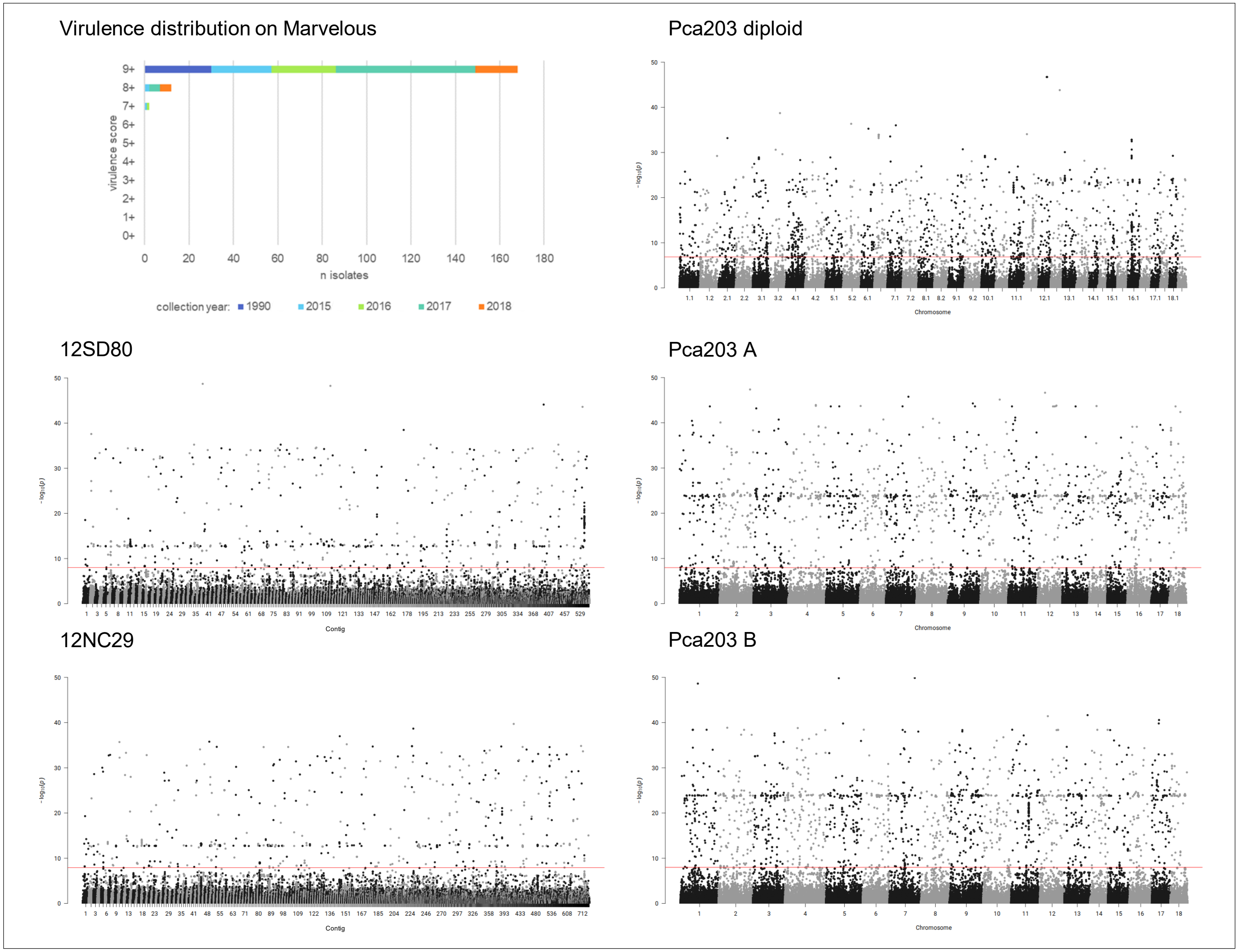

**Fig 7A. quantile-quantile plots of SNP association values for virulence of *Pca* to oat line *Pc14*.** Red lines indicate expected P-value distributions if P-values follow a uniform distribution (null hypothesis).

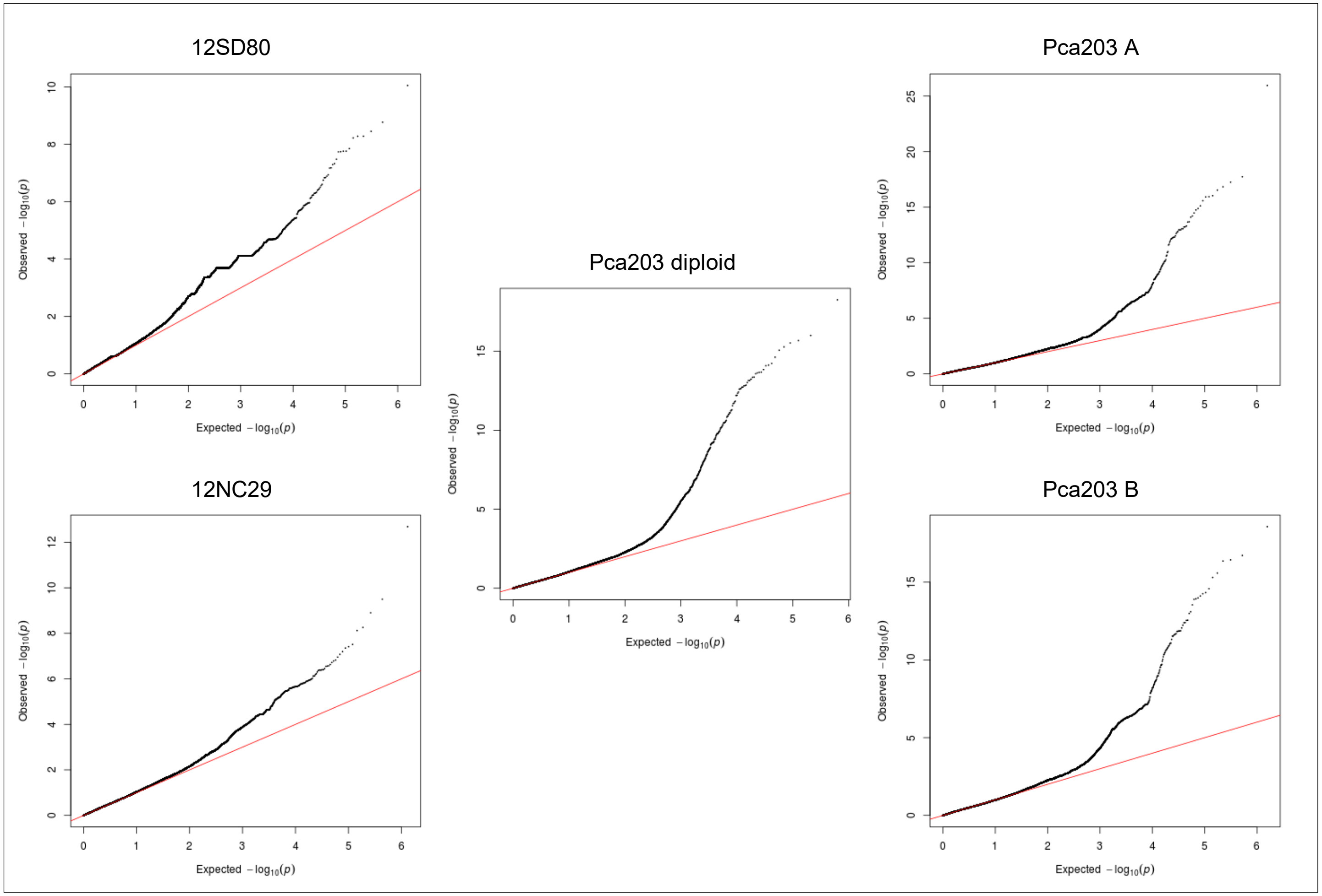

**Fig 7B. *Pca* isolate virulence score distribution and Manhattan plots showing SNP association values for virulence to oat line *Pc14*.** Red horizontal lines denote Bonferroni significance threshold (α = 0.01/total number of markers).

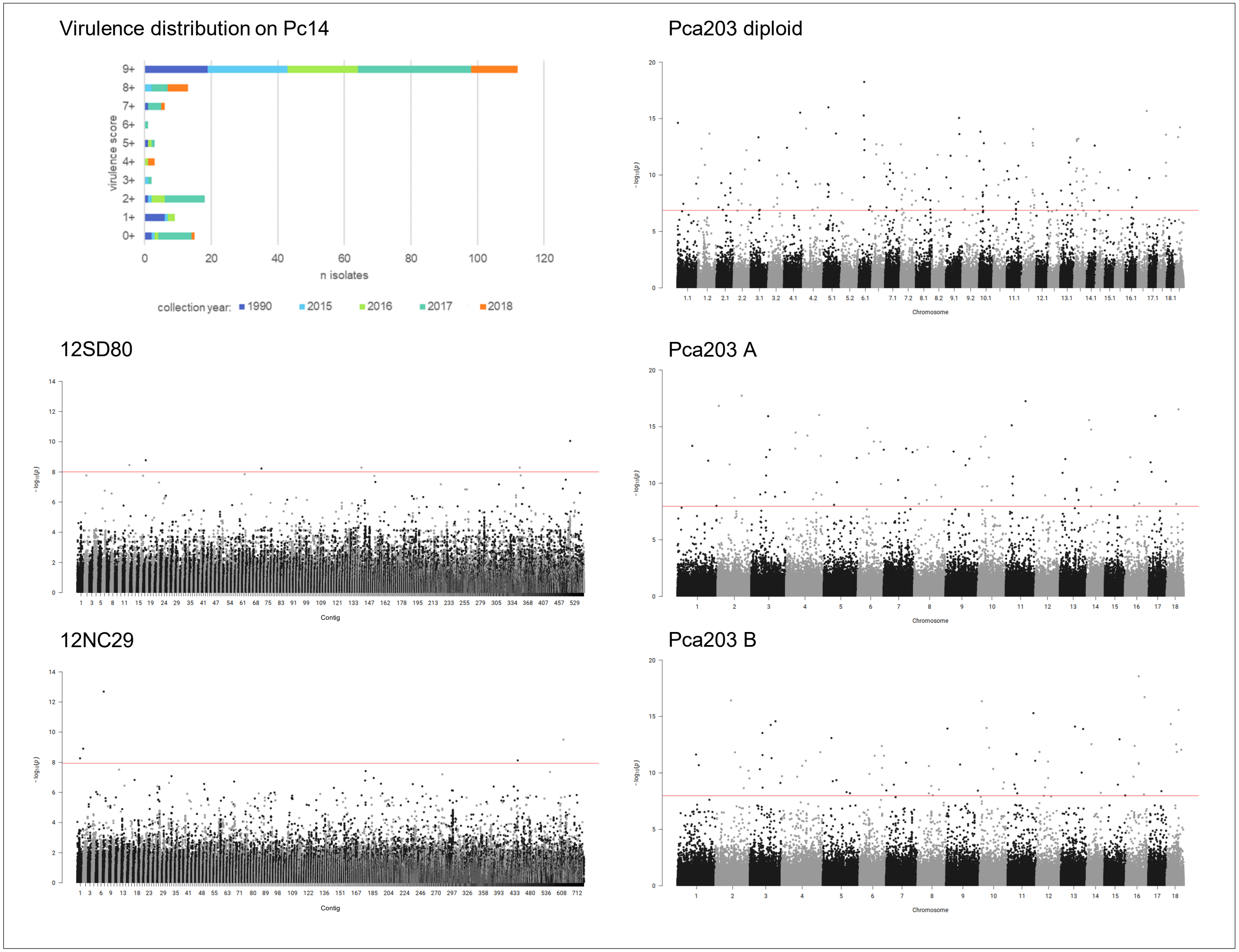

**Fig 8A. quantile-quantile plots of SNP association values for virulence of *Pca* to oat line *Pc35*.** Red lines indicate expected P-value distributions if P-values follow a uniform distribution (null hypothesis).

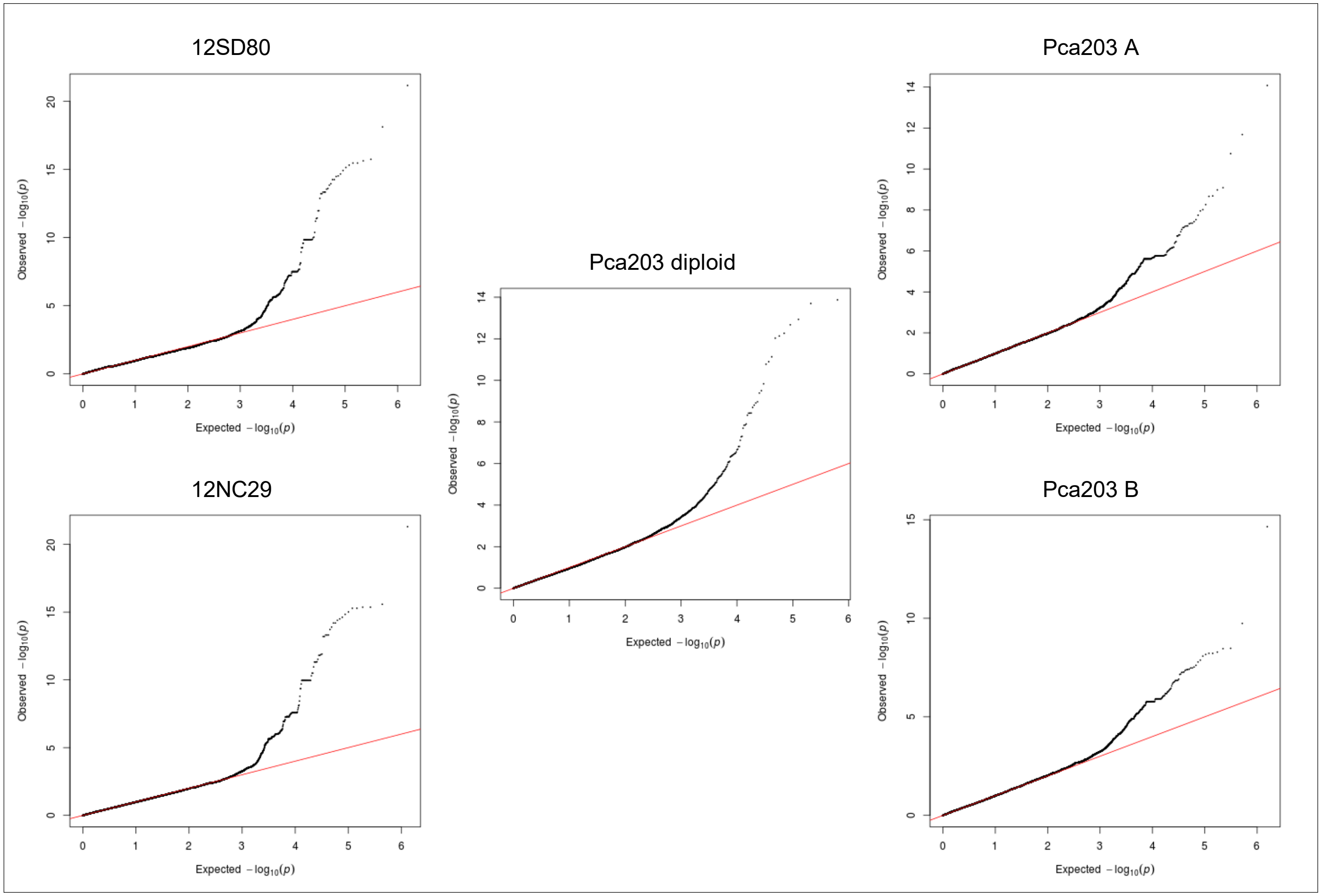

**Fig 8B. *Pca* isolate virulence score distribution and Manhattan plots showing SNP association values for virulence to oat line *Pc35*.** Red and blue horizontal lines denote Bonferroni significance threshold (α = 0.01/total number of markers) and 5% false discovery rate threshold, respectively. Significant association peaks are labelled with contig or chromosome number and assigned VGI number.

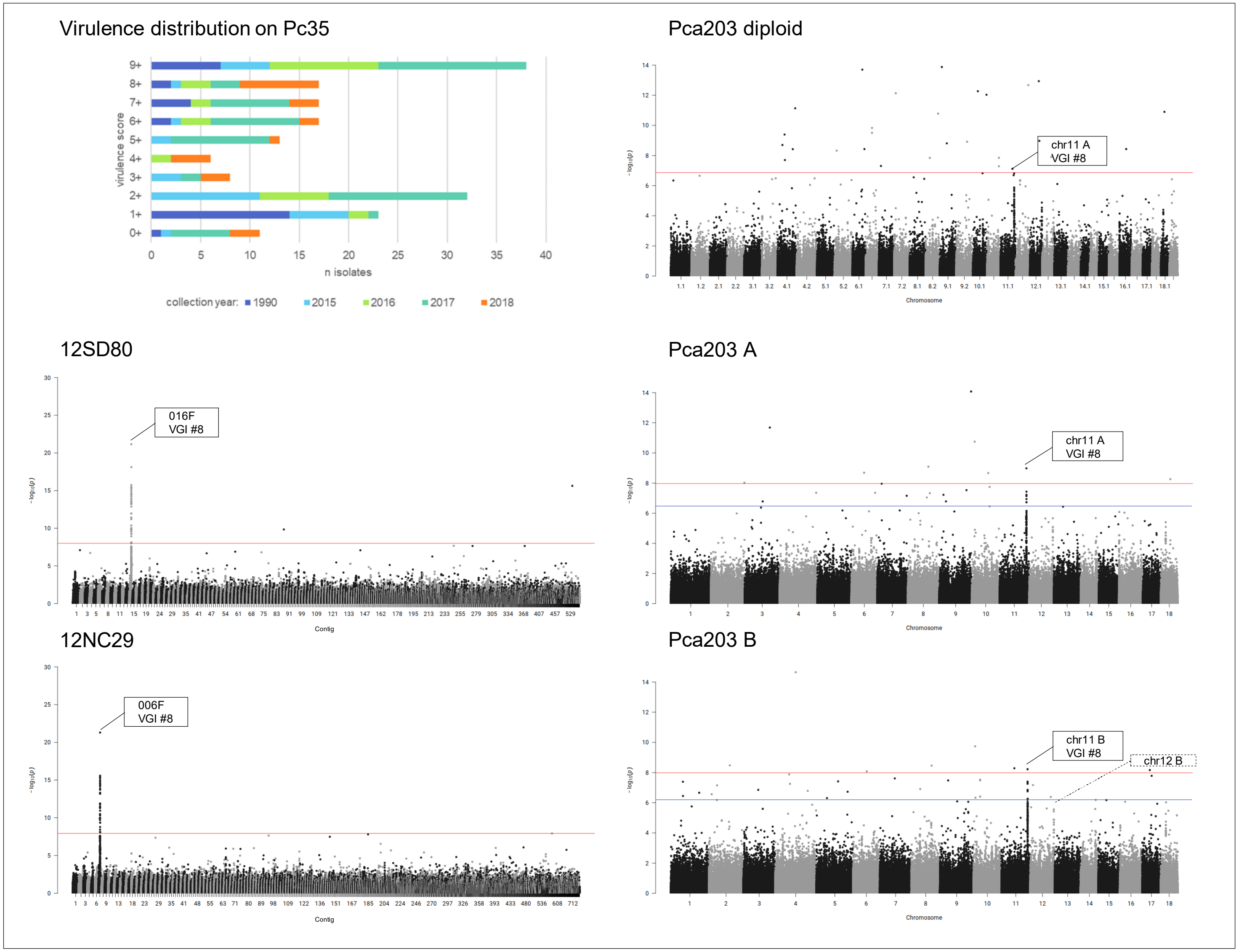

**Fig 9A. quantile-quantile plots of SNP association values for virulence of *Pca* to oat line *Pc36*.** Red lines indicate expected P-value distributions if P-values follow a uniform distribution (null hypothesis).

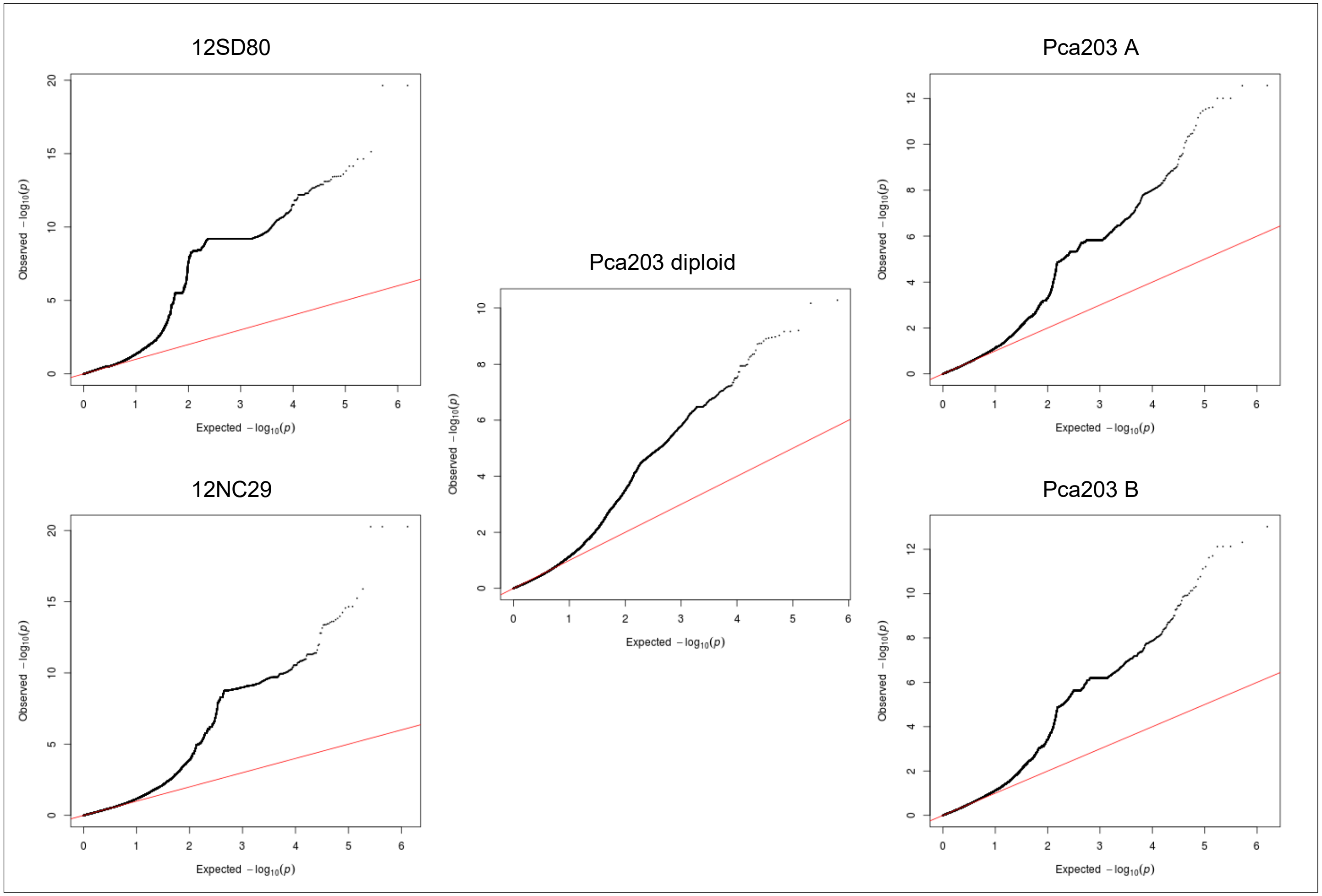

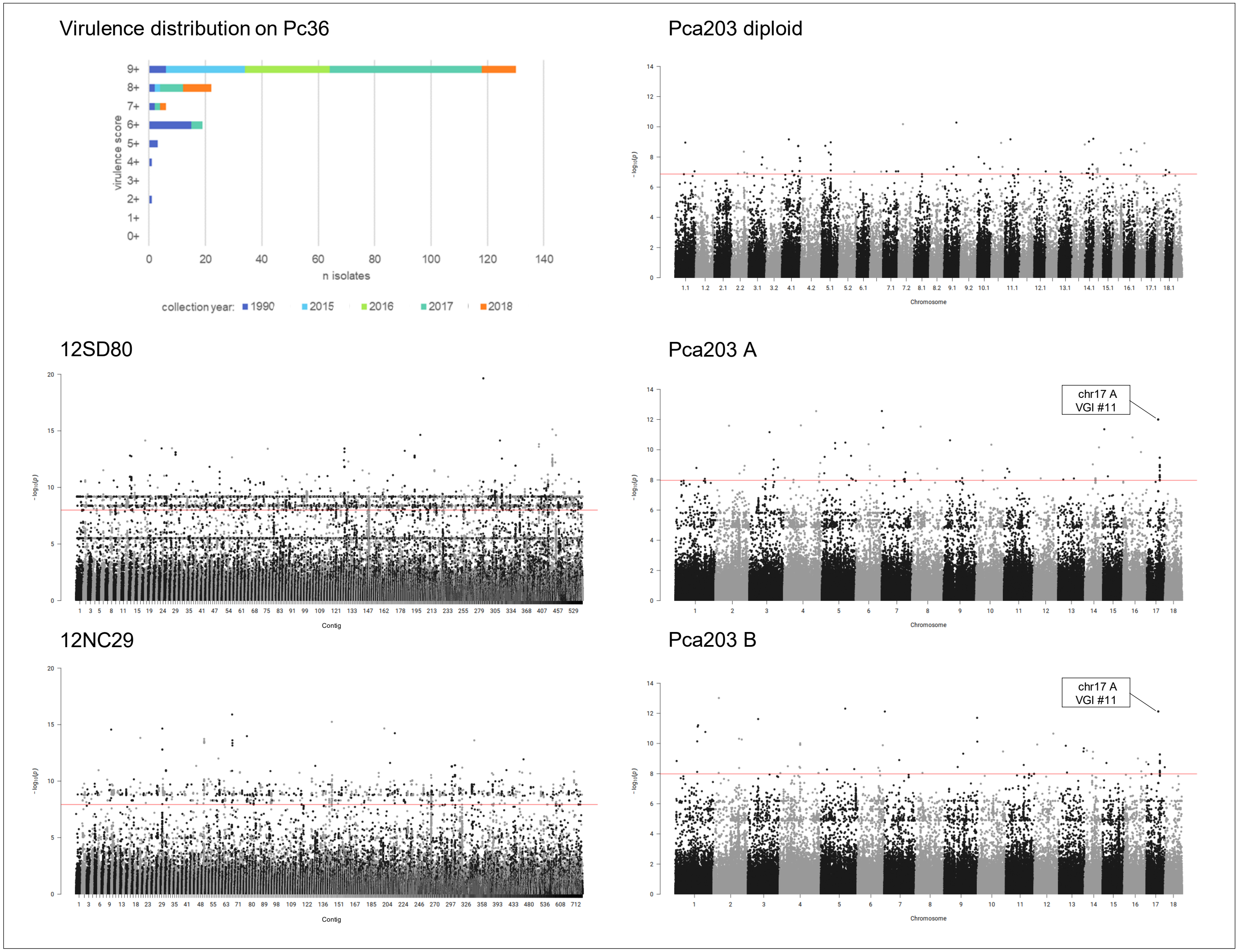


**Fig 9B. *Pca* isolate virulence score distribution and Manhattan plots showing SNP association values for virulence to oat line *Pc36*.** Red horizontal lines denote Bonferroni significance threshold (α = 0.01/total number of markers). Significant association peaks are labelled with contig or chromosome number and assigned VGI number.

**Fig 10A. quantile-quantile plots of SNP association values for virulence of *Pca* to oat line *Pc38*.** Red lines indicate expected P-value distributions if P-values follow a uniform distribution (null hypothesis).

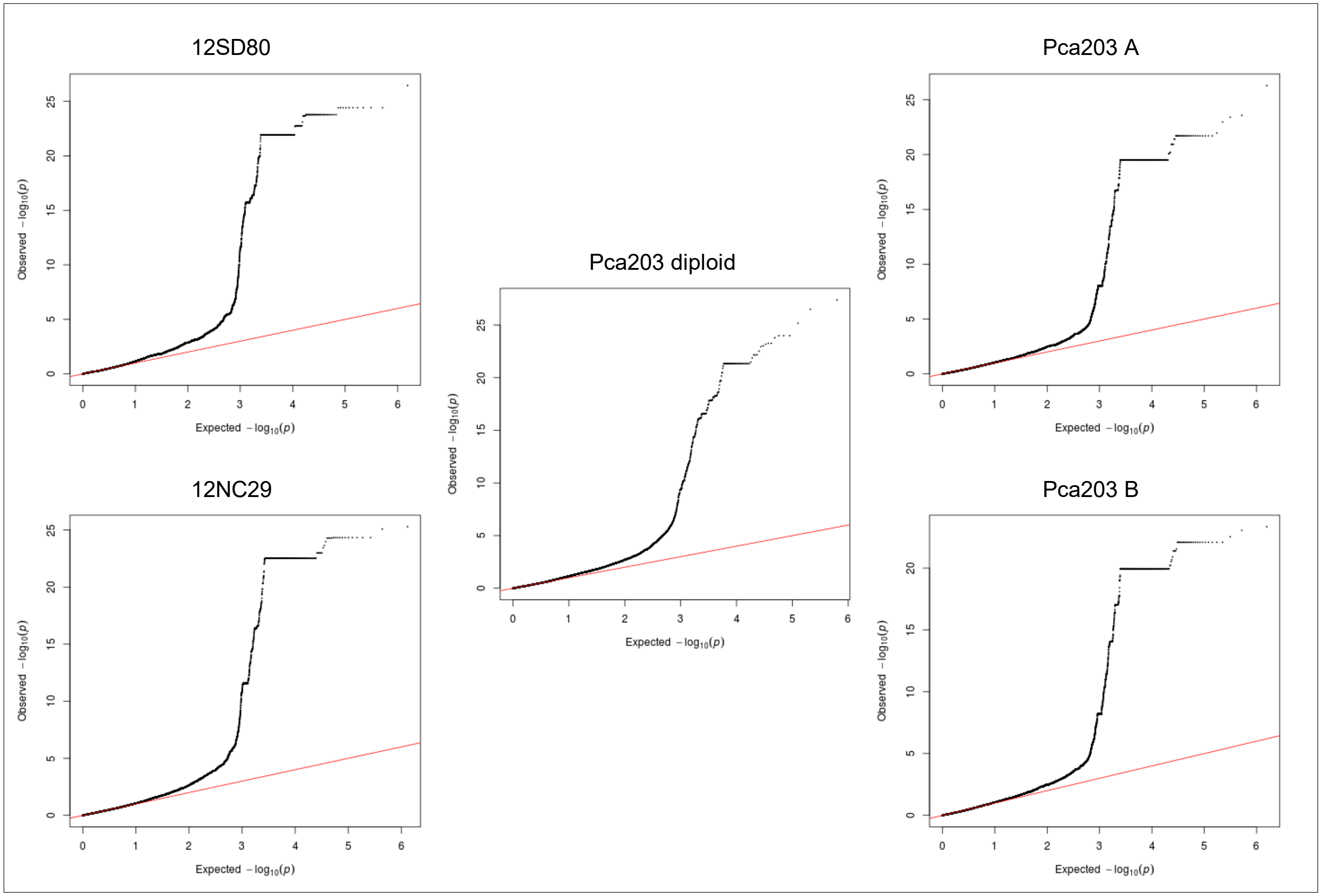

**Fig 10B. *Pca* isolate virulence score distribution and Manhattan plots showing SNP association values for virulence to oat line *Pc38*.** Red horizontal lines denote Bonferroni significance threshold (α = 0.01/total number of markers). Significant association peaks are labelled with contig or chromosome number and assigned VGI number.

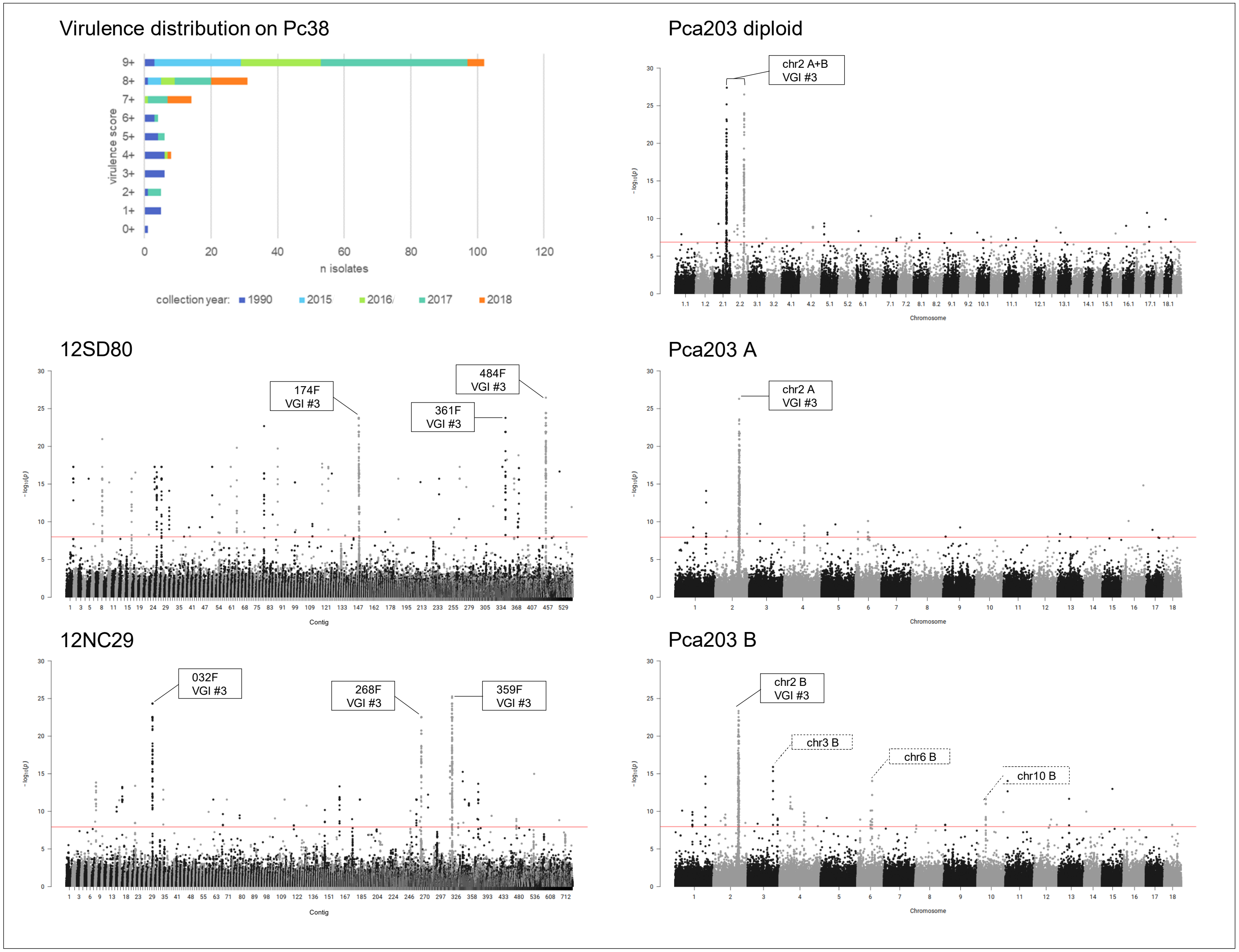

**Fig 11A. quantile-quantile plots of SNP association values for virulence of *Pca* to oat line *Pc39*.** Red lines indicate expected P-value distributions if P-values follow a uniform distribution (null hypothesis).

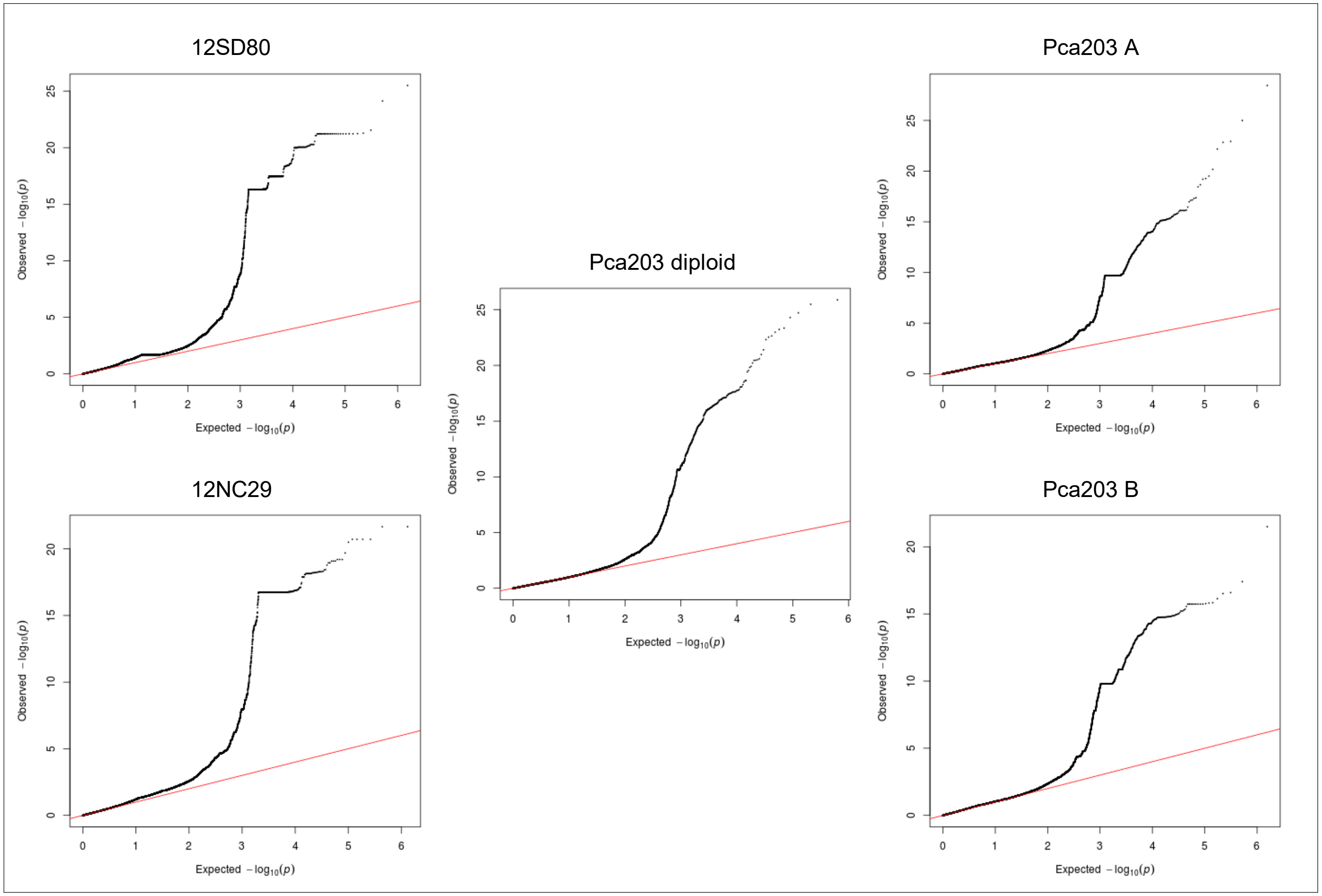

**Fig 11B. *Pca* isolate virulence score distribution and Manhattan plots showing SNP association values for virulence to oat line *Pc39*.** Red horizontal lines denote Bonferroni significance threshold (α = 0.01/total number of markers). Significant association peaks are labelled with contig or chromosome number and assigned VGI number.

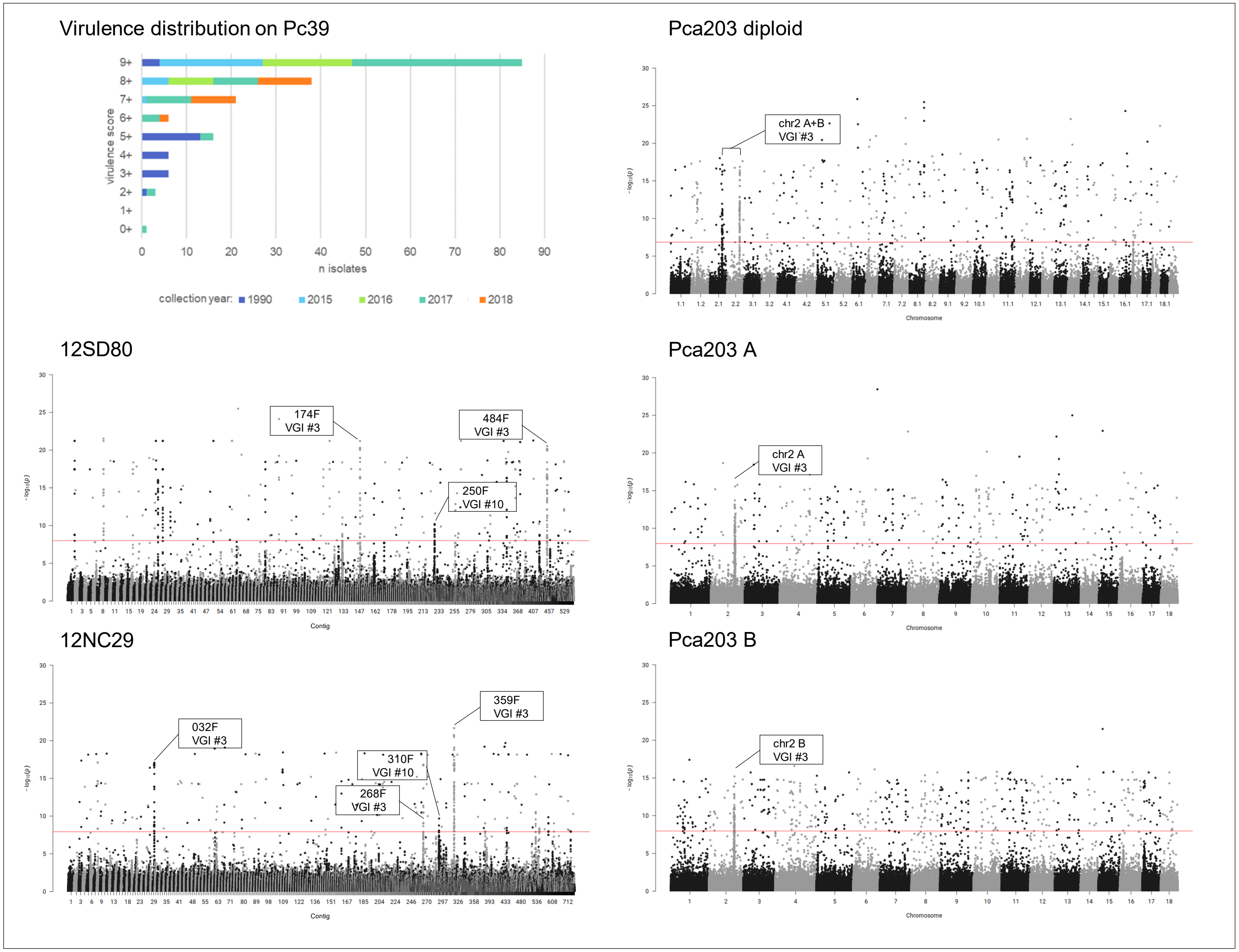

**Fig 12A. quantile-quantile plots of SNP association values for virulence of *Pca* to oat line *Pc40*.** Red lines indicate expected P-value distributions if P-values follow a uniform distribution (null hypothesis).

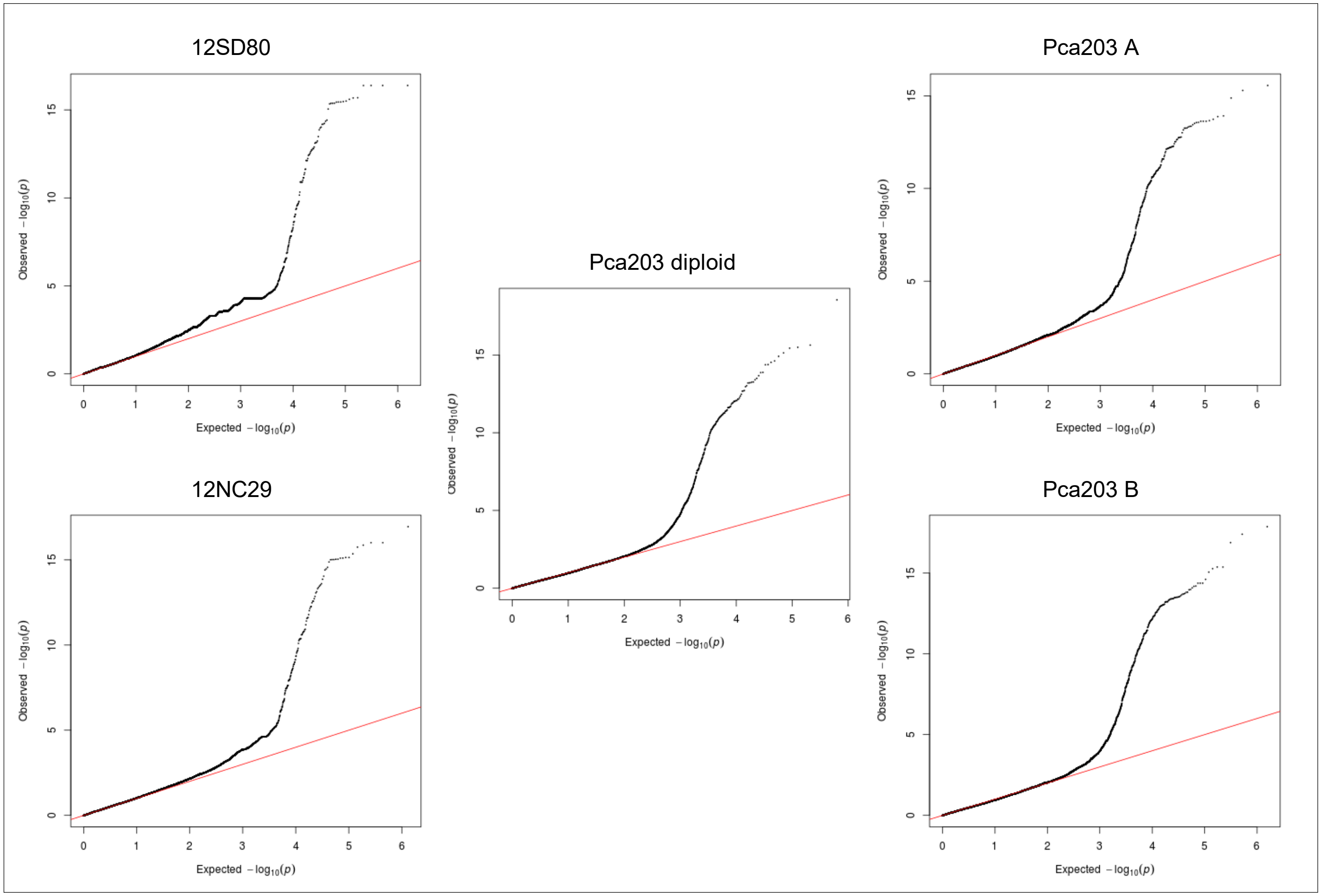

**Fig 12B. *Pca* isolate virulence score distribution and Manhattan plots showing SNP association values for virulence to oat line *Pc40*.** Red horizontal lines denote Bonferroni significance threshold (α = 0.01/total number of markers).

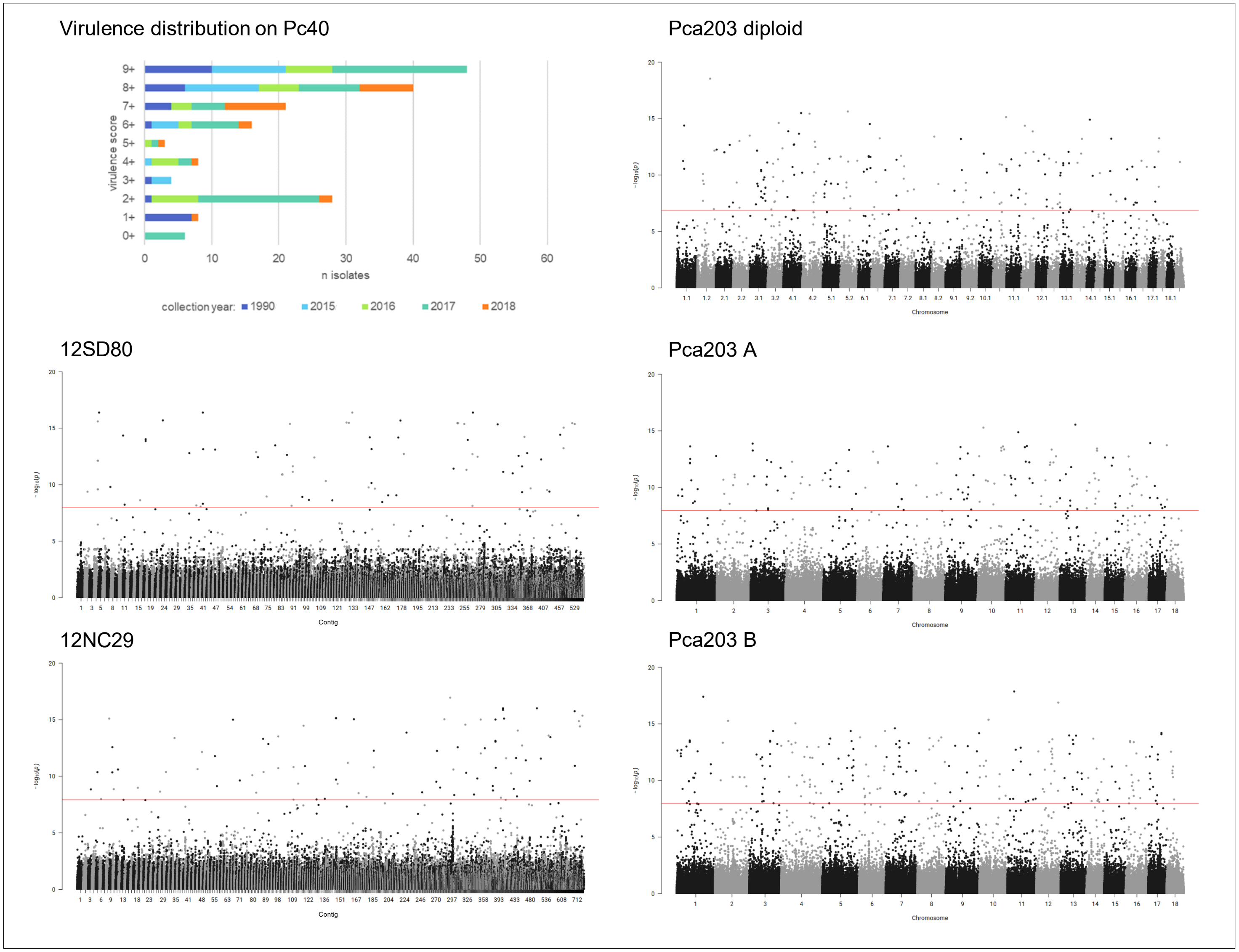

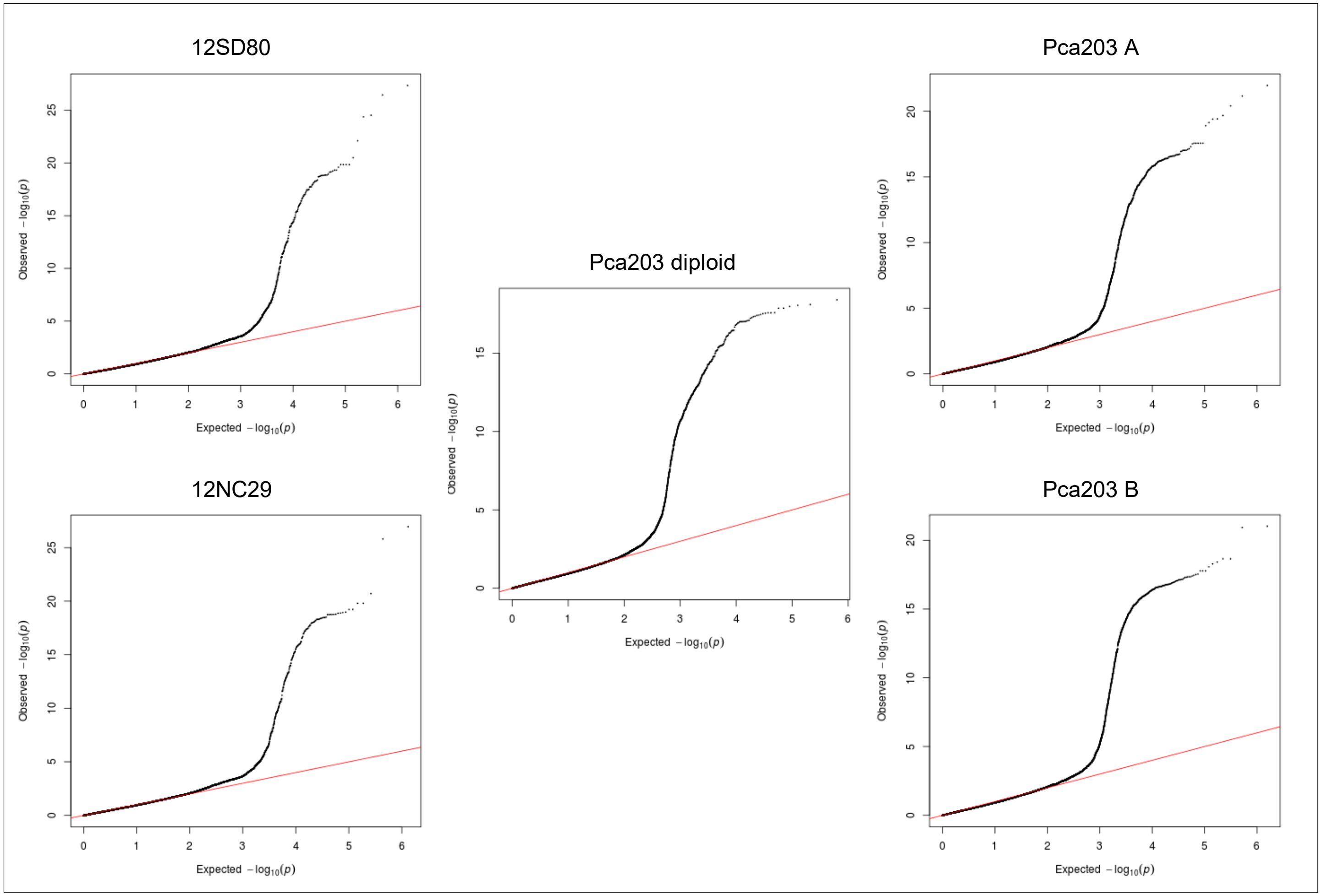


**Fig 13A. quantile-quantile plots of SNP association values for virulence of *Pca* to oat line *Pc45*.** Red lines indicate expected P-value distributions if P-values follow a uniform distribution (null hypothesis).

**Fig 13B.** ***Pca*** **isolate virulence score distribution and Manhattan plots showing SNP association values for virulence to oat line** ***Pc45*****.** Red horizontal lines denote Bonferroni significance threshold (α = 0.01/total number of markers).

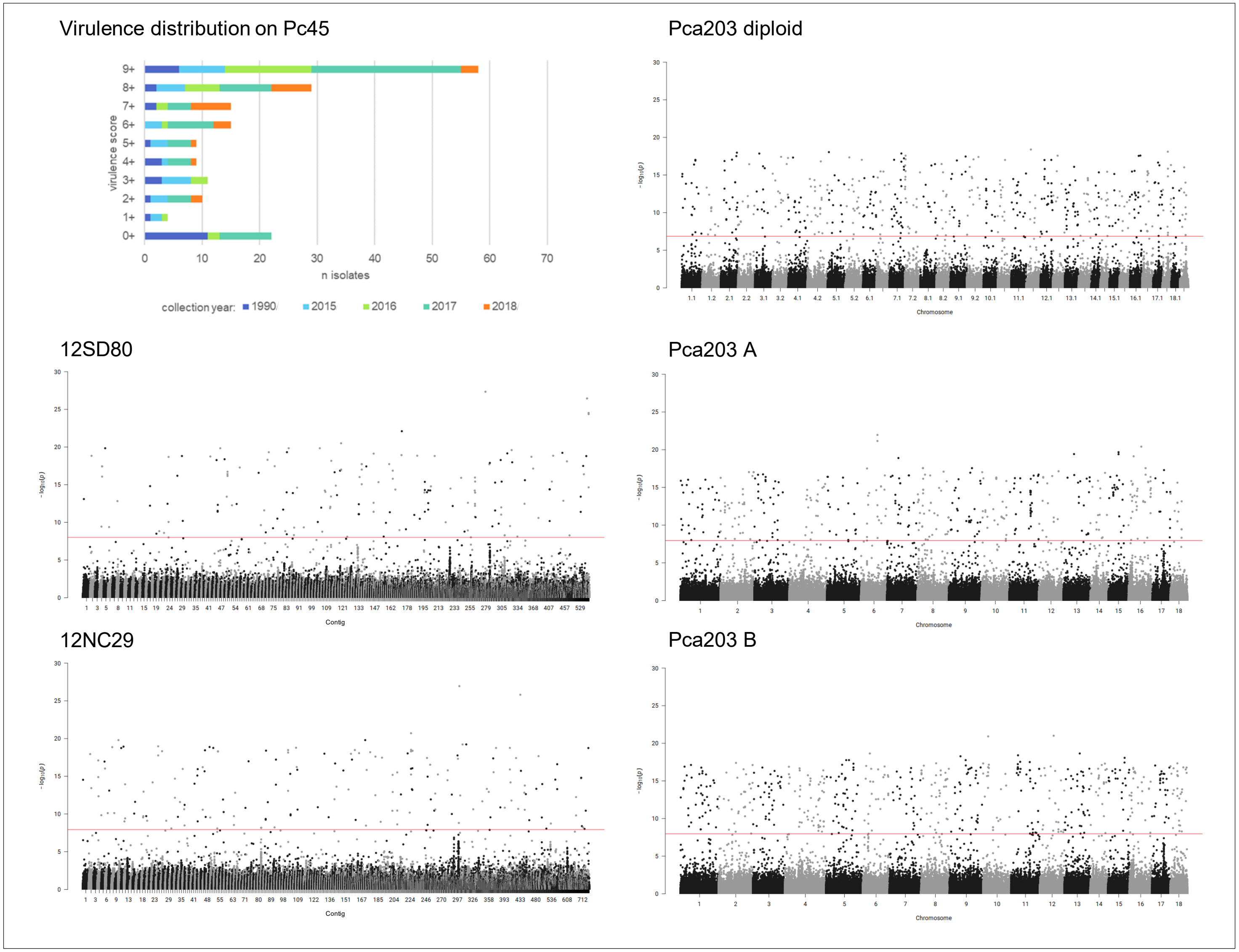

**Fig 14A. quantile-quantile plots of SNP association values for virulence of *Pca* to oat line *Pc46*.** Red lines indicate expected P-value distributions if P-values follow a uniform distribution (null hypothesis).

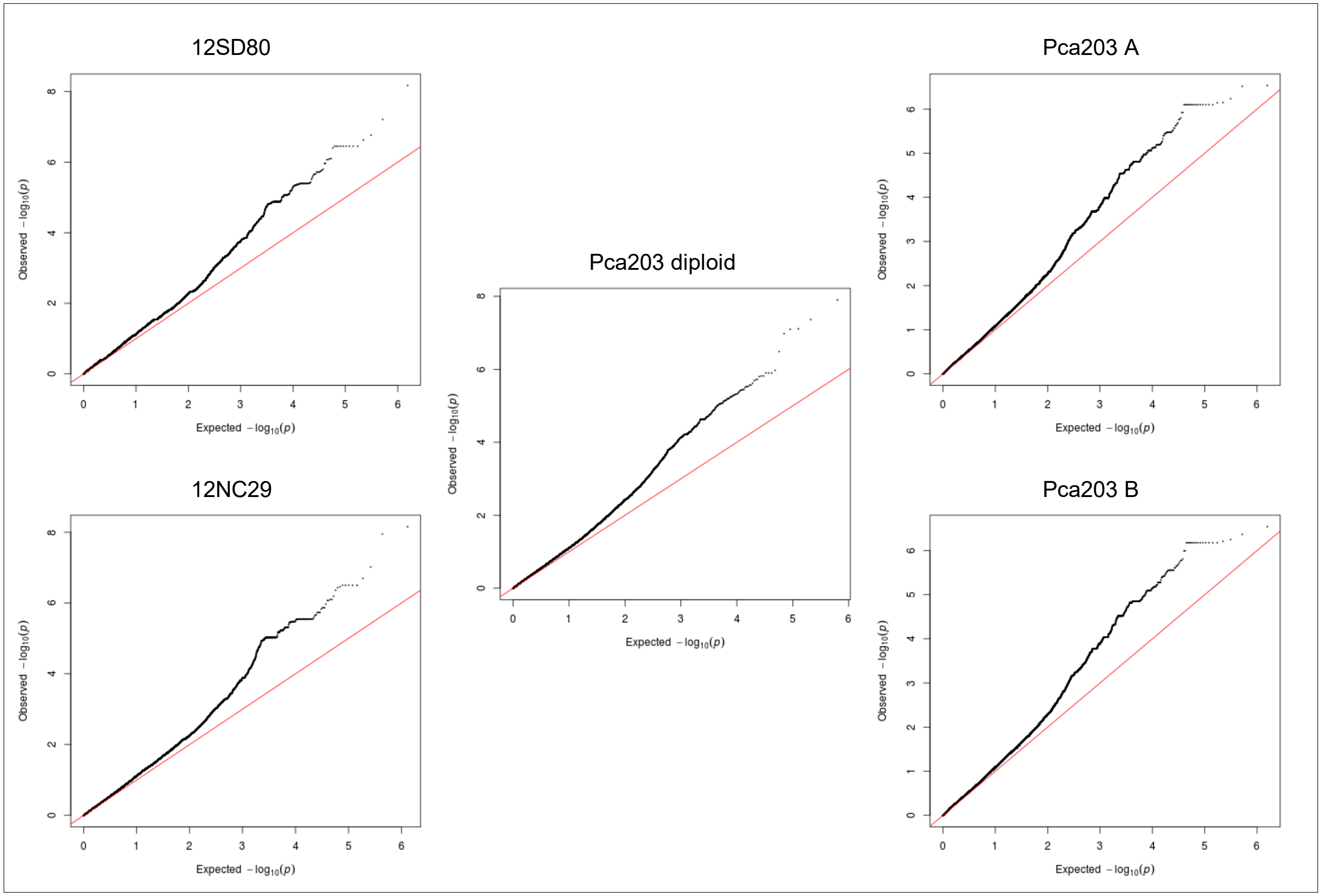

**Fig 14B. *Pca* isolate virulence score distribution and Manhattan plots showing SNP association values for virulence to oat line *Pc46*.** Red horizontal lines denote Bonferroni significance threshold (α = 0.01/total number of markers).

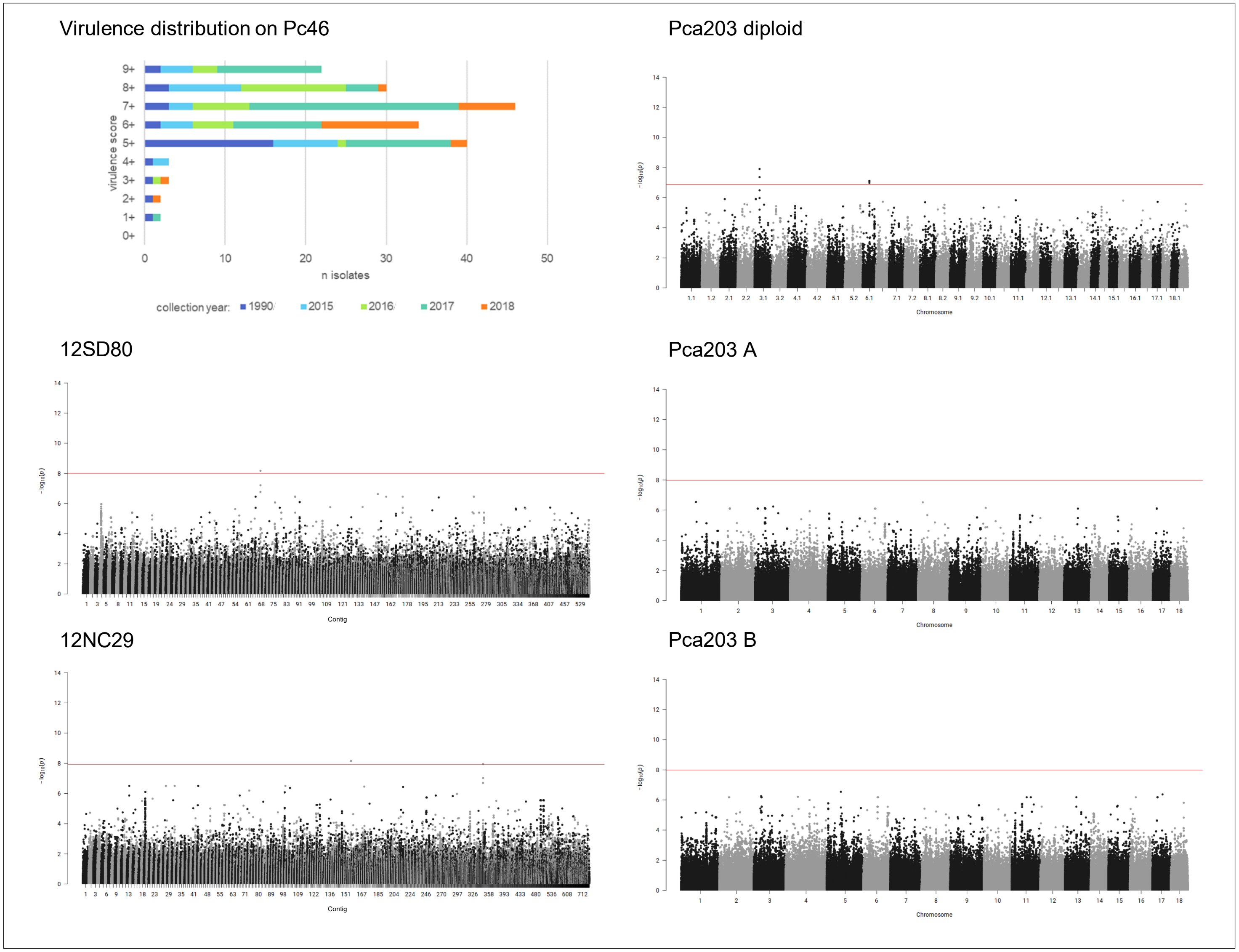

**Fig 15A. quantile-quantile plots of SNP association values for virulence of *Pca* to oat line *Pc48*.** Red lines indicate expected P-value distributions if P-values follow a uniform distribution (null hypothesis).

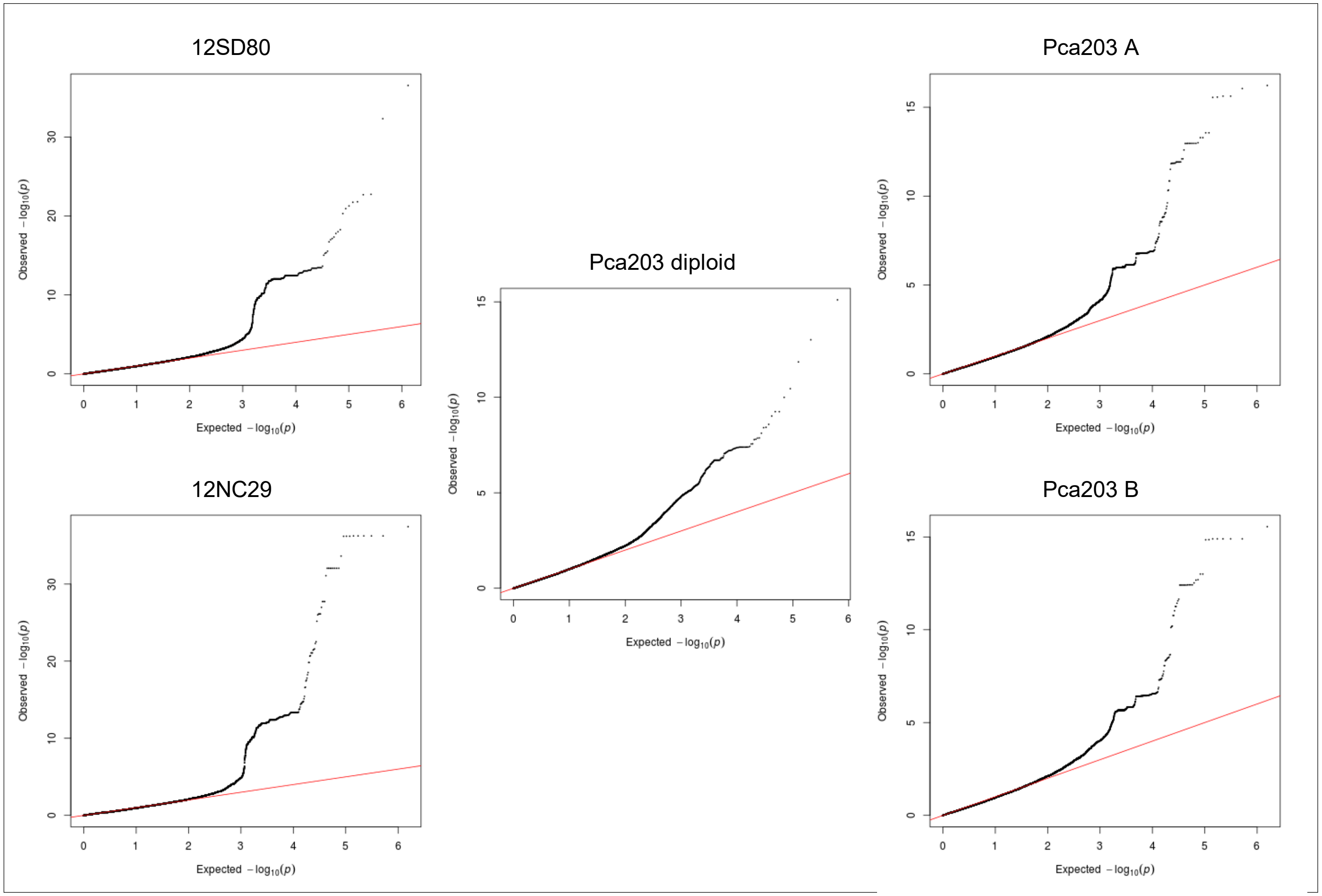

**Fig 15B. *Pca* isolate virulence score distribution and Manhattan plots showing SNP association values for virulence to oat line *Pc48*.** Red horizontal lines denote Bonferroni significance threshold (α = 0.01/total number of markers). Significant association peaks are labelled with contig or chromosome number and assigned VGI number.

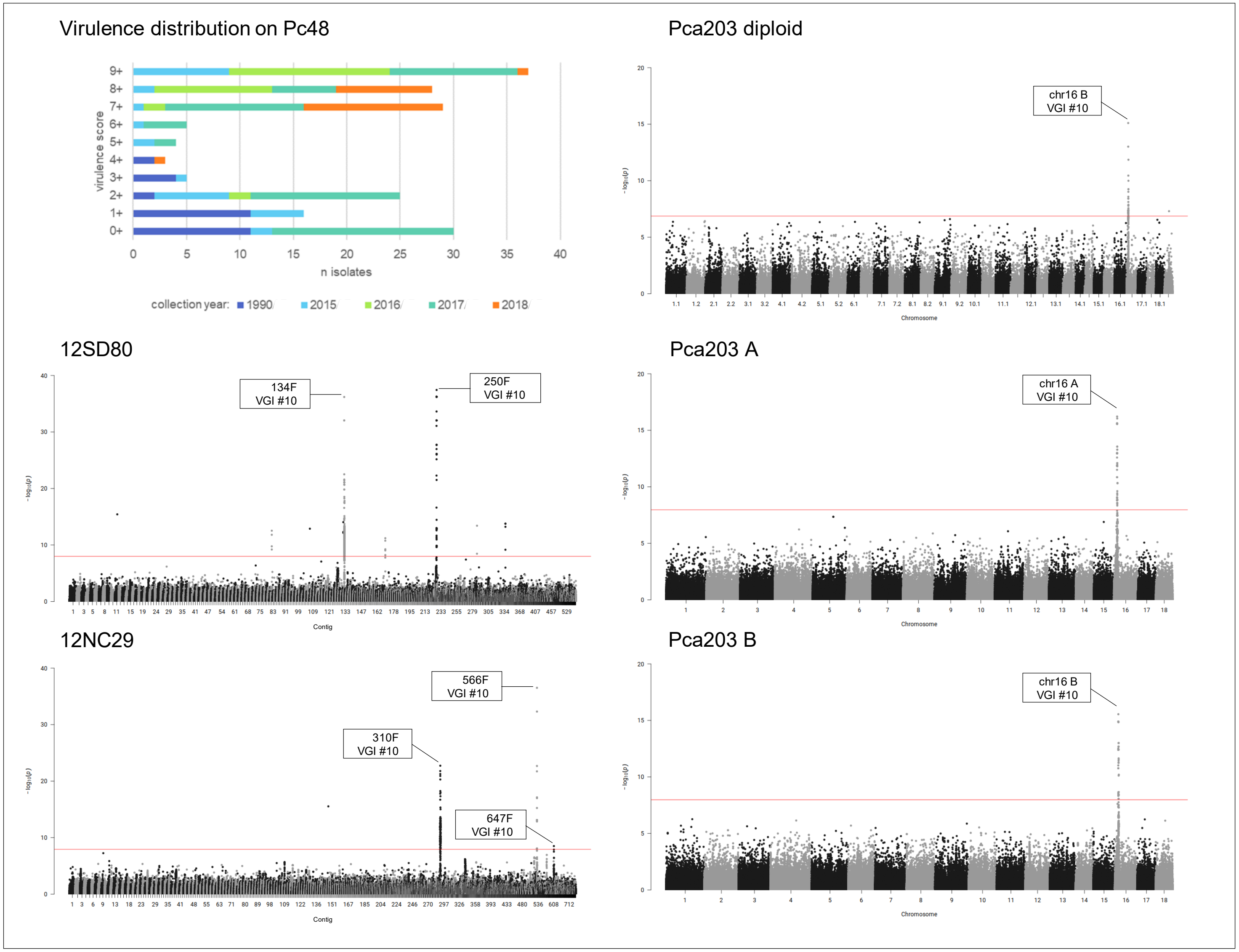

**Fig 16A. quantile-quantile plots of SNP association values for virulence of *Pca* to oat line *Pc50*.** Red lines indicate expected P-value distributions if P-values follow a uniform distribution (null hypothesis).

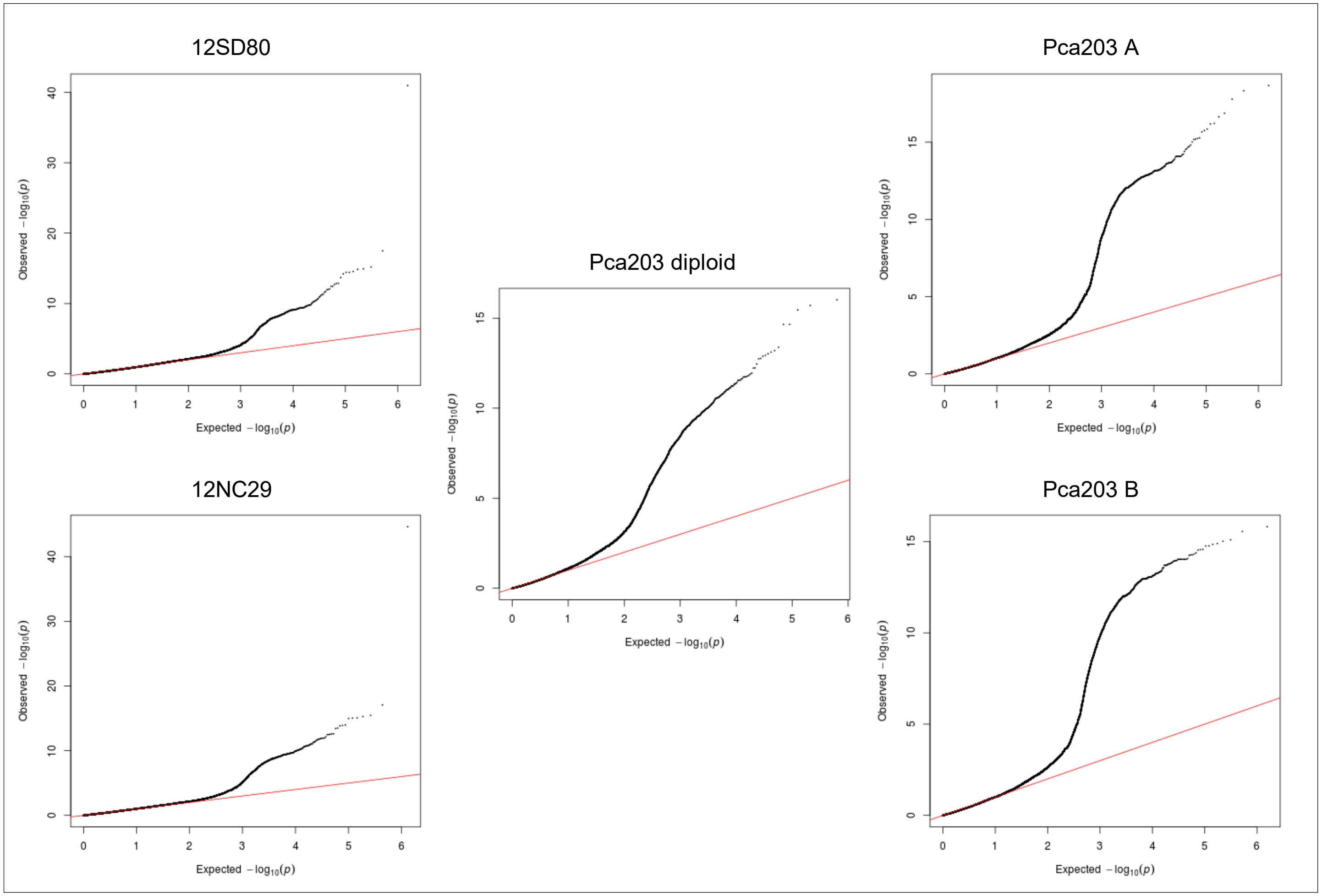

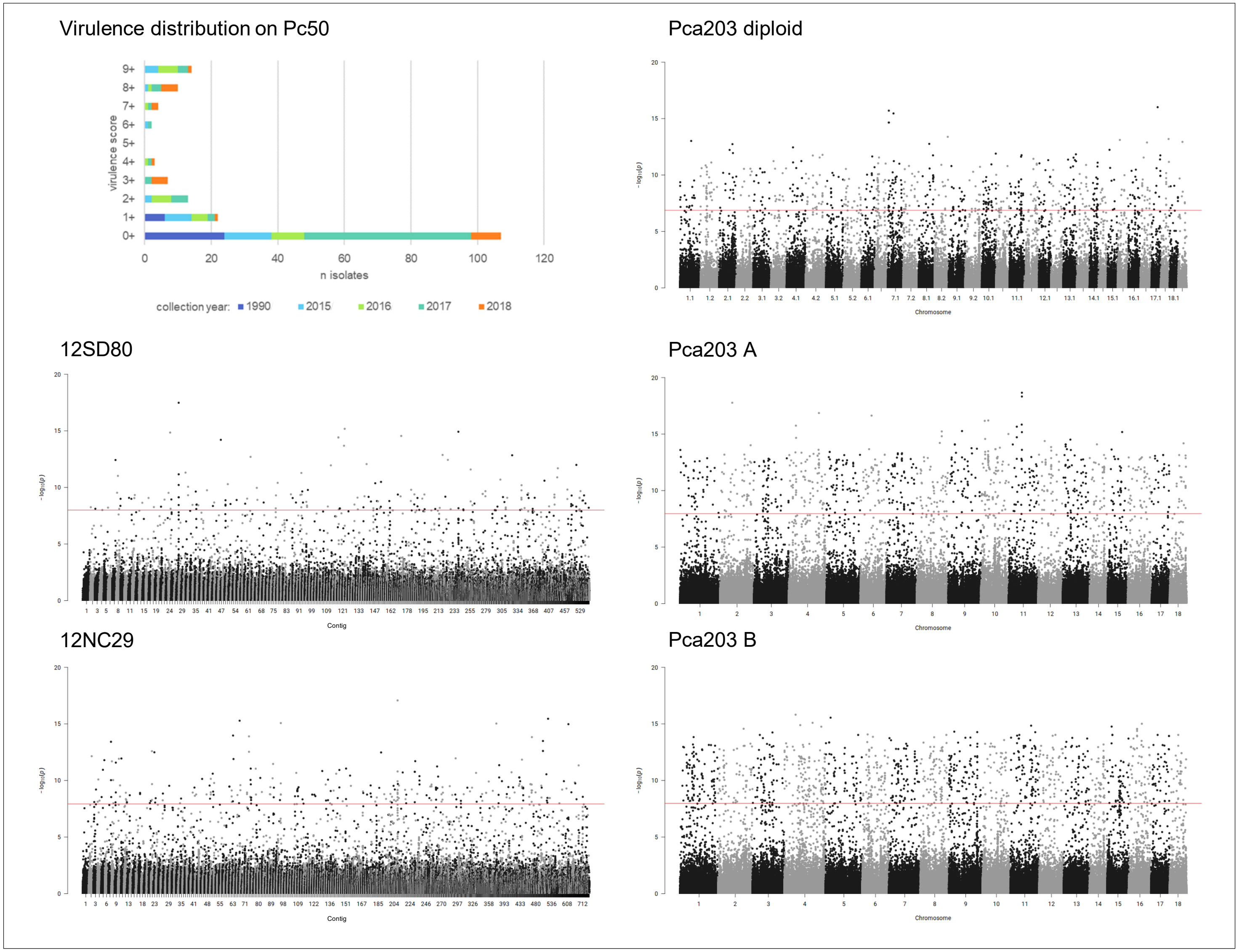


**Fig 16B. *Pca* isolate virulence score distribution and Manhattan plots showing SNP association values for virulence to oat line *Pc50*.** Red horizontal lines denote Bonferroni significance threshold (α = 0.01/total number of markers).

**Fig 17A. quantile-quantile plots of SNP association values for virulence of *Pca* to oat line *Pc51*.** Red lines indicate expected P-value distributions if P-values follow a uniform distribution (null hypothesis).

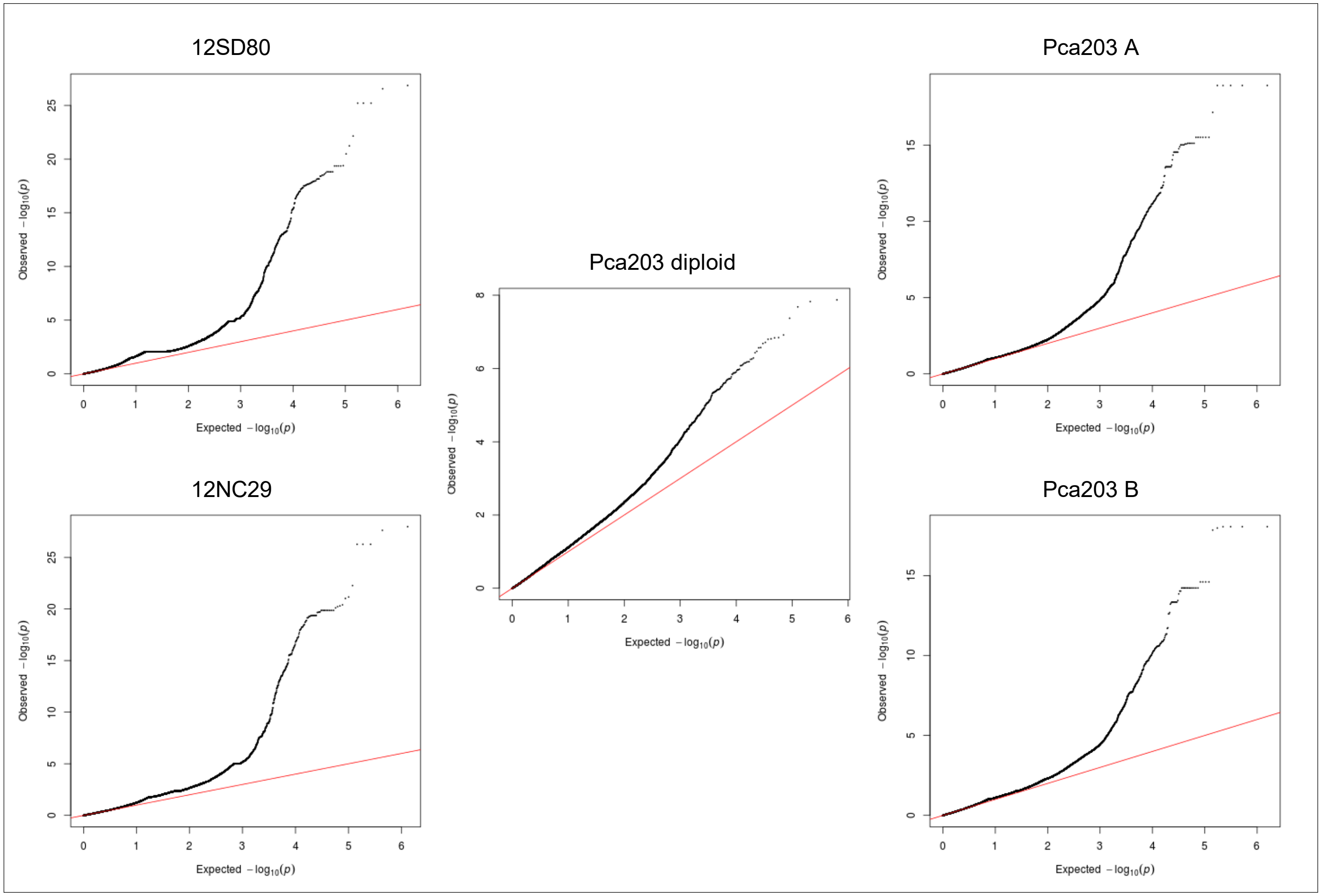

**Fig 17B. *Pca* isolate virulence score distribution and Manhattan plots showing SNP association values for virulence to oat line *Pc51*.** Red horizontal lines denote Bonferroni significance threshold (α = 0.01/total number of markers). Significant association peaks are labelled with contig or chromosome number and assigned VGI number.

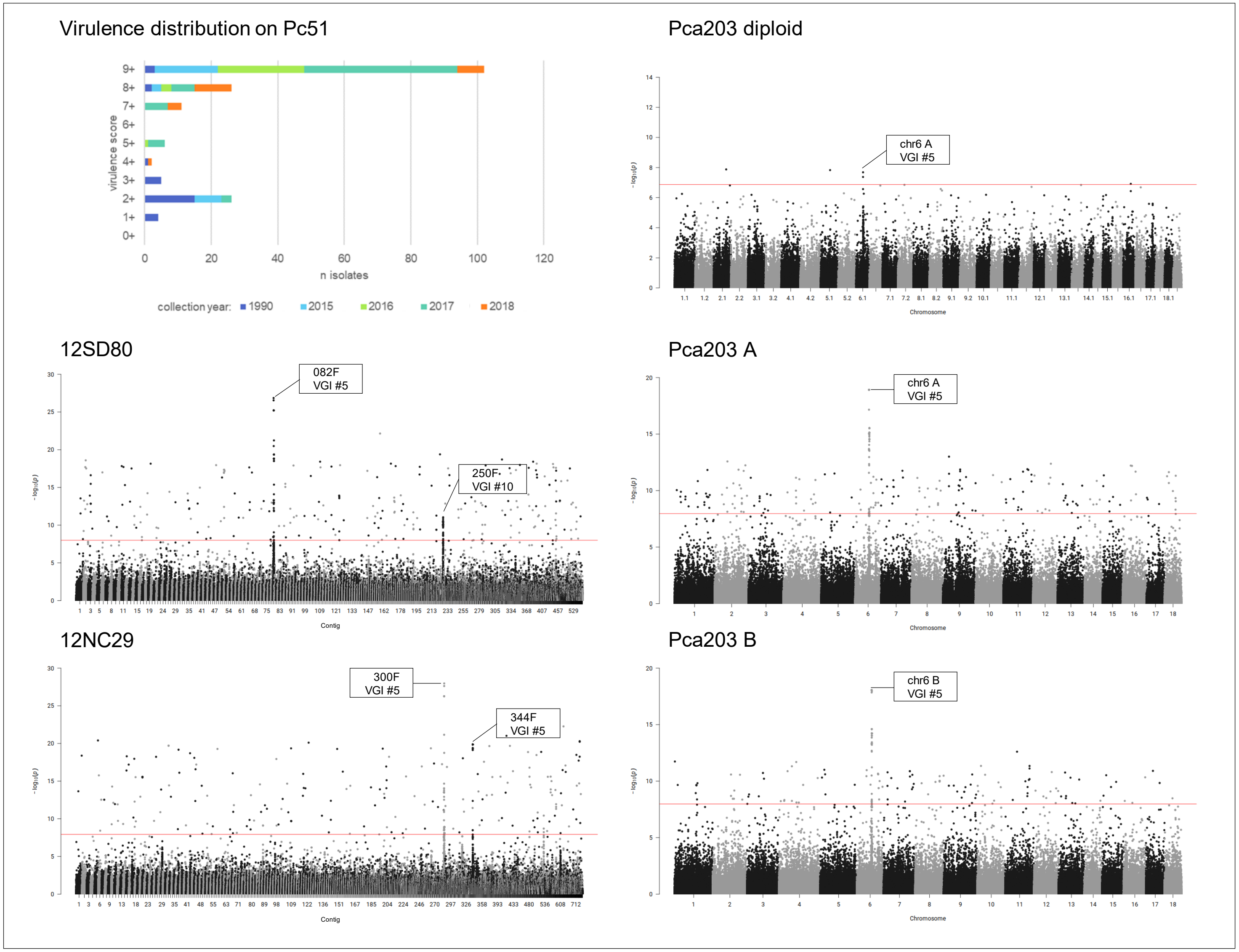

**Fig 18A. quantile-quantile plots of SNP association values for virulence of *Pca* to oat line *Pc52*.** Red lines indicate expected P-value distributions if P-values follow a uniform distribution (null hypothesis).

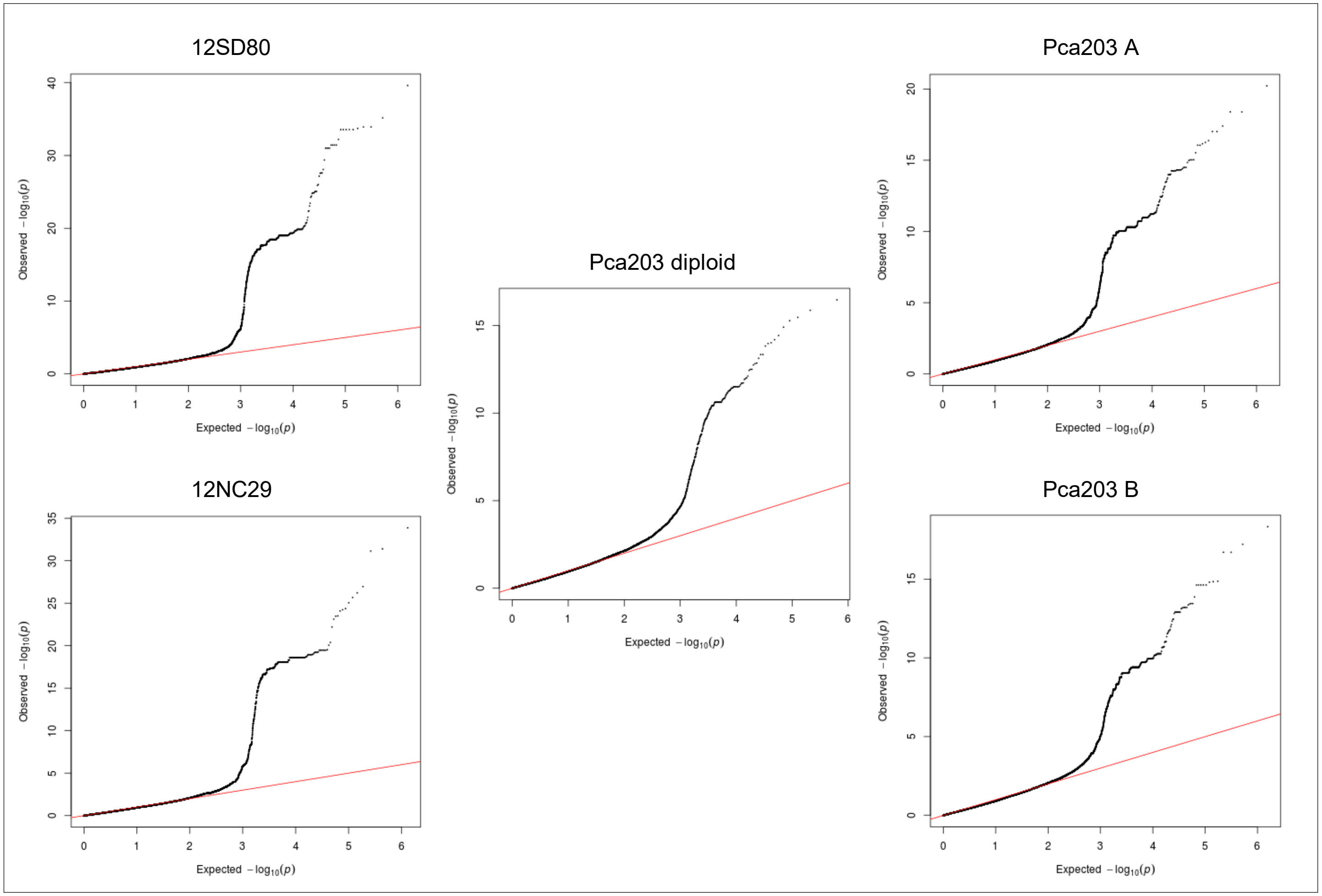

**Fig 18B. *Pca* isolate virulence score distribution and Manhattan plots showing SNP association values for virulence to oat line *Pc52*.** Red horizontal lines denote Bonferroni significance threshold (α = 0.01/total number of markers). Significant association peaks are labelled with contig or chromosome number and assigned VGI number.

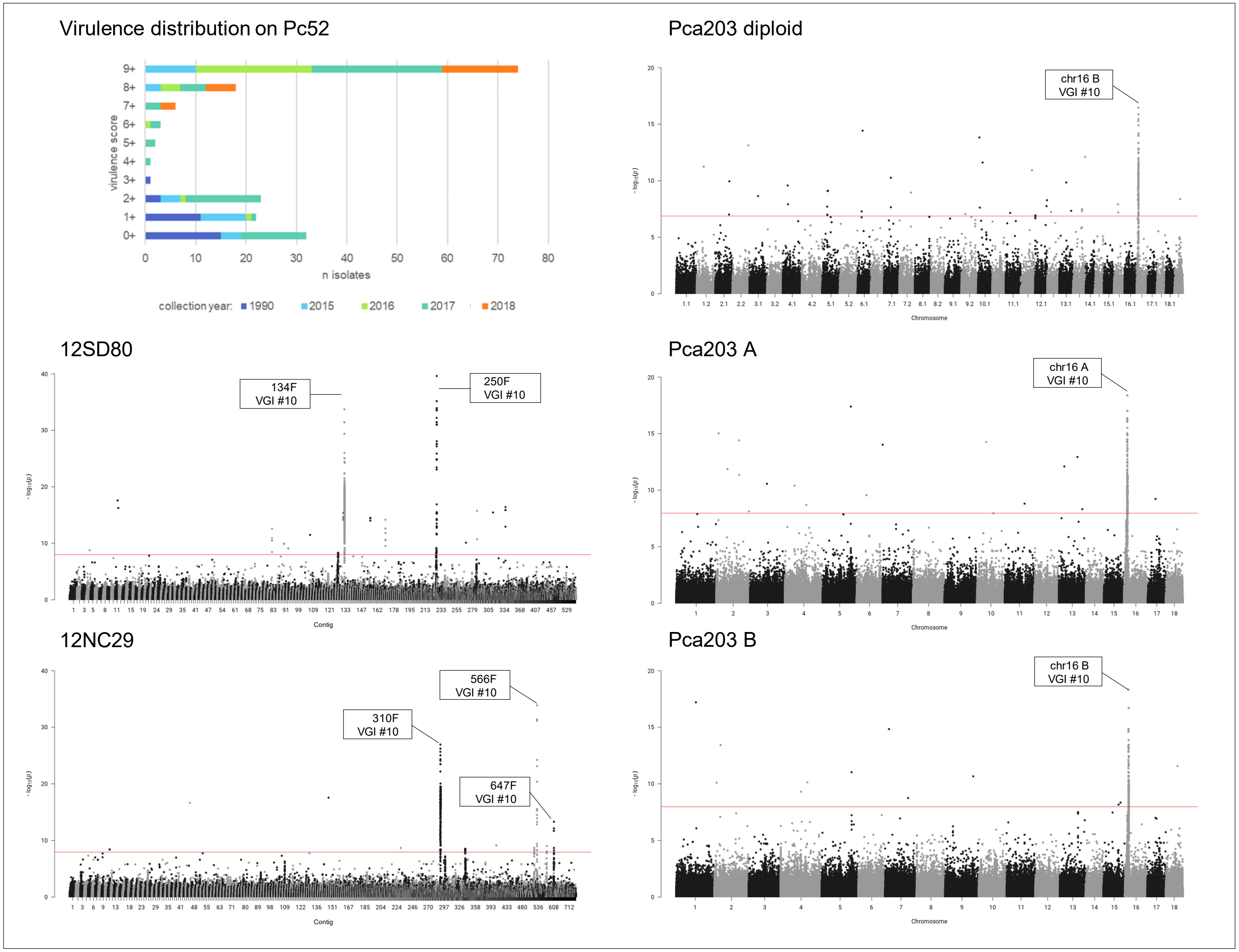

**Fig 19A. quantile-quantile plots of SNP association values for virulence of *Pca* to oat line *Pc53*.** Red lines indicate expected P-value distributions if P-values follow a uniform distribution (null hypothesis).

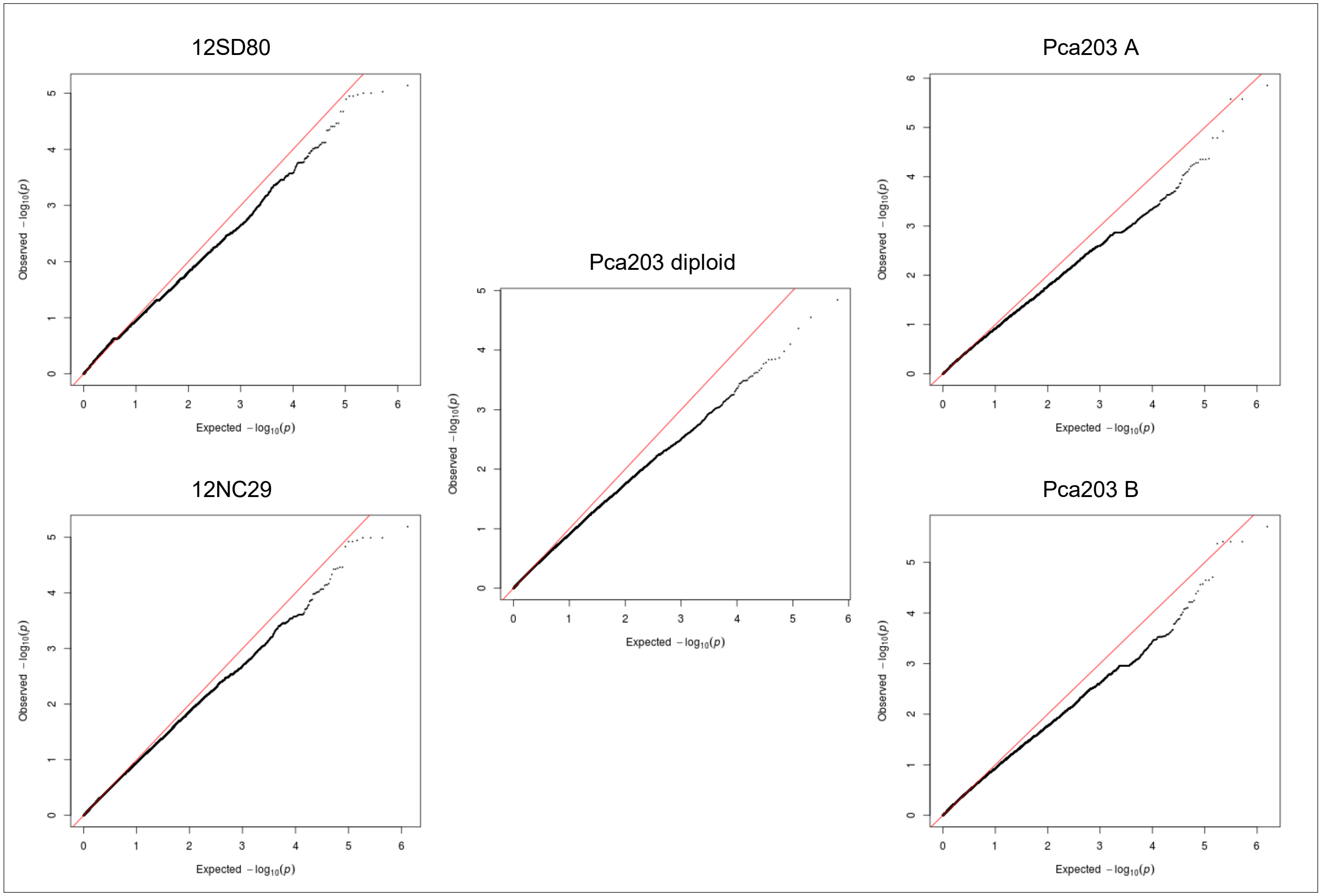

**Fig 19B. *Pca* isolate virulence score distribution and Manhattan plots showing SNP association values for virulence to oat line *Pc53*.** Red and blue horizontal lines denote Bonferroni significance threshold (α = 0.01/total number of markers) and 5% false discovery rate threshold, respectively. Significant association peaks are labelled with contig or chromosome number and assigned VGI number.

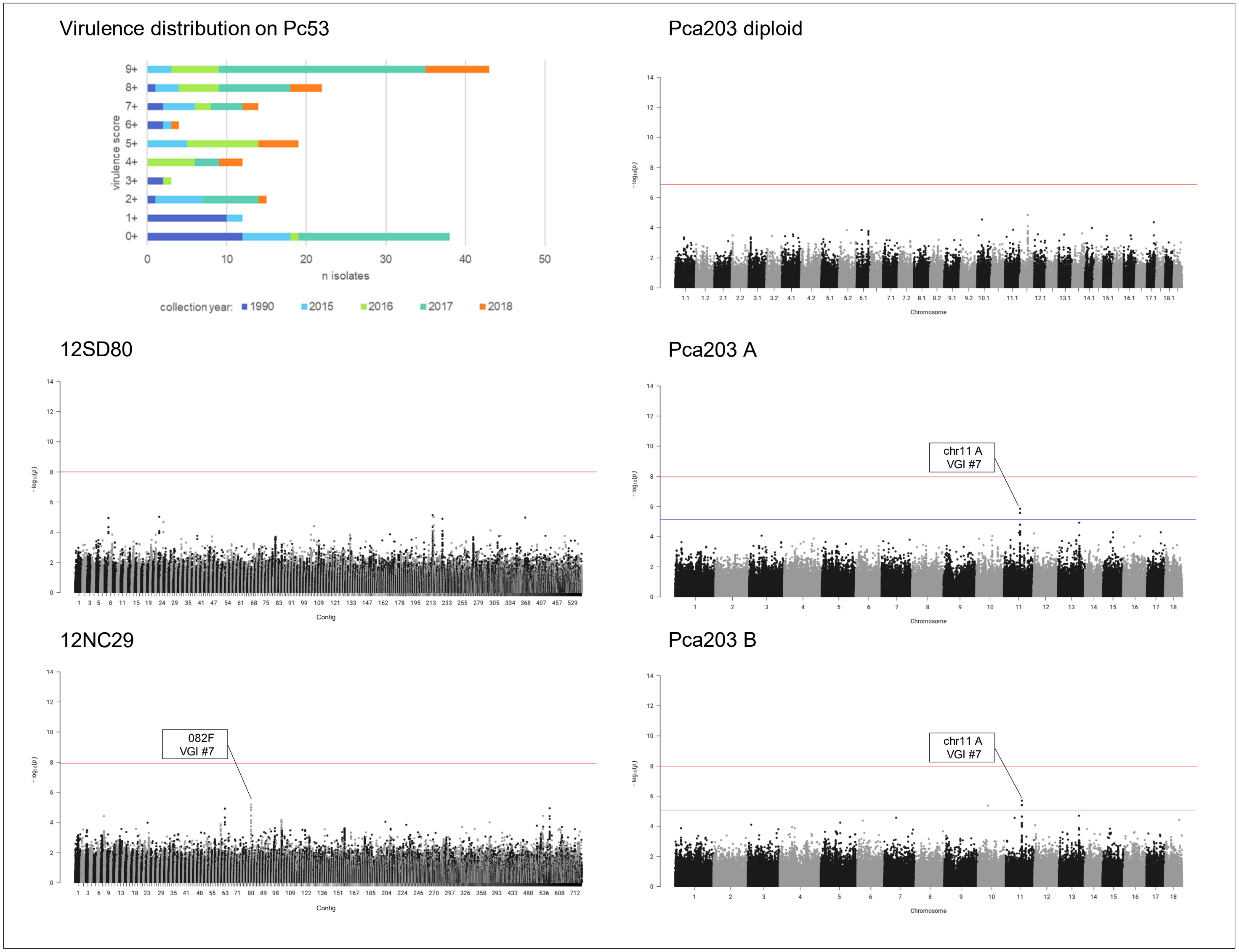

**Fig 20A. quantile-quantile plots of SNP association values for virulence of *Pca* to oat line *Pc54*.** Red lines indicate expected P-value distributions if P-values follow a uniform distribution (null hypothesis).

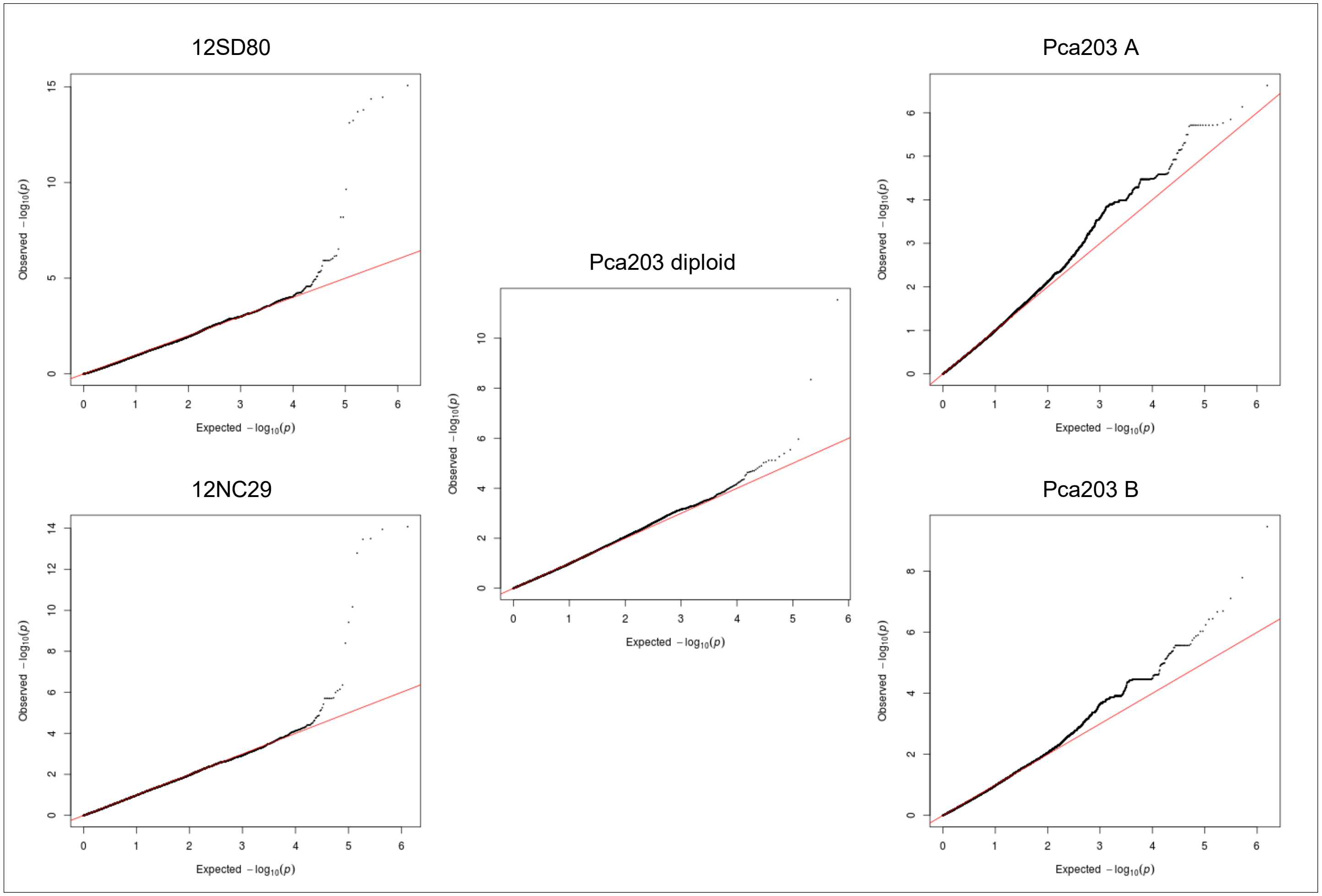

**Fig 20B. *Pca* isolate virulence score distribution and Manhattan plots showing SNP association values for virulence to oat line *Pc54*.** Red horizontal lines denote Bonferroni significance threshold (α = 0.01/total number of markers). Significant association peaks in 12SD80 and 12NC29 are labelled with contig number and assigned VGI number while labels in Pca203 A and B indicate corresponding homologous regions.

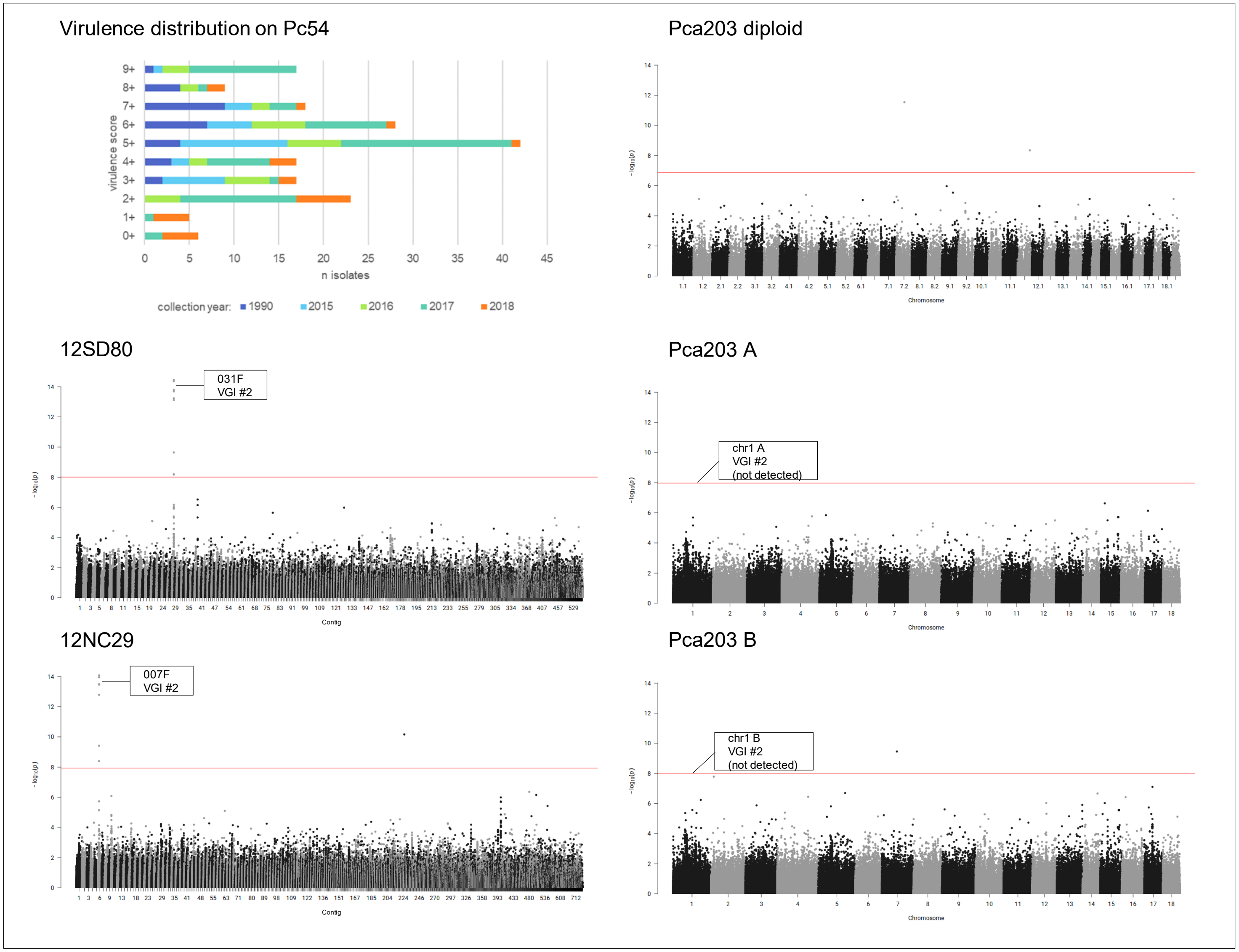

**Fig 21A. quantile-quantile plots of SNP association values for virulence of *Pca* to oat line *Pc55*.** Red lines indicate expected P-value distributions if P-values follow a uniform distribution (null hypothesis).

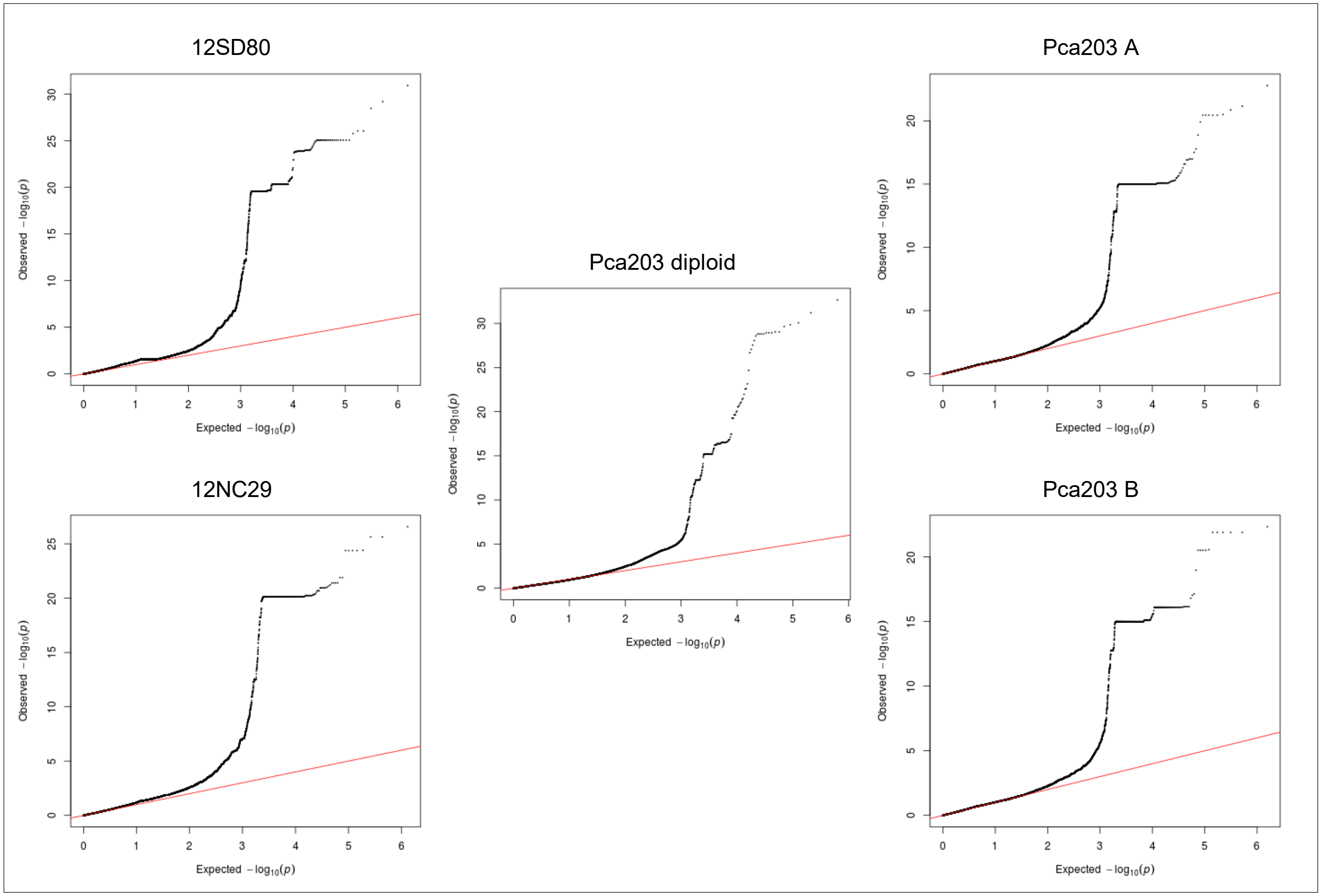

**Fig 21B. *Pca* isolate virulence score distribution and Manhattan plots showing SNP association values for virulence to oat line *Pc55*.** Red horizontal lines denote Bonferroni significance threshold (α = 0.01/total number of markers). Significant association peaks are labelled with contig or chromosome number and assigned VGI number.

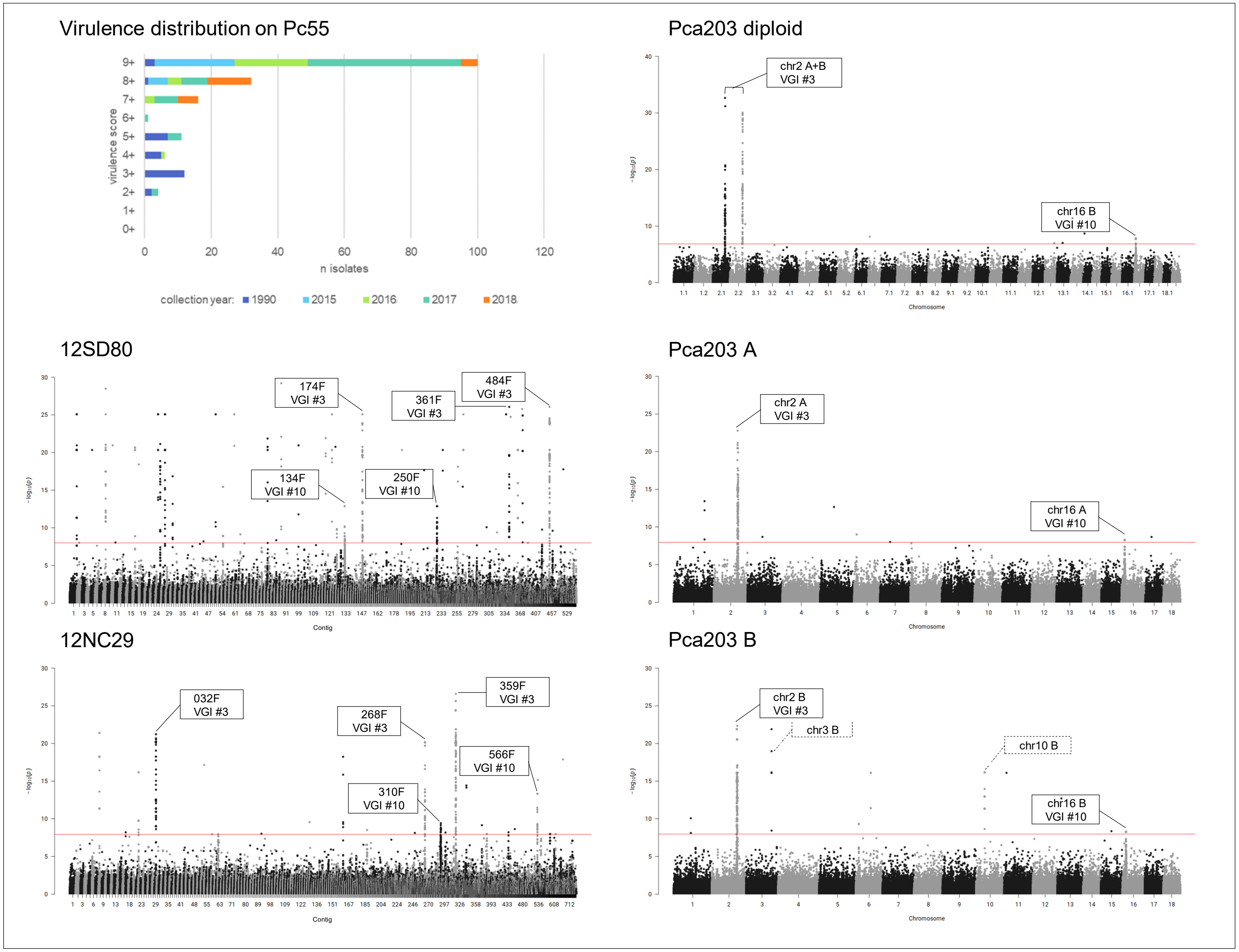

**Fig 22A. quantile-quantile plots of SNP association values for virulence of *Pca* to oat line *Pc56*.** Red lines indicate expected P-value distributions if P-values follow a uniform distribution (null hypothesis).

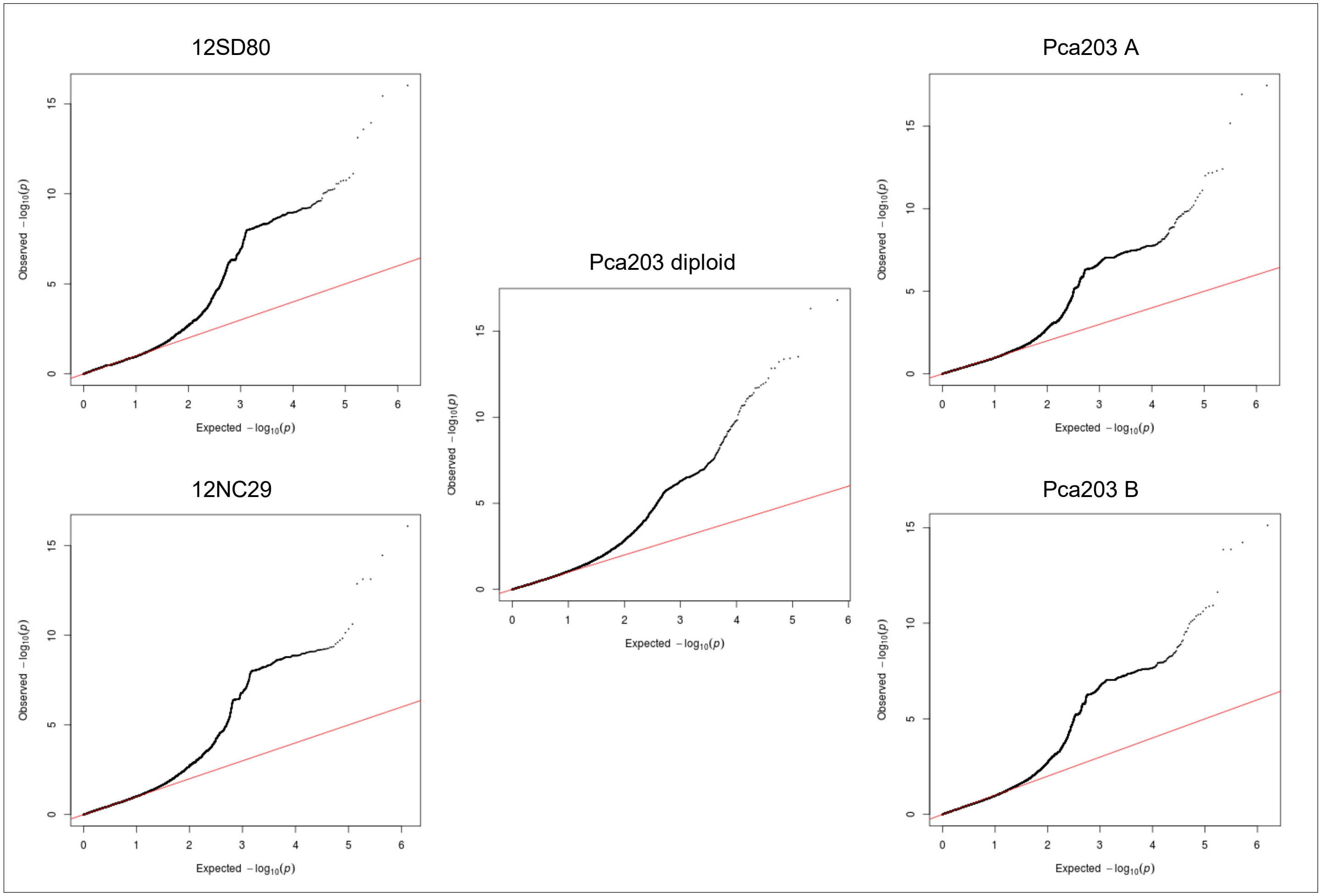

**Fig 22B. *Pca* isolate virulence score distribution and Manhattan plots showing SNP association values for virulence to oat line *Pc56*.** Red horizontal lines denote Bonferroni significance threshold (α = 0.01/total number of markers).

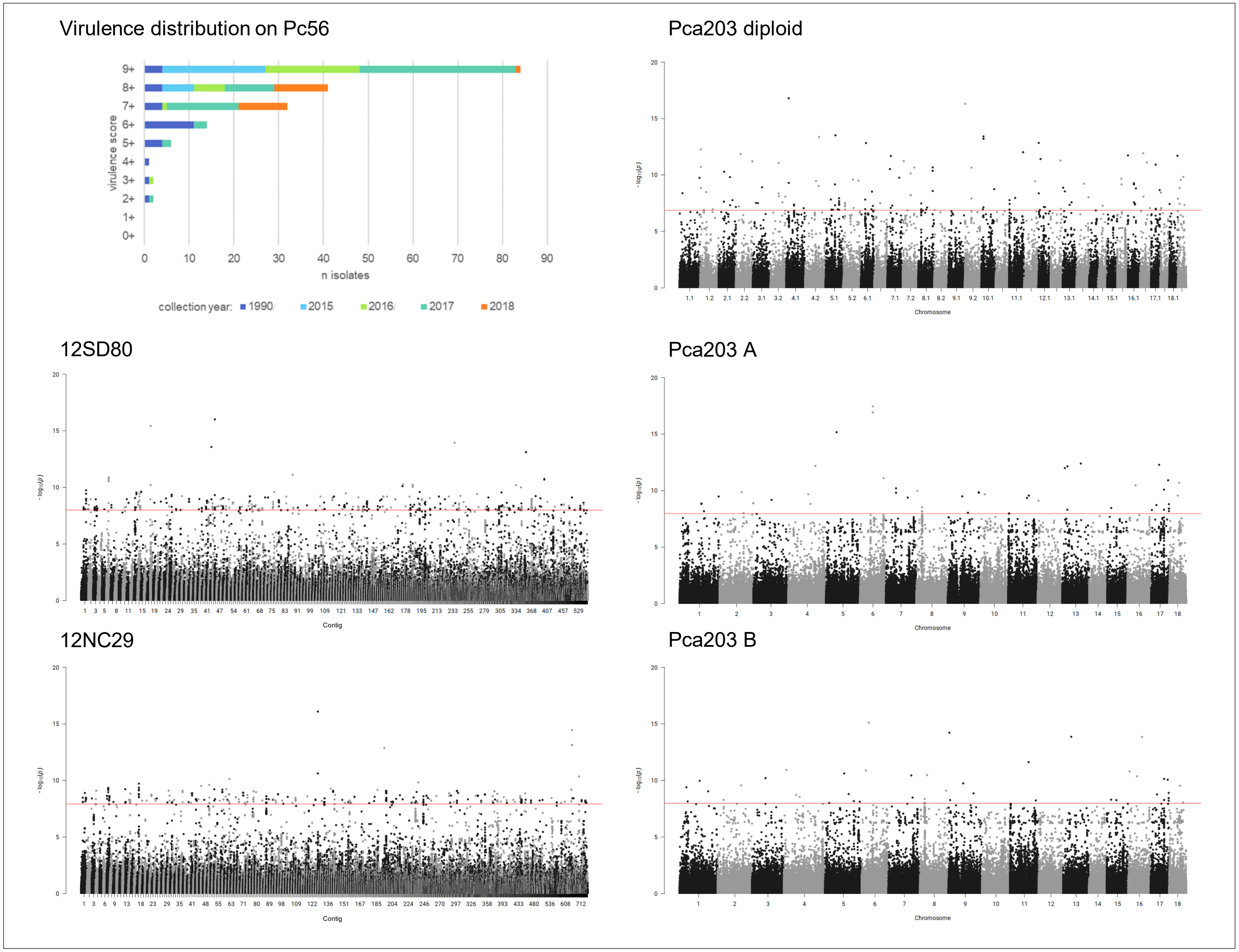

**Fig 23A. quantile-quantile plots of SNP association values for virulence of *Pca* to oat line *Pc57*.** Red lines indicate expected P-value distributions if P-values follow a uniform distribution (null hypothesis).

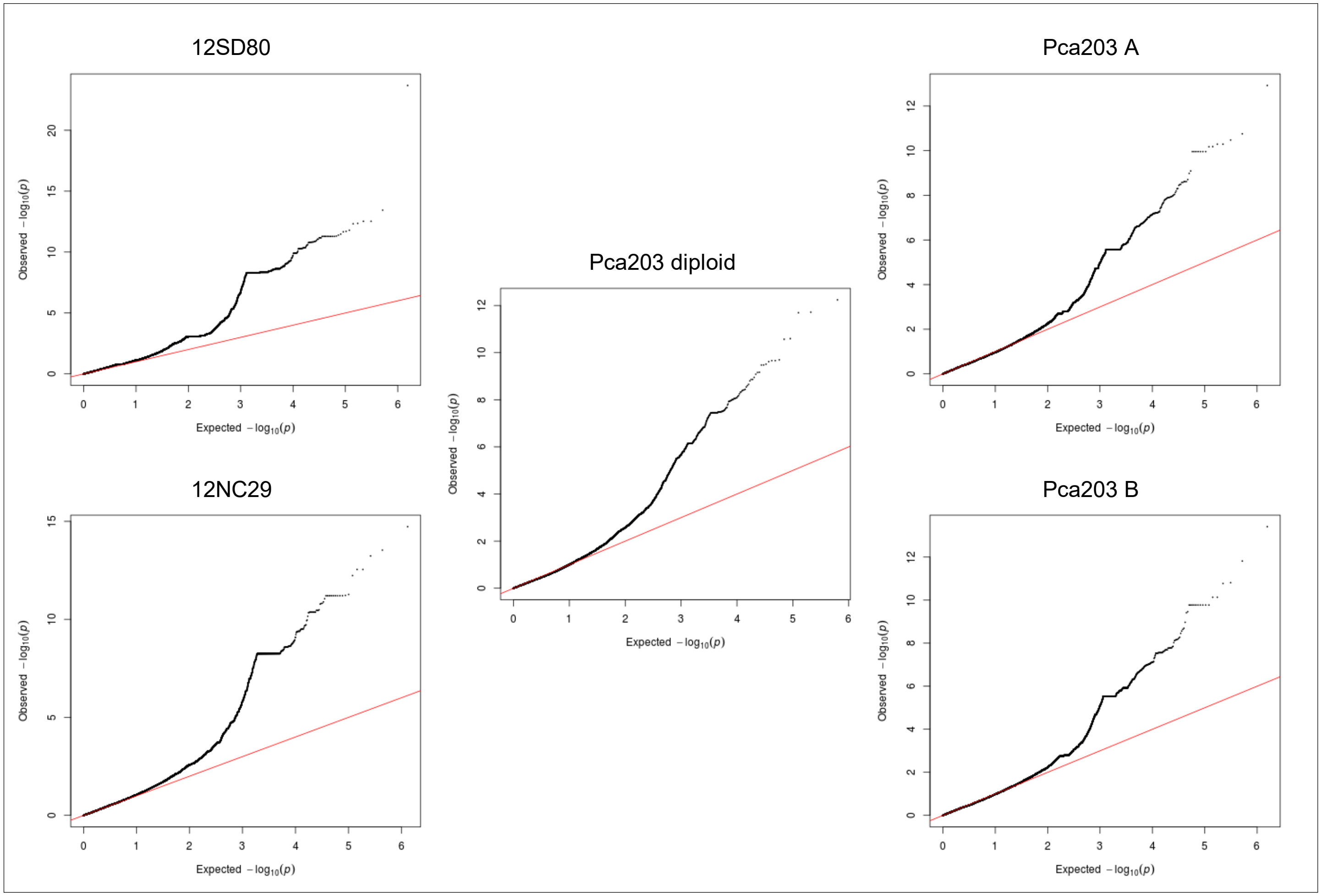

**Fig 23B. *Pca* isolate virulence score distribution and Manhattan plots showing SNP association values for virulence to oat line *Pc57*.** Red and blue horizontal lines denote Bonferroni significance threshold (α = 0.01/total number of markers) and 5% false discovery rate threshold, respectively. Significant association peaks are labelled with contig or chromosome number and assigned VGI number.

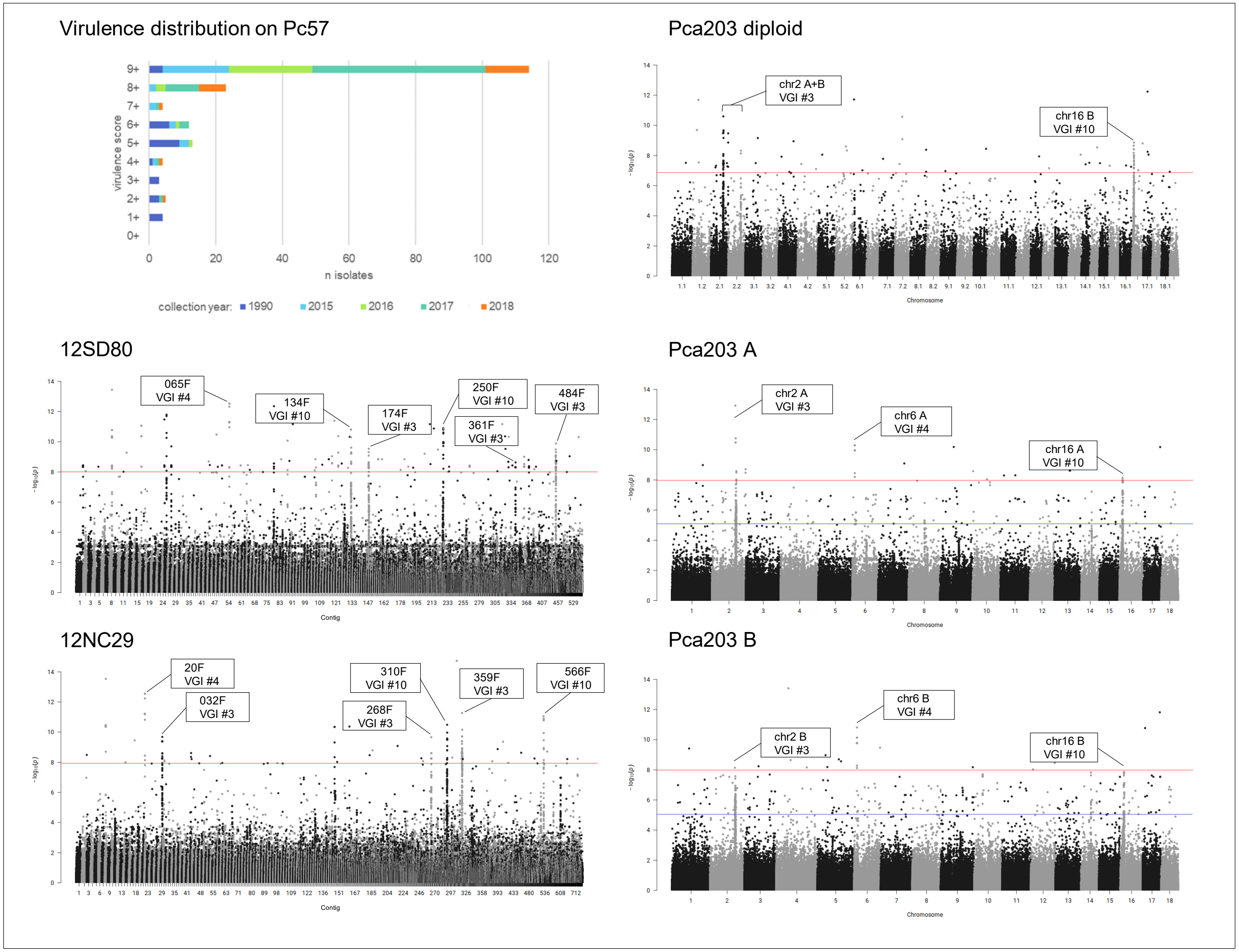

**Fig 24A. quantile-quantile plots of SNP association values for virulence of *Pca* to oat line *Pc58*.** Red lines indicate expected P-value distributions if P-values follow a uniform distribution (null hypothesis).

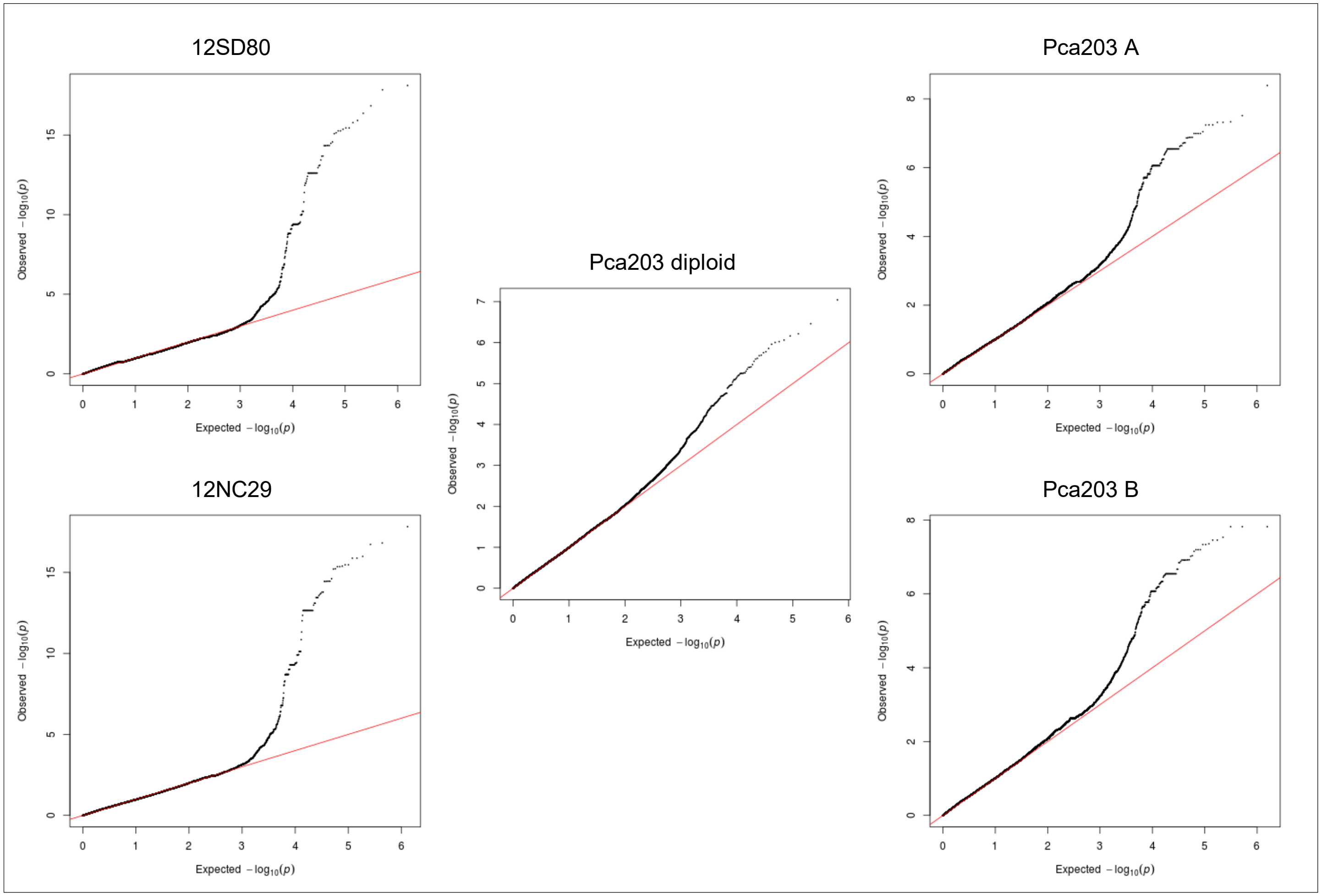

**Fig 24B. *Pca* isolate virulence score distribution and Manhattan plots showing SNP association values for virulence to oat line *Pc58*.** Red and blue horizontal lines denote Bonferroni significance threshold (α = 0.01/total number of markers) and 5% false discovery rate threshold, respectively. Significant association peaks are labelled with contig or chromosome number and assigned VGI number.

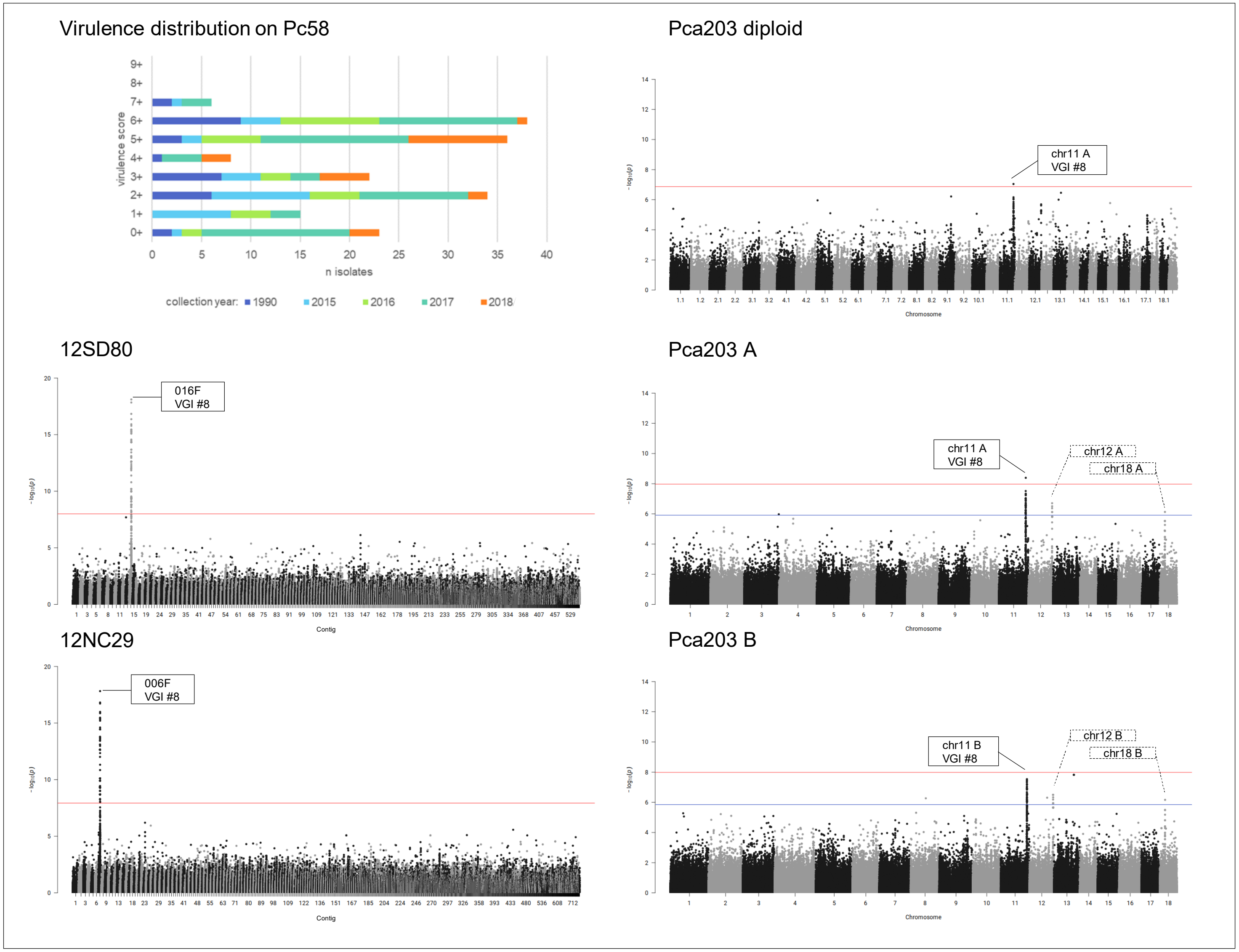

**Fig 25A. quantile-quantile plots of SNP association values for virulence of *Pca* to oat line *Pc59*.** Red lines indicate expected P-value distributions if P-values follow a uniform distribution (null hypothesis).

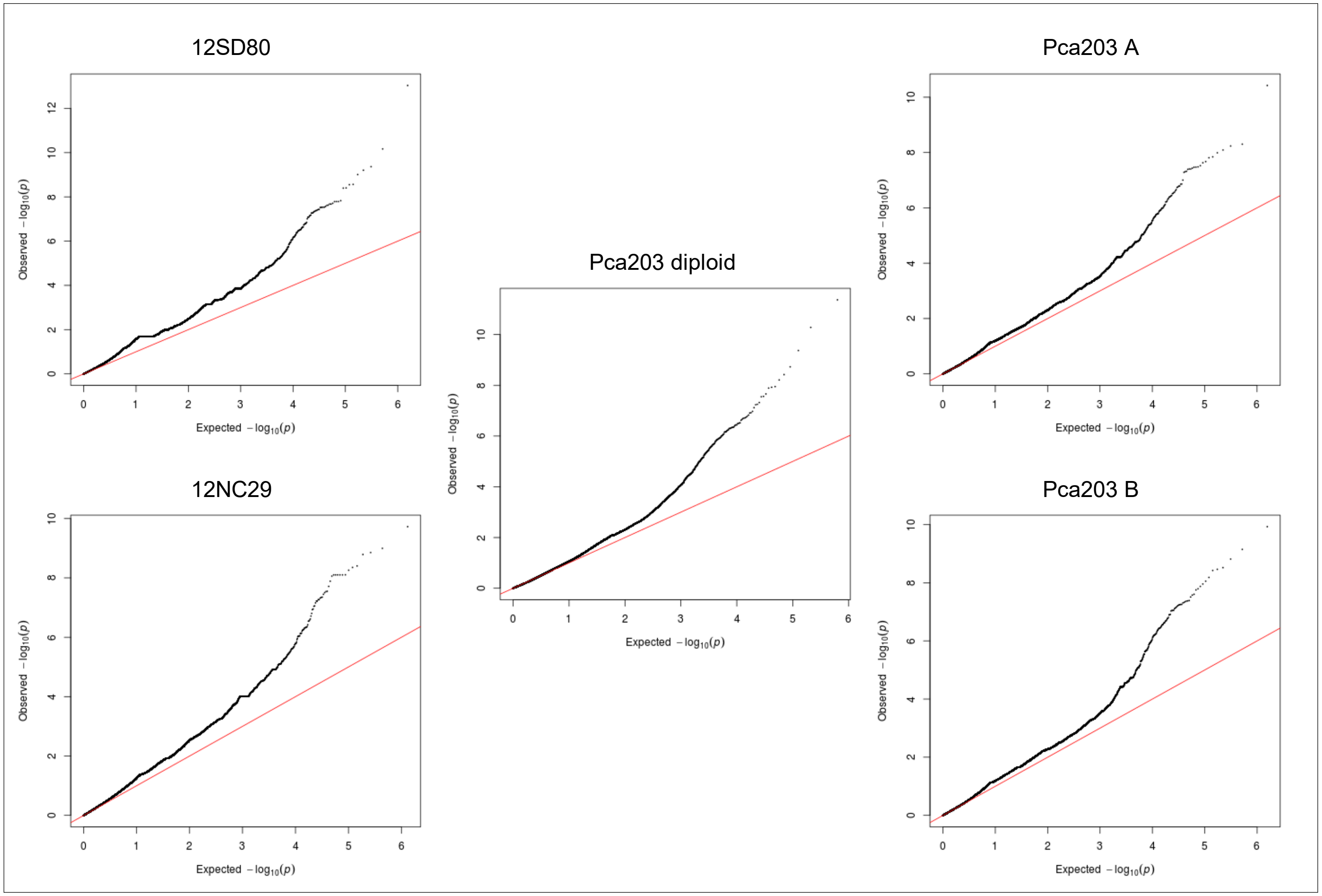

**Fig 25B. *Pca* isolate virulence score distribution and Manhattan plots showing SNP association values for virulence to oat line *Pc59*.** Red horizontal lines denote Bonferroni significance threshold (α = 0.01/total number of markers).

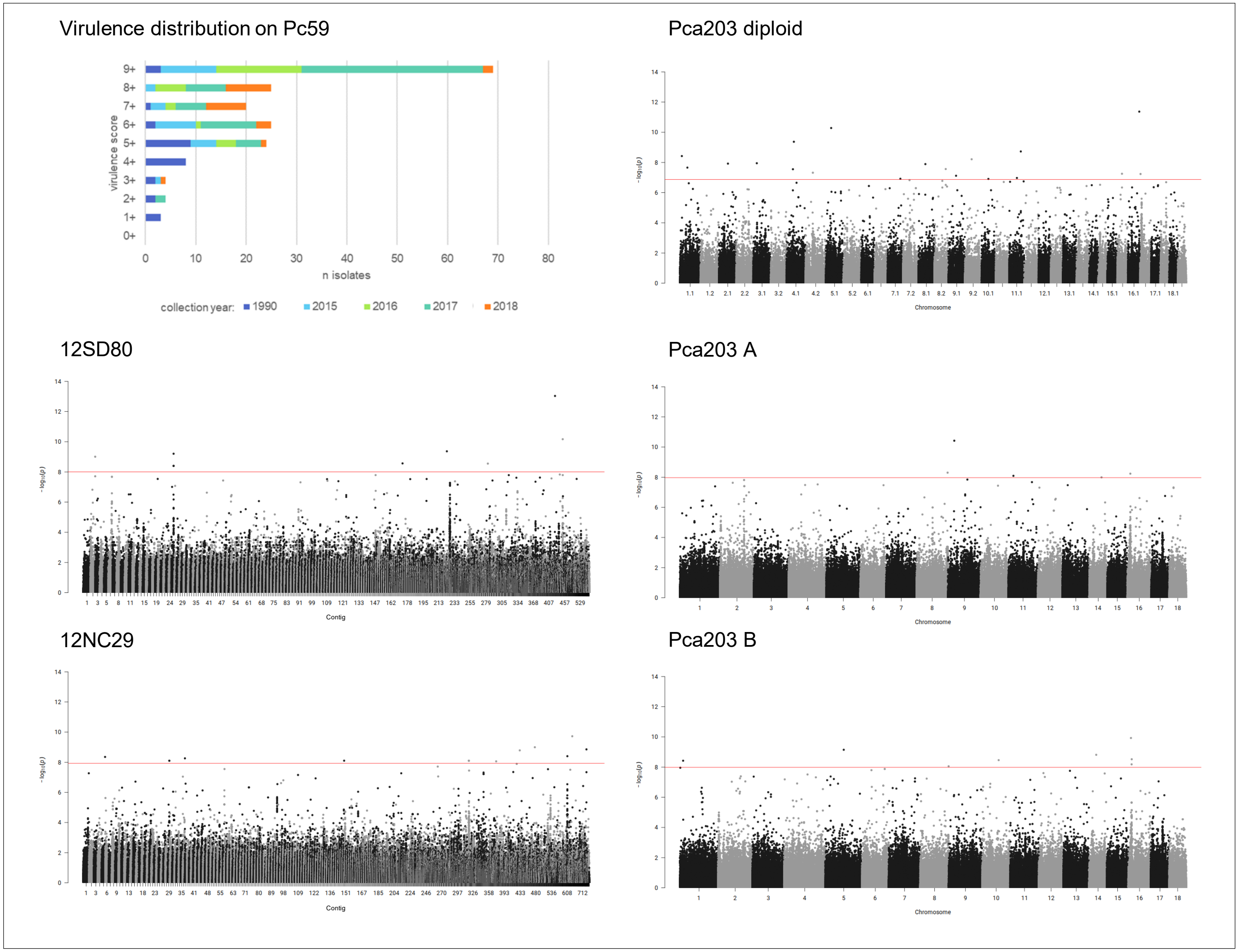

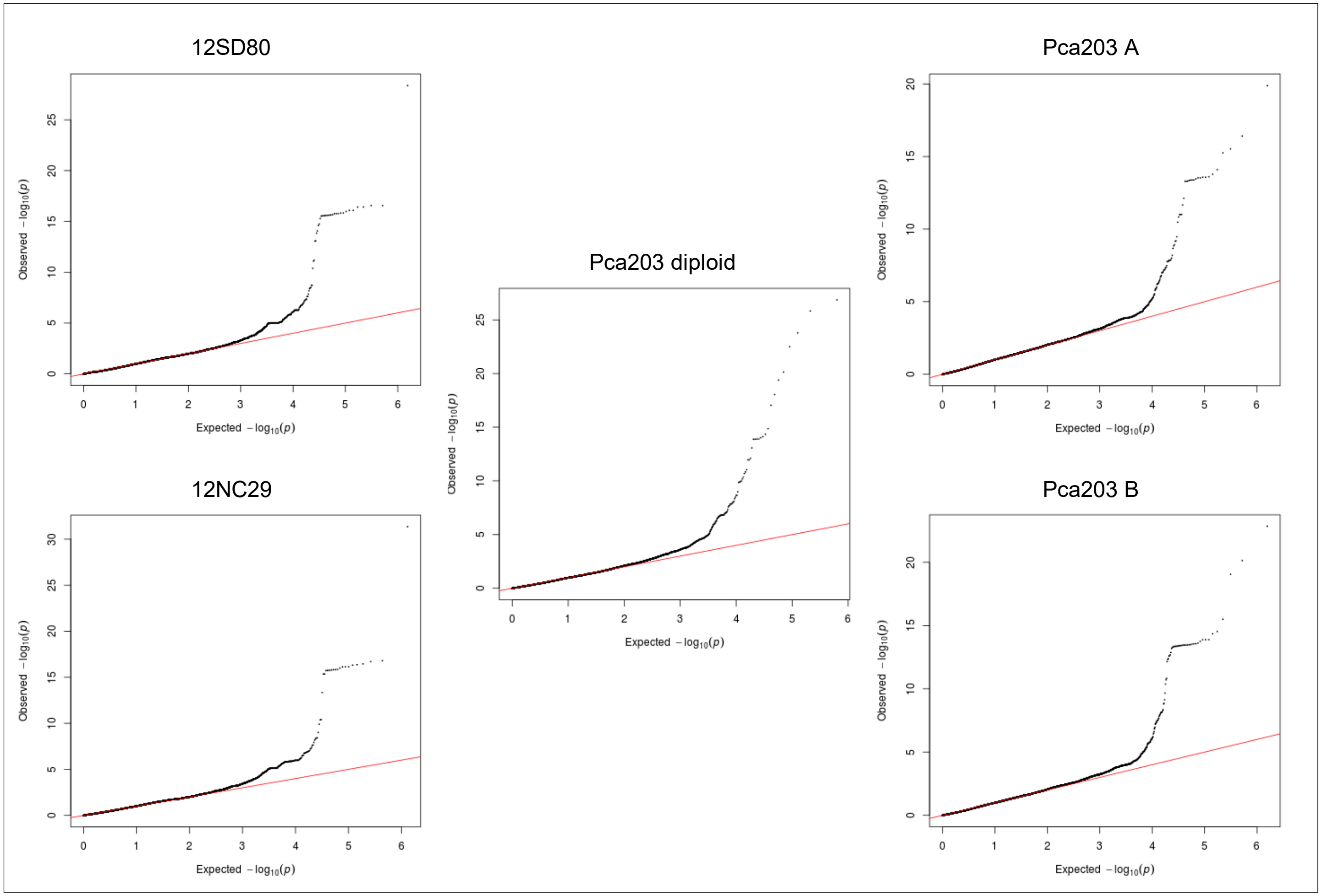


**Fig 26A. quantile-quantile plots of SNP association values for virulence of *Pca* to oat line *Pc60*.** Red lines indicate expected P-value distributions if P-values follow a uniform distribution (null hypothesis).

**Fig 26B. *Pca* isolate virulence score distribution and Manhattan plots showing SNP association values for virulence to oat line *Pc60*.** Red and blue horizontal lines denote Bonferroni significance threshold (α = 0.01/total number of markers) and 5% false discovery rate threshold, respectively. Potentially significant association peaks are labelled with contig or chromosome number.

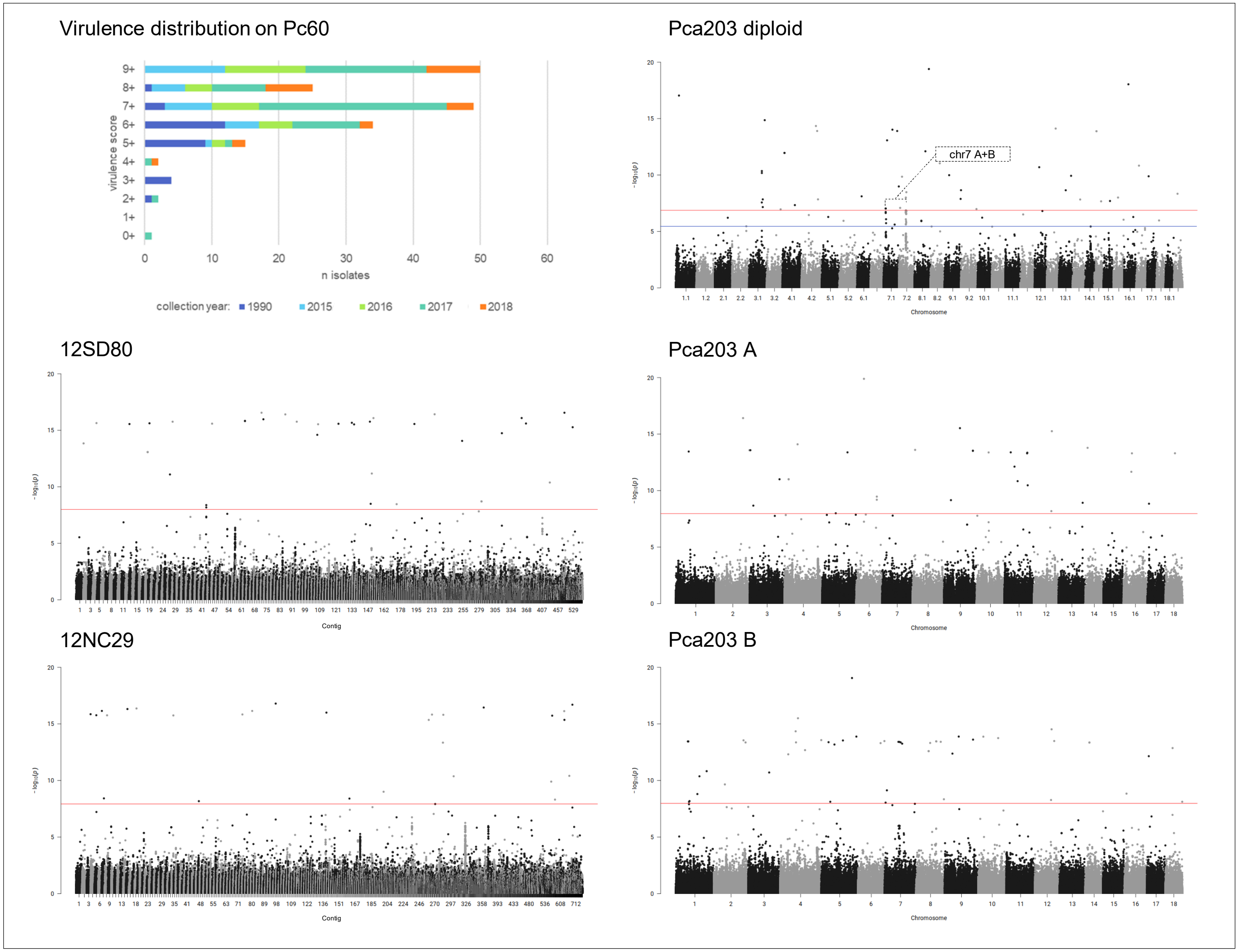

**Fig 27A. quantile-quantile plots of SNP association values for virulence of *Pca* to oat line *Pc61*.** Red lines indicate expected P-value distributions if P-values follow a uniform distribution (null hypothesis).

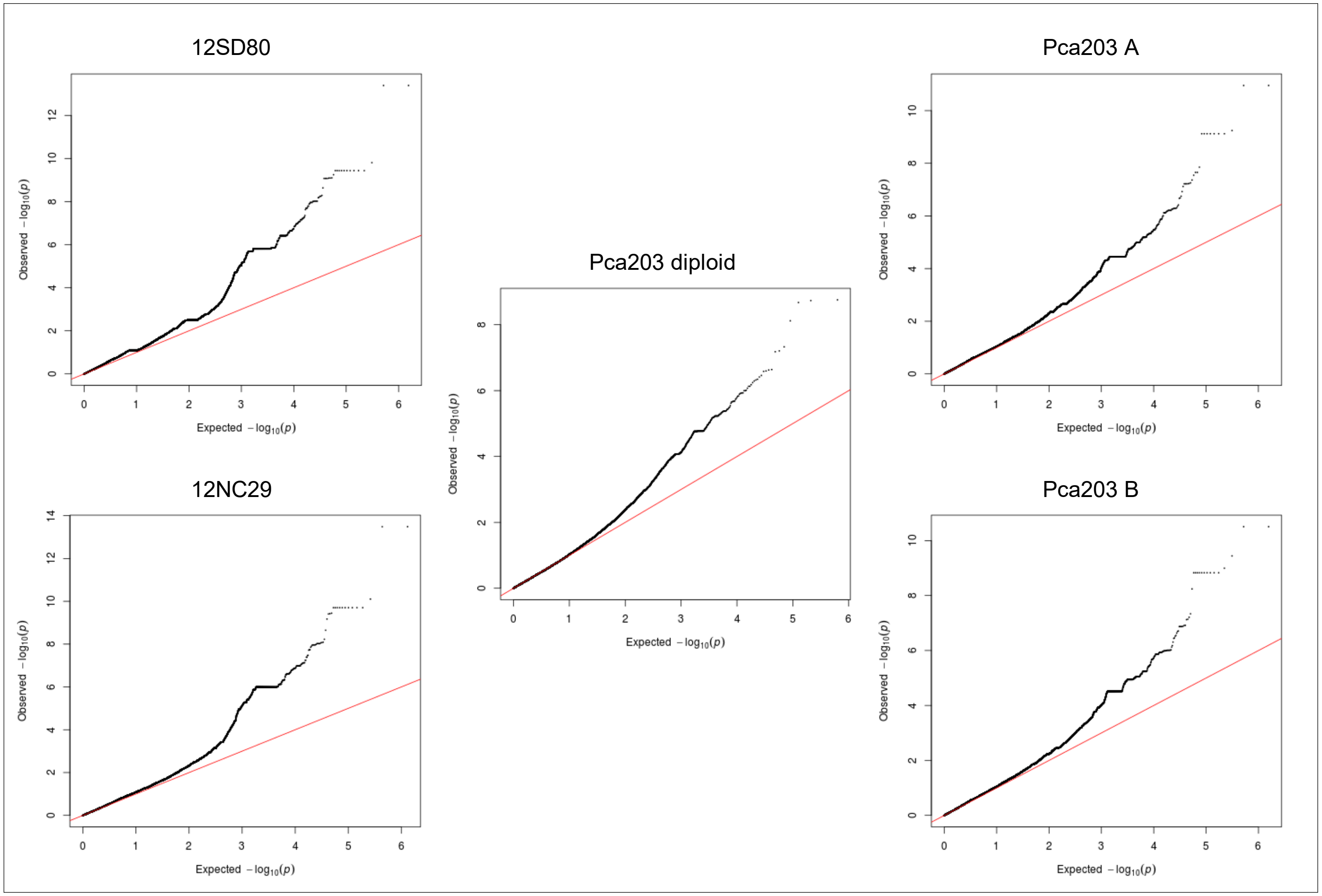

**Fig 27B. *Pca* isolate virulence score distribution and Manhattan plots showing SNP association values for virulence to oat line *Pc61*.** Red and blue horizontal lines denote Bonferroni significance threshold (α = 0.01/total number of markers) and 5% false discovery rate threshold, respectively. Significant association peaks are labelled with contig or chromosome number and assigned VGI number.

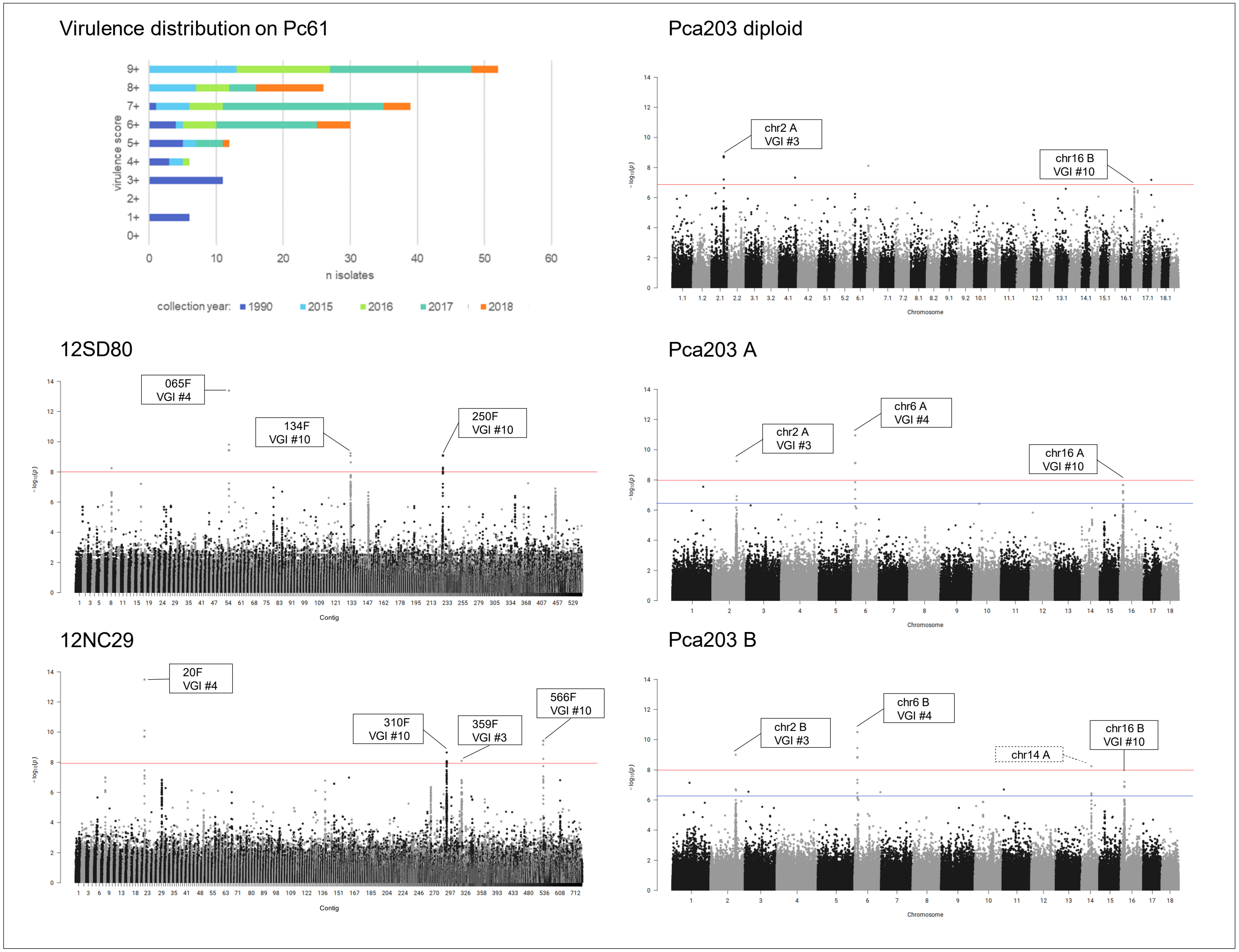

**Fig 28A. quantile-quantile plots of SNP association values for virulence of *Pca* to oat line *Pc62*.** Red lines indicate expected P-value distributions if P-values follow a uniform distribution (null hypothesis).

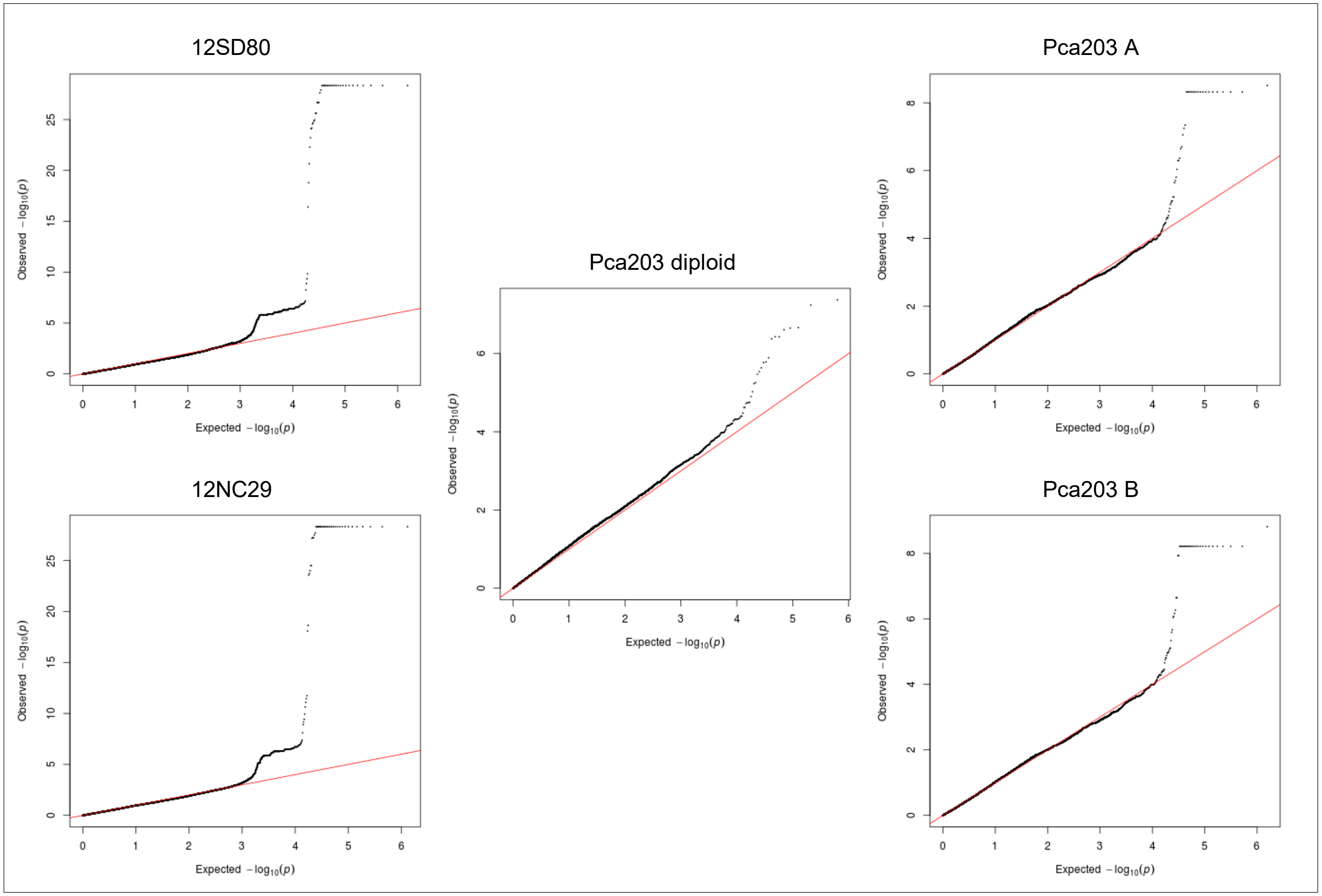

**Fig 28B. *Pca* isolate virulence score distribution and Manhattan plots showing SNP association values for virulence to oat line *Pc62*.** Red horizontal lines denote Bonferroni significance threshold (α = 0.01/total number of markers). Significant association peaks are labelled with contig or chromosome number and assigned VGI number.

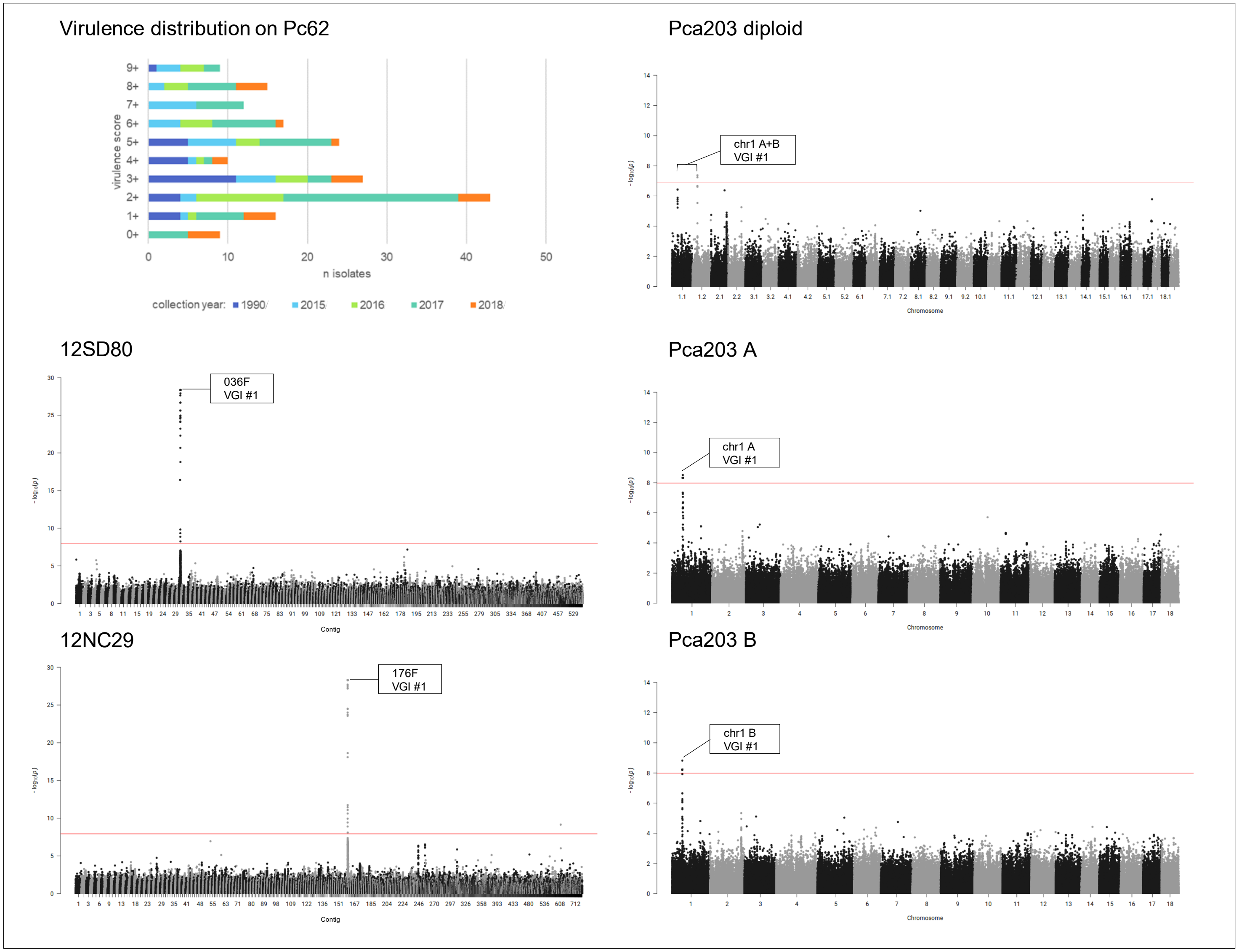

**Fig 29A. quantile-quantile plots of SNP association values for virulence of *Pca* to oat line *Pc63*.** Red lines indicate expected P-value distributions if P-values follow a uniform distribution (null hypothesis).

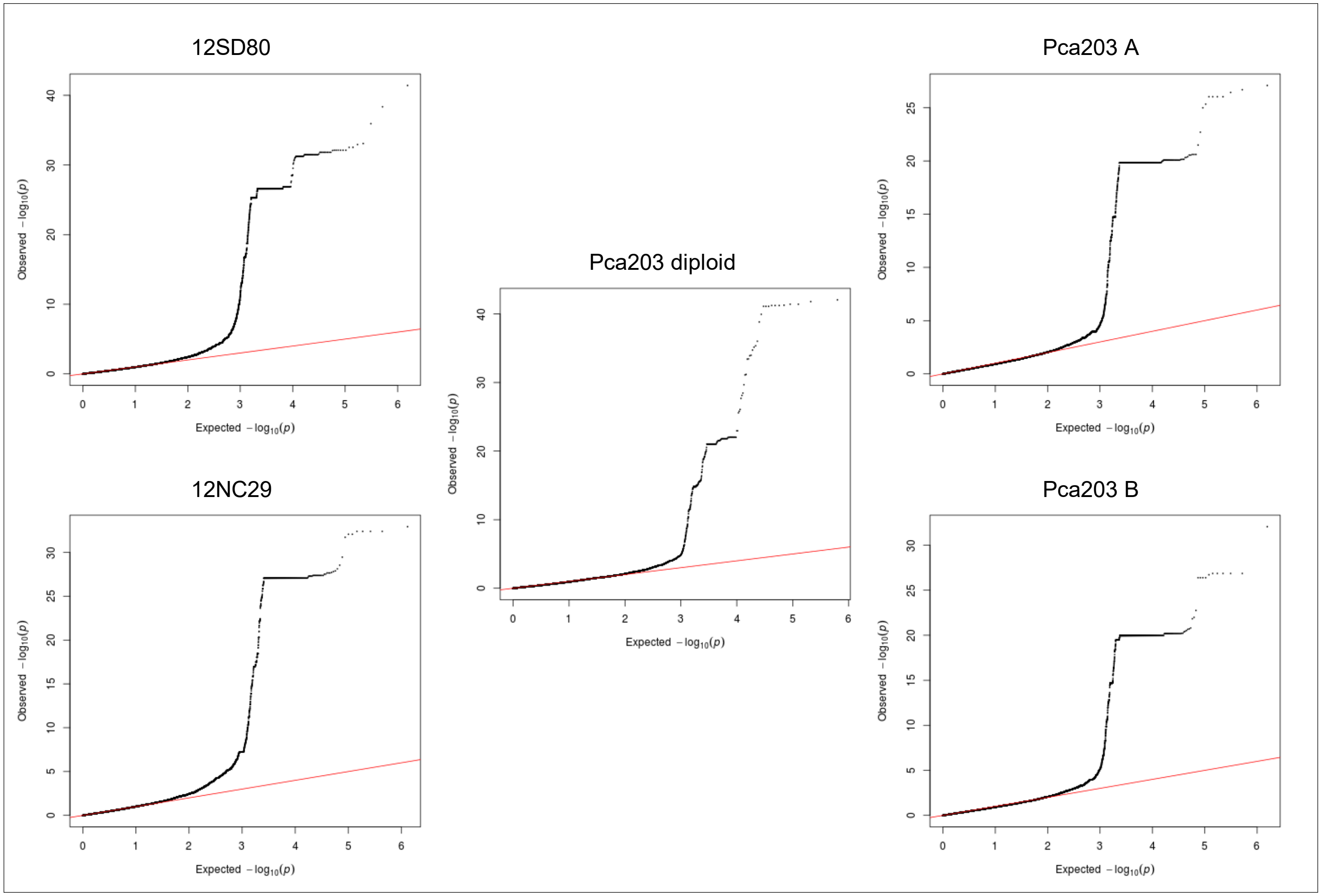

**Fig 29B. *Pca* isolate virulence score distribution and Manhattan plots showing SNP association values for virulence to oat line *Pc63*.** Red and blue horizontal lines denote Bonferroni significance threshold (α = 0.01/total number of markers) and 5% false discovery rate threshold, respectively. Significant association peaks are labelled with contig or chromosome number and assigned VGI number.

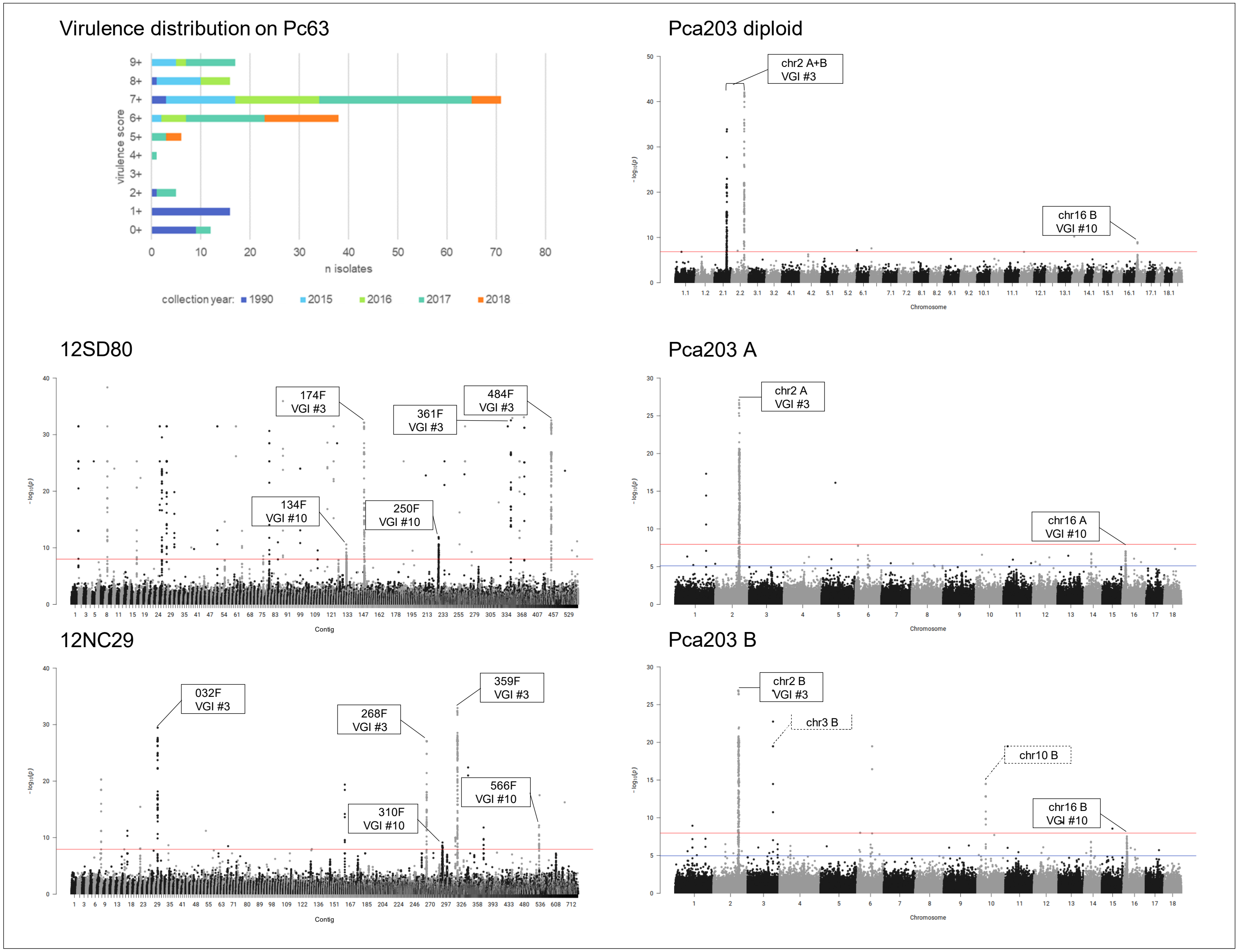

**Fig 30A. quantile-quantile plots of SNP association values for virulence of *Pca* to oat line *Pc64*.** Red lines indicate expected P-value distributions if P-values follow a uniform distribution (null hypothesis).

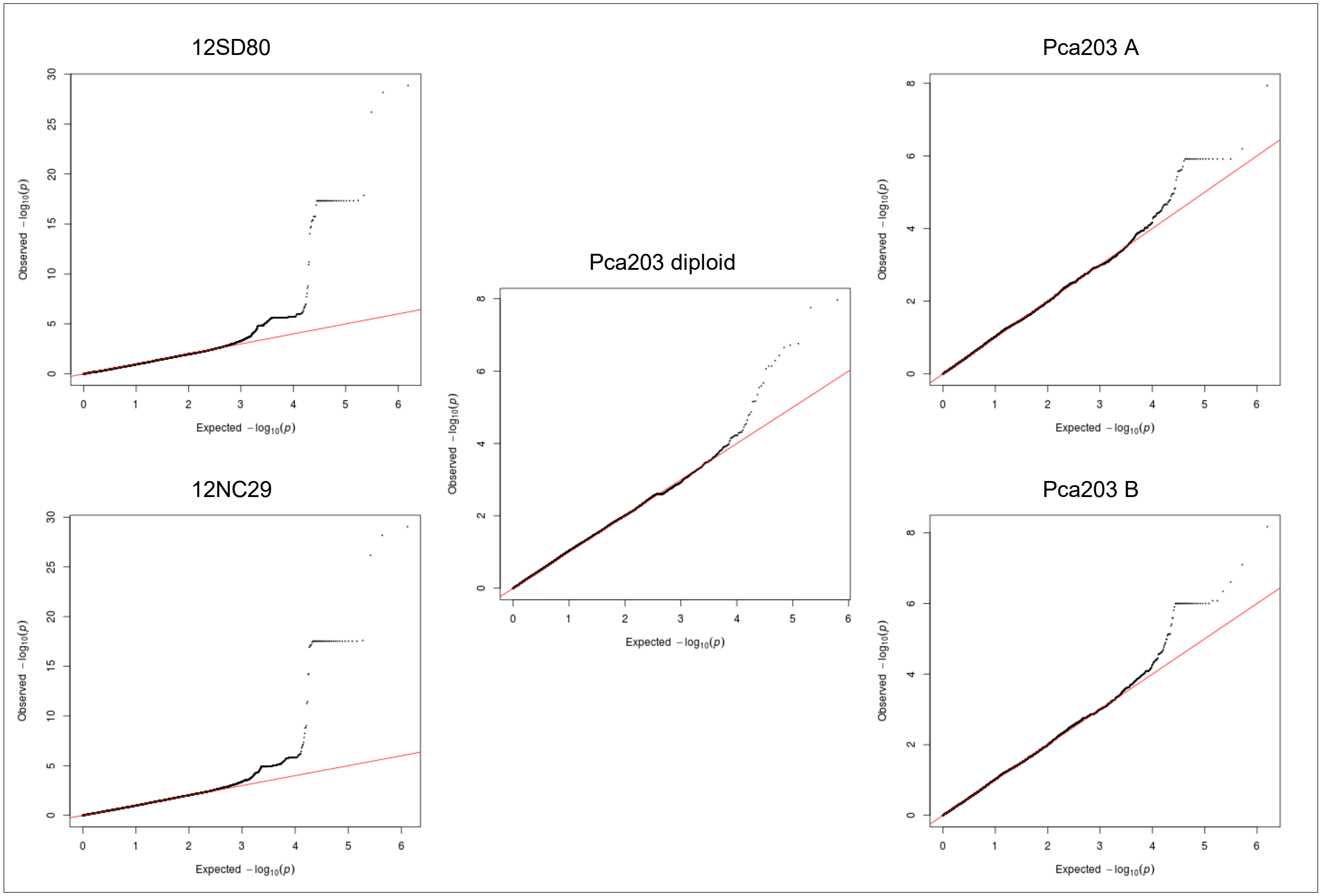

**Fig 30B. *Pca* isolate virulence score distribution and Manhattan plots showing SNP association values for virulence to oat line *Pc64*.** Red and blue horizontal lines denote Bonferroni significance threshold (α = 0.01/total number of markers) and 5% false discovery rate threshold, respectively. Significant association peaks are labelled with contig or chromosome number and assigned VGI number.

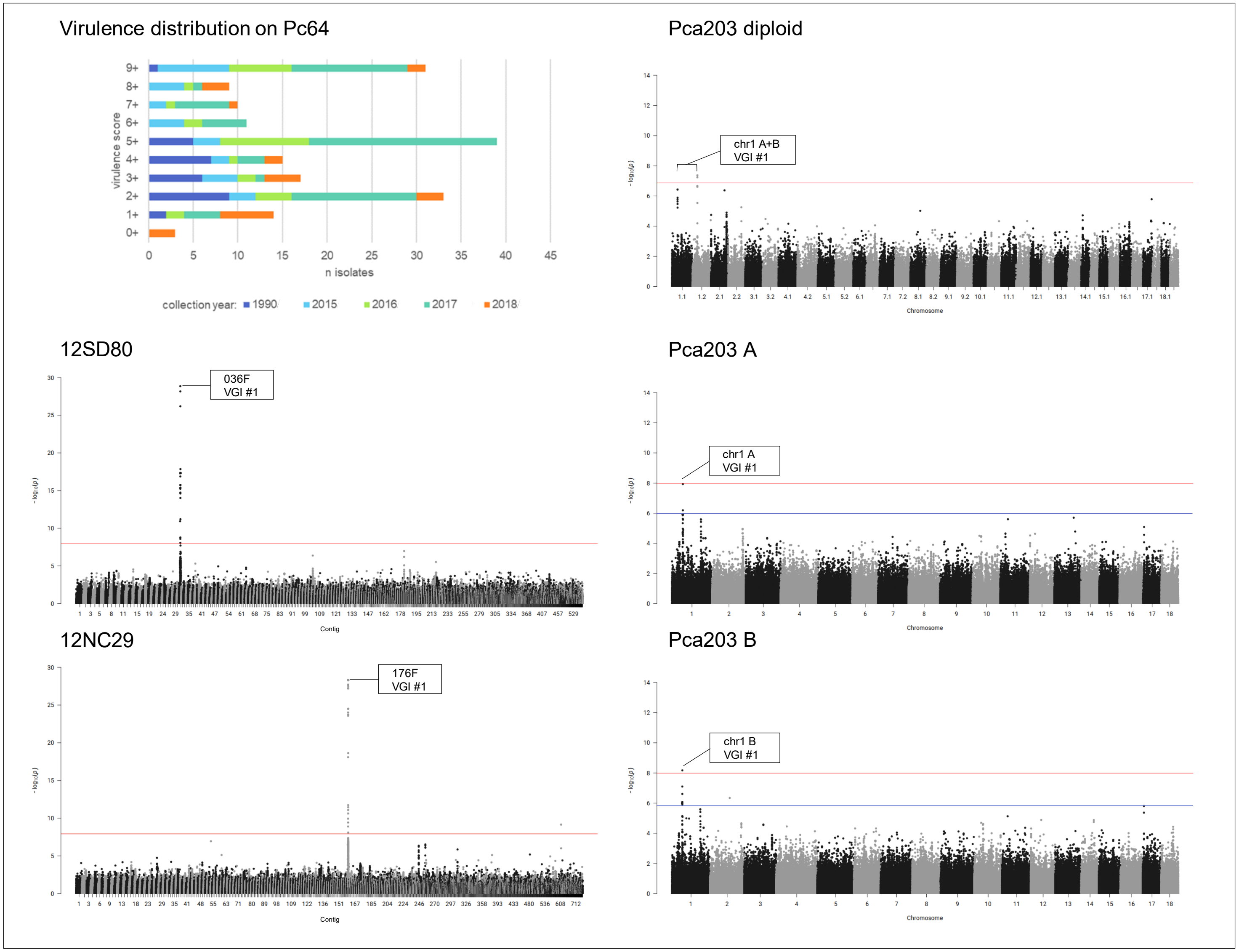

**Fig 31A. quantile-quantile plots of SNP association values for virulence of *Pca* to oat line *Pc67*.** Red lines indicate expected P-value distributions if P-values follow a uniform distribution (null hypothesis).

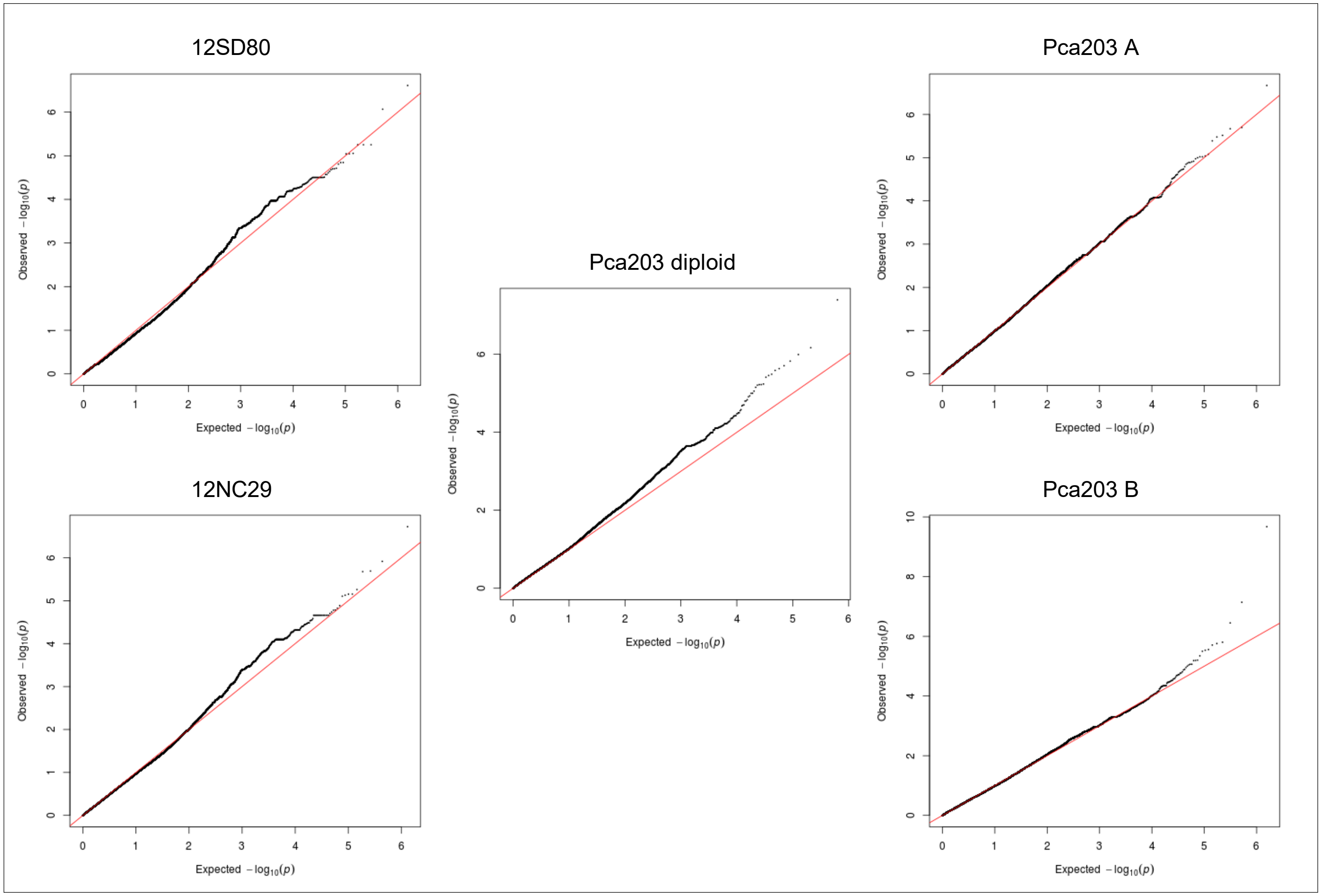

**Fig 31B. *Pca* isolate virulence score distribution and Manhattan plots showing SNP association values for virulence to oat line *Pc67*.** Red horizontal lines denote Bonferroni significance threshold (α = 0.01/total number of markers).

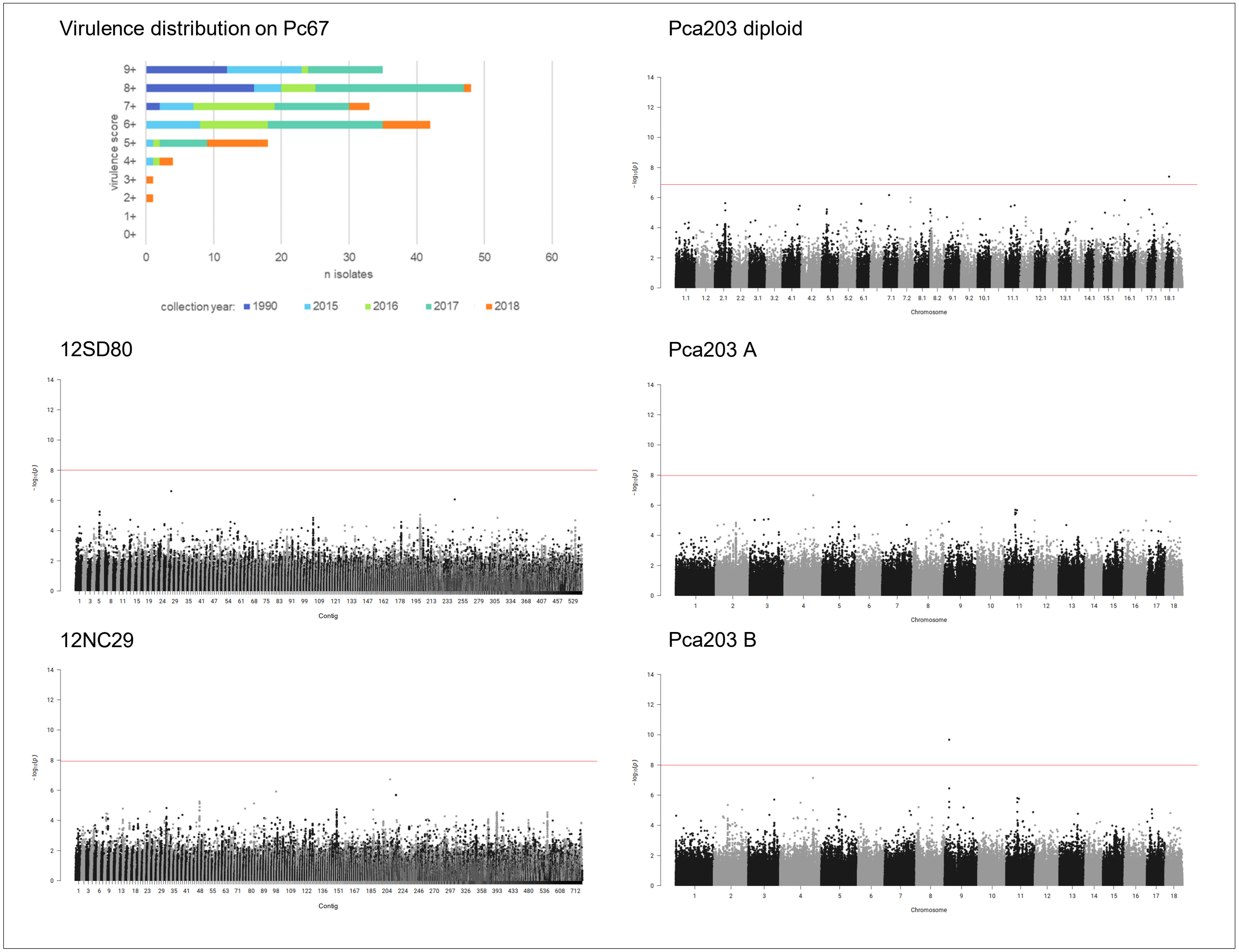

**Fig 32A. quantile-quantile plots of SNP association values for virulence of *Pca* to oat line *Pc68*.** Red lines indicate expected P-value distributions if P-values follow a uniform distribution (null hypothesis).

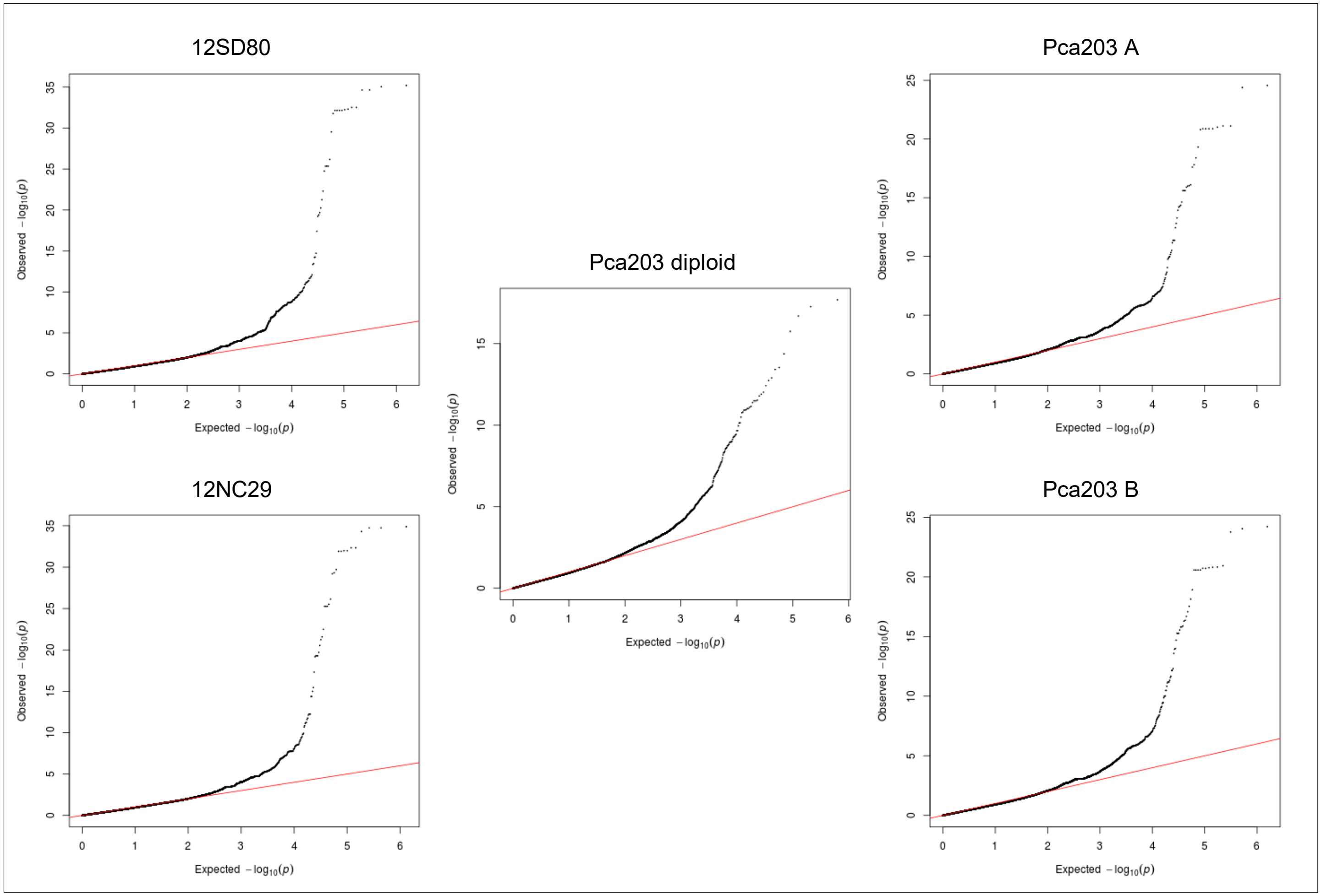

**Fig 32B. *Pca* isolate virulence score distribution and Manhattan plots showing SNP association values for virulence to oat line *Pc68*.** Red and blue horizontal lines denote Bonferroni significance threshold (α = 0.01/total number of markers) and 5% false discovery rate threshold, respectively. Significant association peaks are labelled with contig or chromosome number and assigned VGI number.

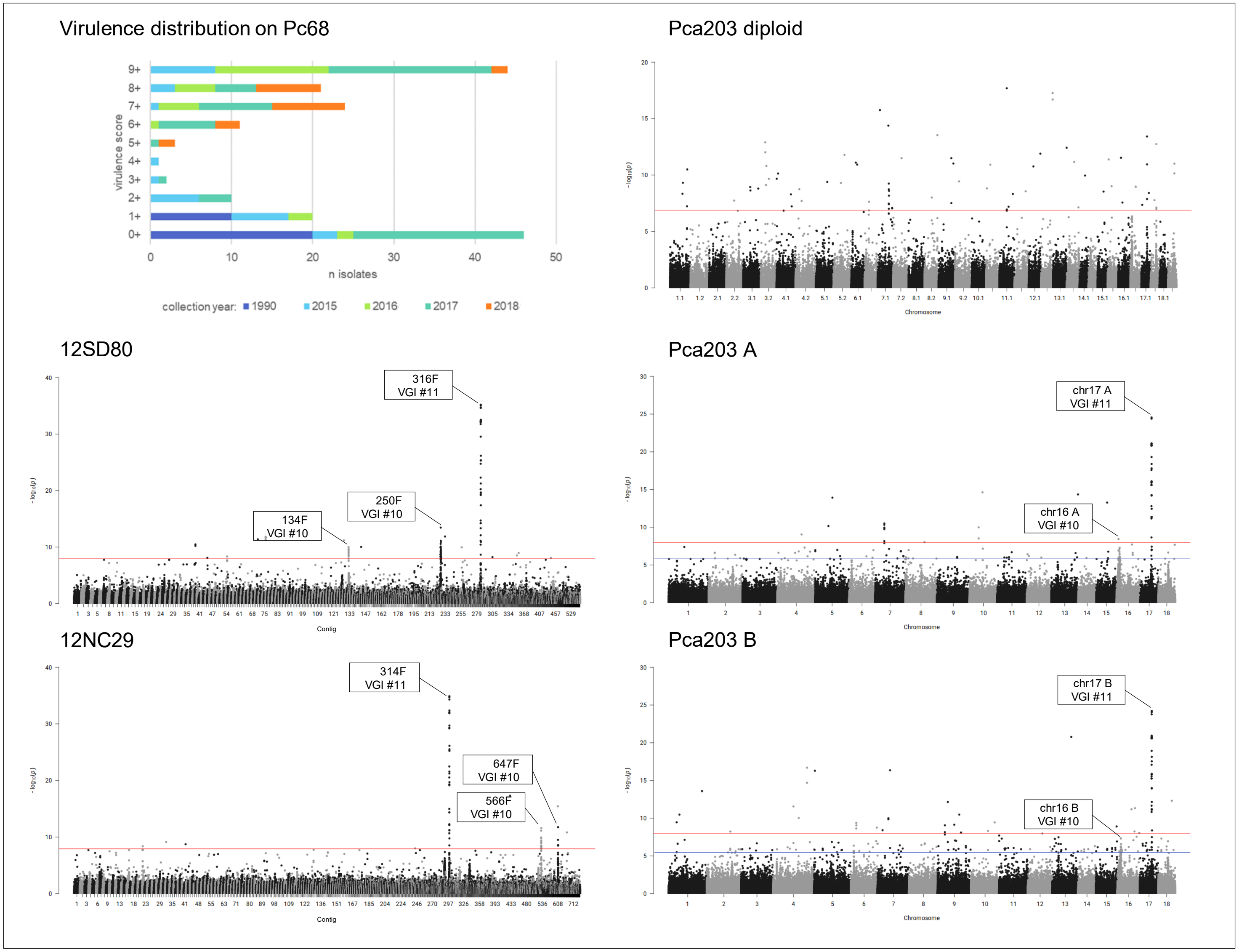

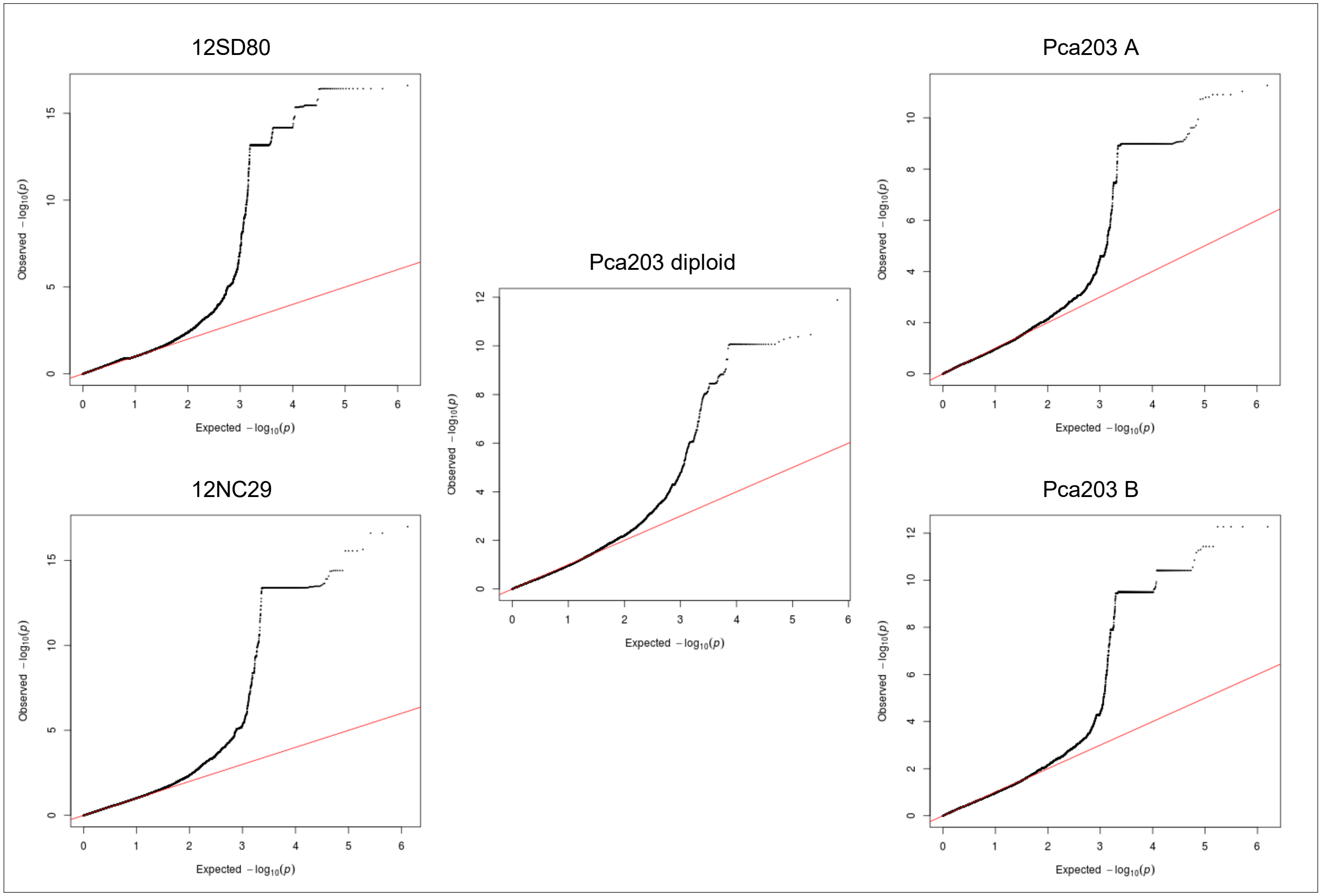


**Fig 33A. quantile-quantile plots of SNP association values for virulence of *Pca* to oat line *Pc70*.** Red lines indicate expected P-value distributions if P-values follow a uniform distribution (null hypothesis).

**Fig 33B. *Pca* isolate virulence score distribution and Manhattan plots showing SNP association values for virulence to oat line *Pc70*.** Red horizontal lines denote Bonferroni significance threshold (α = 0.01/total number of markers). Significant association peaks are labelled with contig or chromosome number and assigned VGI number.

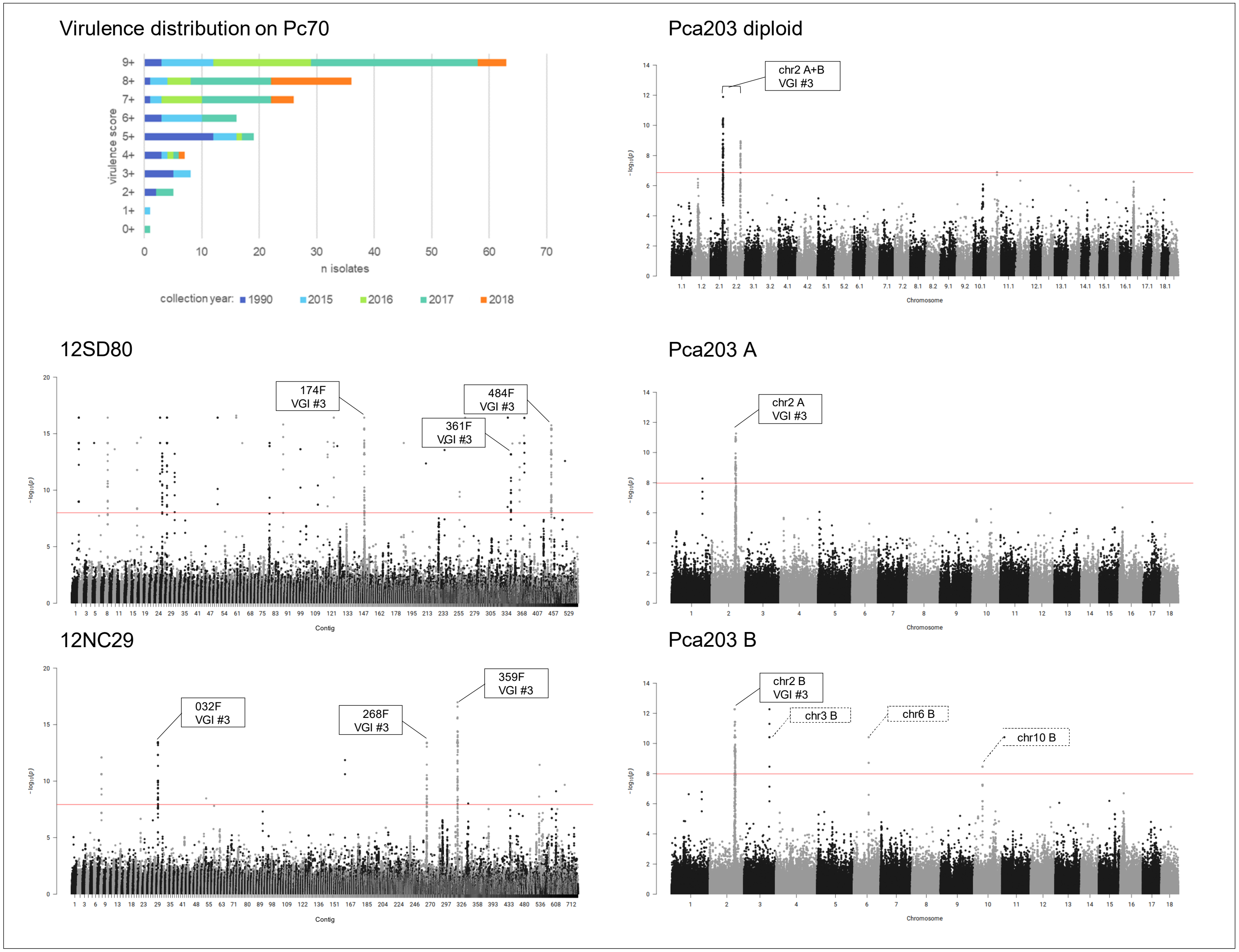

**Fig 34A. quantile-quantile plots of SNP association values for virulence of *Pca* to oat line *Pc71*.** Red lines indicate expected P-value distributions if P-values follow a uniform distribution (null hypothesis).

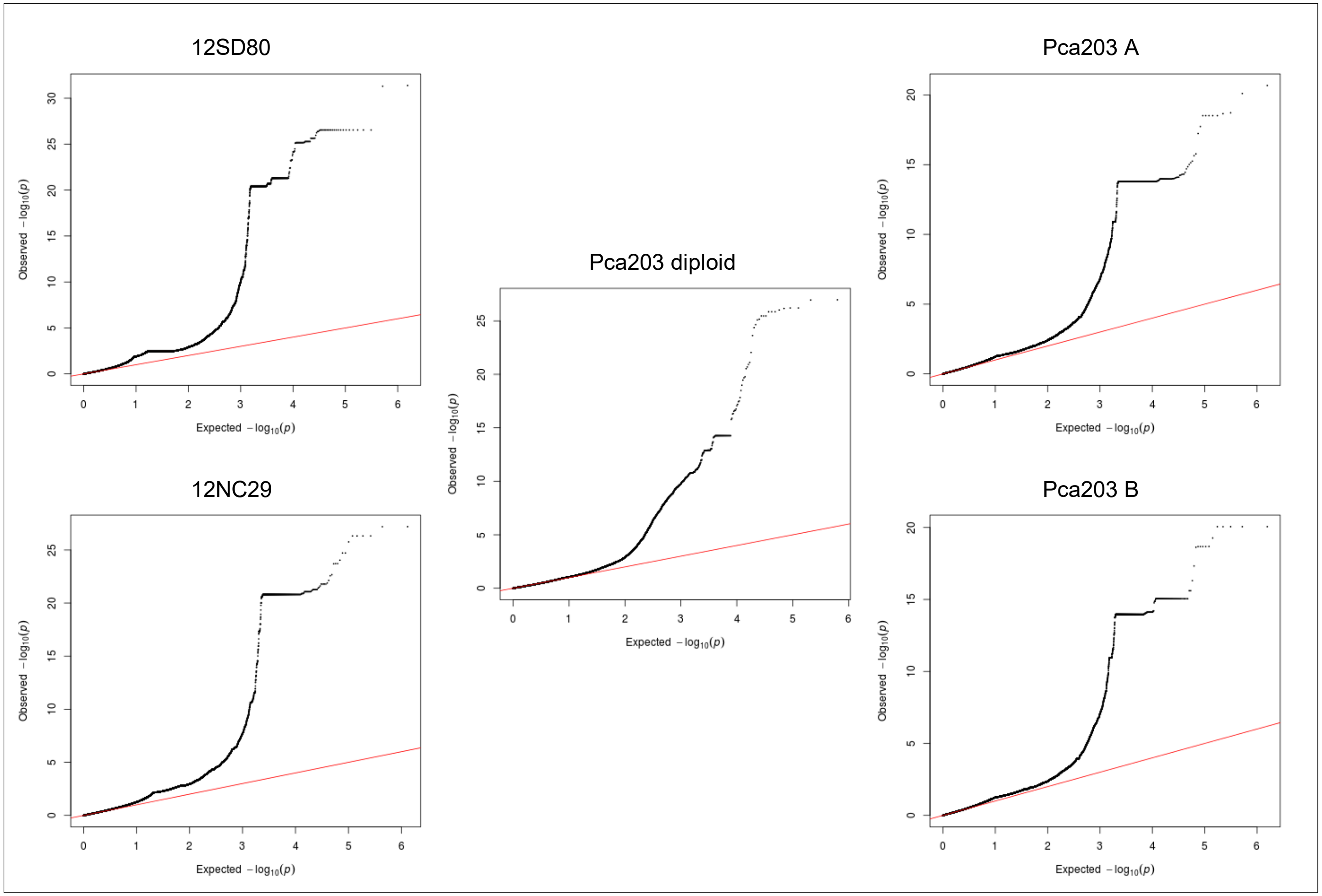

**Fig 34B. *Pca* isolate virulence score distribution and Manhattan plots showing SNP association values for virulence to oat line *Pc71*.** Red horizontal lines denote Bonferroni significance threshold (α = 0.01/total number of markers). Significant association peaks are labelled with contig or chromosome number and assigned VGI number.

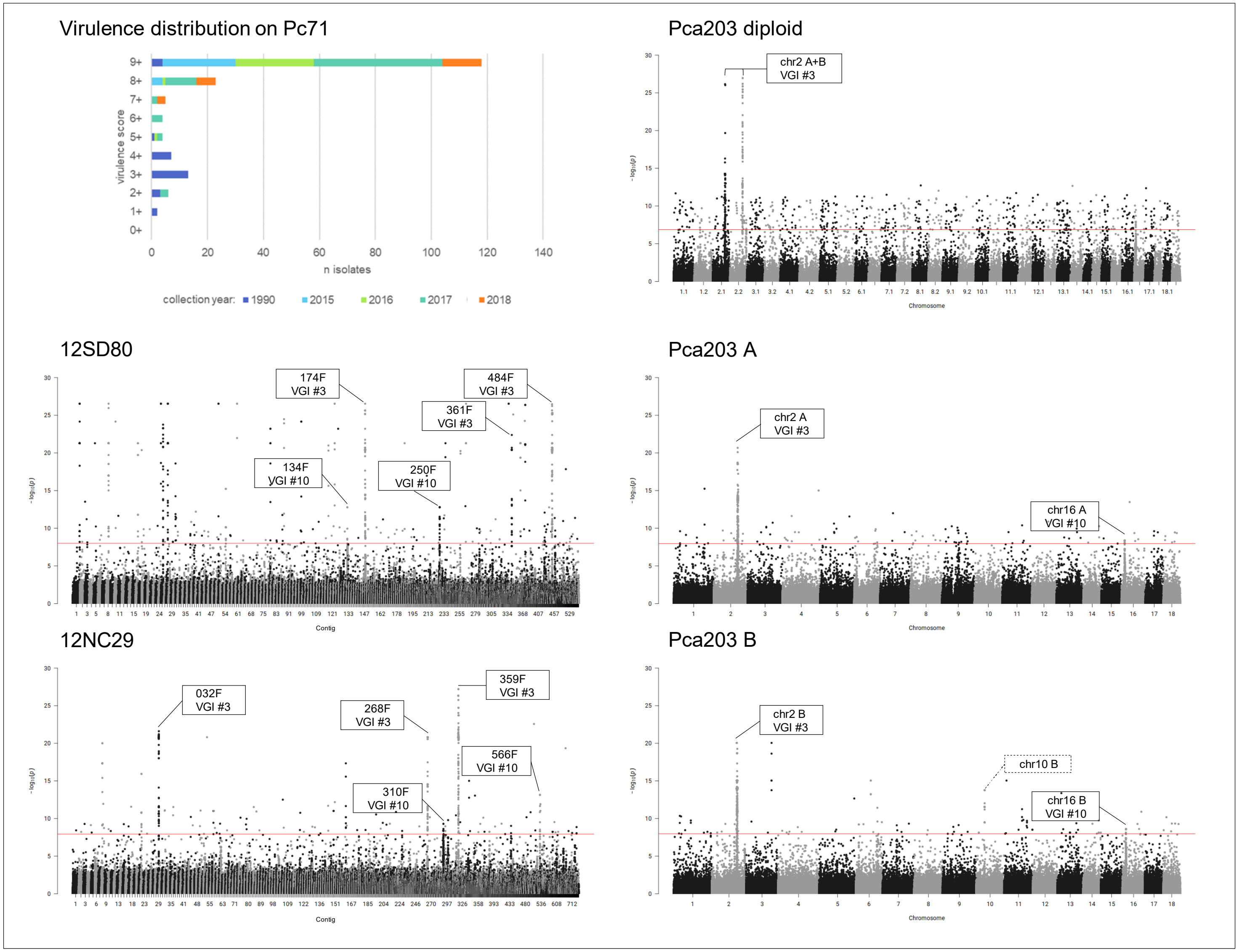

**Fig 35A. quantile-quantile plots of SNP association values for virulence of *Pca* to oat line *Pc91*.** Red lines indicate expected P-value distributions if P-values follow a uniform distribution (null hypothesis).

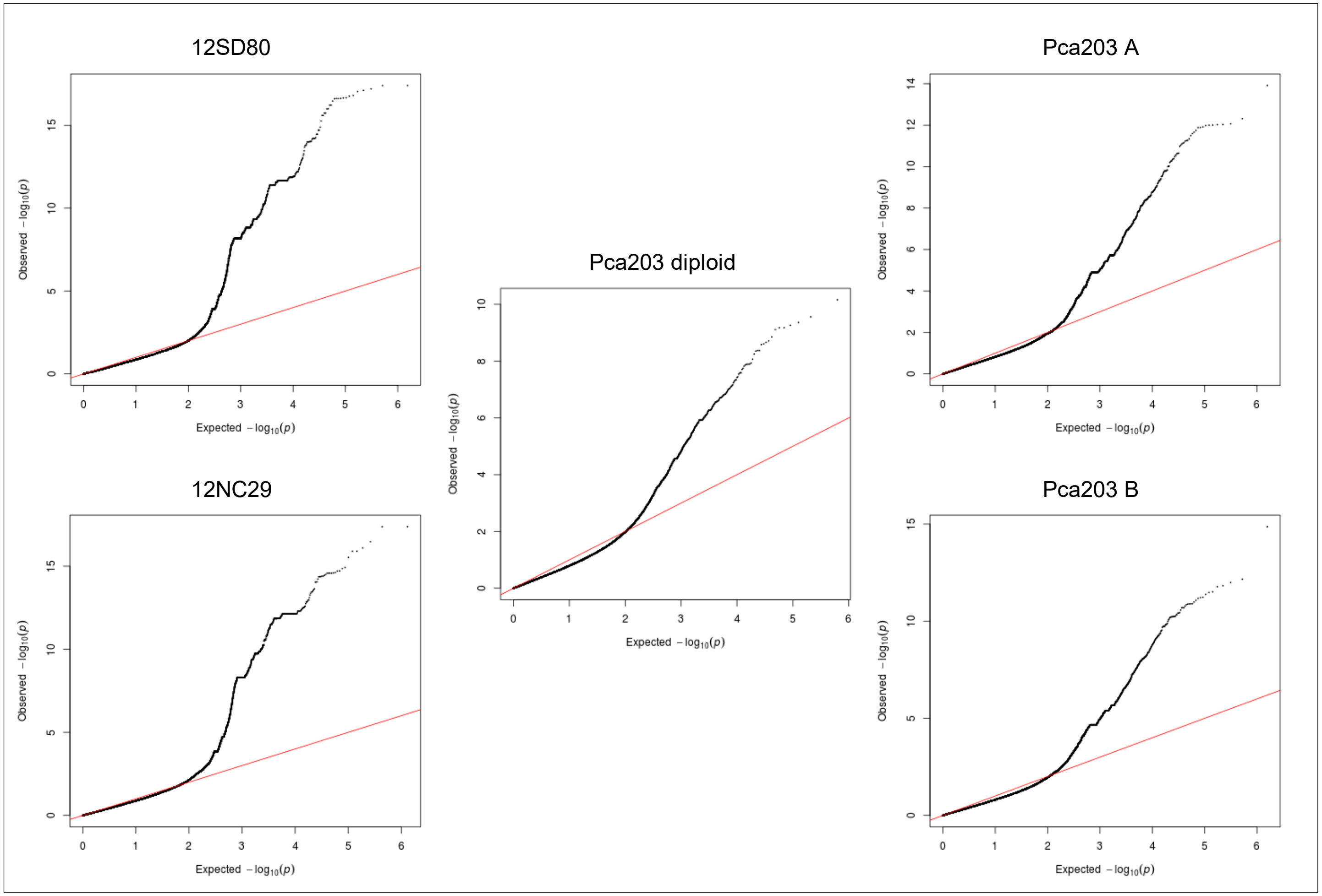

**Fig 35B. *Pca* isolate virulence score distribution and Manhattan plots showing SNP association values for virulence to oat line *Pc91*.** Red and blue horizontal lines denote Bonferroni significance threshold (α = 0.01/total number of markers) and 5% false discovery rate threshold, respectively. Significant association peaks are labelled with contig or chromosome number and assigned VGI number.

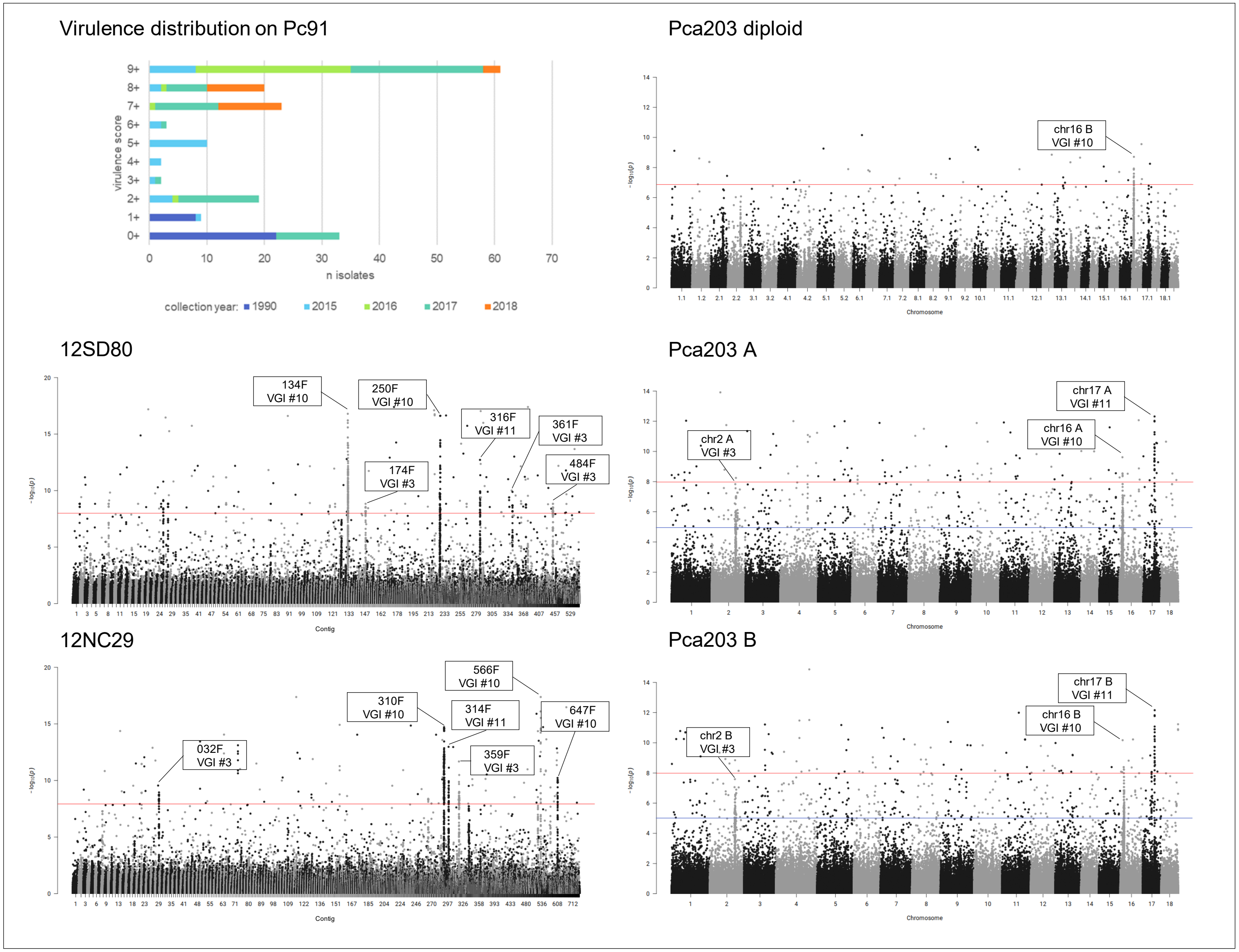

**Fig 36A. quantile-quantile plots of SNP association values for virulence of *Pca* to oat line *Pc94*.** Red lines indicate expected P-value distributions if P-values follow a uniform distribution (null hypothesis).

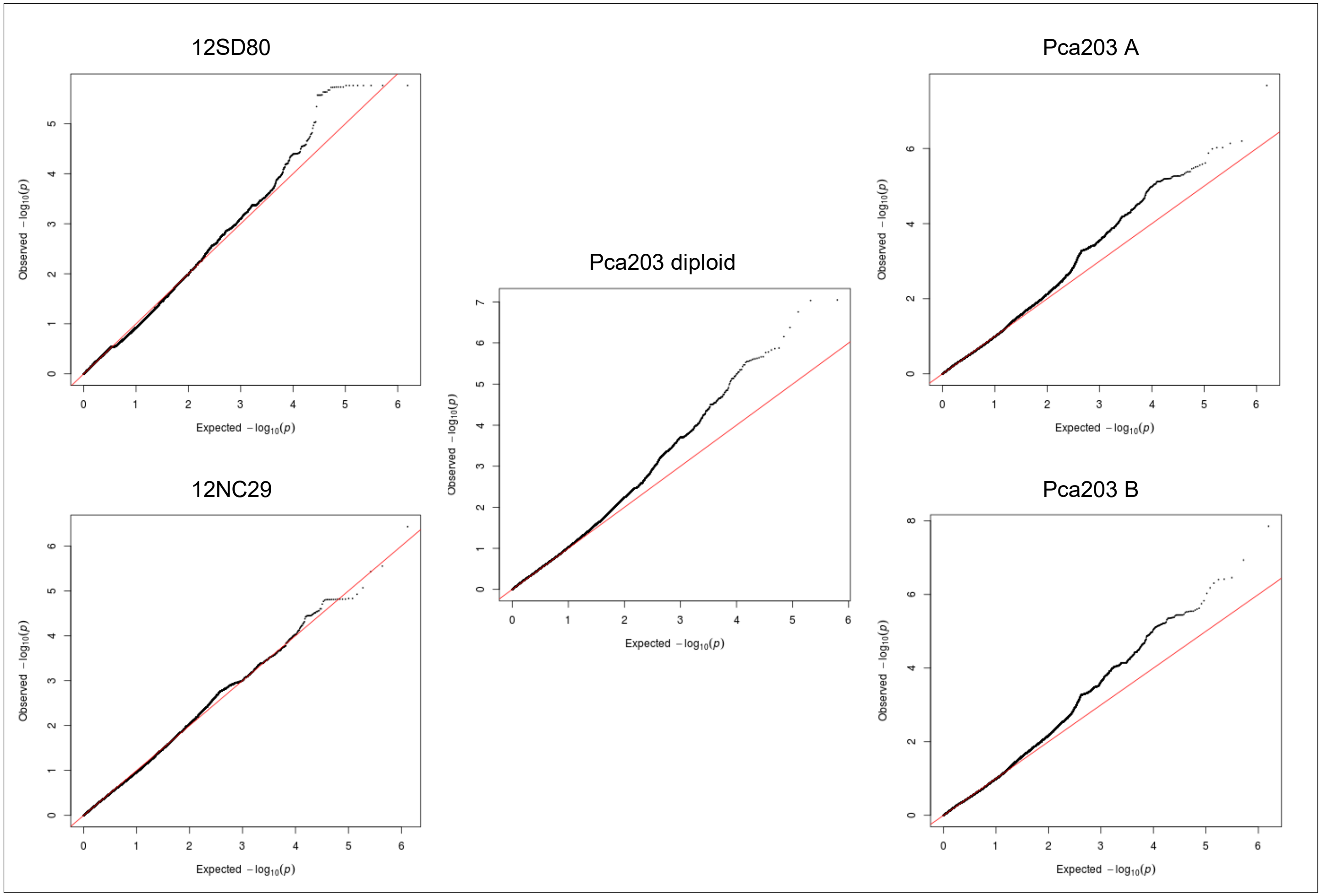

**Fig 36B. *Pca* isolate virulence score distribution and Manhattan plots showing SNP association values for virulence to oat line *Pc94*.** Red horizontal lines denote Bonferroni significance threshold (α = 0.01/total number of markers).

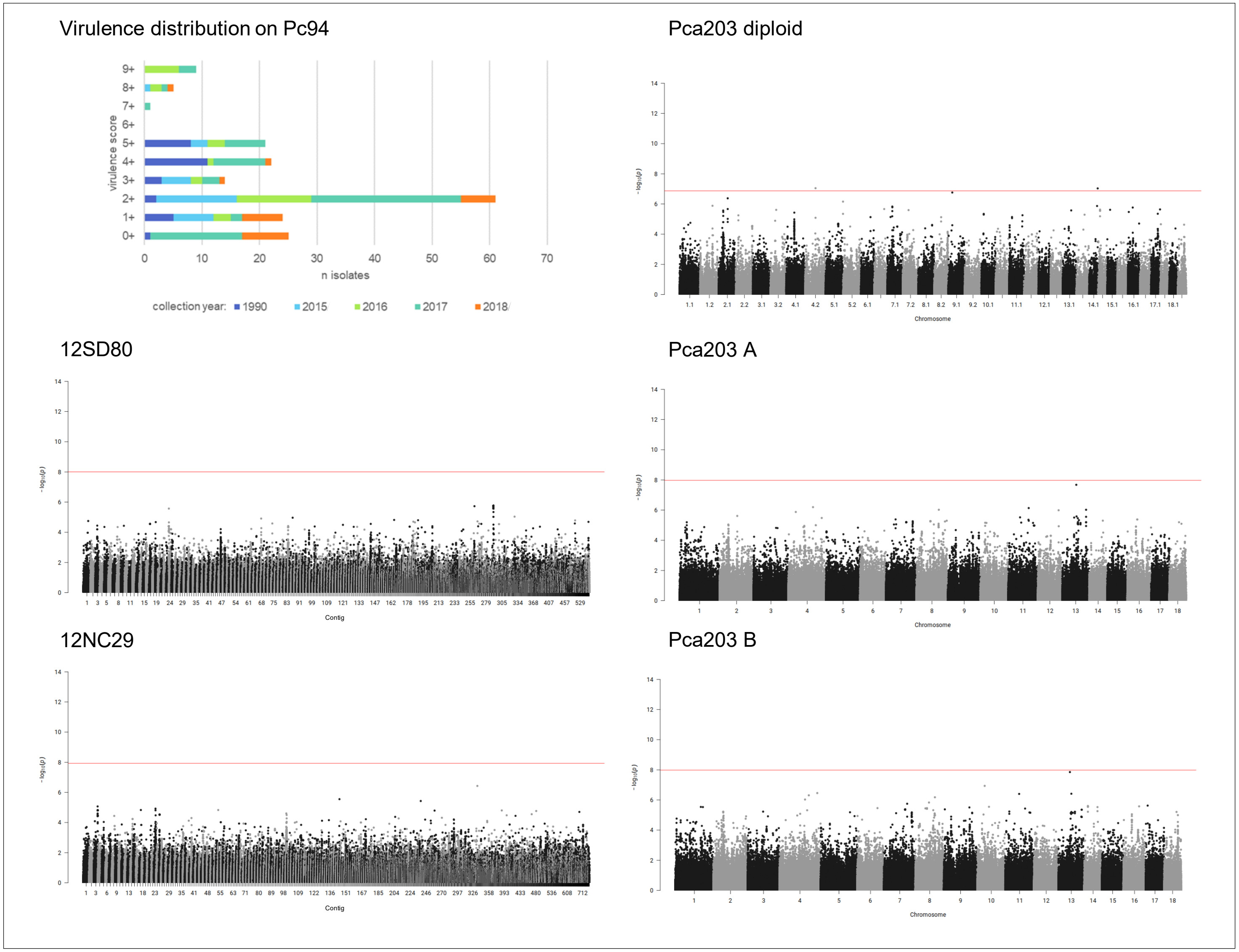

**Fig 37A. quantile-quantile plots of SNP association values for virulence of *Pca* to oat line *Pc96*.** Red lines indicate expected P-value distributions if P-values follow a uniform distribution (null hypothesis).

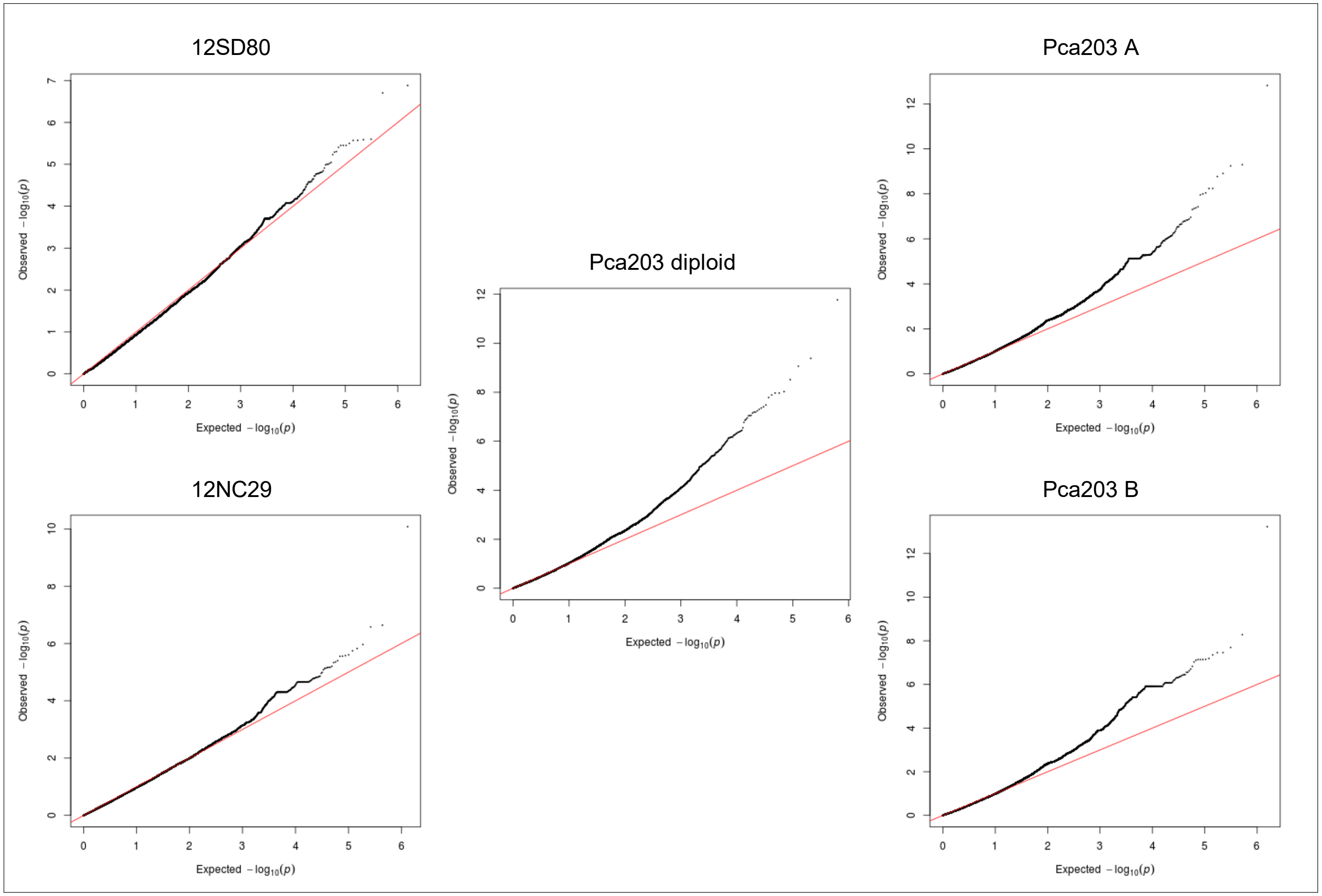

**Fig 37B. *Pca* isolate virulence score distribution and Manhattan plots showing SNP association values for virulence to oat line *Pc96*.** Red and blue horizontal lines denote Bonferroni significance threshold (α = 0.01/total number of markers) and 5% false discovery rate threshold, respectively. Potentially significant association peaks are labelled with contig or chromosome number.

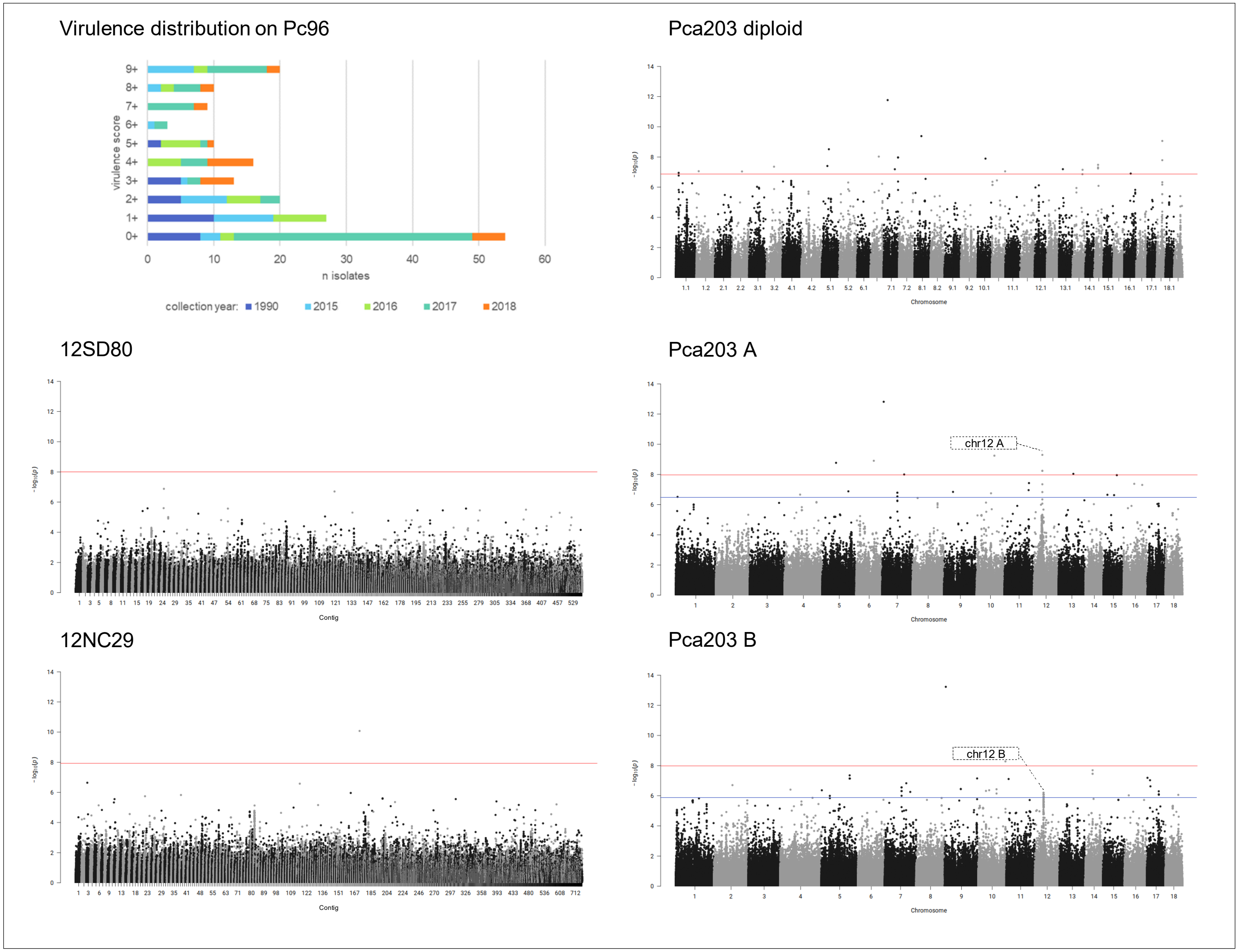

**Fig 38A. quantile-quantile plots of SNP association values for virulence of *Pca* to oat line Stainless.** Red lines indicate expected P-value distributions if P-values follow a uniform distribution (null hypothesis).

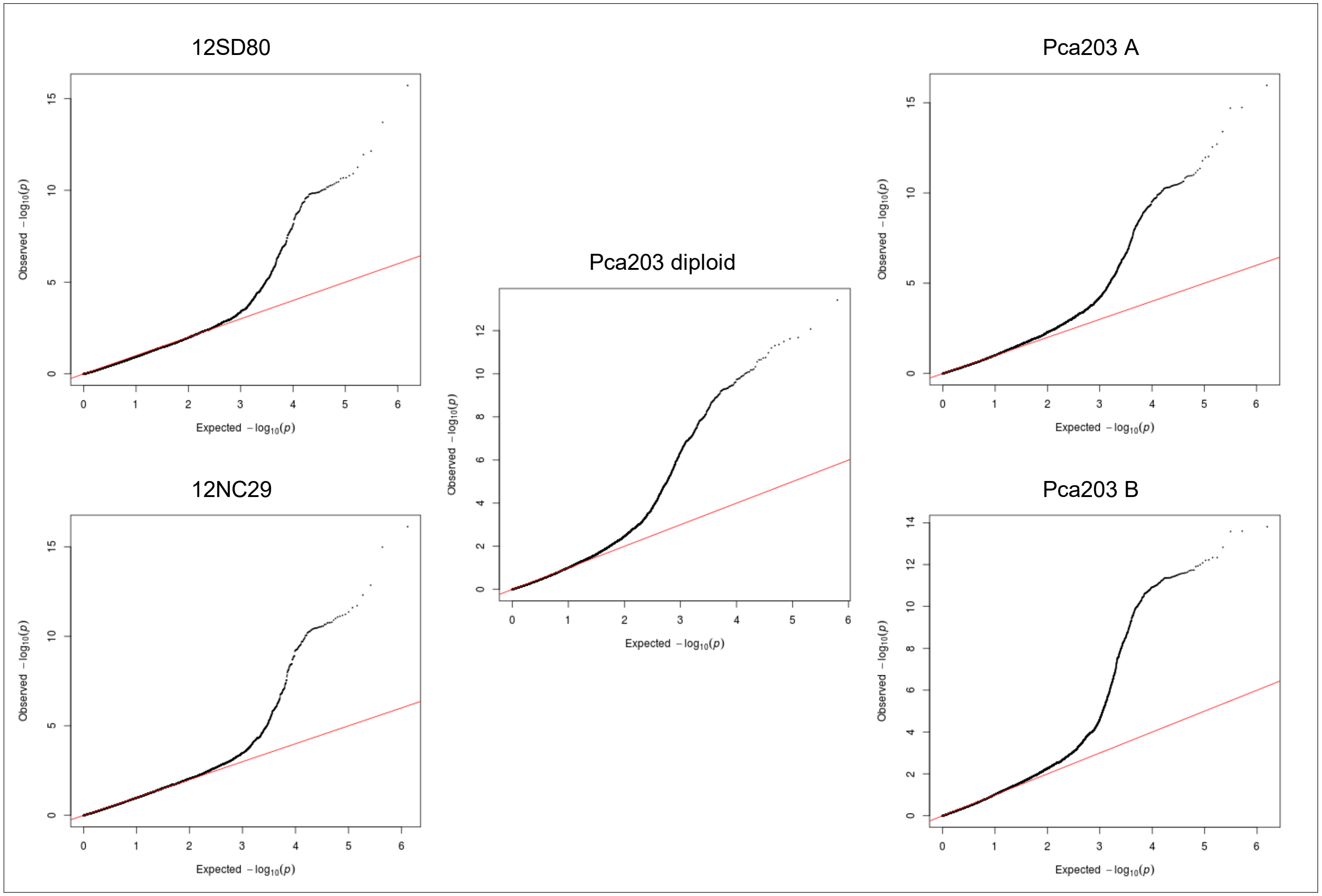

**Fig 38B. *Pca* isolate virulence score distribution and Manhattan plots showing SNP association values for virulence to oat line Stainless.** Red horizontal lines denote Bonferroni significance threshold (α = 0.01/total number of markers).

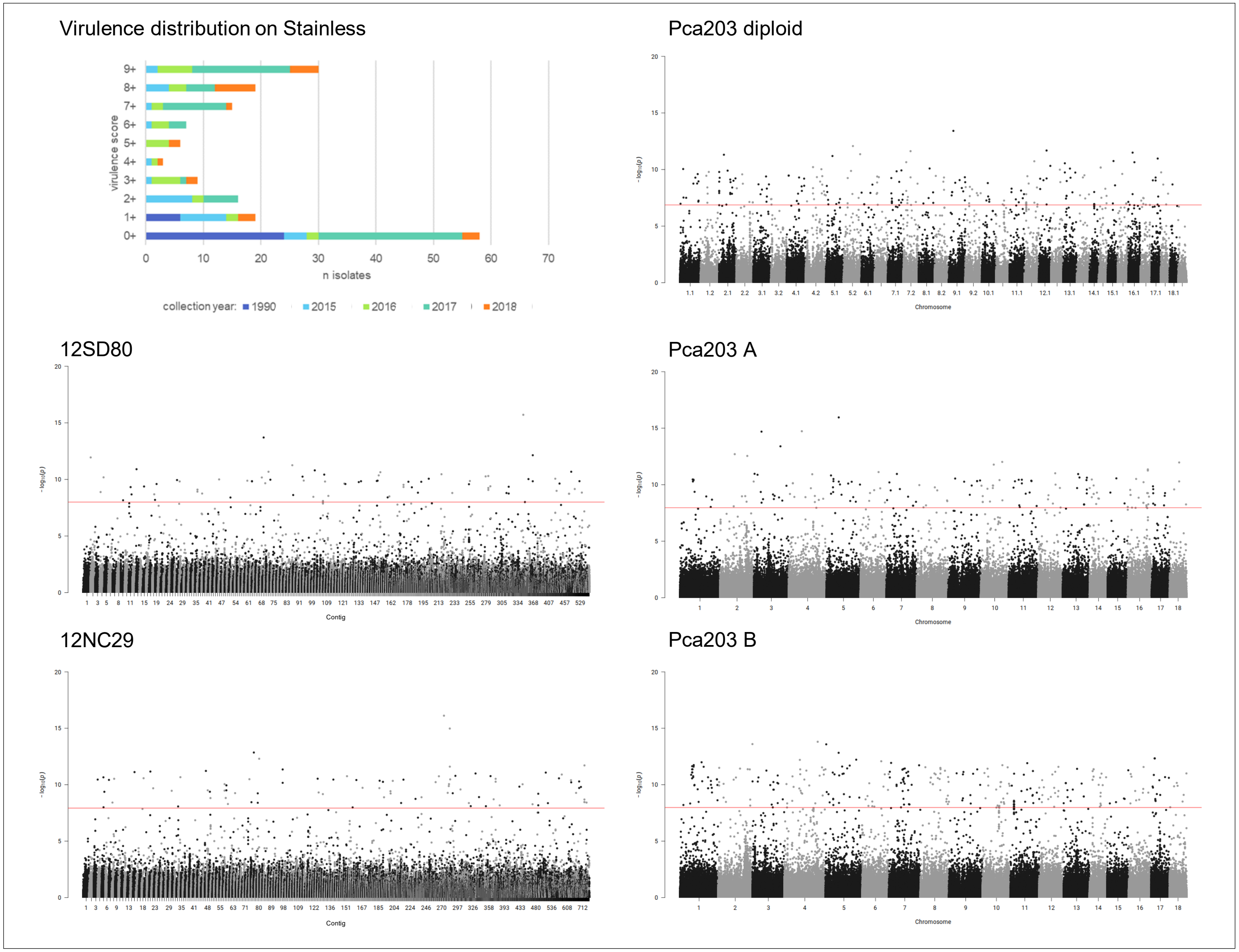

**Fig 39A. quantile-quantile plots of SNP association values for virulence of *Pca* to oat line TAM-O-405.** Red lines indicate expected P-value distributions if P-values follow a uniform distribution (null hypothesis).

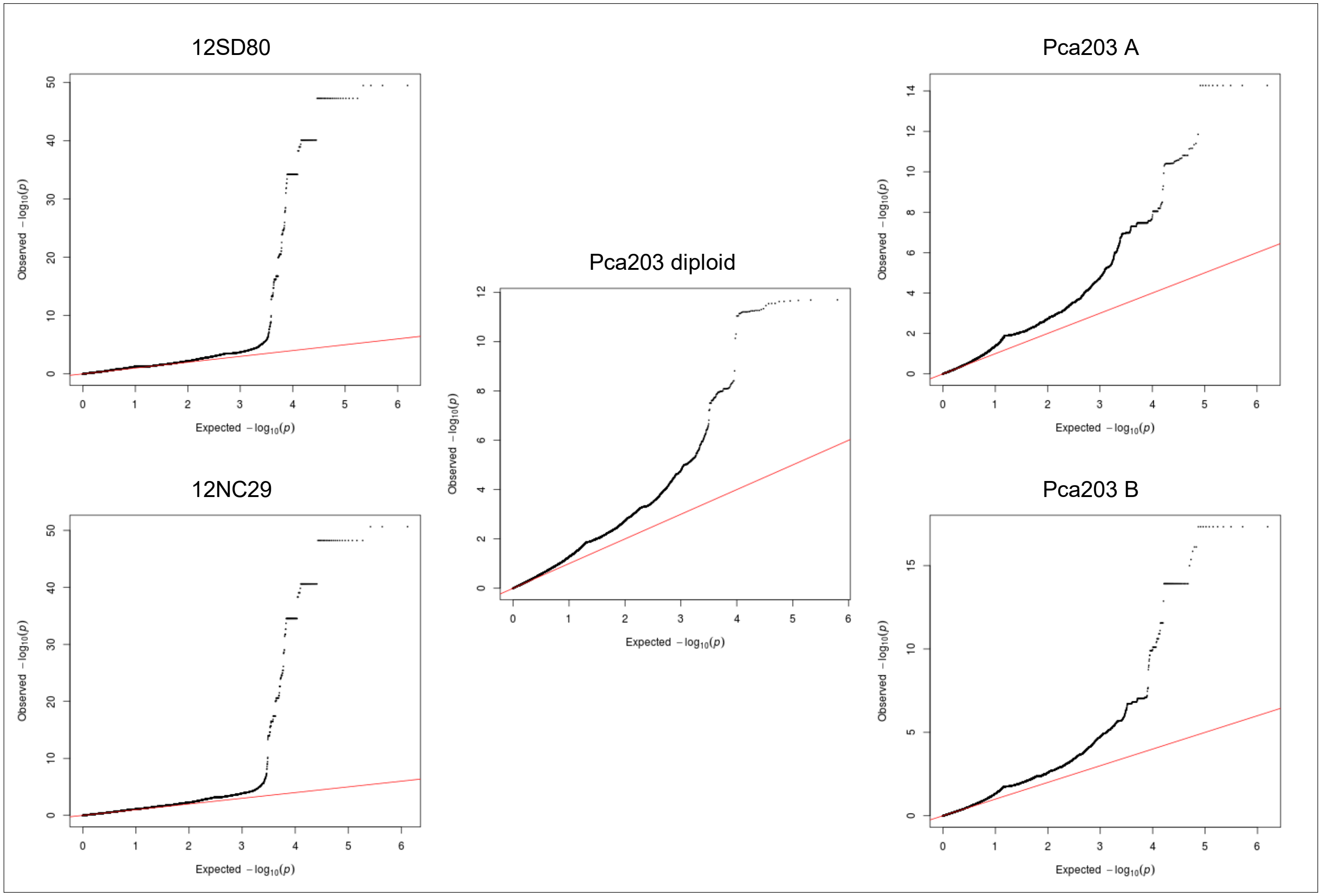

**Fig 39B. *Pca* isolate virulence score distribution and Manhattan plots showing SNP association values for virulence to oat line TAM-O-405.** Red horizontal lines denote Bonferroni significance threshold (α = 0.01/total number of markers). Significant association peaks are labelled with contig or chromosome number and assigned VGI number.

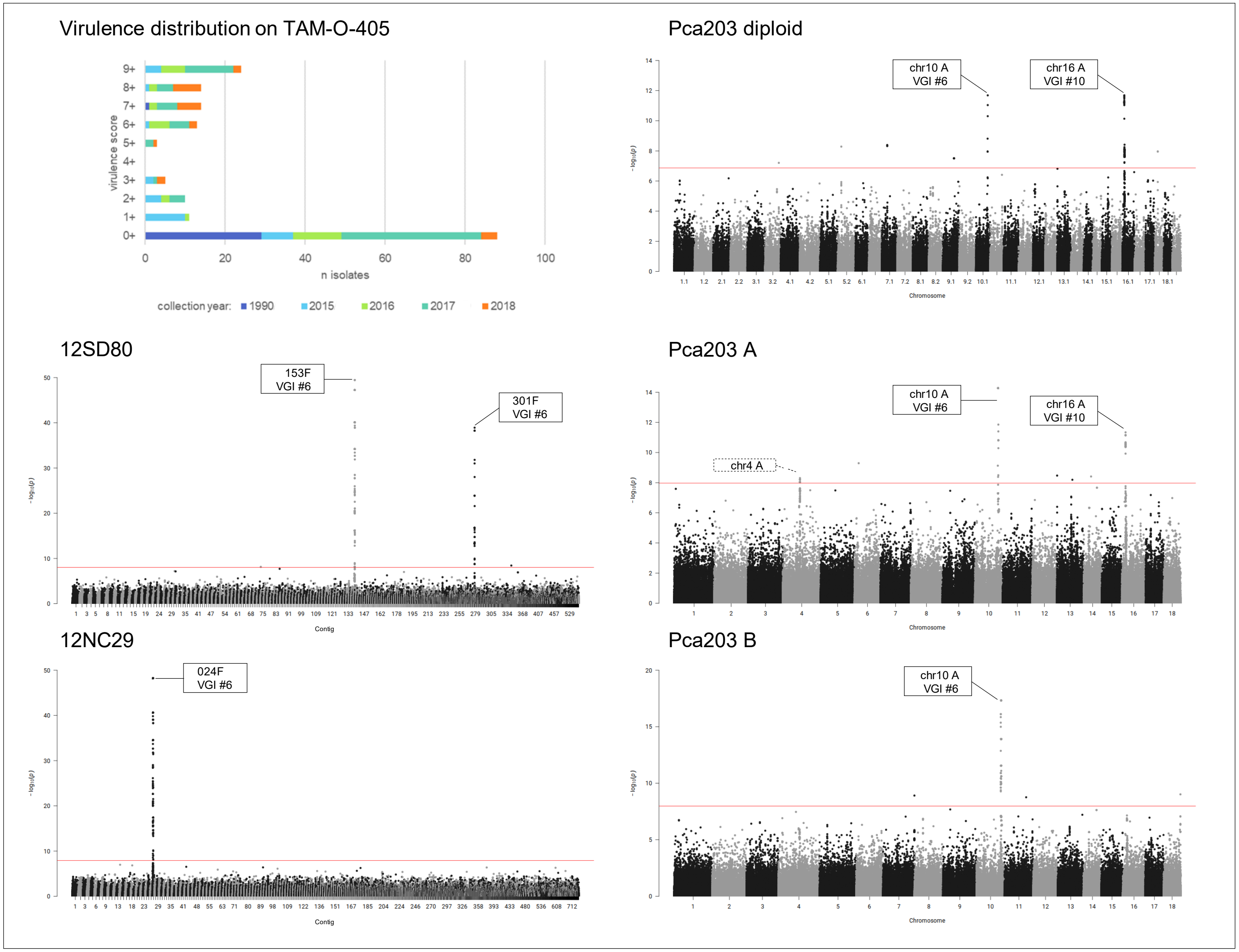

**Fig 40A. quantile-quantile plots of SNP association values for virulence of *Pca* to oat line WIX4361-9.** Red lines indicate expected P-value distributions if P-values follow a uniform distribution (null hypothesis).

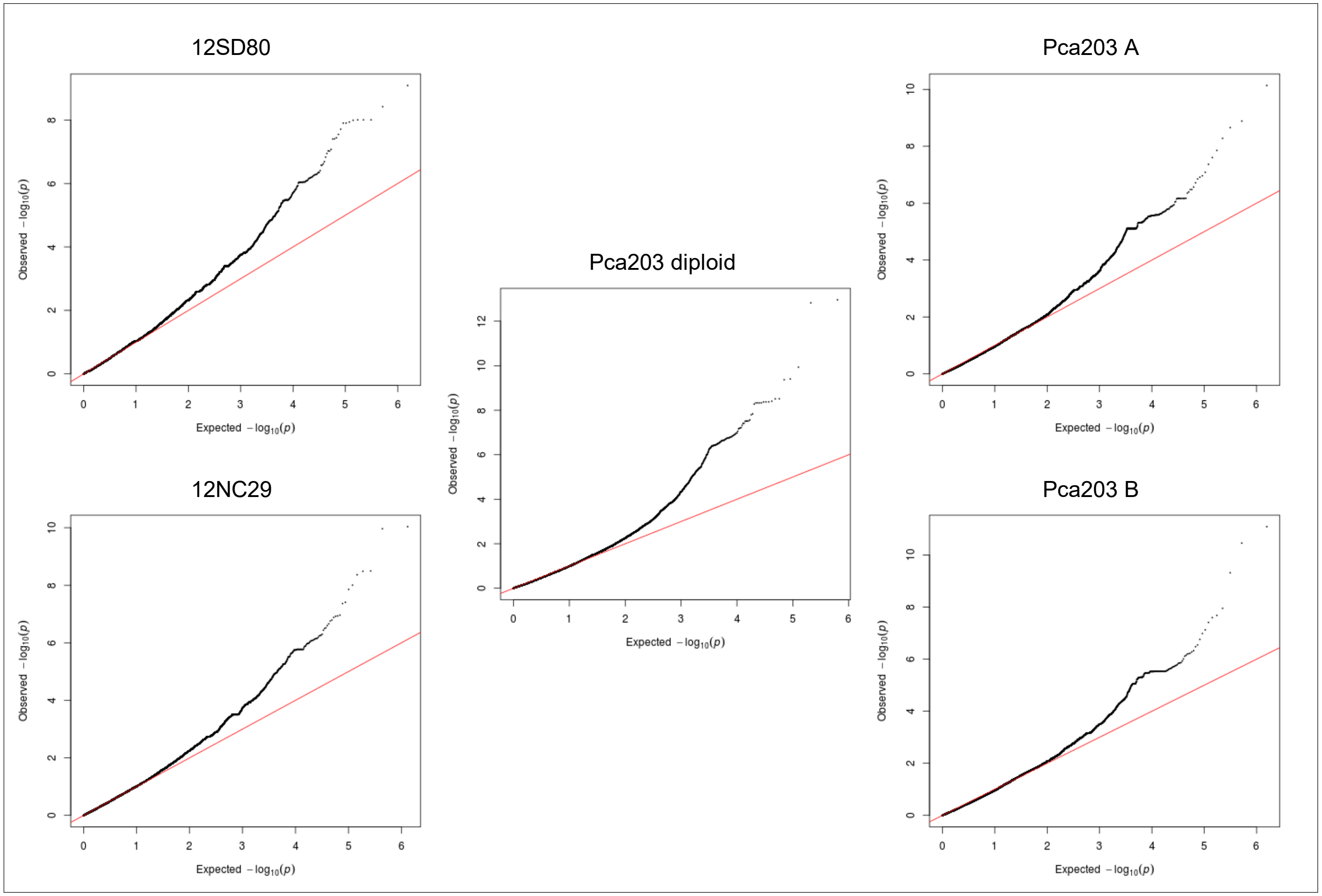

**Fig 40B. *Pca* isolate virulence score distribution and Manhattan plots showing SNP association values for virulence to oat line WIX4361-9.** Red and blue horizontal lines denote Bonferroni significance threshold (α = 0.01/total number of markers) and 5% false discovery rate threshold, respectively. Significant association peaks are labelled with contig or chromosome number and assigned VGI number.

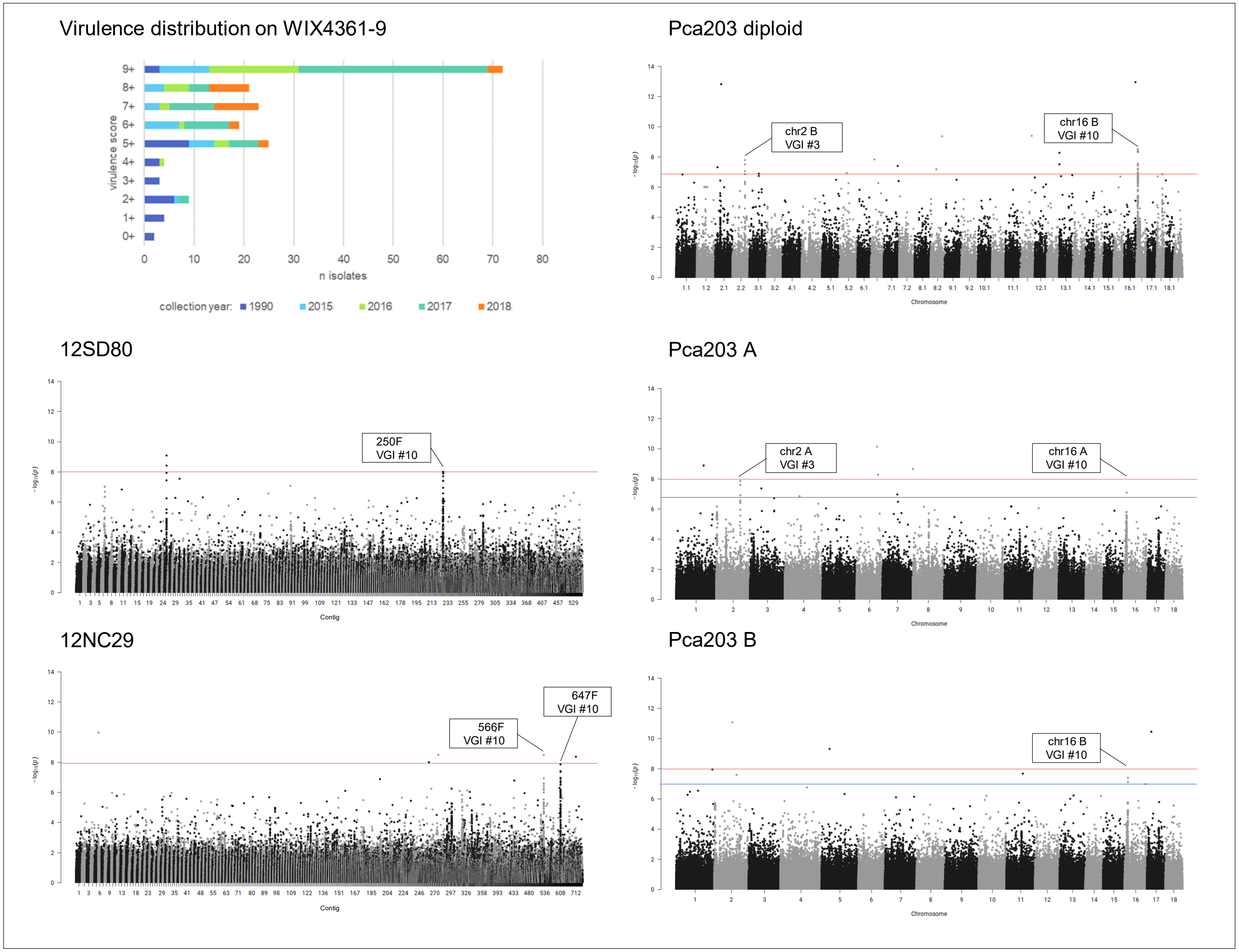
